# *LMNA*-Related Dilated Cardiomyopathy: Single-Cell Transcriptomics during Patient-derived iPSC Differentiation Support Cell type and Lineage-specific Dysregulation of Gene Expression and Development for Cardiomyocytes and Epicardium-Derived Cells with Lamin A/C Haploinsufficiency

**DOI:** 10.1101/2024.06.12.598335

**Authors:** Michael V. Zaragoza, Thuy-Anh Bui, Halida P. Widyastuti, Mehrsa Mehrabi, Zixuan Cang, Yutong Sha, Anna Grosberg, Qing Nie

## Abstract

*LMNA*-Related Dilated Cardiomyopathy (DCM) is an autosomal-dominant genetic condition with cardiomyocyte and conduction system dysfunction often resulting in heart failure or sudden death. The condition is caused by mutation in the Lamin A/C (*LMNA*) gene encoding Type-A nuclear lamin proteins involved in nuclear integrity, epigenetic regulation of gene expression, and differentiation. Molecular mechanisms of disease are not completely understood, and there are no definitive treatments to reverse progression or prevent mortality. We investigated possible mechanisms of *LMNA*-Related DCM using induced pluripotent stem cells derived from a family with a heterozygous *LMNA c.357-2A>G* splice-site mutation. We differentiated one *LMNA* mutant iPSC line derived from an affected female (Patient) and two non-mutant iPSC lines derived from her unaffected sister (Control) and conducted single-cell RNA sequencing for 12 samples (4 Patient and 8 Control) across seven time points: Day 0, 2, 4, 9, 16, 19, and 30. Our bioinformatics workflow identified 125,554 cells in raw data and 110,521 (88%) high-quality cells in sequentially processed data. Unsupervised clustering, cell annotation, and trajectory inference found complex heterogeneity: ten main cell types; many possible subtypes; and lineage bifurcation for Cardiac Progenitors to Cardiomyocytes (CM) and Epicardium-Derived Cells (EPDC). Data integration and comparative analyses of Patient and Control cells found cell type and lineage differentially expressed genes (DEG) with enrichment to support pathway dysregulation. Top DEG and enriched pathways included: 10 *ZNF* genes and RNA polymerase II transcription in Pluripotent cells (PP); *BMP4* and TGF Beta/BMP signaling, sarcomere gene subsets and cardiogenesis, *CDH2* and EMT in CM; *LMNA* and epigenetic regulation and *DDIT4* and mTORC1 signaling in EPDC. Top DEG also included: *XIST* and other X-linked genes, six imprinted genes: *SNRPN*, *PWAR6*, *NDN*, *PEG10*, *MEG3*, *MEG8*, and enriched gene sets in metabolism, proliferation, and homeostasis. We confirmed Lamin A/C haploinsufficiency by allelic expression and Western blot. Our complex Patient-derived iPSC model for Lamin A/C haploinsufficiency in PP, CM, and EPDC provided support for dysregulation of genes and pathways, many previously associated with Lamin A/C defects, such as epigenetic gene expression, signaling, and differentiation. Our findings support disruption of epigenomic developmental programs as proposed in other *LMNA* disease models. We recognized other factors influencing epigenetics and differentiation; thus, our approach needs improvement to further investigate this mechanism in an iPSC-derived model.

## INTRODUCTION

Lamins are intermediate filament proteins of the nuclear lamina that interacts with chromatin and the inner nuclear membrane and are distributed throughout the nucleoplasm [1, 2]. Lamins have diverse roles in both nuclear structure and function including gene expression, chromatin regulation, cell proliferation, and differentiation [1, 2]. Epigenomic roles involve heterochromatin binding and genome organization through Lamina-Associated Domains (LAD) [3, 4]. In humans, nuclear lamins are A-type lamins: lamin A and C isoforms encoded by the *LMNA* gene and B-type lamins: Lamin B1 encoded by *LMNB1* and Lamin B2 and B3 encoded by *LMNB2* [1, 2]. B-type lamins are constitutively expressed, and A-type lamins are expressed temporally during development in most differentiated cells [5, 6].

Despite *LMNA* expression in most differentiated cells, *LMNA* mutations primarily affect mesoderm-derived lineages in diseases collectively called laminopathies [1, 7]. *LMNA*-Related Dilated Cardiomyopathy (DCM), the heart-specific laminopathy, is an autosomal-dominant condition with left ventricular enlargement, systolic dysfunction, and conduction system disease often resulting in heart failure or sudden death [8]. Despite clinical severity, molecular mechanisms of *LMNA*-Related DCM are not completely understood, and there are no definitive treatments to reverse progression or prevent mortality.

For heart and striated muscle laminopathies, two interconnected mechanisms are proposed to explain cell type-specific effects of lamin mutations: the ‘mechanical stress’ hypothesis, based on nuclear structure, and ‘gene expression’ hypothesis, based on nuclear function [1, 7, 9]. In the ‘mechanical stress’ hypothesis, *LMNA* mutations alter lamina structure leading to mechanical fragility of nuclei and cell dysfunction. In the ‘gene expression’ hypothesis, *LMNA* mutations dysregulate nuclear chromatin organization and gene transcription to alter cell function [1, 7, 9]. The ‘gene expression’ hypothesis involves epigenomic defects, affecting normal cell differentiation, in which *LMNA* mutations uncouple LAD from peripheral heterochromatin to alter genome organization and unlock aberrant genes and pathways [10–12].

For *LMNA*-Related DCM, the ‘gene expression’ hypothesis is supported in comparisons of *LMNA* mutant and control cells from three main types of experimental models: mouse models, patient heart tissue, and induced pluripotent stem cell (iPSC) models [1, 7, 9, 11, 12]. From *Lmna* mutant mouse models, evidence supports cell signaling dysregulation including hyperactivation of TGF Beta [13, 14] and AKT-mTORC1 signaling pathways [15] and dysregulated differentiation pathways including Epithelial-Mesenchymal Transition (EMT) [16]. From *LMNA* patient heart tissue, evidence also supports dysregulation of signal transduction and gene expression including TGF Beta/BMP signaling pathway and *BMP4* overexpression [17, 18].

While evidence from numerous mouse models and limited patient heart tissue supports the ‘gene expression’ hypothesis for *LMNA*-Related DCM, another valuable approach involves Patient-derived iPSC models from reprogramming somatic cells of patients into iPSC [19] that can be directly differentiated *in vitro* into cardiomyocytes (iPSC-CM) for analyses [20]. From *LMNA* Patient-derived iPSC, evidence supports dysregulation of PDGF signaling associated with open chromatin [21], non-cardiac lineage expression with chromatin compartment changes [22], and non-myocyte lineage expression with LAD changes [23]. Furthermore, analysis of *LMNA* knockdown iPSC-CM during differentiation found premature increase in cardiogenesis genes [24]. These studies [13–18, 21–24] do provide much support for the ‘gene expression’ hypothesis; however, the results obtained using traditional techniques (e.g., bulk RNA-sequencing) may be inconsistent. For samples with ‘previously unappreciated levels of heterogeneity,’ such techniques, that average measurements across all cells in a sample, may obscure cell specific findings [25].

More recently, measurements at single-cell resolution are used in cardiovascular studies to evaluate cell heterogeneity of cell samples and heart tissues and to dissect cell type-specific mechanisms in normal and disease processes [26]. To study normal processes, many recent studies used single-cell RNA-sequencing (scRNA-seq) to evaluate normal gene expression and developmental pathways in human iPSC [27, 28] and during differentiation to cardiomyocytes [29–36]. In contrast, single-cell analyses to study disease processes as *LMNA-*Related DCM are limited and include our initial study of Patient iPSC-CM [37], a large DCM single nuclei RNA-seq study with heart tissue from 12 *LMNA* patients [38], and scRNA-seq studies in *Lmna* Q353R mutant mouse model and heart tissue from one *LMNA* Q353R patient [39].

To evaluate cell heterogeneity and cell type and lineage-specific mechanisms for *LMNA*-Related DCM, we analyzed a Patient-derived iPSC disease model using scRNA-seq at multiple time points during CM differentiation. This extends our studies focused on a unique family with DCM with a heterozygous *LMNA* c.357-2A>G splice-site mutation [37, 40, 41] by testing the ‘gene expression’ hypothesis that Lamin A/C haploinsufficiency may be associated with dysregulation of epigenomic developmental pathways and expression of genes with important roles in the nuclear function. Here, we describe our scRNA-seq bioinformatic workflow with serial, cell type-specific data processing, report our results from extensive multi-level analyses in differentiation of Control iPSC and *LMNA* Patient iPSC, and discuss our evidence to support the ‘gene expression’ hypothesis along with the challenges of *LMNA* disease modeling using Patient-derived iPSC.

## MATERIALS AND METHODS

### Generation and validation of Control and Patient iPSC lines

Using standard methods, we generated and validated eight iPSC lines (**Table S1**) each derived from dermal fibroblasts as reported [41]. Of eight iPSC lines, we generated seven iPSC lines from the study family [40]: three *LMNA* mutant iPSC lines from three affected members heterozygous for the *LMNA* c.357-2A>G splice-site mutation (Patient) and four non-mutant iPSC lines from three unaffected, *LMNA* mutation-negative members to serve as sex and age-matched controls (Control). For the first Control (CA1), we generated two iPSC lines, CA1-A [41] and CA1-B [37] from independent clones (A and B) as reported. For Control A2, revival of cryopreserved cells had low viability and culturing failed to reach the optimal confluency for differentiation; therefore, we generated Unrelated Control 2 iPSC using purchased human dermal fibroblasts from a healthy unrelated male (CC-2511, Lot No. 0000293971, Lonza, Basel, Switzerland). We conducted fibroblast collection and culturing, DNA extraction, *LMNA* genotyping, and Whole Exome Sequencing (WES) as previously described [40, 42].

To validate each iPSC line, we evaluated independent clones for absence of chromosome abnormalities by karyotype analysis and for normal pluripotency by immunocytochemistry (ICC) staining as reported (**Table S1**) [41]. Validation also included differentiation capability by ICC staining for three germ layer markers in embryoid bodies (EB). For each clone, we harvested iPSC at 70-80% confluency using 0.5 mM EDTA (Thermo Fisher Scientific (TFS), Waltham, MA) and transferred iPSC clumps to non-treated 35-mm culture dishes with Essential 6 Medium (TFS) and 5 µM ROCK inhibitor Y-27632 (Cellagen Technology, San Diego, CA). After four to seven days, we transferred unattached EB to Matrigel-coated chamber slides for spontaneous differentiation. After 14 to 28 days of EB differentiation, we stained cells for markers of endoderm (alpha-fetoprotein or Forkhead Box A2), mesoderm (smooth muscle actin), and ectoderm (beta-III tubulin) using the 3-Germ Layer Immunocytochemistry Kit (A25538, TFS) (**Table S2**).

### *In vitro* iPSC-CM differentiation and cell collection

Using two standard protocols (A and B) (**Fig. S1**), we differentiated three iPSC lines (CA1-A, CA1-B, PA1) for serial scRNA-seq and six iPSC lines (CA1-B, U2, CA3, PA1, PA2, PA3) for Western Blot. Protocol-A used Cardiomyocyte (CM) Differentiation Methods (Version 1.0) from the Allen Institute of Cell Science [43] based on modulation of Wnt/beta-catenin signaling using GSK3 inhibitor CHIR99201 and Wnt inhibitor IWP2 (GiWi protocol) [44]. Protocol-B used Gibco PSC CM Differentiation Kit (A2921201, TFS) as described [37]. To generate iPSC-CM, we used both protocols for single-cell RNA-sequencing (scRNA-seq) and only Protocol-A for Lamin A/C Western Blots (WB).

Although we used standard protocols to generated iPSC-CM (**Fig. S1**), differences between Protocol-A and Protocol-B [37] include iPSC growth substrates and media. Protocol-B [37] involved seeding iPSC on Geltrex (TFS) coated 12-well plates in Essential 8 media (TFS). In contrast, for Protocol-A, we seeded thawed iPSC at passage 12 (P12) or above in Matrigel-coated 6-well plates in mTESR1 media (STEMCELL Technologies (SCT), Vancouver, Canada) and 10 µM ROCK inhibitor. After 24 hours, we replaced media with mTESR1 media and no ROCK inhibitor with daily changes. At 85-90% confluency, we passaged iPSC using ReLeSR (SCT). After at least two passages, we singularized iPSC with TrypLE Select Enzyme (TFS) and seeded cells to Matrigel-coated 12-well or 24-well plates in mTESR1 media and 10 µM ROCK inhibitor.

For CM differentiation, Protocol-A used the GiWi protocol [43, 44]. To determine optimal cell density, we seeded each iPSC clone at different concentrations three to four days prior to differentiation (D-3 or D-4) for growth to approximately 90-95% confluency. On Day 0 of differentiation (D00), we changed mTESR1 media to CM differentiation medium consisting of RPMI 1640 media (TFS) with B27 supplement without insulin (RPMI/B27-) (TFS) and with 7.5 µM CHIR99201 (Cayman Chemical Company, Ann Arbor, MI). After 48 hours (D02), we replaced media with RPMI/B27-with 7.5 µM IWP2 (R&D Systems, Minneapolis, MN). After 48 hours (D04), we replaced media with RPMI/B27-. On Day 6 of differentiation (D06), we replaced media with RPMI 1640 media supplemented with B27 supplement with insulin (RPMI/B27+) (TFS). The protocol produced beating clusters on D08 to D10.

To collect iPSC-CM at selected time points, we continued culturing in RPMI/B27+ media with changes every other day and used the STEMdiff CM Dissociation Kit (SCT) for harvesting. After washing wells twice with DPBS, we harvested cells by incubation in pre-warmed Dissociation Medium for 10 to 12 minutes at 37C, resuspended dissociated cells in Support Medium by gentle trituration with a 10 mL pipette, transferred the cell suspension to a conical tube with Support Medium, and centrifuged at 300 x *g* for 5 minutes. Finally, to check concentration and viability, we analyzed resuspended cells using trypan blue staining and Countess II Automated Cell Counter (TFS).

### Serial scRNA-seq studies using 10x Genomics platform

To study possible mechanisms of the *LMNA* mutation, we conducted scRNA-seq studies using two Control iPSC lines (two clones: CA1-A and CA1-B), derived from dermal fibroblasts of the same Control individual CA1, and one Patient iPSC line (one clone: PA1) (**Graphical Abstract**). For scRNA-seq studies, we differentiated three iPSC lines and collected cell samples at seven time points (Day 0, 2, 4, 9, 16, 19, and 30). We collected a total of 12 cell samples, 8 Control and 4 Patient cell samples, consisting of two sample sets, Set-A and Set-B derived from two CM differentiation protocols (**Fig. S1**), Protocol-A (Zaragoza Lab) and Protocol-B (Grosberg Lab), respectively. Set-A consisted of four unpaired Control samples derived from iPSC line CA1-A collected at Day 2, 4, 9, and 30. Set-B consisted of four paired samples derived from iPSC lines CA1-B (Control) and PA1 (Patient) differentiated in parallel and collected at Day 0, 9, 16, and 19.

For Sample Set-A (n=4 samples, CA1-A unpaired: D02, D04, D09A, D30), we harvested cells from two wells of one 12-well plate using TrypLE Select Enzyme and filtered using a 35 µM cell strainer to create single-cell suspensions. We counted cells and checked for viability, washed cells twice using DPBS with 0.04% bovine serum albumin, and recounted cells for the final cell count. We resuspended cells in Eppendorf DNA LoBind tubes (TFS) to a target cell concentration of 7 x 10(5) to 1 x 10(6) cells/mL and cell viability of at least 80%.

For Sample Set-B (n=8 samples, CA1-B and PA1 pairs: D00, D09B, D16, D19), we first harvested iPSC (P15) at ∼70-80% confluency from three 12-well plates (36 total wells) and pooled cells for each sample (CA1-B and PA1: D00). We also reseeded iPSC (P16) for CM differentiation. At Day 9, 16, and 19, we harvested cells from six 12-well plates (72 total wells) and pooled cells for each sample (CA1-B: D09B, D16, D19 and PA1: D09B, D16, D19). For Day 16 and 19, we collected cells after metabolic selection by glucose deprivation and lactate supplementation [45, 46] for four days (D10 to D14) using the Gibco enrichment protocol (MAN0014828, TFS).

For scRNA-seq, we transferred cell samples on ice immediately after collection to the UCI Genomics High-Throughput Facility. Sample processing used the droplet-based Chromium system [47] and Chromium Single Cell 3’ Gene Expression reagent kits v3 (10X Genomics, Pleasanton, CA). The Facility quantified single-cell cDNA libraries and multiplexed libraries by Illumina paired-end sequencing for 28 cycles for Read 1 (cell barcode + UMI tag), 8 cycles for sample index, and 100 cycles for Read 2 (cDNA insert). Library sequencing involved first, shallow sequencing using HiSeq 4000 with the goal of obtaining 3,000 to 9,000 raw reads per cell to estimate the number of cells captured for each sample, and then, deeper sequencing using NovaSeq 6000 with the goal of obtaining at least 50,000 raw reads per cell. Using Illumina bcl2fastq2 conversion software, the Facility processed raw data files (BCL) for index reads, demultiplexed data into sample specific FASTQ files for Read 1, Read 2, and sample index, and then stored FASTQ sequence files on the UCI High Performance Computing Cluster.

### scRNA-seq Bioinformatics Workflow

Our workflow (**Fig. 1**) used standard pipelines [48–50] and bioinformatics software packages (**Table S3**) involving three steps: Step-I. Data Processing: to obtain high-quality cells; Step-II. Data Analysis-Cluster, Annotate, and Subset: to identify main cell types and possible subtypes; Step-III. Data Combining and Comparative Analyses: to align cells from different samples for cell type and lineage differential gene expression and pathway enrichment. These three steps used two general types of data: 1. Single Sample Data: Individual data for each of the 12 samples; 2. Combined Data: Merged (Non-Integrated) and Integrated Data from combining multiple sets or subsets of data.

**Figure 1.**
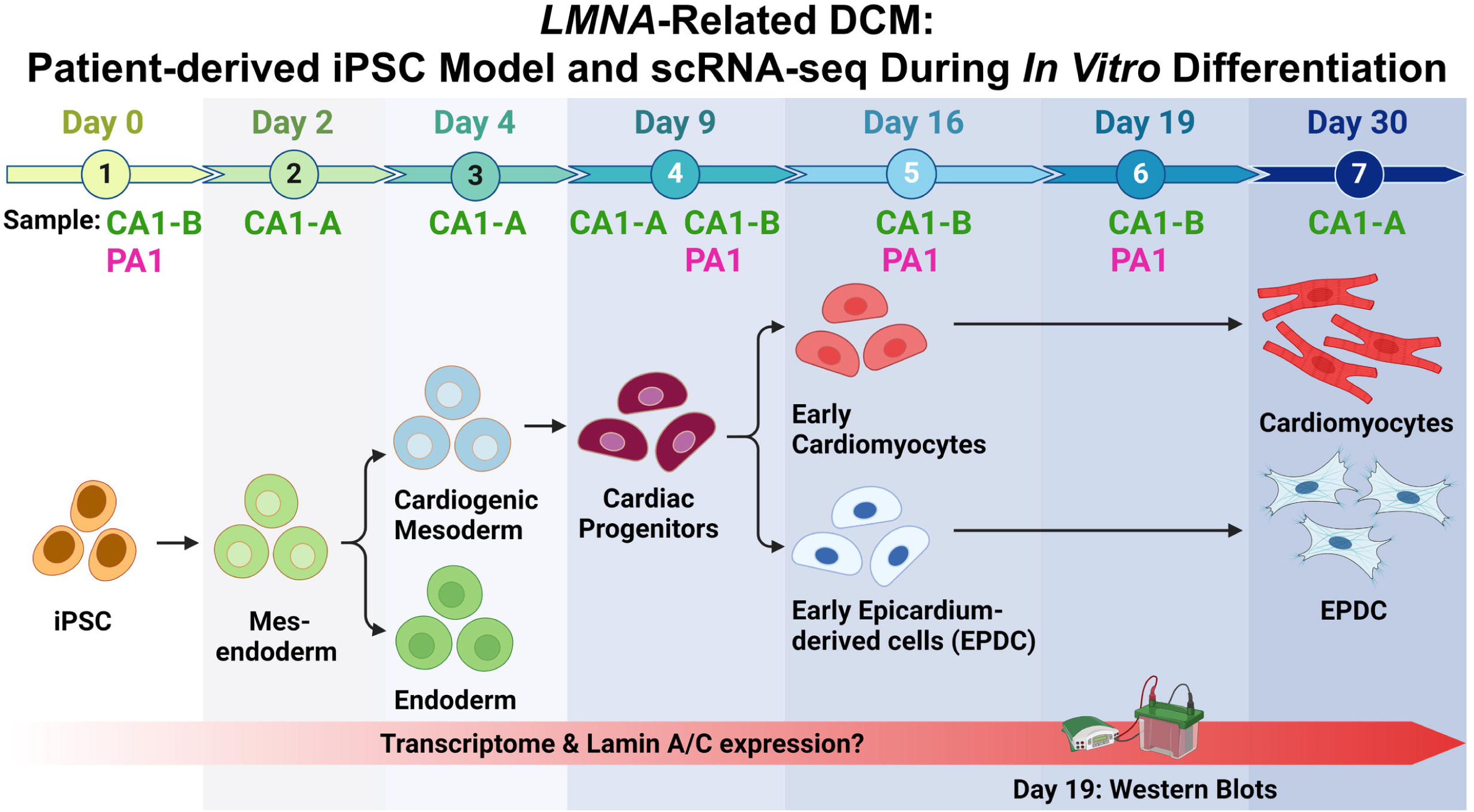
scRNA-seq Bioinformatics Workflow. We used standard pipelines and software packages for data processing and analyses. Three main steps involved: I. Data Processing for initial read processing/mapping and sequential quality control processing of raw data sets; II. Data Analysis for identification of main cell types and possible cell subtypes by unsupervised clustering, annotation, and subset analyses; III. Data Combining and Comparative Analyses of samples (Patient vs. Control) for cell type and lineage Differentially Expressed Genes (DEG) and gene set enrichment.

### Step-I. Data Processing

For this first step in our Workflow (**Fig. 1**), we processed Single Sample Data using Cell Ranger (10X Genomics) for read processing and mapping, transcript and cell quantification, and generation of summary metrics (**Table S4**). Using the ‘cellranger count’ pipeline, we processed FASTQ files of each sample to identify high-quality reads with valid cell barcodes and UMIs and mapped these reads to human reference genome (GRCh38-3.0.0) to produce BAM files. Using the number of confidently mapped reads to the transcriptome in each cell, we quantified transcripts to generate a gene-barcode matrix, an output table of cell barcodes and gene UMI counts defined by matrix.mtx, barcode.tsv, and features.tsv files for each sample. We used both raw unfiltered gene-barcode matrix (raw_feature_bc_matrix) and raw filtered gene-barcode matrix (filtered_feature_bc_matrix) in our analysis. In addition, we used Cell Ranger output BAM files (possorted_genome_bam.bam and possorted_genome_bam.bam.bai) for each sample and the Integrative Genomics Viewer (IGV) [51] to visualize aligned reads to the genome and to evaluate expressed Single Nucleotide Variants (SNV). We reported results of initial analysis for two samples, CA1-B and PA1 at Day 19 [37].

To obtain high-quality cells for clustering and downstream analysis, we conducted QC processing for Single Sample Data in three consecutive stages to identify and remove background RNA contamination, low-quality cells, and cell doublets (**Fig. 1**). We used R and RStudio version for Windows to run three software packages at each stage: A. SoupX [52]; B. Seurat [53]; and C. DoubletFinder [54] (**Fig. 1**). For each sample, QC processing generated four types of data matrices (gene-barcode matrices): 1. Raw Data Matrix (Cell Ranger filtered gene-barcode matrix), 2. Corrected Data Matrix (SoupX background-corrected gene-barcode matrix), 3. Filtered Data Matrix (Seurat gene-barcode matrix after removal of low-quality cells), 4. Singlet Data Matrix (DoubletFinder gene-barcode matrix after removal of doublet cells). To visualize effects of QC processing at each stage, we compared pre- and post-processing data by examining QC covariates and/or gene expression using frequency histograms, DimPlot, FeaturePlot, and VlnPlot functions in Seurat. The parameters and results at each stage are provided for Single Sample (**Table S5**) and Combined Data (**Table S6**).

The first stage of QC processing used SoupX [52] to process Raw Data Matrix into Corrected Data Matrix for each sample by estimating levels of background RNAs in each sample from empty droplets, estimating cell-specific contamination fraction (rho) using clustering information and negative gene markers, and producing a background-corrected gene-barcode matrix for each of the 12 samples. First, we processed the Raw Data Matrix (filtered_feature_bc_matrix) into a data object using the standard clustering pipeline in Seurat (see below). We then used the SoupX ‘SoupChannel’ function to create an object from the table of counts (toc) and the table of droplets (tod), derived from the filtered_feature_bc_matrix and unfiltered raw_feature_bc_matrix Cell Ranger output files, respectively, and to characterize the background RNAs levels (soup expression profile). Next, we estimated contamination fraction (rho), the fraction of UMIs originating from the background in each cell using two methods in SoupX: the automatic ‘autoEstCont’ function and the manual ‘calculateContaminationFraction’ function by providing a selected list of cell type-specific negative gene markers (expected non-expressed genes in at least one cluster for the sample). To remove a greater portion of the contamination, we used the greater rho value from the two methods. Possible overcorrection for background contamination likely has minimal negative effects [52]. For Day 0, we did not conduct manual calculation because the iPSC clusters had similar expression profiles and lacked appropriate negative marker genes; thus, we used the automatically estimated global rho value. For CA1 Day 0, this value was low (0.01) with low complexity (<10 marker genes found); therefore, the greater rho value (0.13) for PA1 Day 0 was used. Finally, we applied ‘adjustCounts’ to produce a background-corrected gene-barcode matrix (Corrected Data Matrix).

The second stage of QC processing used Seurat [53] to process Corrected Data Matrix into Filtered Data Matrix for each sample by identifying characteristics of low-quality cells using three common QC covariates: nFeature (numbers of genes detected), nCount (the number of UMIs), and %Mt (percentage of reads mapped to mitochondrial (Mt) genome) [48]. For each sample, we evaluated distributions of QC covariates for each cell using frequency histograms, scatter plots, and violin plots [48]. To excluded low-quality cells [55], we selected appropriate thresholds separately for each sample to filter out outlier cells that may represent dying cells with small numbers of genes (nFeature) or potential doublet cells with high number of UMIs (nCount). To filter out low-quality cells that may represent broken cells [55], thresholds for high %Mt reads also are used. However, since %Mt reads can be variable depending on cell type and energy requirements with heart having a higher Mt content [56, 57], we decided to use thresholds for %Mt reads only after clustering and cell annotation to data subsets with Non-CM cells.

The third stage of QC processing used DoubletFinder (DF) [54] to process Filtered Data Matrix into Singlet Data Matrix for each sample by predicting gene expression features of simulated doublet cells, classifying cells as either singlet and doublet, and removing heterotypic doublets. First, we processed the Filtered Data Matrix (high-quality cells) into a data object using the standard clustering pipeline in Seurat (see below). Next, we used the DF ‘paramSweep_v3’ function (PCs = 1:30) to estimate optimal pK values for each sample. Based on number of cells captured for each sample (**Table S4**), we selected values for expected proportion of doublets (Exp) from the Multiplet Rate Table provided in the User Guide for the Single Cell 3’ Gene Expression v3 assay (CG000204, 10X Genomics). Based on estimated number of cells loaded for each sample, we calculated number of expected doublets (nExp). We adjusted this number (nExp-adj) using the DF ‘modelHomotypic’ function to estimate proportion of homotypic doublets. Finally, we used these parameters in doubletFinder_v3 (pN = 0.25) to classify each cell as either a predicted singlet or doublet cell and removed Doublet cells to generate post-DF data of high-quality cells (Singlet Data Matrix).

### Step-II. Data Analysis-Cluster, Annotate, and Subset

For this second step in our Workflow (**Fig. 1**), we focused on determining cell identities for each sample in two stages: A. Individual Analyses of Singlet Data for Main Cell Types and B. Subcluster Analyses of Subset Data for Possible Cell Subtypes. For both stages, we used the standard clustering analysis workflow in Seurat: data normalization, calculation of variable features, data scaling with and without regressing out potential confounding variables (regression covariates), dimensionality reduction by Principal Component Analysis (PCA), unsupervised clustering, and visualization. We used the functions (parameters): ‘NormalizeData’ (method = "LogNormalize"), ‘FindVariableFeatures’ (nfeatures = 3000), ‘ScaleData’ (vars.to.regress), ‘RunPCA’ (npcs = 50), ‘FindNeighbors’, ‘FindClusters,’ and ‘RunUMAP.’ For dimension value (dims), we used 1:30 for Single Sample Data and 1:50 for the larger Combined Data. We tested a range of values for clustering resolution and chose final values on results with similar clusters and cell types between paired samples and on comparable results described in previous scRNA-seq studies [29–31, 33, 34, 58]. The clustering parameters used are provided for Single Sample Data (**Table S7**).

To evaluate potential confounding biological factors (regression covariates), our analysis also included cell quantification of cell cycle (CC) phase and Mt content [50, 56, 59, 60]. For each cell, we calculated scores for CC phase ("G2M.Score" and "S.Score") using the Seurat ‘CellCycleScoring’ function based on expression of 97 canonical G2/M-phase and S-phase markers [53, 60]. Using CC scores, we categorized cells as G2/M-phase, S-phase, G1-phase. For each cell, we calculated %Mt using the Seurat ‘PercentageFeatureSet’ function. Using quartiles values calculated by the ‘summary’ function in R [61], cells were categorized as "Low", "Medium", "Medium high", or "High" %Mt. After categorization for each sample, we corrected data using the function ‘ScaleData’ (vars.to.regress) to regress out %Mt, CC Scores, and both covariates and compared clustering results with and without covariate regression. The covariates used are provided for Single Sample Data (**Table S7**).

After covariate regression and clustering, we annotated clusters by Individual Analyses of Singlet Data for Main Cell Types using expression panels of known markers (**Table S8**) and Cluster DEG in Seurat. We used a Primary Marker Panel (25 genes) to identify the main cell type and Expanded Marker Panels (up to 38 genes) to confirm cell type and evaluate for possible cell subtypes among similar clusters. These Marker Panels consisted of genes expressed in reported cell types and subtypes during iPSC-CM differentiation and maturation [20, 62–65] and in previous scRNA-seq studies for iPSC [28], endoderm [66, 67], iPSC-CM or hES-CM [29–31, 33, 34, 36, 58], and human tissues [68]. To detect residual undifferentiated cells, we included the *CNMD* gene [69]. After cell annotation, we confirmed our results by evaluating Cluster DEG for known cell type markers and signatures by gene set enrichment analysis (see below). First, we found Cluster DEG using ‘FindAllMarkers’ (test.use= “wilcox”, logfc.threshold = 0.25, min.pct = 0.1) and then ranked top DEG for each cluster by log fold-change of the average expression relative to all other clusters. We compared and visualized cluster expression for known markers and top DEG using DimPlot, VlnPlot, FeaturePlot, and DoHeatmap functions in Seurat. If cell type was not clearly established after evaluation, we annotated the cluster as “Unknown.”

After annotation for main cell types, we conducted Subcluster Analyses of Subset Data for Possible Cell Subtypes, using similar steps in our Individual Analyses as described above. First, we divided annotated Single Sample Data into separate subsets by cell type(s) using the ‘subset’ function in Seurat. We separated CM, Non-CM, and Unknown cells into different subsets for cell type-specific QC processing and removal of Unknown cells prior to data integration and Comparative Analyses. Our subset data analysis involved: 1. Evaluation for common QC covariates; 2. Cell type-specific QC processing that used high threshold levels for %Mt to identify low-quality cells in Non-CM cells; 3. Correction by regressing out CC Scores that might confound trajectory inference [48]; 4. Unsupervised clustering using the standard clustering pipeline in Seurat; and 5. Cell annotation using Marker Panels (**Table S8**) and Subcluster DEG. In addition, we used two Cell Subtype Marker Panels to identify possible CM subtypes (27 genes for ventricular, atrial, and nodal cells) [30, 63, 70] and EPDC subtypes (27 genes for EPDC Progenitors, Cardiac Fibroblast, Vascular Smooth Muscle, and Angioblasts/Endothelial cells) [34, 58, 63] (**Table S8**). For Unknown cells, we classified cells into possible subtypes using Marker Panels, Subcluster DEG, and common QC covariates: nFeature, nCount, and %Mt. The parameters for cell type-specific QC processing and subclustering are provided for Single Sample Data (**Table S7**).

### Step-III. Data Combining and Comparative Analyses

For the third step in our Workflow (**Fig. 1**), we created two types of Combined Data from the 12 samples: Integrated Data primarily to compare paired samples, Patient versus (vs.) Control at D00, D09B, D16, and D19, by cell type and lineage differential gene expression and pathway enrichment and Merged Data (Non-Integrated) to summarize and evaluate our results. To evaluate data integration, we compared clustering results between Merged Data without integration and Integrated Data with integration using Seurat. In Combined Data, we also compared the “imbalance score” for each cell to test whether nearby cells have the same condition using the ‘imbalance_score (k = 20, smooth = 40)’ function in Condiments [71]. We defined ‘balanced’ cell types and subtypes as shared cells with low imbalance scores compared to the other cells in Combined Data. The parameters are provided for integration and evaluation of Combined Data for Paired Sample Data and Subset Data (**Table S9**).

To identify ‘balanced’ cell types/subtypes for Comparative Analyses between conditions, we first analyzed Combined Data of paired samples (D00, D09B, D16, and D19). Like our analyses for Single Sample Data, we conducted both Individual and Subcluster Analyses of Combined Data of Paired Sample Data (**Table S9**). First, we created Merged Data using the ‘merge’ function to combine two processed Seurat objects from CA1 and PA1: Paired Sample Data (Singlet) (n=4 pairs) or Paired Subset Data (n=11 pairs). Next, we created Integrated Data (**Table S9**) first by splitting Merged Data into a list of objects by condition (Patient and Control) using the ‘SplitObject’ function (split.by = "orig.ident"). After normalization, we applied the standard canonical correlation analysis (CCA) integration pipeline [53]: CCA to identify shared variation between conditions, identification of “anchors,” and alignment of Control and Patient cells into an integrated Seurat object. We used the functions (parameters): ‘SelectIntegrationFeatures’ (nfeatures = 3000), ‘FindIntegrationAnchors,’ and ‘IntegrateData.’ After standard clustering analysis, we annotated as described above using Marker Panels (**Table S8**) and Cluster DEG. For Integrated Data, we identified Cluster DEG conserved between conditions using the function ‘FindConservedMarkers’ (grouping.var = "Sample", test.use = "wilcox", logfc.threshold = 0.25, min.pct = 0.1). We ranked Cluster DEG by average fold change of expression (avg_fc = (PAT_avg_log2FC + CTRL_avg_log2FC) /2)), visualized top Cluster DEG using ‘DoHeatmap’ in Seurat, and confirmed cell types by Over-Representation Analysis (ORA) [72] as described below.

To identify cell type-specific differentially expressed genes (Cell Type DEG) between conditions, we conducted Comparative Analyses (Patient vs. Control) in ‘balanced’ cell subtypes of Integrated Data. First, we identified all Cell Type DEG between conditions using the function ‘FindMarkers’ (ident.1 = “patient”, ident.2 = “control”, test.use= “wilcox”, logfc.threshold = 0). Next, we divided Cell Type DEG by direction of change in Patient cells compared to Control cells: overexpressed genes (positive DEG, logfc.threshold > 0) and underexpressed genes (negative DEG, logfc.threshold < 0). We identified and ranked top Cell Type DEG using chosen thresholds for statistical significance and difference in average expression (adjusted p-value < 10e-50 and I Average Log2 FoldChange I > 0.25 (Fold Change = 1.2x)). We visualized and compared Cell Type DEG using ‘EnhancedVolcano’ [73] and ‘VennDiagram’ [74].

To identify biological process or pathways altered between conditions, we evaluated Cell Type DEG by enrichment analysis using two common approaches: Over-Representation Analysis (ORA) [72] and Gene Set Enrichment Analysis (GSEA) [75]. For each cell subtype, we tested individually the lists of threshold DEG (overexpressed or underexpressed genes with minimum DEG = 3) for overrepresentation in the Gene Ontology (GO) biological processes gene sets (n = 7751) [76] using the ‘EnrichGo’ function (adjusted p-value < 0.05, q-value cutoff = 0.20) in Cluster Profiler [77]. To evaluate all DEG (selected without a threshold), we first ordered the complete lists of DEG (overexpressed and underexpressed genes) by GSEA metric (= -log10(p_val_adj) * sign(avg_log2FC)). We then tested ranked lists for enrichment in the Molecular Signatures Database (MSigDB) Hallmark gene sets (n = 50) [75, 78] using the ‘GSEA’ function (GSEA (minGSSize = 10, maxGSSize = 500, eps = 1e-10, p-value Cutoff = 0.05) in Cluster Profiler [77]. We visualized enrichment results using ‘dotplot’, ‘emapplot’, and ‘gseaplot2’ in Cluster Profiler [77] and used Module Scoring with ‘AddModuleScore’ function in Seurat to quantify, compare, and visualize patterns of expression of key DEG in significant ORA and GSEA gene sets.

To evaluate lineage-specific differential expression, we conducted trajectory inference using Slingshot [79] and expression analysis using TradeSeq [80]. To determine possible cell lineages in each sample, we first analyzed Single Subset Data (**Table S10**) for selected cell subtypes and adjacent subtypes of one condition (Control or Patient). First, we converted each processed object to a SingleCellExperiment (SCE) data class using the ‘as.SingleCellExperiment’ function in Seurat. Next, in Slingshot, we conducted trajectory inference with semi-supervision with ‘slingshot’ or ‘getLineages’ and ‘getCurves’ (reducedDim = ’UMAP’, approx_points = FALSE, stretch = 0, extend = "n") and ‘slingMST.’ These functions identified lineage topology (global lineage structure) by constructing a minimum spanning tree (MST) on clusters, fitted a principal curve through data that defined a trajectory, and calculated a pseudotime variable for each cell. For each sample, we selected starting (start.clus) and ending (end.clus) clusters based on progression of cell types and subtypes during iPSC-CM differentiation [20, 62] and cell lineages reported in previous scRNA-seq studies for iPSC [28], endoderm [67], and iPSC-CM or hES-CM [29, 31, 32, 34–36]. We visualized results in UMAP plots generated using ‘ggplot’ and base R ‘plot’ and density plots for progression.

Next, to identify differential expressed genes within each lineage (Lineage DEG), we analyzed SCE objects in TradeSeq [80] (**Table S10**). First, we used the ‘fitGAM’ function to make a general additive model (GAM) of relationship between expression of each gene and pseudotime. Using the ‘evaluateK’ function (k = 3:10, nGenes = 200), we selected the number of knots. We tested genes for two types of Lineage DEG (Threshold: FDR <0.05 & Fold Change > 2x): Association Test (AT) DEG and Start-End Test (SET) DEG. Using the ‘associationTest’ function, we identified AT Lineage DEG for which average gene expression significantly changed along pseudotime of the lineage. Using the ‘startVsEndTest’ function, we identified SET Lineage DEG for which average gene expression significantly changed between starting and ending pseudotime points of the lineage. For samples with a bifurcating trajectory, we identified Global AT and SET Lineage DEG by testing across both lineages (global = TRUE). We visualized and compared expression patterns of top Lineage DEG and known marker genes using ‘pheatmap’ and Tradeseq functions ‘plotSmoothers’ and ‘plotGeneCount’.

After integrating Paired Subset Data (**Table S11**), we conducted Comparative Analyses for lineage differential expression across conditions (Patient vs. Control) and pathway enrichment analyses. As described above for Single Subset Data, we analyzed Integrated Subset Data for trajectory inference and lineage differential expression using Slingshot [79] and Tradeseq [80] (**Table S11**). Using Condiments [71], we then tested for differences in lineage topology, progression, differentiation, and expression between conditions. Using the ‘topologyTest’ (rep = 100) function, we tested for differential topology: different individual trajectories for each condition. Using ‘progressionTest’ (global = TRUE, lineages = TRUE), we tested for differential progression: different distributions of cells found along the trajectories for each condition. For bifurcating trajectories, we used ‘differentiationTest’ (global = TRUE, pairwise = TRUE) for differential differentiation: different cell distributions between lineages for each condition.

For Integrated Subset Data (**Table S11**), we then tested for two types of Lineage DEG (Threshold: FDR <0.05 & Fold Change > 2x): Association Test (AT) and Condition Test (CT) Lineage DEG. Using the ‘associationTest’ function, we identified AT Lineage DEG for which average gene expression significantly changed along pseudotime of the lineage for both conditions. Using the ‘conditionTest’, we identified CT Lineage DEG for which average gene expression significantly changed along pseudotime of the lineage between conditions. For bifurcating trajectories, we tested for differential expression between conditions to identify Global AT and CT Lineage DEG that changed for both lineages (global = TRUE) and CT Lineage-Specific DEG that changed in only one lineage (lineages = TRUE).

As described above for Cell Type DEG, we conducted enrichment analyses for Lineage DEG by ORA [72] using Cluster Profiler [77] to identify biological processes or pathways that changed with pseudotime in Single Subset Data (**Table S10**) and Integrated Subset Data (**Table S11**). We conducted ORA using top 100 AT, SET, and CT Lineage DEG (ranked by Wald statistic) and smaller groups of DEG obtained by hierarchical cluster analysis using the functions ‘hclust’ and ‘cutree’ (k=2 to 4). In addition, like Cell Type DEG, we evaluated all CT Lineage DEG (FDR <0.05 & WaldStat > 0.0) using GSEA (scoreType = "pos") for enrichment of MSigDB Hallmark gene sets [75, 78] in Cluster Profiler [77].

### Lamin A/C Western Blot

To compare Lamin A and C proteins levels in Control and Patient cells, we conducted WB as described [81] and quantitative analyses [82]. Using Protocol-A (**Fig. S1**), we differentiated three iPSC pairs as biological replicates (BR): three Patient-derived iPSC (PA1, PA2, and PA3) and three Control iPSC (CA1-B, U2, and CA3) (**Table S1**) to Day 19. Cell samples were collected and lysed using RIPA Lysis and Extraction Buffer with Inhibitor Cocktail (TFS) at a ratio of 100 μL for every 1,000,000 cells. Using the Pierce BCA Protein Assay Kit (TFS), we determined protein concentration of cell lysate. We combined 20 μg of total protein lysate, Bolt LDS Sample Buffer and Reducing Agent (TFS), and distilled water in 30 μL total that was heated and loaded on to Bolt 4-12% Bis-Tris Plus Gels (TFS). After gel electrophoresis, we used the Mini Blot Module (TFS) to transfer proteins to a PVDF membrane that was processed using the iBind Western Device and iBind Fluorescent Detection solutions (TFS). To detect target proteins, we used primary antibodies for the Lamin A/C N-terminus (sc-376248, Santa Cruz Biotechnology, Dallas, TX) and Beta-Actin as the internal loading control and fluorophore-conjugated secondary antibodies for visualization (**Table S2**). As a positive control for Lamin A/C expression, we included protein lysate from Control (CA1) fibroblast. We repeated gel electrophoresis and immunoblotting to produce three technical replicates (TR).

For protein visualization and quantification of band signal, we used the Azure c600 imaging system and AzureSpot software (Azure Biosystems, Dublin, CA). For each TR blot, we first determined protein levels as normalized ratios (NR) of Lamin A, Lamin C, and total Lamin A+C protein for each individual sample (n=6: CA1-B, U2, CA3, PA1, PA2, PA3) by dividing the Lamin A/C band volume (BV) with the corresponding Beta-Actin BV. From the three TR blots, we then calculated mean protein levels (Mean NR of TR) of Lamin A, Lamin C, and Lamin A/C for each sample, and from the three biological replicates (BR), we calculated mean protein levels (Mean NR of BR) for Control samples (n=3: CA1-B, U2, CA3) and Patient samples (n=3: PA1, PA2, PA3). We also compared Patient samples pairwise to biologically (sex and age) matched Control samples (PA1 to CA1, PA2 to U2, PA3 to CA3) using Relative Normalized Ratio (RNR) by dividing the Lamin A/C protein level (NR) of the Patient sample with the Control sample to calculate mean fold change (FC) and percent change (% change) of TR. To evaluate technical variability between blots for pairwise comparisons, we calculated coefficient of variation (CV= SD of TR/ Mean FC of TR) and % change/CV. For statistical comparisons of Patient and Control data, we used the independent samples t-test and defined significance as *p* < 0.05.

## RESULTS

### Validated iPSC and collected 12 cell samples (8 Control and 4 Patient) during differentiation for scRNA-seq

We used eight validated iPSC lines including three *LMNA* Patient (PA1, PA2, PA3) and five Control (CA1-A, CA1-B, CA2, CA3, U2) iPSC lines (**Table S1**). Validation results included normal karyotype at passage number 9 or above, normal pluripotency by ICC staining for pluripotent stem cell markers [41], and normal ICC staining for germ layer markers in EB for all eight iPSC lines (**Fig. S2**). We differentiated seven iPSC lines and excluded one line (CA2) due to poor cell viability.

During iPSC differentiation, we collected 12 cell samples for scRNA-seq consisting of four Patient and eight Control cell samples. To identify the main cell types, possible cell subtypes, and cell lineages during non-mutant iPSC differentiation into CM, we differentiated two Control iPSC lines (CA1-A and CA1-B) and collected eight Control cell samples across seven time points (D00, D02, D04, D09, D16, D19, D30) for scRNA-seq. To study possible disease mechanisms, we differentiated in parallel one Patient iPSC line (PA1) and collected four Patient cell samples across four time points (D00, D09, D16, D19) for scRNA-seq. Results for allelic expression of 22 coding SNV (10 autosomal and 12 XL SNV), compared to fibroblast genotype, confirmed the identification of the 12 cell samples collected for scRNA-seq (**Fig. S3**).

### Processed data: 110,521 (88%) high-quality cells of 125,554 cells collected (Workflow Step-I)

To identify high-quality cells for clustering and downstream analysis, our bioinformatics workflow (**Fig. 1**) began with read processing and mapping followed by sequential quality control (QC) processing for each the 12 cell samples. For Single Sample Data and Combined Data, the parameters and results are provided in (**Table S4-6**, **Fig. S4**).

First, we processed FASTQ sequence files for the 12 cell samples using Cell Ranger to obtain summary metrics for read mapping and cell/gene quantification (**Table S4**). The metrics showed that deep sequencing estimated 125,554 total cells captured with 40,978 mean reads per cell and 93.2% Reads Mapped Confidently to Genome. The eight Control samples had greater total number of captured cells with similar values for % Reads Mapped compared to the four Patient samples. These results are consistent with high-quality sequencing data [83] and sufficient read depth for an initial analysis to identify main cell types in both Control and Patient cell samples [84]

Next, we conducted QC processing for Single Sample Data in three consecutive stages to identify and remove background RNA contamination using SoupX, low-quality cells using Seurat, and doublet cells using DoubletFinder that generated four types of data (gene-barcode matrices): 1. Raw Data Matrix, 2. Corrected Data Matrix, 3. Filtered Data Matrix, 4. Singlet Data Matrix for each sample (**Table S6**) and for Combined Data (**Fig. S4**). QC processing of Raw Data Matrices identified 110,521 (88%) high-quality cells of 125,554 total cells and similar proportions of high-quality cells in eight Control samples and four Patient samples (**Fig. S4A**). Of 15,032 (12%) total cells removed, 8,272 (55%) were identified as low-quality cells and 6,761 (45%) as doublet cells. By comparing Merged Data for all 12 samples after each stage, our results demonstrate significant effects of QC processing not only in the total number of cells at each stage but also in clustering and gene expression patterns after background RNA removal (**Fig. S4E**) and in distributions of QC covariates (nFeature, nCount, and %Mt) between different samples and clusters (**Fig. S4BCD**), between low-quality and high-quality cells (**Fig. S4F**), and between singlet and doublet cells (**Fig. S4F**). For example, although Control and Patient samples had similar median values of the QC covariates for each cell (**Fig. S4B**), distributions of QC covariates varied by sample (**Fig. S4C**) and by cluster (**Fig. S4D**). Overall, these results emphasize the importance of comprehensive QC processing with assessment after each stage and using parameters that were determined separately for each sample [48] (**Table S5**) and for different cell types (CM vs. Non-CM) (**Table S7**).

### Complex heterogeneity with ten main cell types in Control samples and eight shared cell types between paired Patient and Control samples (Workflow Step-IIA)

To identify main cell types and possible subtypes, the second step in our bioinformatics workflow (**Fig. 1**) involved unsupervised clustering, annotation, and subsetting using Marker Panels (88 total genes, **Table S8**) of processed data for the 12 samples (8 Control and 4 Patient samples: 110,521 total cells). For Single Sample Data (**Table S7**, **Fig. S5A**) and Combined Data (**Fig. 2**, **Fig. S5B)**, results are provided as two stages: Individual Analyses of Singlet Data for Main Cell Types (**Fig. S6**) and Subcluster Analyses of Subset Data for Possible Cell Subtypes (**Fig. S7**).

**Figure 2.**
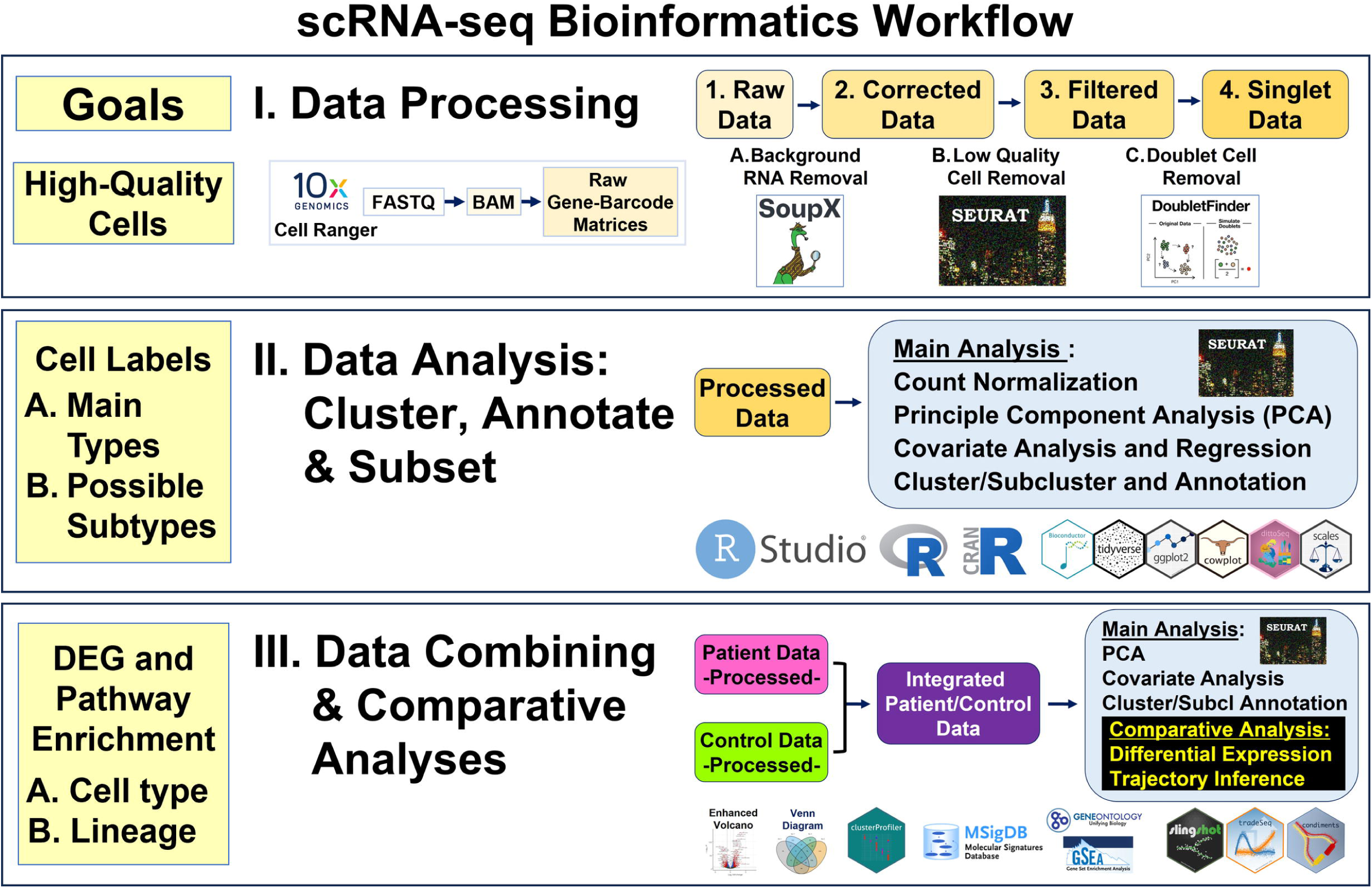
Complex Heterogeneity: Main Cell Types and Shared Subtypes. Our scRNA-seq bioinformatics workflow included analyses of Combined Data: Merged Data (Top Panel) and Integrated Data (Bottom Panel). Each panel consists of four UMAP plots from left to right with clusters marked and colored by time point, pre-annotation cluster number, main cell type or shared subtype, and imbalance score. To summarize results from our Individual Analyses, we created Merged (Non-Integrated) Data (Top Panel) by combining processed Single Sample Data for all 12 samples (110,521 total cells) across all seven time points: D00, D02, D04, D09A/B, D16, D19, and D30 (First plot). To compare Patient and Control samples, we created Integrated Data (Bottom Panel) by combining processed Subset Data for six pairs of ‘balanced’ cell types/subtypes (75,330 total cells) at four time points: D00, D09B, D16, and D19 (First plot). Unsupervised clustering (2^nd^ plots) found 49 clusters in Merged Data and 17 clusters in Integrated Data. Using known markers (3rd plots), we identified 10 major cell types: Pluripotent cells (PP), Mesendoderm (ME), Cardiogenic Mesoderm (CMESO), Endoderm (ENDO), Ectoderm (ECTO), Endothelium (ENDOTH), Cardiac Progenitors (CP), Cardiomyocyte (CM), Epicardium-Derived Cells (EPDC), and Unknown (UNK) in Merged Data and multiple possible shared subtypes in Integrated Data. Using Condiments [71], we calculated the “Imbalance Score” of each cell to assess if nearby cells had the same condition (Last plots). We defined ‘balanced’ cell types/subtypes, as shared cells with low Imbalance Scores, that were then selected for Comparative Analyses between Patient and Control cells.

From our Individual Analyses for the 12 samples, we identified ten main cell types in Control samples and eight conserved cell types between paired samples (**Fig. 2**, **Fig. S5A**, **Fig. S6**, **Table S7**). In Control samples (**Fig. S6B**) collected at seven time points (D00, D02, D04, D09, D16, D19, D30), we identified ten main cell types: 1. Pluripotent cells (PP), 2. Mesendoderm (ME), 3. Cardiogenic Mesoderm (CMESO), 4. Endoderm (ENDO), 5. Ectoderm (ECTO), 6. Endothelium (ENDOTH), 7. Cardiac Progenitors (CP), 8. Cardiomyocyte (CM), 9. Epicardium-Derived Cells (EPDC), and 10. Unknown (UNK). Of these ten cell types, we found eight (PP, ENDO, ECTO, ENDOTH, CP, CM, EPDC, UNK) conserved in the four Patient samples (D00, D09B, D16, D19) (**Fig. S6C**) and two cell types (ME and CMESO) identified only in Control samples (D02 and D04). For all 12 samples, we found clusters with unclear results and assigned these cells into an “Unknown” category (UNK). Furthermore, at this initial stage, we found CM consisting of two possible subtypes: clusters with high expression of multiple known CM markers (CM-A) and clusters with lower expression of fewer CM markers except *TTN* and higher levels of %Mt (CM/UNK-B).

Overall, cell type heterogeneity increased during early iPSC differentiation with all ten cell types identified from Day 0 to Day 9; thereafter, CM and EPDC became the predominant cell types (**Fig. S6**). At Day 0, PP cells (89%) served as the predominate cell type with expression of three marker genes, *POU5F1*, *SOX2*, and *NANOG*. At Day 2, three early differentiated cell types (96%) predominated that were identified by marker expression of *EOMES* for ME, *MESP1* for CMESO, and *SOX17* for ENDO. At Day 4, nearly all cells (98%) were identified as either PP, CMESO, ENDO, or CP with expression of *HAND1*, *HAPLN1*, and *TMEM88* to mark CP cells. At Day 9, differentiation of two independent clones (CA1-A and CA1-B) showed increased cell heterogeneity with most cells (69-88%) as CP cells or CP-derived cells (CM-A, CM/UNK-B, and EPDC) with multiple marker expression that included *MYL7*, *TNNI1*, and *NKX2-5* for CM and *COL3A1*, *LUM*, and *FBN1* for EPDC. Day 9 also consisted of ENDO cells with *FOXA2* expression and ENDOTH cells with *EGFL2* expression, two cell types derived from both clones and ECTO cells with *PAX6* expression, a cell type derived from one clone (CA1-B). At Day 16 and Day 19 (post-metabolic selection), we identified most cells (∼75%) as CM (CM-A and CM/UNK-B) and a smaller subset as EPDC (13%). At Day 30 (without metabolic selection), these two types of cells comprised most cells (82%); however, CM (38%) were found in a lower proportion compared to EPDC (44%). For all samples, we confirmed cell type by finding known marker genes among top DEG for the cluster (Cluster DEG) (**Fig. S6**, **Table S7**, **Excel Table S1-1**).

### CM and EPDC differentiated into possible subtypes with differences between paired Patient and Control samples (Workflow Step-IIB)

In our Subcluster Analyses for all 12 samples (8 Control and 4 Patient samples), we subsetted Single Sample Data into 32 total subsets and found 30 total possible subtypes in eight cell types (PP, ME, ENDO, ECTO, CP, CM, EPDC, UNK) (**Fig. S5A**, **Fig. S7**, **Table S7**, **Excel Table S1-2**). For eight paired samples (D00, D09B, D16, D19), we found 19 possible subtypes in six cell types (PP, ECTO, CP, CM, EPDC, UNK): 13 subtypes conserved in both conditions and 6 subtypes in one condition.

For Control samples, both CM and EPDC showed maturation and then differentiation into cell subtypes. For Control CM, we found four possible subtypes: an early pair (CM-A, CM/UNK-B) at Days 9 to 19 and a further differentiated pair: Atrial CM (ATR-CM) and Ventricular CM (VENTR-CM) at Day 30 (**Fig. S7B**). Similarly, for Control EPDC, we found seven possible subtypes: an early set at Day 16 (EPDC-A, EPDC-B, EPDC-C) and a further differentiated set: Epicardial Progenitor (EPI), Cardiac Fibroblast (CFIBRO), Vascular Smooth Muscle (VSM), and an Unspecified EPDC subtype (EPDC-UNSP) at Day 30 (**Fig. S7B**). From Days 9 to 19, both CM and EPDC had results consistent with cell type maturation as increased proportions of cells with expression of later markers (CM: *TTN*, *MYH7*, *MYL2;* EPDC: *COL3A1*, *LUM*, *FBN1*) compared to early markers (CM: *MYL7*, *TNNI1*, *NKX2-5*; EPDC: *TBX18*, *TCF21*, *WT1*). At Day 30, early subtypes were not found; instead, we identified the six later differentiated subtypes: ATR-CM, VENTR-CM, EPI, CFIBRO, VSM, an Unspecified EPDC subtype with differentiating patterns of gene expression in CM and EPDC Subtype Marker Panels (**Fig. S7B**).

For eight paired Control and Patient samples, we found 19 total possible subtypes in six cell types (PP, ECTO, CP, CM, EPDC, UNK): 13 subtypes conserved and six subtypes in one condition (**Fig. S7**). For PP and CM, we found only conserved subtypes. Both Control and Patient PP cells (D00: Subset-A) had four possible subtypes: PP-A (88-92%), PP-B, PP-C, and PP-D with variable expression levels of ten PP marker genes (*POU5F1*, *SOX2*, *NANOG*, *DNMT3B*, *NODAL*, *UTF1*, *LIN28A*, *LEFTY1*, *GDF3*, and *SDC2*) and one undifferentiated cell marker: *CMND*. Likewise, both Control and Patient early CM (D09B: Subset-A) and post-metabolic selection CM (D16 and D19: Subset-A) had two possible subtypes (CM-A and CM/UNK-B) with distinguishing features as seen in our Individual Analyses. In addition to lower expression levels of Mt DNA genes, CM-A cells had higher expression of Ribosomal Protein Genes (*RPL* and *RPS*) compared to CM/UNK-B cells.

In contrast to PP and CM, we found subtype differences for four cell types: CP, EPDC, ECTO, and UNK between paired Control and Patient samples (**Fig. S7**). While both Control and Patient CP cells (D09B: Subset-A) had two possible subtypes (CP-A and CP-B), we identified a third subtype (CP-C) in only the Patient sample using nine CP marker genes (*HAND1*, *HAPLN1*, *TMEM88*, *GATA4*, *HCN4*, *TBX5*, *ISL1*, *TBX1*, and *HAND2*). Likewise, both Control and Patient EPDC (D16: Subset-B) had two possible subtypes (EPDC-A and EPDC-B) and a third subtype (EPDC-C) in only the Patient sample. For ECTO (D09B: Subset-B), we identified four possible subtypes: ECTO-A, ECTO-B, ECTO-C, ECTO-D with later two subtypes in only Patient cells; however, these subtypes each had low cell number (∼130 to 250 cells) using only two ECTO markers: *PAX6*, *SOX1*. For Unknown cells, we found three possible subtypes: UNK-A, UNK-B, and UNK-C using levels of covariates and marker expression. We defined UNK-A as cells with low nCount levels, UNK-B as cells with low nCounts levels and highest %Mt levels, and UNK-C as cells with higher nCount and Marker expression levels compared to the other possible subtypes. While both Control and Patient cells (D00: Subset-B and D09B, D16, D19: Subset-C) had the UNK-A subtype, the other two subtypes had subsets with only Control cells (UNK-B D16:Subset-C) or Patient cells (UNK-C D16 and D19: Subset-C).

### Four shared main cell types: PP, CP, CM, EPDC with primarily ‘balanced’ cell subtypes in Integrated Patient and Control Data (Workflow Step-III)

The third step in our bioinformatics workflow (**Fig. 1**) focused on creation and evaluation of two types of Combined Data: Integrated Data of paired samples for differential expression analysis and Merged Data (Non-Integrated) to summarize and evaluate our results. For Combined Data of Paired Sample Data (**Fig. S8A**, **Table S9**), results are provided for Individual Analyses (n=4 pairs: 89,269 total cells) in **Fig. S9** and Subcluster Analyses (n=11 pairs: 88,420 total cells) in **Fig. S10**.

To identify ‘balanced’ cell types/subtypes for differential expression analysis between conditions, we created Combined Data: Integrated and Merged Data for each pair of Patient and Control samples (D00, D09B, D16, D19) and conducted both Individual and Subcluster Analyses for shared cell types and subtypes (**Fig. 2B**, **Fig. S8AB**). Overall, our analyses found Patient and Control cells clustered separately by condition in Merged Data and clustered together by shared cell type/subtype in Integrated Data (**Fig. S9A**, **Fig. S10A**). From our Individual Analyses of Integrated Data (**Fig. S8AB**, **Fig. S9B**, **Table S9, Excel Table S1-3, S1-5, S1-6)**, we identified eight shared cell types (PP, ENDO, ECTO, ENDOTH, CP, CM, EPDC, UNK), the same cell types conserved in Single Sample Data (**Fig. S6, Table S7**, **Excel Table S1-1)**. From our Subcluster Analyses (**Fig. S10BC**, **Table S9**, **Excel Table S1-4**), we also found the same 19 possible subtypes in six cell types (PP, ECTO, CP, CM, EPDC, UNK) in both Single Sample (**Fig. S7, Table S7**, **Excel Table S1-2)** and Integrated Data. However, of eight shared cell types, imbalance scoring identified one cell type (CP) with all ‘balanced’ subtypes, four cell types (PP, ECTO, CM, EPDC) with both ‘balanced’ and ‘imbalanced’ subtypes, and three cell types (ENDO, ENDOTH, UNK) with all ‘imbalanced’ subtypes (**Table S9B**). Overall, four shared cell types (PP, CP, CM, EPDC) in Integrated Data had mostly ‘balanced’ cell subtypes.

### Cell type-specific differential expression: *LMNA* DCM-related gene, X-linked genes including the non-coding *XIST* RNA, and multiple imprinted genes (Workflow Step-IIIA)

From four shared main cell types (PP, CP, CM, EPDC) in Integrated Data, we selected 14 primarily ‘balanced’ subtypes (71,541 total cells) (**Table S9B**) for differential expression and enrichment analyses (**Table, Fig. S11ABC**). Across these 14 subtypes, we found similar average proportions of Patient (52%) and Control (48%) cells and total proportions of overexpressed DEG (25,551 of 53,548: 48%) and underexpressed DEG (27,997 of 53,548: 52%) in Patient cells compared to Control cells.

Among top Cell Type DEG, we found our DCM study gene *LMNA* underexpressed in Patient cells compared to Control cells (**Fig. 3**, **Fig. S11D**, **Excel Table S2-1**, **S2-2**). We found *LMNA* underexpressed in 12 of the 14 shared cell subtypes and as a top ten threshold DEG in two subtypes: D19 CM-A1 and D19 EPDC. Across all 14 subtypes from Days 0 to 19, *LMNA* expression increased with highest expression levels in CM and EPDC. At D19, both Patient CM and EPDC had lower *LMNA* expression compared to Control cells. In addition, Patient cells showed primarily monoallelic expression from only the normal allele when we examined the heterozygous *LMNA* coding SNV (C>T, rs4641) (**Table S1**, **Fig. S3A**, **Excel Table S4-1**). In contrast, we did not identify B-type Lamin genes, *LMNB1* and *LMNB2*, as threshold DEG. Expression was limited to PP and EPDC/CP cells for *LMNB1* and to PP for *LMNB2* with no differences between Patient and Control cells (**Fig. S11D**) and with biallelic *LMNB2* expression in Patient cells (**Fig. S3A**).

**Figure 3.**
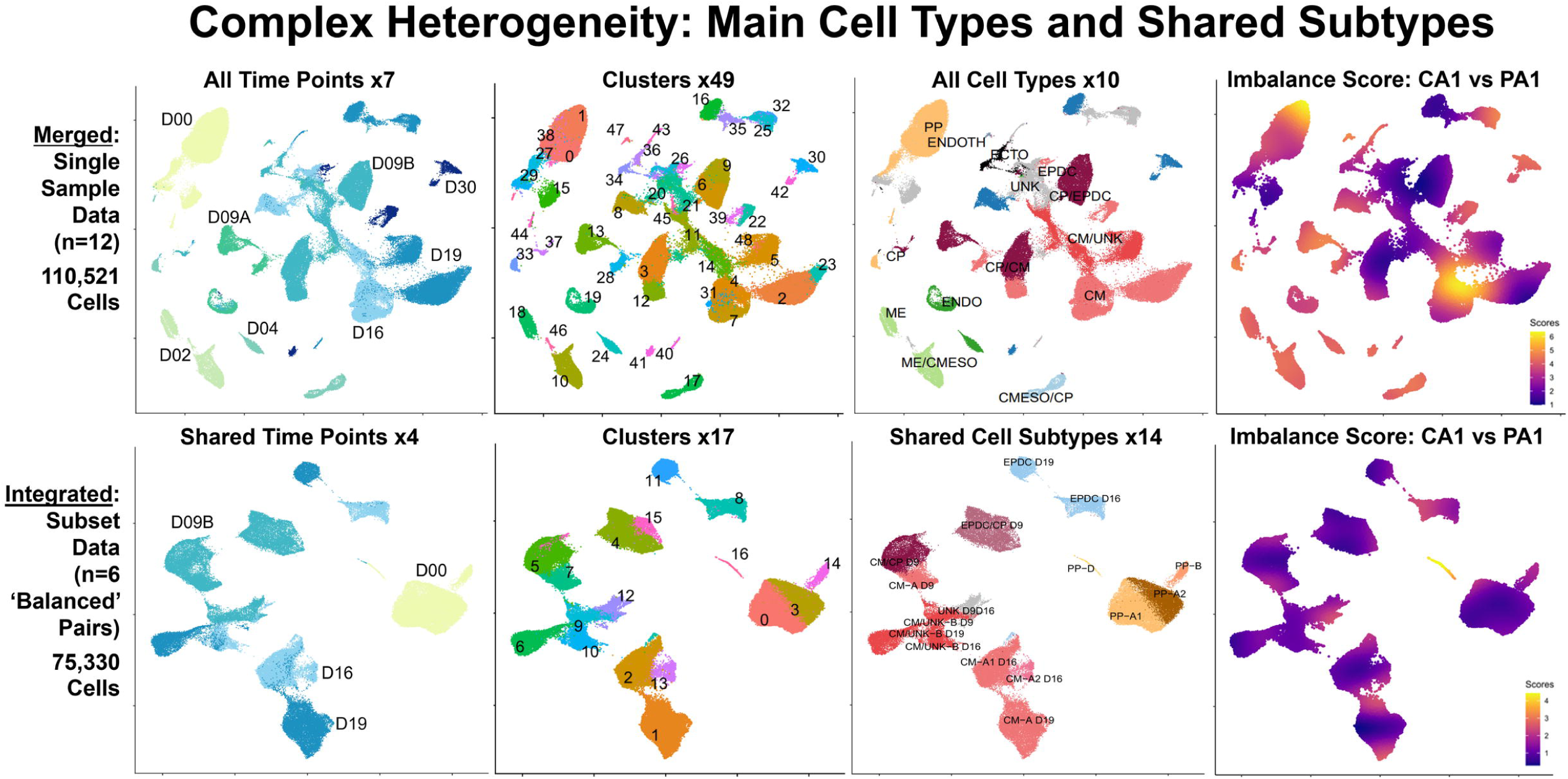
Cell Type-Specific Differentially Expressed Genes (Cell Type DEG): Top DEG. Our Comparative Analyses (Patient vs. Control cells) included cell type-specific differential expression and enrichment using Integrated Data at four time points: D00, D09B, D16, and D19. We identified Cell Type DEG in six pairs of ‘balanced’ cell types/subtypes with 75,330 total cells. Left Panel: Heat map for top DEG in main cell types for 38,150 (51%) Control cells and 37,180 (49%) Patient Cells. Top DEG included *LMNA*, 11 X-linked (XL) genes, and six imprinted genes. Right Panel: Violin Plots with boxplot by Sample, Day, and Cell Type for *LMNA*, XL genes: *XIST* and *GPC3*, and imprinted genes: *SNRPN* and *MEG3*. Overall, these results show Patient compared to Control cells had *LMNA* underexpressed in both CM and EPDC at Day 19, limited *XIST* expression for all time points and shared cell types, XL gene *GPC3* overexpressed in CM/CP and EPDC, *SNRPN* underexpressed for all time points and most shared cell types, and *MEG3* overexpressed in PP and EPDC.

Top Cell Type DEG also included underexpressed X-linked (XL) genes including *XIST*, the non-coding RNA gene that initiates X chromosome inactivation (XCI) [85], and *PIN4*, an isomerase encoding gene with roles in cell cycle progression and chromatin remodeling [86] (**Fig. 3**, **Fig. S11D**, **Excel Table S2-1**, **S2-2**). Of 47 genes found as a top ten underexpressed threshold DEG, we found two (4%) XL genes: *XIST* and *PIN4*. We found *PIN4* underexpressed in 11 of the 14 shared cell subtypes and as a top ten threshold DEG in only D00 PP-A1A2, the cell subtype with highest expression levels (**Fig. S11D**). In contrast, we found *XIST* as the only gene ranked in the top ten threshold DEG shared in all 14 subtypes. For all time points and shared cell types, we detected no expression or lower expression levels of *XIST* in Patient cells compared to Control cells (**Fig. 3**, **Fig. S11D**). In addition, despite limited *XIST* expression in Patient cells, we detected primarily monoallelic expression using heterozygous coding SNV in seven XL genes known to be subject to XCI (**Fig. S3B**, **Excel Table S4-2**).

Top Cell Type DEG also included multiple overexpressed XL genes including *GPC3*, encoding Glypican Proteoglycan 3, a cell-surface glycoprotein involved in growth regulation [87]. Of 71 genes found as a top ten overexpressed threshold DEG, we found nine (13%) XL genes: *PCSK1N*, *SMPX*, *RBBP7 (TXLNG)*, *CD99*, *RPS4X*, *DIAPH2*, *MORF4L2*, *PRPS1*, and *GPC3* across 7 of the 14 shared cell subtypes (**Excel Table S2-2**). For overexpressed XL DEG, expression patterns varied across seven cell subtypes. For example, we found as a threshold DEG: *GPC3* in two cell subtypes: D09B CM-A and D19 EPDC (**Fig. 3, Fig. S11D**), four genes: *RBBP7 (TXLNG)*, *DIAPH2*, *MORF4L2*, and *PRPS1* in only one subtype D00 PP-A1A2, and both *CD99* and *RPS4X* in three subtypes: D09B CP-A, D09A CM-A, and D19 EPDC (**Excel Table S2-1**, **S2-2**). In addition, we detected biallelic expression of *CD99* in Patient cells using a heterozygous coding SNV (C>T, rs1136447) (**Fig. S3B**). Since *CD99* is known to escape XCI [88], we might expect this result; however, the proportion of allelic expression differed. In Patients cells, we detected equal total expression of each allele (Total read count= 3528, C 51% and T 49%) compared to skewed expression of one allele (Total read count= 3342, C: 81% and T:19%) in Control cells (**Excel Table S4-2**). We also found similar results for another gene known to escape XCI, *EIF2S3* and for two genes with variable XCI escape, *PLS3* and *TMEM187* (**Fig. S3B**, **Excel Table S4-2**).

Top Cell Type DEG also included five imprinted genes including *SNRPN* at chr15q11.2 and *MEG3* at chr14q32.2 (**Fig. 3**, **Excel Table S2-1**, **S2-2**) [87]. Of 47 genes found as a top ten underexpressed threshold DEG, we found four (9%) paternally expressed imprinted genes: *SNRPN*, *PWAR6*, *NDN*, and *PEG10* with *SNRPN* shared in nine subtypes (**Excel Table S2-2**). Of 71 genes found as a top ten overexpressed threshold DEG, we found one (1%) maternally expressed imprinted gene *MEG3* in two subtypes: D00 PP-A1A2 and D19 EPDC (**Excel Table S2-2**). Across all 14 subtypes from Days 0 to 19, PP cells had highest *SNRPN* expression that then decreased in other shared cell types (**Fig. S11D**). Like *XIST* expression, for all time points and shared cell types, we detected lower expression levels of *SNRPN* in Patient compared to Control cells except for CM/UNK cells (**Fig. 3**). Likewise, for two other imprinted genes at chr15q11.2, we found *PWAR6* as an underexpressed DEG in 13 of 14 subtypes, a threshold DEG in six subtypes, and a top ten DEG in four cell subtypes, and *NDN* as an underexpressed DEG in 11 of 14 subtypes, a threshold DEG in seven subtypes, and a top ten DEG in four cell subtypes (**Excel Table S2-2**). For *PEG10* at 7q21, we found variable results across 14 subtypes with underexpression in 7 and overexpression in 4 subtypes and identification as a top ten threshold underexpressed DEG in only D16 CM-A1A2 (**Excel Table S2-2**). For *MEG3*, despite limited expression at all time points (**Fig. 3**, **Fig. S11D**), we found *MEG3* as a top ten overexpressed threshold DEG in two subtypes: D00 PP-A1A2 and D19 EPDC (**Excel Table S2-2**). For these imprinted genes, we could not determine allelic expression due to the lack of informative coding SNV.

### Cell type-specific pathway enrichment: dysregulation of gene expression and development of Cardiac Progenitors and Cardiomyocytes (Workflow Step-IIIA)

Using enrichment analysis of top Cell Type DEG between Patient and Control cells (**Table**, **Fig. S11AC**), we found evidence for dysregulation of gene expression and developmental pathways by ORA (**Excel Table S2-3**) and GSEA (**Excel Table S2-4**).

For most shared cell types (**Table**, **Fig. S11AC**), our results by ORA supported differences in gene regulation (**Excel Table S2-3**) involving dosage compensation, epigenetic regulation, and heterochromatin formation attributed to underexpression of three DEG: *LMNA*, *XIST*, and *NDN* in Patient cells. Specifically, D00 PP-B and D09B EPDC enriched for dosage compensation by inactivation of X chromosome (GO:0009048, GO:0007549) as expected, attributed to *XIST* underexpression. Similarly, D09B CP-B and D19 EPDC enriched for epigenetic regulation of gene expression (GO:0040029) attributed to underexpression of up to three DEG: *XIST*, *NDN*, and *LMNA*. For D19 EPDC, underexpression of *NDN* and *LMNA* also resulted in enrichment for heterochromatin formation (GO:0031507, GO:0070828).

Using enrichment analysis of top DEG, we also found evidence for differences in developmental pathways between Patient and Control CP and CM (**Table**, **Fig. S11AC**) attributed to DEG subsets encoding transcription factors, signaling proteins, and structural components of cardiac muscle cells and sarcomeres [87, 89]. First, we found differences in CP subtypes at Day 9 by ORA (**Excel Table S2-3**). D09B CP-A enriched for ventricular cardiac muscle cell development or differentiation (GO:0055015, GO:0055012) attributed to underexpression of two DEG, our DCM-study gene *LMNA* and *NKX2-5*. In contrast, for D09B CP-B, overexpression of two DEG, *KRT19*, encoding a keratin intermediate filament protein, and *MYL9*, encoding a myosin light chain, resulted in enrichment for myofibril assembly (GO:0030239) and striated muscle cell development (GO:0055002).

For CM at Days 16 and 19, our results also supported differences in developmental and differentiation pathways (**Table**, **Fig. 4**, **Fig. S11AC**). For D16 CM-A1A2, among top enriched gene sets by ORA (**Excel Table S2-3**), we found muscle organ development (GO:0007517) attributed to underexpression of 13 DEG including *NKX2-5* and *WNT2*, encoding a highly expressed WNT signaling protein during cardiac differentiation [90] and cardiac muscle contraction (GO:0060048, GO:0006941) and actin filament-based movement (GO:0030048) attributed to the underexpression of 8 to 11 DEG including *CACNA1D* and *CACNA1G*, encoding voltage-sensitive calcium channels [87]. For D16 CM-A1A2, we also found muscle cell development and differentiation (GO:0055001, GO:0042692, GO:0051146) by ORA attributed to overexpression of 23 to 31 genes including four sarcomere protein encoding genes *ACTC1*, *ACTN2*, *CSRP3*, and *MYH7* and Hallmark EMT by GSEA (NES= 1.497658595, FDR= 0.0074101) (**Excel Table S2-4**) attributed to upregulation of 23 DEG including known EMT genes: *CDH2* and XL *TIMP1* [91]. For all CM-A subtypes, Patient cells had higher median module score for EMT compared to Control cells at Day 16 (**Fig. 4A**, **Fig. S11E**).

**Figure 4.**
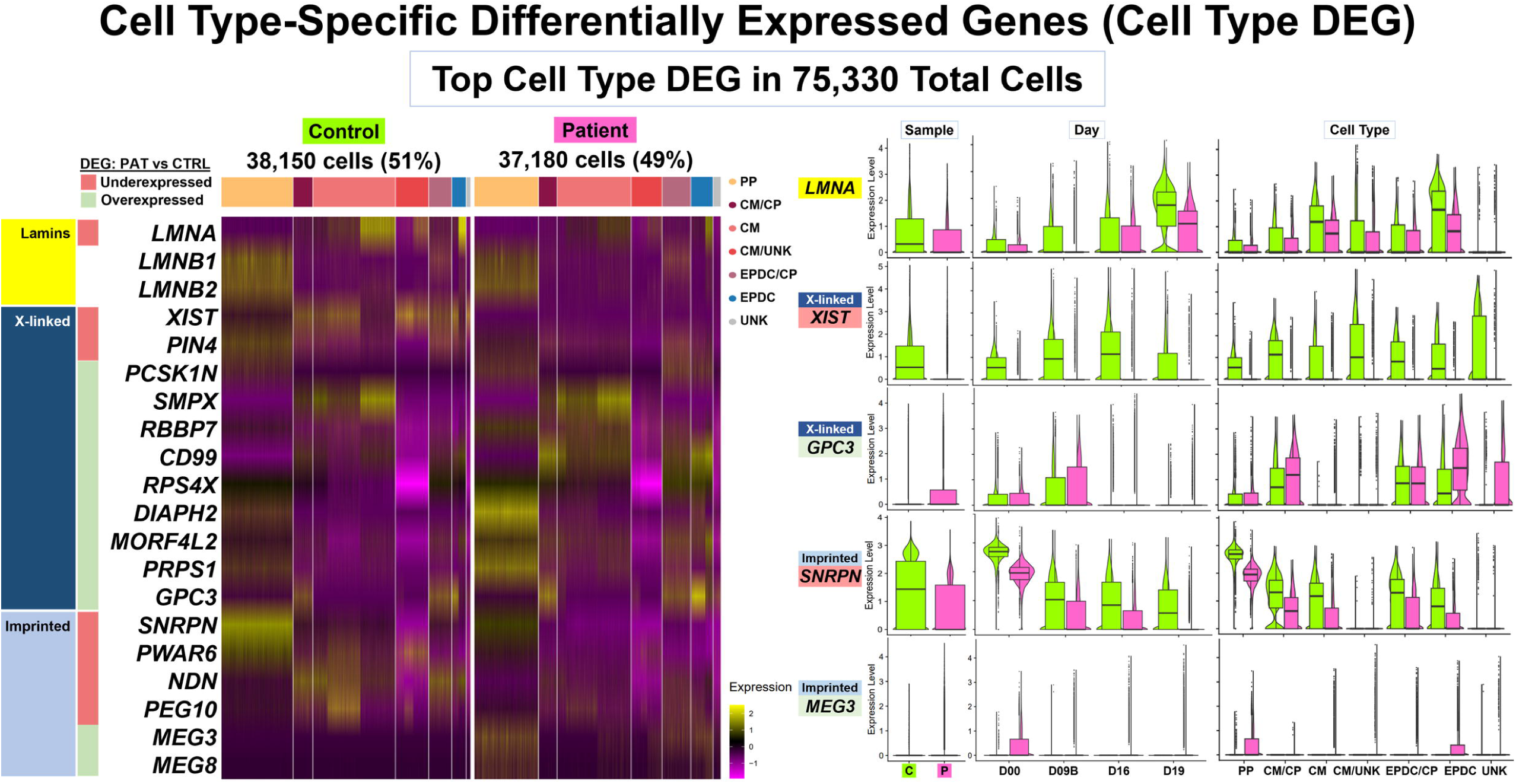
Cell Type-Specific Differentially Expressed Genes (Cell Type DEG): Enrichment. Our Comparative Analyses (Patient vs. Control cells) included cell type-specific differential expression and enrichment using Integrated Data at four time points: D00, D09B, D16, and D19. We identified Cell Type DEG in 14 pairs of ‘balanced’ cell types/subtypes with 71,541 total cells. (**A**) Day 16 Enrichment. Shown are GSEA [75] results using Cluster Profiler [77] (Left Panel) for 8,639 CM-A1A2 cells (Top) and 3,133 EPDC (Bottom) with numbers of DEG and enriched MSigDB Hallmark gene sets, dot plots with normalized enrichment score (NES) and adjusted p-values, and GSEA enrichment plots. We used Module Scoring in Seurat (Right Panel) to compare expression patterns of key DEG in enriched gene sets for Subset-A Data (12,700 CM) and Subset-B Data (3,133 EPDC). For each gene set, shown are UMAP plots with Cell Subtype and Module Score and violin plots with boxplot by Cell Subtype, Condition, and Cell Subtype-Condition. Overall, these results at Day 16 support dysregulation of TGF Beta signaling, EMT developmental pathway, oxidative phosphorylation and glycolysis metabolism, and cell proliferation (via MYC target genes) in Patient CM and mTORC1 signaling in Patient EPDC. (**B**) Day 19 Enrichment. Shown are Cell Type DEG results using Seurat and EnhancedVolcano [73] (Top Panel) and ORA [72] results using Cluster Profiler [77] (Bottom Panel) for 9,978 CM-A1 cells (Left) and 2,256 EPDC (Right) with UMAP plot of cell subtypes, number of Threshold DEG and volcano plot [73] with top ten DEG labeled, number of under/over-expressed DEG and enriched GO biological processes gene sets, dot plots with Gene Ratio and adjusted p-values, and Enrichment Maps. Overall, these results at Day 19 support dysregulation of cardiogenesis with both underexpressed DEG including *LMNA* and overexpressed DEG including sarcomere genes in Patient CM and epigenetic/heterochromatin dysregulation with underexpression of *XIST*, *NDN*, and *LMNA* in Patient EPDC.

For D19 CM-A1 (**Fig. 4B**), among top enriched gene sets, we found striated muscle cell differentiation (GO:0051146) and myofibril assembly/sarcomere organization (GO:0030239, GO:0045214) by ORA (**Excel Table S2-3**) attributed to underexpression of 7 to 15 DEG including *LMNA* and two genes *SYNPO2L* and *MYOZ2* encoding actin binding proteins. For D19 CM-A1, we also found cardiac muscle and heart contraction (GO:0060048, GO:0006941, GO:0003015, GO:0060047, GO:0006936) by ORA attributed to the overexpression of 14 to 21 DEG including five sarcomere genes: *MYBPC3*, *MYL2*, *MYL3*, *TCAP*, and *TNNI3* and Hallmark Myogenesis by GSEA (NES= 1.59608061, FDR= 0.007430091) (**Excel Table S2-4**) attributed to upregulation of 37 DEG including the XL *DMD* Dystrophin gene and seven sarcomere protein encoding genes: *LDB3*, *MYBPC3*, *MYL2*, *MYL3*, *TCAP*, *TNNC1*, and *TNNT2*. For all CM-A subtypes, Patient cells had a higher median module score for Myogenesis compared to Control cells at Day 19 (**Fig. S11E**).

### Cell type-specific pathway enrichment: dysregulation of cell signaling, metabolism, and proliferation (Workflow Step-IIIA)

Using enrichment analysis of top Cell Type DEG between Patient and Control cells (**Table**, **Fig. S11AC**), we also found evidence for dysregulation of signaling, metabolism, and cell cycle control by ORA (**Excel Table S2-3**) and GSEA (**Excel Table S2-4**).

For cell signaling, our results included enriched pathways involving TGF Beta signaling for CM and mTORC1 signaling for EPDC at Day 16 (**Fig. 4A**) attributed to DEG subsets encoding secreted proteins, regulatory factors, and downstream targets [87]. Among top enriched gene sets in D16 CM-A1A2, we found transmembrane receptor serine/threonine kinase signaling pathway (GO:0007178) by ORA (**Excel Table S2-3**) and Hallmark TGF Beta signaling by GSEA (NES= -1.60, FDR= 0.039) (**Excel Table S2-4**) attributed to underexpression or downregulation of 7 to 13 DEG including *SMAD6*, *ID1*, *ID2*, and *ID3* encoding negative regulatory factors [87]. For CM-A1 and A2, Patient cells had a lower median module score for TGF Beta signaling compared to Control cells at Day 16 (**Fig. 4A**, **Fig. S11E**). In contrast, D16 EPDC enriched for Hallmark Mtorc1 signaling by GSEA (NES= 1.35, FDR= 0.011) attributed to upregulation of 43 DEG including mTORC1 regulation gene *DDIT4* (*REDD1*) and metabolic genes: *HK2*, *PGK1*, *ENO1*, and *GAPDH* for glucose, *LDHA* for lactate, and *PHGDH* and *SHMT2* for amino acid metabolism. For EPDC-A1, A2, and B, Patient cells had a higher median module score for Mtorc1 signaling compared to Control cells at Day 16 (**Fig. 4A**, **Fig. S11E**).

Our enrichment results also supported dysregulation in cell metabolism involving oxidative phosphorylation, glucose, and nucleotides for different cell types and time points (**Table**, **Fig. S11AC**). At Days 0 and 9, our results supported decreased oxidative phosphorylation for Patient PP and CP and increased glycolysis for Patient EPDC (**Fig. S11C**) compared to Control cells attributed to DEG subsets encoding key enzymes, components, and regulators of the electron transport chain (ETC) and glycolysis [87]. For D00 PP-A1A2, we found mitochondrial oxidative metabolism (GO:0019646, GO:0042773, GO:0042775, GO:0006120, GO:0022904) among top enriched gene sets by ORA (**Excel Table S2-3**) attributed to three underexpressed DEG: *DNAJC15*, *NDUFA3*, and *NDUFA6*. Similarly, D09B CP-A enriched for Hallmark oxidative metabolism by GSEA (NES= -1.74, FDR= 0.00016) (**Excel Table S2-4)** attributed to downregulation of 46 DEG including multiple ETC encoding genes. For D09B CP-A, Patient cells had a lower median module score for oxidative metabolism compared to Control cells (**Fig. S11E**). In contrast, for D09B EPDC, we found glycolytic metabolism (GO:0061621, GO:0061718, GO:0006735) by ORA attributed to overexpression of *PFKP* encoding phosphofructokinase in the first committing step of glycolysis.

At Day 16, our results for CM supported (**Fig. 4A**) metabolic differences consistent with increased glucose metabolism and oxidative phosphorylation for Patient compared to Control CM. For D16 CM-A1A2 using GSEA (**Excel Table S2-4)**, we found Hallmark glycolysis (NES= 1.45, FDR= 0.020) attributed to upregulation of 27 DEG including *HK2* and *ENO2* encoding enzymes for glucose metabolism and Hallmark oxidative metabolism (NES= 1.57, FDR= 3.86E-05) attributed to upregulation of 72 DEG including multiple ETC encoding genes and three DEG overlapping glycolysis. For all CM-A subtypes, Patient cells had a higher median module score for both glycolysis and oxidative metabolism compared to Control cells at Day 16 (**Fig. 4A**, **Fig. S11E**). Likewise, for D16 CM-A1A2 by ORA (**Excel Table S2-3**), we found mitochondrial oxidative metabolism (GO:0019646, GO:0042773) attributed to 16 overexpressed ETC encoding genes including *COX6A2*, *COX7A1*, and *CYCS*, not found by GSEA.

In contrast to CM at Day 16, our results for EPDC supported increased glucose and nucleotide metabolism for Patient compared to Control EPDC consistent with metabolic selection by lactate enrichment [45]. For D16 EPDC (**Fig. S11C**), we found purine ribonucleotide metabolism (GO:0009150, GO:0006163, GO:0072521, GO:0019693,) and generation of precursor metabolites and energy (GO:0006091) by ORA (**Excel Table S2-3**) attributed to upregulation of 11 to 12 DEG including *PFKP*, *TPI1*, *PKM*, and *GAPDH* encoding enzymes for glycolysis and gluconeogenesis and *TKT*, *HINT1*, and *PAICS* encoding enzymes for nucleotide metabolism.

In addition, for CM at Day 16, we found differences in cell proliferation (**Fig. 4A**). D16 CM-A1A2 enriched for Hallmark Myc targets V1 by GSEA (NES= 1.47, FDR= 0.004) (**Excel Table S2-4**) attributed to upregulation of 36 DEG encoding targets of the MYC transcription factor that promote cell growth and proliferation including ten spliceosome genes: *HNRNPA2B1*, *HNRNPA3*, *HNRNPD*, *HNRNPR*, *LSM7*, *SNRPD2*, *SNRPG*, *SRSF2*, *SRSF3*, *TXNL4A* [78, 87]. For CM-A1 and A2, Patient cells had a higher median module score for Myc targets V1 compared to Control cells at Day 16 (**Fig. 4A**, **Fig. S11E**).

### Single lineage for Pluripotent cells and lineage bifurcation for Cardiac Progenitors to Cardiomyocytes and EPDC in Patient and Control Data (Workflow Step-IIIB)

For Single Subset Data (n=7: 22,326 total cells) for selected cell subtypes of one condition (**Table S10**) and for Integrated Subset Data (n=2: 62,488 total cells) of 18 paired subtypes (**Fig. S8A**, **Table S11**), we conducted trajectory inference, testing for Lineage DEG, and enrichment analyses (**Table**, **Fig. 5 - Fig. 7**, **Fig. S12**, **Fig. S13, Excel Table S3**). Results for Control Single Subset Data included three single trajectories (PP-A to PP-B, CMESO to CP, CP-A to CM-A) and one bifurcating trajectory (ME to CMESO and ENDO) (**Fig. S12A)**. Results for Integrated Subset Data consisted of two parts: D00 PP cells (19,346 total cells) with a single trajectory from PP-A to PP-B (PP Lineage) (**Fig. 5**, **Fig. S13BC**) and D09B, D16, and D19 CP, CM, and EPDC cells (43,142 total cells) with a bifurcating trajectory starting at CP-1 and ending at CM-2 (CP Lineage-1) and EPDC (CP Lineage-2) (**Fig. 6 – Fig. 7**, **Fig. S13BD**). Between conditions, we found differences in topology and progression for all three lineages (PP and CP lineages) and in rate of differentiation between CP Lineage-1 for CM and CP Lineage-2 for EPDC (**Fig. S13A**).

**Figure 5.**
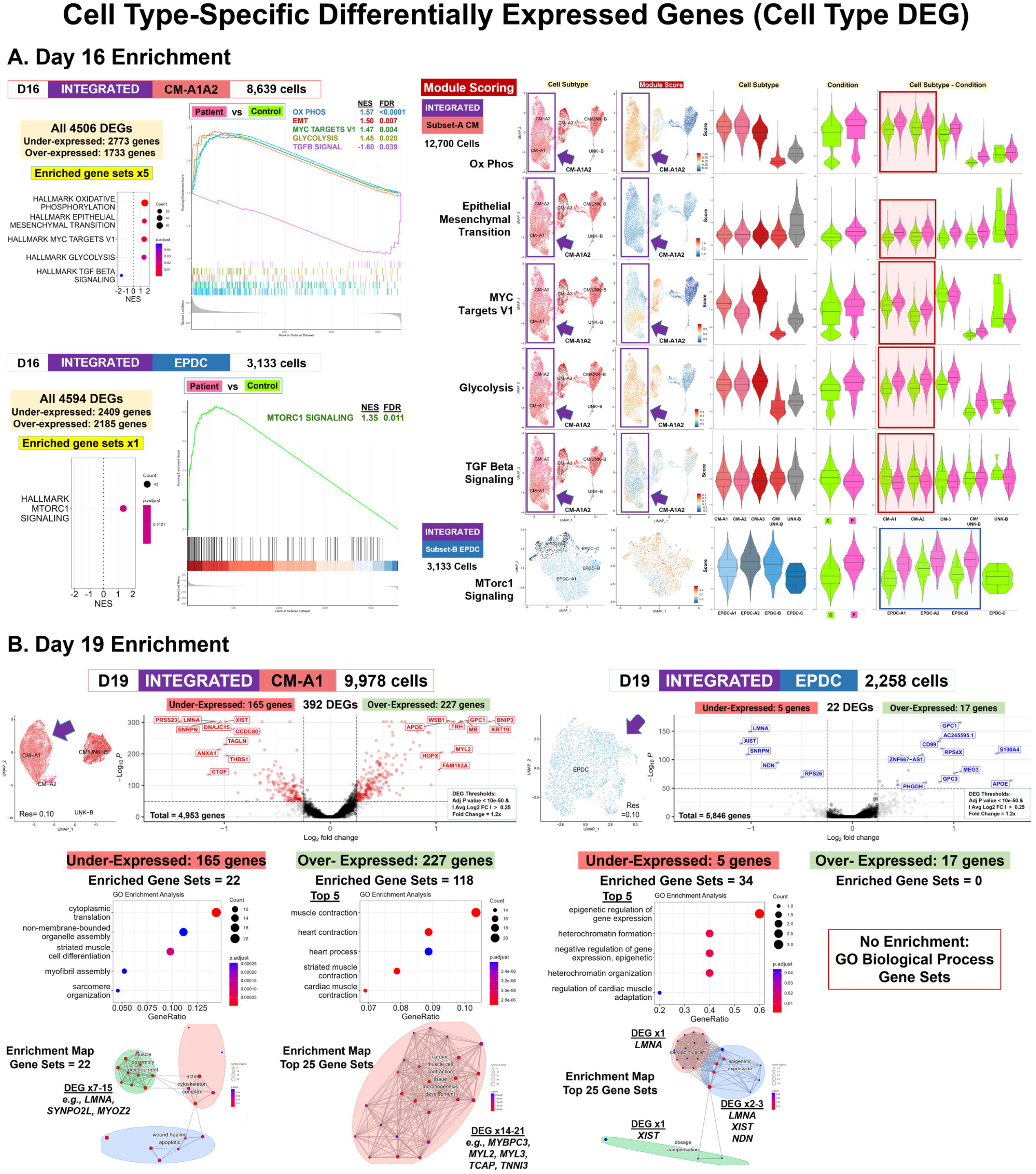
Lineage-Specific Differentially Expressed Genes (Lineage DEG): Pluripotent Lineage-Top DEG and Enrichment. Our Comparative Analyses (Patient vs. Control cells) included lineage-specific differential expression and enrichment using Integrated Data of 18 shared subtypes (62,488 total cells). This included D00 Pluripotent (PP) subtypes for the PP Lineage (19,346 total cells), a single trajectory from PP-A to PP-B. Shown are UMAP plots with trajectory and principal curves using Slingshot [79] (Left Panel), heat map by condition along pseudotime points using pheatmap and Condiments [71] (Middle Panel), and Smoother plots and Gene Count plots using TradeSeq [80] (Right Panel). (**A**) PP Lineage: Top DEG in 19,346 Total Cells. We identified 97 Conditional Test DEG that clustered into two Groups: A (74 genes) and B (23 genes). Top DEG included *XIST* (#1) underexpressed and imprinted gene *MEG8* overexpressed in Patient PP along all pseudotime points. *LMNA* was not found as a Lineage DEG; however, *LMNA* expression was lower in Patient compared to Control PP at all pseudotime points. (**B**) PP Lineage: Enrichment. Shown are ORA enrichment results using Cluster Profiler [77] (Left Panel) with numbers of DEG by Group and enriched GO BP gene sets (GS), Top GS, key DEG, and Enrichment Maps. To compare expression of key DEG in enriched GS, we used Module Scoring in Seurat (Right Panel) and violin plots with boxplot. These results support dysregulation of Metal Homeostasis with overexpression of 5 to 11 genes from DEG Group-A including four metallothionein proteins (*MT*) genes and RNA polymerase II transcription with underexpression of 12 DEG from DEG Group-B including 10 Zinc-finger protein (*ZNF)* genes in Patient PP.

**Figure 6.**
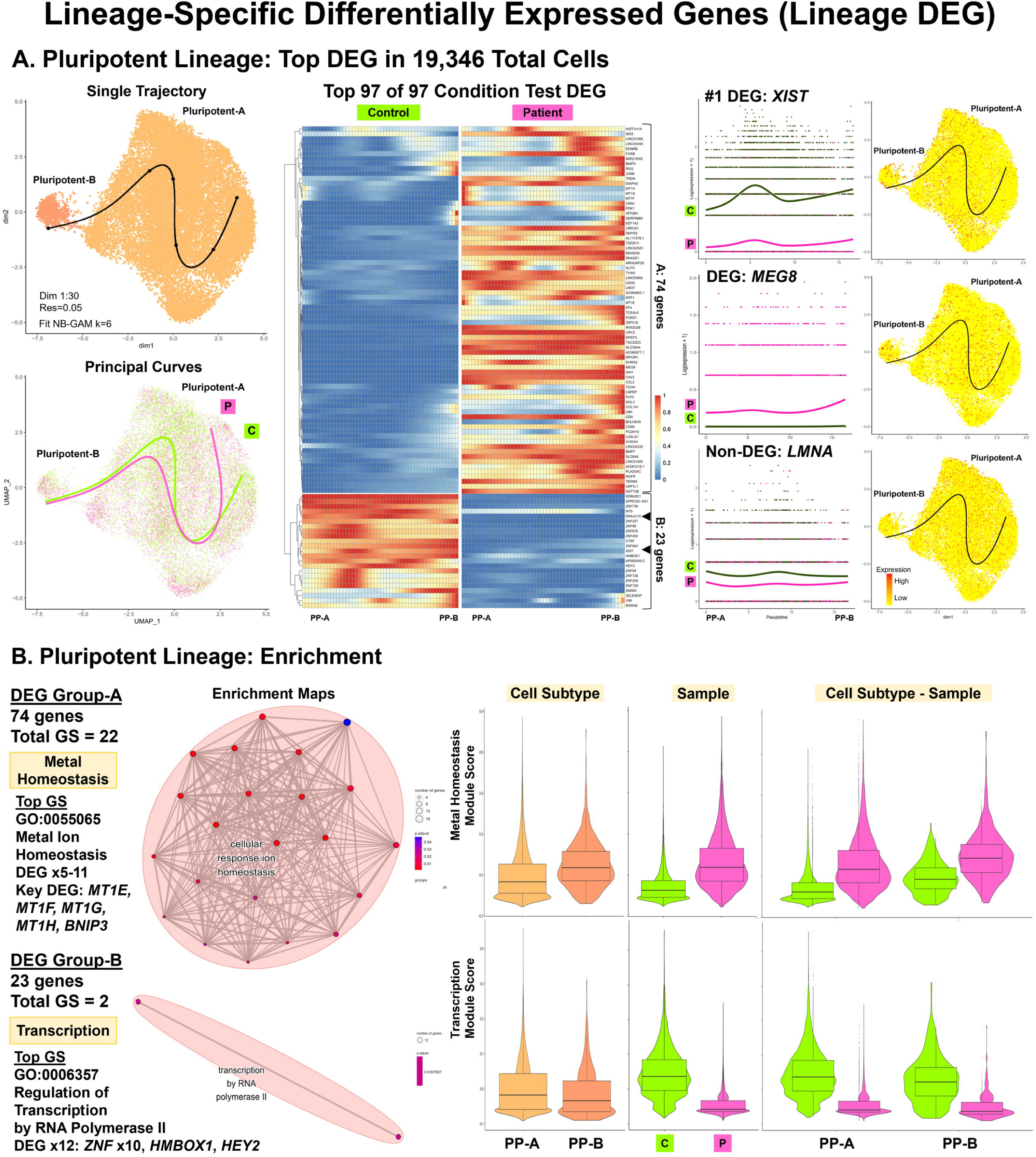
Lineage-Specific Differentially Expressed Genes (Lineage DEG): Cardiac Progenitor Lineages-Top DEG. Our Comparative Analyses (Patient vs. Control cells) included lineage-specific differential expression and enrichment using Integrated Data of 18 shared subtypes (62,488 total cells). This included D09B, D16, and D19 Cardiac Progenitor (CP), CM, and EPDC subtypes for the CP Lineages (43,142 total cells), a bifurcating trajectory starting at CP-1 and ending at CM-2 (Lineage-1: CP to CM) and EPDC (Lineage-2: CP to EPDC). Shown are UMAP plots with trajectory and principal curves using Slingshot [79] (Left Panel), heat map by condition along pseudotime points using pheatmap and Condiments [71] (Middle Panel), and Smoother plots and Gene Count plots using TradeSeq [80] (Right Panel). (**A**) CP Lineage-1: CP to CM-Top DEG. We identified 391 total Conditional Test DEG with Top 100 DEG clustered into four Groups: A (10 genes), B (10 genes), C (59 genes), and D (21 genes). Top DEG included *XIST* (#1) underexpressed and *GPC1* (#2) overexpressed in Patient cells along all pseudotime points. (**B**) CP Lineage-2: CP to EPDC-Top DEG. We identified 25 total Conditional Test DEG that clustered into three Groups: A (19 genes), B (3 genes), and C (3 genes). Top DEG included XL *DIAPH2* overexpressed and *ZNF208* underexpressed in Patient cells along all pseudotime points and cell cycle gene *CCNB2* overexpressed in Patient CP cells. Like the PP lineage, *LMNA* was not as a DEG in both CP lineages with lower expression in Patient compared to Control cells at all pseudotime points.

**Figure 7.**
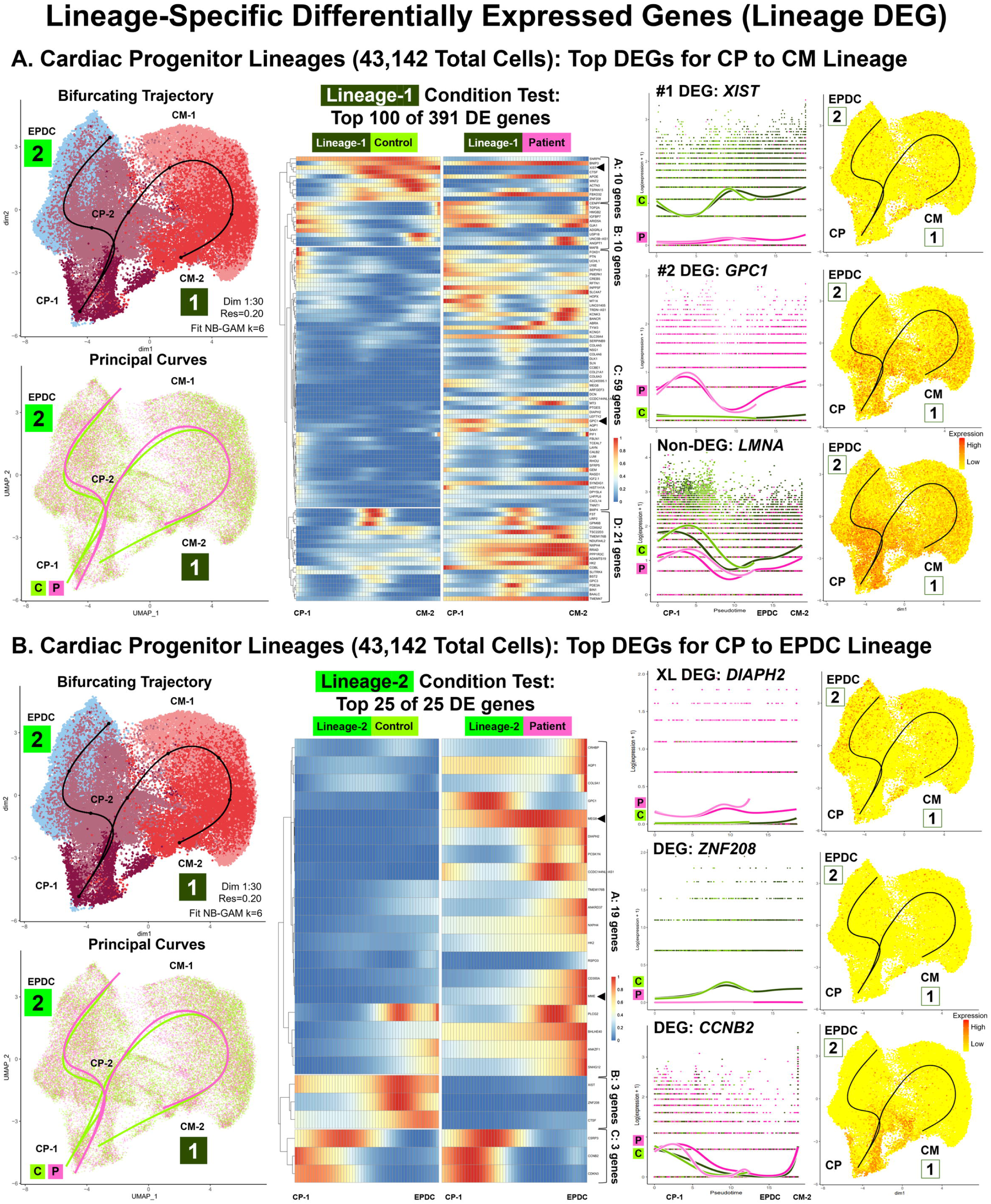
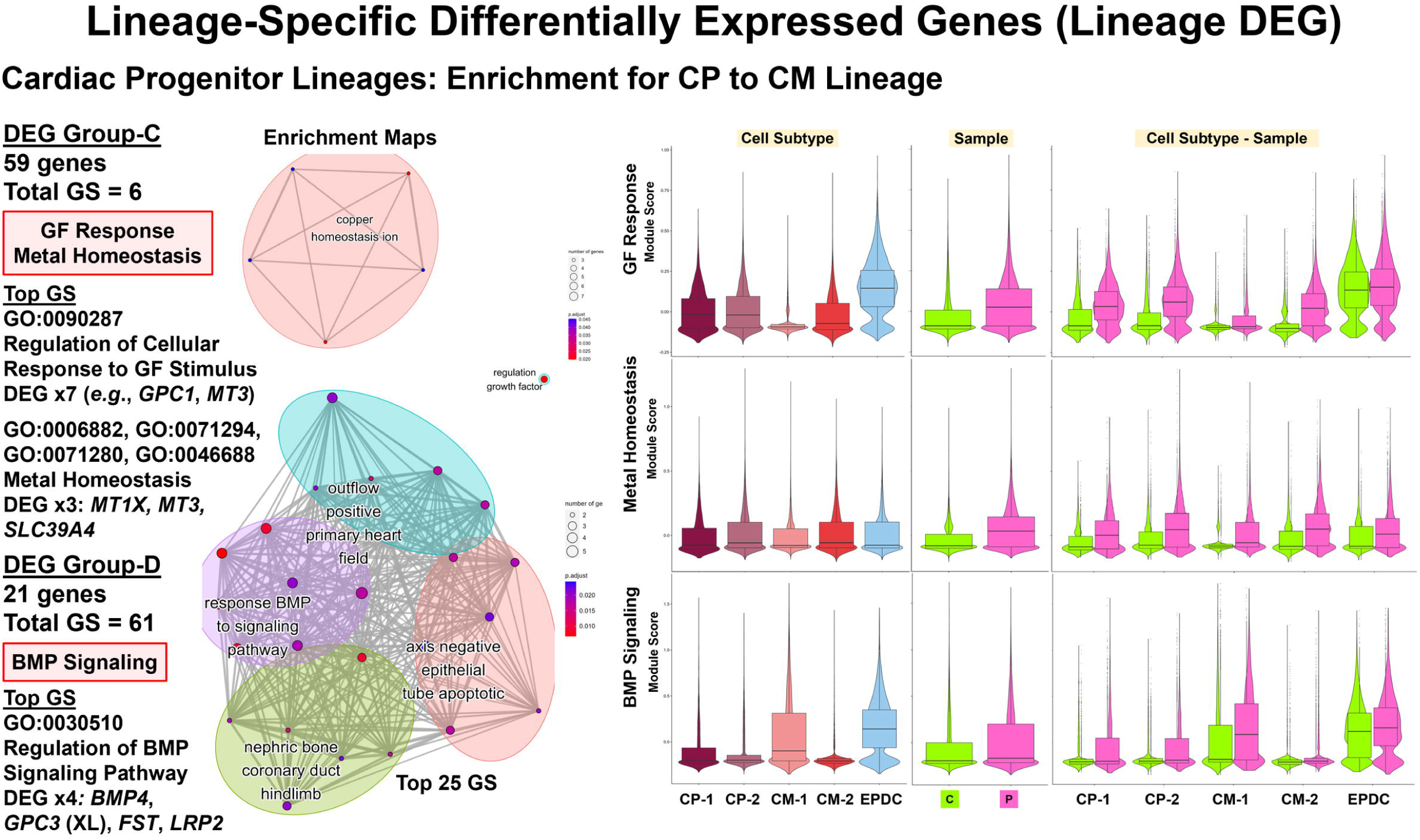
Lineage-Specific Differentially Expressed Genes (Lineage DEG): Cardiac Progenitor Lineages-Enrichment. For Lineage-1: CP to CM, shown are ORA enrichment results using Cluster Profiler [77] (Left Panel) with numbers of DEG by Group and enriched GO BP gene sets (GS), Top GS, key DEG, and Enrichment Maps. To compare expression of key DEG in enriched GS, we used Module Scoring in Seurat (Right Panel) and violin plots with boxplot. For Lineage-1: CP to CM, these results support dysregulation of Growth Factor (GF) Response with overexpression of seven Group-C DEG including *GPC1* and *MT3*, Metal Homeostasis with overexpression of three Group-C DEG including two metallothionein proteins (*MT1X* and *MT3*) genes, and BMP Signaling Pathway with overexpression of four Group-D DEG: *BMP4*, XL *GPC3*, *FST*, and *LRP2*.

### Lineage-specific differential expression: non-coding RNA *XIST*, sixth imprinted gene *MEG8*, and cell-surface glycoprotein encoding gene *GPC1* (Workflow Step-IIIB)

Across the three lineages, we found equal proportions of Patient (50%) and Control (50%) cells (**Table S11**) and variable numbers of Lineage DEG between conditions (**Fig. S13A**). For example, we identified fewer total DEG between conditions (CT Lineage DEG) for the PP lineage (924 DEG) compared to the two CP lineages (2,335 Global DEG). Likewise, between the two CP lineages, we found fewer total DEG between conditions (CT Lineage DEG) for CP Lineage-1 to CM (1,795 DEG) compared to CP Lineage-2 to EPDC (2,216 DEG).

For all three lineages, consistent with our Cell Type DEG results, we found the *XIST* non-coding RNA gene as the highest ranked or among top 25 CT Lineage DEG (**Excel Table S3-2**). For all three lineages, we found underexpression of *XIST* in Patient compared to Control PP cells along all pseudotime points (**Fig. 5A**, **Fig. 6A**). In addition, we found a sixth imprinted gene *MEG8* at 14q32.2 differentially expressed between Patient and Control cells [87]. For the PP lineage, we found *MEG8* among top CT Lineage DEG and for the CP to EPDC lineage, as the highest ranked CT Lineage DEG (**Excel Table S3-2**) with overexpression of *MEG8* in Patient compared to Control PP cells along all pseudotime points (**Fig. 5A**). For both CP lineages (**Excel Table S3-3**), top CT Lineage DEG included *GPC1* encoding Glypican Proteoglycan 1, a cell-surface glycoprotein for growth regulation [87], with overexpression of *GPC1* in Patient compared to Control cells along all pseudotime points (**Fig. 6A**). For all three lineages, *LMNA* expression was lower in Patient compared to Control cells at most pseudotime points (**Fig. 5A**, **Fig. 6A**); however, top CT Lineage DEG did not include *LMNA*.

### Lineage-specific pathway enrichment: regulation of gene expression, cell signaling, proliferation, and homeostasis (Workflow Step-III: Lineage Enrichment)

Using enrichment analysis of top Lineage DEG between Patient and Control cells (**Table**, **Fig. 5 – Fig. 7**, **Fig. S13A**), we found evidence for dysregulation of pathways consistent with our analyses of top Cell Type DEG. For the PP lineage, we found four enriched gene sets compared to none for CP lineages by ORA and GSEA. For the CP lineages, we found 19 enriched gene sets for Lineage-1 for CM compared to none for Lineage-2 for EPDC by ORA and GSEA. Furthermore, we found additional enriched gene sets after clustering top Lineage DEG into smaller groups by hierarchical cluster analysis.

Consistent with our Cell Type DEG results, we also found evidence for differences in gene regulation between Patient and Control cells for two lineages: PP lineage and CP to EPDC lineage (**Table**, **Excel Table S3-4**). The PP lineage (**Fig. 5B**, **Fig. S13C**) enriched for regulation of transcription by RNA polymerase II (GO:0006357, GO:0006366) using CT Lineage DEG Group-B attributed to underexpression of 12 genes encoding DNA-binding proteins: 10 Zinc-finger proteins (ZNFs), *HMBOX1*, and *HEY2* [87]. For PP-A and PP-B, Patient cells had a lower median module score based on expression of the 12 genes compared to Control cells (**Fig. 5B**). For the CP to EPDC lineage, Top DEG included *ZNF208* underexpressed in Patient cells along all pseudotime points (**Fig. 6B**). For the CP to EPDC lineage, we also found enrichment for dosage compensation (GO:0007549, GO:0009048) using CT Lineage-2 DEG Group-B attributed to *XIST* underexpression (**Fig. S13D**).

In each CP lineage, we also found evidence for differences in cell signaling and proliferation between Patient and Control cells by ORA (**Table**, **Excel Table S3-4**). The CP to CM lineage (**Fig. 7**, **Fig. S13D**) enriched for regulation of BMP signaling pathway (GO:0030510, GO:0030509, GO:0071773, GO:0071772) using CT Lineage-1 DEG Group-D attributed to overexpression of four DEG: *BMP4*, XL *GPC3*, *FST*, and *LRP2* encoding four regulatory proteins [87]. For CM-1, Patient cells had a greater median module score based on *BMP4*, *GPC3*, *FST* and *LRP2* expression compared to Control cells (**Fig. 7**) consistent with our Cell Type DEG results for TGF Beta signaling in D16 CM-A1A2 (**Fig. 4A**). The CP to CM lineage also enriched for regulation of cellular response to growth factor stimulus (GO:0090287) using CT Lineage-1 DEG Group-C attributed to overexpression of seven DEG such as *GPC1* and *MT3*, encoding Metallothionein 3, a growth inhibitory factor [87]. For CP-1, CP-2, and CM-2 cells, Patient compared to Control cells had a greater median module score based on the expression of these seven DEG (**Fig. S13D**). For the CP to EPDC lineage (**Fig. S13C**), we found enrichment for regulation of cyclin-dependent protein serine/threonine kinase activity (GO:0000079, GO:0071900, GO:1904029) using CT Lineage-2 DEG Group-C attributed to overexpression, primarily in CP cells, of two DEG: *CDKN3* and *CCNB2* encoding Cyclin B2 in TGF Beta-mediated cell cycle control [87].

Finally, for two lineages: PP lineage and CP lineage-1, our results included evidence for differences in cell homeostasis between Patient and Control cells by ORA (**Table**, **Excel Table S3-4**). The PP lineage (**Fig. 5B**, **Fig. S13CD**), enriched for zinc/metal ion homeostasis (GO:0055065, GO:0055069, GO:0072507, GO:0006875, GO:0072503) using CT Lineage DEG Group-A attributed to overexpression of 5 to 11 DEG including *MT1E*, *MT1F*, *MT1G*, *and MT1H*, encoding four metallothionein proteins that bind heavy metals, and *BNIP3*, encoding BCL2 Interacting Protein 3, a pro-apoptotic factor [87]. For PP-A and PP-B, Patient compared to Control cells had a higher median module score based on the expression of the 11 genes (**Fig. 5B**). Likewise, the CP to CM lineage (**Fig. 7**, **Fig. S13D**) enriched for metal homeostasis and response to zinc ion (GO:0006882, GO:0071294) and copper ion (GO:0071280, GO:0046688) using CT Lineage-1 DEG Group-C attributed to overexpression of three DEG such as *MT1X* and *MT3* also encoding metallothionein proteins [87]. Like the PP lineage, Patient compared to Control cells had a higher median module score based on the expression of the five genes (**Fig. 7**).

### Decreased Lamin A/C protein levels for Patient cells at Day 19

To compare Lamin A/C protein levels, we differentiated three iPSC pairs as biological replicates (BR): three Patient-derived iPSC (PA1, PA2, and PA3) and three Control iPSC (CA1-B, U2, and CA3), obtained protein lysates from differentiated cells at Day 19, repeated Western Blot three times (TR blots x3) (**Fig. 8A**, **Fig. S14**), and quantified protein levels for Lamin A, Lamin C, and total Lamin A+C as Normalized Ratios (NR) (**Table S12A**). Comparison of the Control (n=3: CA1-B, U2, CA3) vs. Patient (n=3: PA1, PA2, PA3) data showed a decrease in mean protein levels for Lamin A, Lamin C, and total Lamin A+C in the Patient compared to Control cells (**Fig. 8B**, **Table S12B**); however, these differences were not statistically significant (Lamin A+C: t = 1.66, p = 0.172, Lamin A: t = 1.59, p = 0.187, Lamin C: t = 1.68, p = 0.168). In our pairwise comparisons, statistical analysis of fold change revealed a significant decrease in protein level for Lamin A+C for PA1 (-81%) compared to CA1 (t = 27.1, p < 0.0001) and PA2 (-51%) compared to U2 (t = 10.2, p < 0.001) but no significant difference for PA3 (-0.34%) compared to CA3 (t = 0.028, p = 0.98) (**Fig. 8C**, **Table S12C**). Our analyses also showed similar, acceptable CV for all the pairs (Mean CV = 21.6% ± 4.5%) and % changes twice as large CV for PA1 and PA2 but not for PA3 (**Table S12C**).

**Figure 8.**
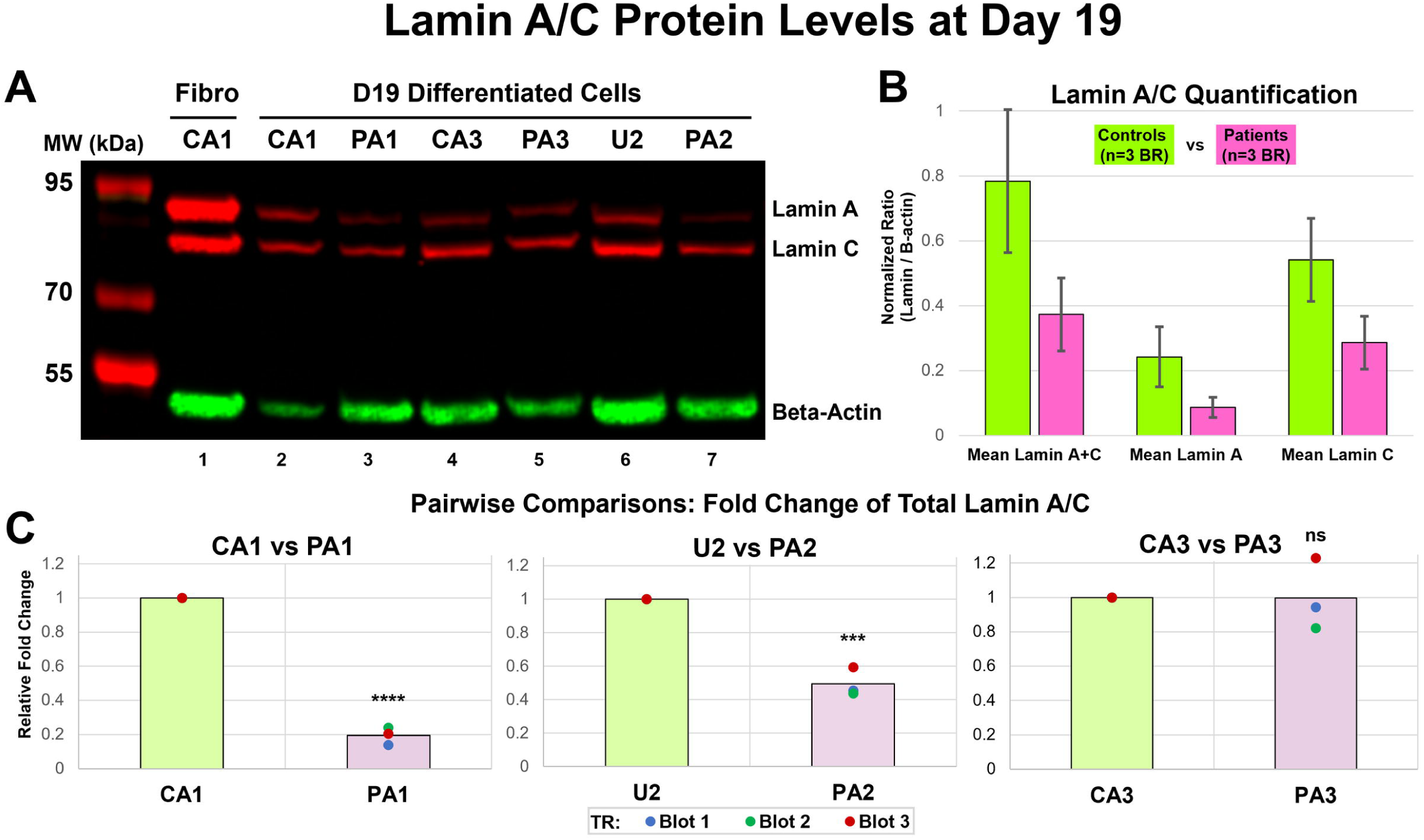
Lamin A/C Protein Levels at Day 19. (**A**) Western blot (WB). We performed immunoblotting on lysates from differentiated cells with three biological replicates (BR) at Day 19 for Patient (PA1, PA2, and PA3) and Control (CA1-B, U2, and CA3) iPSC lines. Image (Blot 3) shown is taken using the Azure c600 imaging system with fluorescent detection in red for Lamin A (74 kDa) and Lamin C (62 kDa) and in green for Beta-Actin (42 kDa) from Control fibroblast (CA1) (lane 1, positive control) and the three age and sex-matched pairs: CA1 and PA1 (lane 2 and 3), CA3 and PA3 (lane 4 and 5), and U2 and PA2 (lane 6 and 7). Positions and sizes of molecular weight (MW) marker proteins are given on the left margin. (**B**) Lamin A/C Quantification: Controls vs. Patients. We measured the band volume (BV) for Lamin A, Lamin C, and Beta-Actin using AzureSpot software and calculated normalized ratio (NR= Lamin BV/Beta-Actin BV) of each sample (CA1-B, PA1, U2, PA2, CA3, and PA3) for three WB experiments (Blot 1-3) as technical replicates (TR). Comparison of the Control (n=3 BR: CA1-B, U2, CA3) vs. Patient (n=3 BR: PA1, PA2, PA3) data showed decrease in mean protein levels for Lamin A, Lamin C, Lamin A+C in the Patient compared to Control cells; these differences were not statistically significant (p > 0.05). Error bars denote SD of the mean protein levels of BR. (**C**) Pairwise Comparisons: Fold Change of Total Lamin A/C. For each matched pair, we calculated fold change for Lamin A+C by dividing Control and Patient protein levels each with Control protein level. Each chart shows Fold Change from three TR (Blot 1-3) as individual data points and mean Fold Change shown as column. Statistical analysis of the Fold Change revealed a significant decreases in protein level for Lamin A+C of PA1 (****p<0.0001) and PA2 (***p<0.001) but no significant difference for PA3 (p = 0.98) compared to matched Control.

## DISCUSSION

To investigate possible molecular mechanisms of *LMNA*-Related DCM due to a *c.357-2A>G* splice-site mutation, we used a Patient-derived iPSC model of CM differentiation, serial scRNA-seq studies, and a comprehensive bioinformatics workflow. Our main findings include: 1. Model complexity with heterogenous cell types, possible subtypes, and lineages after sequential QC data processing and multi-level analyses; 2. Differential expression of genes including *LMNA* and enriched gene sets for cell signaling, cardiogenesis, and EMT involved in gene regulation and cell development in specific *LMNA* mutant cell types and lineages; 3. Differential expression of XL genes including *XIST*, six imprinted genes, and gene sets for regulation of metabolism, homeostasis, and proliferation. In our iPSC-derived model for Lamin A/C haploinsufficiency, our findings support a possible molecular mechanism involving cell type and lineage-specific dysregulation of gene expression in epigenomic developmental programs for PP, CM, and EPDC. We also discuss the challenges of data interpretation in our iPSC-derived model.

### Complexity of iPSC-derived model and comparisons to previous serial scRNA-seq studies of normal CM differentiation

After iPSC validation, we differentiated one *LMNA* mutant line derived from an affected female (Patient) and two non-mutant lines derived from her unaffected sister (Control) for scRNA-seq. We collected 12 total samples (4 Patient and 8 Control samples) across seven time points: D00, D02, D04, D09, D16, D19, and D30 with paired samples at D00, D09B, D16, and D19. After sequential QC data processing to remove of ambient RNA, low-quality cells, and doublets, we identified 110,521 total high-quality cells. We conducted multi-level analyses of high-quality cells by unsupervised clustering and annotation using known markers (88 total genes) to identify main cell types and possible cell subtypes in Single Sample, Merged, and Integrated Data and Subset Data. Here, we compared our results to previous scRNA-seq studies [27–36, 58].

For our Control samples (65,324 high-quality cells) collected across all seven time points (D00 to D30), we identified ten main cell types: PP, ME, CMESO, ENDO, ECTO, ENDOTH, CP, CM, EPDC, and UNK. These main cell types are consistent with seven serial scRNA-seq studies on control human iPSC [29, 30, 32, 33, 35, 36] and human ESC [31] differentiated to CM using the Wnt small molecule method. One exception was ECTO identified as a main cell type in only two [30, 33] of seven studies. Absence of ECTO in five studies [29, 32, 33, 35, 36] may be due to the small proportion of ECTO cells (2% for D09B CA1), lack of cell annotation markers like *PAX6* or *SOX1* [63], and/or biological variability of different control iPSC lines.

By subclustering our Control data, we identified many possible cell subtypes including four PP subtypes, two CP subtypes, two CM subtypes, and six further differentiated CM and EPDC subtypes at Day 30. Among our PP subtypes at Day 0, the predominate PP-A subtype (88%) represents a combined cluster, after regression of CC scores of cells, of two reported main PP subpopulations: Core and Proliferative PP [27, 28]. For our three smaller subtypes, PP-B (3%) and PP-C (6%) might be like the reported smaller PP subpopulations: Early and Late Primed for Differentiation [27] while PP-D (3%) comprises partially differentiated cells with lower expression of a core PP marker *POU5F1* [92] and the undifferentiated cell marker *CMND* [69]. Within our CP cell type at Day 9A and 9B, CP cells subclustered into subtype CP-A with higher expression of *HAND1* and *HAPLN1* and similar expression pattern to early CM and into subtype CP-B with higher expression of *GATA4* and *HAND2* and similar expression pattern to early EPDC. Thus, these two subtypes comprise distinct progenitor cells for CM (CP-A) and EPDC (CP-B) like two reported progenitor subpopulations at Day 5: CM Precursor (D5:S1) and Cardiovascular Progenitor (D5:S3) [29].

We also identified possible cell subtypes for CM and EPDC by Subcluster Analyses. Within our CM cell type at Days 9B, 16, and 19, cells subclustered into subtype CM-A (70%) with higher expression of multiple CM markers and Ribosomal Protein Genes and subtype CM/UNK-B (30%) with lower expression of most CM markers except *TTN* and greater levels of %Mt. The predominate CM subtype CM-A is consistent with reported single CM populations, Committed CM (D15:S2) [29] and D11-15 Early CM [36]. In contrast, subtype CM/UNK-B is unique to our study likely due to cell type-specific QC processing; we used high %Mt threshold levels for low-quality cells for only Non-CM cells and not CM cells because of higher Mt content of heart tissues [56, 57]. Thus, our approach allowed identification of subtype CM/UNK-B, cells typically discarded prior to annotation; subtype CM/UNK-B will need to be evaluated further as either low-quality/dying CM or a possible unique CM subtype characterized by high mitochondrial content. Finally, at Day 30, we identified later differentiated CM subtypes: ATR-CM and VENTR-CM by recognizable gene expression patterns. One of three similar scRNA-seq studies with Day 30 reported six distinct cell populations as atrial and ventricular cells using the Fluidigm C1 system for 43 *TNNT2*+ *ACTC1*+ subclustered cells [30]. Without subclustering at Day 30, two other studies reported a single CM cell type as Late CM [36] and as Definitive CM (D30:S2) [29]. Our results demonstrate the importance of Subcluster Analyses for CM and later time points for identification and further analysis of differentiated subtypes [34].

Likewise, Subcluster Analyses for EPDC at Day 30, without metabolic enrichment, identified four possible subtypes: EPI, CFIBRO, VSM, an Unspecified EPDC subtype with recognizable gene expression patterns. These patterns were not observed for EPDC at earlier time points, Day 16 and 19 with metabolic enrichment. The three similar scRNA-seq studies with Day 30 did not report these EPDC subtypes after metabolic enrichment [29, 30, 36]. Like CM, the two studies without Subcluster Analyses reported a single EPDC cell type as Epicardial Cells [36] and non-contractile cells (D30:S1) [29]. Taken together, our results provide first, additional evidence for continued presence of EPDC [58] despite metabolic enrichment for CM by glucose deprivation/lactate enrichment [45] and second, at later time points, four possible EPDC subtypes requiring further analysis.

After identification of cell types and subtypes, we analyzed Single and Paired Subset Data using Slingshot and found multiple lineages with single trajectories and two bifurcating trajectories. For Control Single Subset Data, we identified three single trajectories (PP-A to PP-B, CMESO to CP, CP-A to CM-A) and one bifurcating trajectory (ME to CMESO and ENDO). For Paired Subset Data of primarily “balanced” subtypes (62,488 total cells), we also identified the single trajectory at Day 0 (PP-A to PP-B) for both Control and Patient cells and a second bifurcating trajectory (CP into CM and EPDC). Our first lineage bifurcation at Day 2 (ME to CMESO and ENDO) is like reported lineages using different trajectory methods including Scdiff [29], Monocle 2 [31], and STREAM [36]. Our second lineage bifurcation of CP at Day 9 (CP into CM and EPDC) is consistent with previous reported lineages: Cardiac Precursor cells (D5:S1 and D5:S3) to Committed CM (D15:S2) and non-contractile cells (D15:S1) [29], D5 CP to D7-D15 CM and CFIBRO [35], and D5-D7 Late CP to D11-D15 Early CM and D5-D15 Epicardial Progenitors/Epicardial cells [36]. Overall, our results for the main cell types, several subtypes, and lineages with at least two bifurcations are consistent with previous scRNA-seq studies that support the validity of our complex iPSC-derived model.

### Patient vs. Control: Differential expression of *LMNA* and gene sets for cell signaling, cardiogenesis, and EMT involved in epigenetic gene regulation and cell development for CM and EPDC with Lamin A/C haploinsufficiency

Using Integrated Data, we conducted Comparative Analyses that identified two types of DEG for Patient compared to Control cells: Cell Type DEG and Lineage DEG. We identified Cell Type DEG using four shared cell types (PP, CP, CM, EPDC) in 14 primarily ‘balanced’ subtypes (71,541 total cells). We identified Lineage DEG using 18 paired subtypes (62,488 total cells) that derived three lineages: a single trajectory from PP-A to PP-B at Day 0, and a bifurcating trajectory from CP to CM (CP Lineage-1) and EPDC (CP Lineage-2) across Days 9, 16, and 19. Using top Cell Type and Lineage DEG, pathway enrichment by ORA and GSEA provided evidence for dysregulation of multiple biological processes and important insight into *LMNA*-related DCM pathogenesis.

Among top Cell Type DEG, we found our DCM study gene *LMNA* underexpressed in Patient cells compared to Control cells for shared cell types and lineages. In Control PP at Day 0, we detected limited *LMNA* expression that increased along normal CM differentiation as reported [22–24]. In Patient PP, we observed a similar pattern but at lower levels with the greatest differences in *LMNA* expression at Day 19 CM-A1 and EPDC. We confirmed decreased Lamin A/C protein expression at Day 19 by WB using two additional affected Patients from the family with matched Controls. In addition, for all shared cell types, we found monoallelic expression of the normal *LMNA* allele in Patient cells like our previous results on *LMNA* mutant dermal fibroblasts derived from the study family [81]. Our results support a molecular mechanism of *LMNA* haploinsufficiency for both cell types, CM and EPDC in our iPSC-derived model.

To evaluate possible effects of *LMNA* haploinsufficiency and other top Cell Type DEG, we used pathway enrichment that provided evidence for differences in regulation of epigenetic gene expression and cell development between conditions. First, EPDC progenitor cells at Day 9 (CP-B) enriched for epigenetic gene expression attributed to decreased expression of the non-coding *XIST* RNA gene that initiates XCI [85], and *NDN*, the paternally expressed imprinted gene [93], and for muscle development attributed to increased expression of keratin and myosin light chain protein encoding genes, *KRT19* and *MYL9.* EPDC at Day 19 also enriched for epigenetic expression and heterochromatin formation attributed to decreased expression of *LMNA*, *NDN*, and/or *XIST*.

Using top Lineage DEG, pathway enrichment also found evidence for differences in the regulation of transcription in all three lineages between conditions. The PP-A to PP-B lineage enriched for regulation of transcription by RNA polymerase II with underexpression of 12 genes including ten Zinc-finger proteins (ZNFs) genes such as *ZNF208* [87]. For both CP lineages, our analysis also identified *ZNF208* as the top Global Lineage DEG with limited expression in Patient compared to Control CP along CM and EPDC lineages. Zinc-finger proteins bind DNA, regulate gene transcription, and play roles in many cellular processes including cell proliferation, differentiation, and cancer progression [94]. For example, ZNF208 may serve as a tumor suppressor with an epigenetic mechanism of differentiation to adenocarcinoma from *ZNF208* gene silencing due to promoter hypermethylation and *ZNF208* underexpression [95]. Thus, it is possible that *LMNA* haploinsufficiency results in epigenomic dysregulation of *ZNF* gene expression during differentiation of CP to EPDC, a similar pathogenic mechanism proposed for defective myogenesis in patients with Emery-Dreifuss muscular dystrophy associated with *LMNA* R453W missense mutation [11, 96].

For CM progenitor cells and CM, our enrichment results also provided evidence for differences between conditions in developmental pathways including both cardiogenesis and EMT. CP-A cells first enriched for muscle cell differentiation attributed to decreased *LMNA* and *NKX2-5* encoding a key transcription factor in cardiogenesis and myocardial regeneration [97, 98]. The predominate CM subtype at Day 16 then enriched for muscle contraction and development attributed to underexpression of 20 different DEG including *NKX2-5* and WNT signaling protein encoding gene *WNT2* and for muscle development and differentiation attributed to overexpression of 31 different DEG including sarcomere protein encoding gene *ACTC1.* At Day 16, CM also enriched for EMT attributed to the upregulation of 23 DEG including mesenchymal marker *CDH2* and known EMT and XL gene *TIMP1* [91]. At Day 19, CM continued enrichment for muscle differentiation attributed to underexpression of 15 different genes including three DEG encoding key components and regulators for sarcomere development: *LMNA* [99], *MYOZ2* [100], and *SYNPO2L* [101] and for myogenesis and muscle contraction attributed to overexpression of 49 different DEG including the XL *DMD* gene and sarcomere protein encoding genes: *MYBPC3*, *MYL2*, *MYL3*, *and TNNT2*.

These results for *LMNA* haploinsufficiency in our iPSC-derived model suggest cardiomyocyte dysfunction due to disruption of gene expression in EPDC and CM as reported in two other iPSC-derived models using bulk RNA-seq for *LMNA* K117fs [21] at Day 40 and *LMNA* R225X at Day 14 [22]. Our results are also like those from studies of Lamin A/C loss-of-function that reported gene expression changes with upregulation for EMT in Lmna -/- mouse heart tissue using bulk RNA-seq [16]. Another recent study reported precocious CM development with upregulation of *Nkx2-5* and sarcomere genes in *Lmna* -/- mouse ESC-derived CP and CM and in *LMNA* knockdown iPSC-derived CM using bulk RNA-seq and qPCR [24]. Like for these studies, our results for *LMNA* haploinsufficiency in our iPSC-derived model supports cell type-specific disruption of epigenomic developmental pathways for cardiogenesis and EMT in *LMNA* mutant CP differentiation into EPDC and CM.

Using enrichment analysis, we also found evidence for differences in cell signaling pathways involving TGF Beta superfamily for CM and mTORC1 for EPDC at Day 16. The predominant CM cell type enriched for RSTK and TGF Beta signaling pathways attributed to underexpression of multiple DEG including *SMAD6* encoding signaling inhibitor [102], and *ID1*, *ID2*, and *ID3* encoding three Inhibitor Of DNA Binding proteins that regulate signaling in heart development [103]. The CP to CM lineage also enriched for regulation of BMP signaling pathway attributed to overexpression of four DEG encoding regulatory proteins involved in heart growth and development: *BMP4* [104], XL gene *GPC3* [105], *FST* [106], and *LRP2* [107] and regulation of response to growth factor attributed to overexpression of seven DEG such as *GPC1*, encoding Glypican Proteoglycan 1, a modulator of signaling pathways that regulate cell growth, motility, and differentiation [108]. In contrast to the CP to CM lineage, EPDC at Day 16 enriched for the mTORC1 signaling pathway attributed to the upregulation of genes involved in metabolism of glucose, lactate, and amino acid metabolism.

These results for *LMNA* haploinsufficiency in our iPSC-derived model suggest both CM and EPDC dysfunction due to dysregulation of TGF Beta and mTORC1 signaling as reported in previous studies using *Lmna* mutant mouse models and Patient samples. These studies reported hyperactivation of TGF Beta signaling in *Lmna* -/- MEF [13] and in *Lmna* H222P/H222P mouse cardiomyocytes and heart tissue with myocardial fibrosis and increased collagen expression [14]. Another study reported hyperactivation of AKT-mTORC1 in heart tissue from *Lmna* H222P/H222P mice and from three Patients with *LMNA* mutation [15]. A recent study also provided evidence for the dysregulation of TGF Beta/BMP pathway and *BMP4* overexpression using LAD analysis in *LMNA*-Related DCM heart tissue [17, 18]. Similarly, the enrichment results in our iPSC-derived model for *LMNA* haploinsufficiency support altered cell specific responses to signaling pathways involving TGF Beta/BMP in mutant CM and mTORC1 in mutant EPDC.

### Patient vs. Control: Differential expression of X-linked genes including *XIST*, six imprinted genes, and gene sets involved in regulation of metabolism, homeostasis, and proliferation

Among top Cell Type and Lineage DEG, we found the non-coding *XIST* RNA gene that initiates XCI [85]. For XIST, we detected no expression or lower expression levels of *XIST* in Patient compared to Control cells at all time points and shared cell types. Despite limited *XIST* expression in Patient cells, we detected primarily monoallelic expression for seven XL genes (*PRICKLE3*, *MAGED2*, *MAGEH1*, *KIF4A*, *ARMCX4*, *HTATSF1*, and *IDS*) known to be subject to XCI [88] consistent with reported retention of clonal X-inactivation during fibroblast reprogramming and iPSC differentiation [109]. In contrast, for four of five XL genes (*CD99*, *EIF2S3*, *PLS3*, *TMEM187*) with consistent or variable XCI escape [88], we detected biallelic expression of equal proportions across Patient cells, compared to skewed expression across Control cells, during iPSC differentiation to CM and EPDC to suggest possible partial X chromosome reactivation or XCI erosion. Since the nuclear lamina plays an important role in *XIST* mediated X chromosome gene silencing [110, 111], we might speculate our findings for altered *XIST* and XL gene expression may be evidence for disruption of epigenomic developmental pathways due to Lamin A/C haploinsufficiency.

In addition to *XIST*, among top Cell Type and Lineage DEG, we found six imprinted genes: *SNRPN*, *PWAR6*, *NDN*, *PEG10*, *MEG3*, and *MEG8* located at known imprinted regions. In Patient compared to Control cells, we found decrease expression for four paternally expressed genes: *SNRPN*, *PWAR6*, *NDN* at chr15q11.2 and *PEG10* at 7q21 and increased for two maternally expressed genes: *MEG3* and *MEG8* at chr14q32.2. Since these imprinted genes lacked informative coding SNV, we could not determine biallelic vs. monoallelic expression. In a genomic imprinting model using mouse ESC, biallelic expression of maternally expressed *Meg3* occurred further away from the nuclear periphery [112]. Similarly, we might speculate our findings in human cells with Lamin A/C haploinsufficiency is due to disruption of epigenomic developmental pathways, altered genome organization, and subsequent altered or loss of imprinting; further studies on a possible role of Lamin A/C in maintenance of XCI and genomic imprinting are needed.

Using enrichment analysis of top DEG, we also found metabolic gene signatures consistent with both cell type and metabolic selection as reported [45, 113] but also differences in cell metabolism between Patient and Control cells by cell type and lineage. First, the predominant PP subtype at Day 0 enriched for oxidative phosphorylation with ETC genes *DNAJC15*, *NDUFA3*, and *NDUFA6* underexpressed in Patient cells. For CP subtypes at Day 9, CM progenitors enriched for oxidative metabolism with 46 DEG downregulated in Patient cells while EPDC progenitors enriched for glycolytic metabolism with the phosphofructokinase encoding gene *PFKP* overexpressed in Patient cells. For both Control and Patient CM and EPDC, we also found unique metabolic gene signatures consistent metabolic selection for CM using lactate oxidation as reported [45, 113] but also EPDC survival using alternative metabolic pathways. For CM and EPDC at Day 16 (post-metabolic selection), CM enriched for glucose metabolism and oxidative phosphorylation with overexpression of 98 DEG while EPDC enriched for glucose and nucleotide metabolism with overexpression of 18 DEG. Only three DEG (*ATP5ME*, *COX4I1*, *P4HA2*) were overexpressed in both CM and EPDC.

These results in our iPSC-derived model demonstrated the ability to detect gene signatures associated with specific cell type, metabolic selection, and condition (Patient vs. Control). Two previous studies reported gene expression changes using bulk RNA-seq with downregulation of oxidative phosphorylation in Lmna -/- mouse heart tissue [16] and mitochondrial dysfunction in *LMNA* E342K iPSC-CM [114]. Using scRNA-seq to distinguish between CP, CM, and EPDC, our results in our iPSC-derived model with Lamin A/C haploinsufficiency might also suggest an association with metabolic activity dependent on mutant cell type and lineage. For example, based on our findings, metabolic activity might decrease in CM progenitors and increase in EPDC progenitors with Lamin A/C haploinsufficiency.

In addition, our enrichment results provided evidence for differences in cell homeostasis and response between Patient and Control cells for two lineages. The PP-A to PP-B and CP to CM lineages enriched for metal homeostasis attributed to the overexpression of *BNIP3*, encoding a pro-apoptotic mitochondrial protein, and metallothionein protein genes (*MT*): *MT1E*, *MT1F*, *MT1G*, *MT1H*, *MT1X*, and *MT3* encoding various heavy metal binding proteins [87]. The BNIP3 and MT proteins play important roles in the regulation of normal cell processes including cell growth, differentiation, mitochondrial function, and responses to oxidative stress [115, 116]. A study on *Bnip3* overexpression mouse ESC and iPSC reported decreased ROS levels and DNA damage and increased mitochondrial respiration and DNA repair [117]. Another study reported *LMNA* S143P iPSC-CM had increased sarcomere disorganization with increased cellular stress due to hypoxia [118]. Similarly, our enrichment results in our iPSC-derived model for *LMNA* haploinsufficiency may suggest dysregulation of cellular homeostasis and responses to oxidative stress or hypoxia in *LMNA* mutant PP and CP to CM lineages.

Finally, our enrichment results provided evidence for differences in cell proliferation for CM at Day 16 and the CP to EPDC lineage. The predominant CM subtype enriched for Hallmark Myc targets V1 with upregulation of 36 DEG including ten spliceosome genes, *TYMS*, a canonical S-phase marker [60], and other proto-oncogene targets for activation of growth-related genes [78, 87]. The CP to EPDC lineage enriched for cyclin-dependent kinase activity attributed to overexpression, primarily in CP cells, of two CC genes, *CDKN3*, encoding a cyclin-dependent kinase inhibitor, and *CCNB2*, encoding Cyclin B2 required for control at the G2/M transition [87]. Our results, supporting increased proliferation of CM and EPDC progenitors, agree with reported increased proliferation and altered CC progression in *Lmna* -/- MEF [13] and increased *CCNB2* expression in cancer cell lines with *LMNA* knockdown [119]. However, our results disagree with reported restricted proliferation and delayed CC activity of *Lmna* -/- mouse CM [120] and downregulation of CC genes in *Lmna* -/- mouse ESC-derived CP and CM and in *LMNA* knockdown iPSC-CM [24]. These discrepancies may involve *LMNA* genotype (haploinsufficiency vs. loss of function), species (human vs. mouse), growth conditions including metabolic selection, and possible biological and technical variables in our iPSC-derived model that require further evaluations using single-cell analyses to define cell type-specific changes in cell proliferation and CC control.

## CONCLUSIONS AND CHALLENGES

In our on-going studies, focused on a multi-generation family with a heterozygous *LMNA* c.357-2A>G splice-site mutation to identify possible molecular mechanism of *LMNA*-Related DCM [37, 40, 41], we generated and differentiated iPSC lines, derived from an affected female (Patient) and her unaffected sister (Control) and conducted scRNA-seq across multiple time points. To evaluate cell heterogeneity, we used a comprehensive scRNA-seq bioinformatic workflow with serial, cell type-specific data processing. After extensive multi-level analyses of 110,521 high-quality cells, we found substantial complexity in our iPSC-derived model: ten main cell types, many possible subtypes, and several lineages including bifurcation of CP into CM and EPDC lineages consistent with previous scRNA-seq studies of normal iPSC differentiation. To identify cell type and lineage-specific mechanisms, we used Comparative Analyses of Patient and Control cells. In our analyses of top Cell Type and Lineage DEG, we found evidence for the ‘gene expression’ hypothesis as proposed in other *LMNA* disease models. Overall, using our Patient-derived iPSC model and single-cell transcriptomics, our study supports an association with Lamin A/C haploinsufficiency and cell type and lineage-specific disruption of epigenomic developmental programs such as gene transcription, cell signaling, and cardiogenesis and EMT differentiation pathways.

Although our evidence supports the ‘gene expression’ hypothesis and pathway dysregulation as proposed in other *LMNA* disease models, we acknowledge the challenges of data interpretation in our iPSC-derived model due to variability from biological and technical factors that can influence gene expression, epigenetic regulation, and cell differentiation; these confounding factors include reported variability between different iPSC lines and clones [121, 122] and variability from the susceptibility for epigenomic aberrations of human iPSC [123–125].

Thus, to validate proposed molecular mechanisms of *LMNA*-Related DCM using our Patient-derived iPSC disease model, we first will need to resolve or reduce confounding factors. To address variability between iPSC lines and clones, we will need to conduct a larger scRNA-seq study with both biological replicates and isogenic Control iPSC lines. Biological replicates, multiple *LMNA* iPSC derived from different family members, are needed to determine the levels of variability in CM differentiation and for gene expression between different *LMNA* mutant iPSC lines and clones [121]. In addition, to minimize variability due to differences in genomic background [122], we will need to create a matched set of isogenic Control iPSC lines by repairing the *LMNA* mutation in each Patient iPSC line using genome editing tools such as CRISPR-Cas9 or Base Editing [126, 127].

To improve our Patient-derived iPSC disease model, we also will need to address the susceptibility of normal iPSC lines for epigenomic aberrations that may influence multiple biological processes. Reported epigenomic aberrations include altered *XIST* expression, XCI erosion, and upregulation of XL genes [125], altered DNA methylation, and loss of imprinting for many genes such as *SNRPN*, *NDN*, *MEG3*, *PEG10* [123] associated with somatic reprogramming, molecular memory of cell origin, and cell culturing [125]. A recent study showed the effects of molecular memory in human iPSC of fibroblast origin with expression retained for fibroblast-specific genes and increased for mesoderm progenitor markers such as *BMP4* compared with human ESC [128]. To address the biological effects of molecular memory, we may try to wipe away fibroblast epigenomic memory in our *LMNA* Patient and Control fibroblast or iPSC lines to create cells in the naïve pluripotency state (naïve iPSC), that more closely resemble human ESC [129], using a new method called transient-naïve-treatment [128].

By improving our Patient-derived iPSC model using biological replicates, isogenic Controls, and naïve iPSC, it might be possible to separate the consequences of *LMNA* haploinsufficiency from confounding factors and to continue testing the ‘gene expression’ hypothesis for *LMNA*-related DCM. Finally, with the newly described role of Lamin A/C in naïve pluripotency and cardiovascular cell fate [12, 24], successfully addressing the limitations may enable new investigations to evaluate other possible mechanisms, including epigenomic dysregulation, using naïve iPSC-based models for *LMNA*-related DCM.

## SUPPLEMENTARY MATERIALS

### Supplementary Figures (S1-S14)

**Figure S1. Cardiomyocyte Differentiation Protocols A & B**

**Figure S2. iPSC Validation Results: Differentiation Capability by ICC Staining**

**Figure S3. Allelic Expression Using Coding Single Nucleotide Variants (SNVs)**

A. Autosomal Genes including *LMNA*

B. X-Linked Genes with X-Chromosome Inactivation (XCI) Status and Summary

**Figure S4. Workflow Step-I: Data Processing Results**

A. Summary: Sequential QC Processing of Raw-Corrected-Filtered-Singlet Data

B. Merged Data for All Single Sample Samples: QC Covariate Plots

C. Merged Data for All Single Sample Samples: QC Covariate Plots-Control vs. Patient

D. Merged Data for All Single Sample Samples: QC Covariate Plots by Cluster

E. Background RNA: Removal Using SoupX

F. Cell Quality: Removal of Low-Quality Cells Using Seurat

G. Doublets: Identification and Removal Using DoubletFinder (DF)

**Figure S5. Workflow Step-II: Clustering and Annotation**

A. Summary: Single Sample Data for Cell Annotation

B. Summary: Merged Single Sample Data for Cell Annotation (110,521 Total Cells)

**Figure S6. Workflow Step-II: Single Sample Data Results**

- Individual Analyses of Singlet Data for Main Cell Types

A. Summary: Raw Data to Annotated Clusters

B. Single Sample Data Analyses: Control Samples (n=8)

C. Single Sample Data Analyses: Patient Samples (n=4)

**Figure S7. Workflow Step-II: Single Sample Data Results**

- Subcluster Analyses of Subset Data for Possible Cell Subtypes

A. Summary: Annotated Clusters to Annotated Subsets

B. Subcluster Analyses: Control Samples (n=8) to Annotated Subset Data (n=21)

C. Subcluster Analyses: Patient Samples (n=4) to Annotated Subset Data (n=11)

**Figure S8. Workflow Step-III: Data Combining & Comparative Analyses**

A. Summary: Combined Data for Paired Sample Data

B. Summary: Combined Data for ‘Balanced’ Paired Subsets (n=6 Prs: 75,330 Total Cells)

**Figure S9. Workflow Step-III: Paired Sample Data Results**

- Individual Analyses of Combined Singlet Data for Shared Cell Types

A. Summary: Merged vs. Integrated Singlet Data-Clusters and Imbalance

B. Summary: Cell Annotation of Integrated Singlet Data (n=4 Prs: 89,269 Total Cells)

C. Paired Sample Data Analyses: Integrated Singlet Data-Patient vs. Control (n= 4 Prs)

**Figure S10.Workflow Step-III: Paired Sample Data Results**

- Individual Subcluster Analyses of Combined Subset Data for Possible Shared Subtypes

A. Summary: Merged vs. Integrated Subset Data-Clusters and Imbalance

B. Summary: Cell Annotation of Integrated Subset Data (n=11 Prs: 88,420 Total Cells)

C. Paired Sample Data Analyses: Integrated Subset Data-Patient vs. Control (n= 11 Prs)

**Figure S11.Workflow Step-III: Paired Sample Data Results**

- Comparative Analyses for Cell Type-Specific DE

A. Summary: Cell Type Differentially Expressed Genes (Cell Type DEG)

B. Summary: Volcano Plots and Cell Type DEG (n=14 Cell Subtypes: 71,541 Total Cells)

C. Individual Analyses of Integrated Subsets: Cell Type DEG and Enrichment (n=14)

D. Cell Type DEG: LMNA, X-Linked Genes, and Imprinted Genes Across 14 Subtypes

E. Cell Type DEG Enrichment: Module Scoring of GSEA Significant Gene Sets

**Figure S12.Workflow Step-III: Single Subset Data Results**

- Trajectory Analyses for Lineage-Specific DE and Enrichment

A. Summary: Annotated Subset Data to Cell Lineages

B. Single Subset Data: Trajectory, Lineage DEG, and Enrichment Analyses (n=7)

**Figure S13.Workflow Step-III: Paired Subset Data Results**

- Trajectory Analyses for Lineage-Specific DE and Enrichment

A. Summary: Lineage Differentially Expressed Genes (Lineage DEG)

B. Summary: UMAP Plots and Cell Lineages (n=2: 62,488 Total Cells)

C. Paired Subset Data Analyses: Pluripotent Cell Lineage (19,346 Total Cells)

D. Paired Subset Data Analyses: Cardiac Progenitor Lineages (43,142 Total Cells)

**Figure S14. Western blots (n=3)**

### Supplementary Tables (S1-S12)

Table S1. Human subjects and results for fibroblast LMNA sequencing and iPSC validation.

Table S2. Methods: Primary and secondary antibodies.

Table S3. Methods: Main software packages.

Table S4. scRNA-seq Summary Metrics using Cell Ranger.

Table S5. Single Sample Data: QC data processing using SoupX, Seurat, and DoubletFinder.

Table S6. Single Sample and Combined Data: QC processing of Raw to Singlet Data Matrices.

Table S7. Single Sample Data: Individual and Subcluster Analyses for Cell Annotation.

Table S8. Gene marker panels for cell annotation.

Table S9. Paired Sample Data: Individual and Subcluster Analyses.

Table S10. Single Subset Data: Trajectory Inference, Lineage DEG, and Enrichment.

Table S11. Paired Subset Data: Integration, Trajectory Inference, Lineage DEG, and Enrichment.

Table S12. Lamin A/C Western Blot Quantification Data and Statistical Analyses.

### Supplementary Excel Tables (S1-S4) [<u>Not included in BioRxiv pre-print</u>]

#### Excel Table S1. Cluster.Subcluster.DEG

Tab-1. Single Sample Data: All Cluster DEG for Cell Types

Tab-2. Single Sample Data: Top Cluster DEG for Possible Subtypes

Tab-3. Paired Sample Data: Top Conserved Cluster DEG for Shared Cell Types

Tab-4. Paired Sample Data: Top Conserved Cluster DEG for Shared Possible Subtypes

Tab-5. Combined Data: All Conserved Cluster DEG for Shared Cell Types

Tab-6. Combined Data: Enrichment Results by ORA for Shared Cell Types

#### Excel Table S2. CellType.DEG

Tab-1. Integrated Subset Data: Cell Type DEG in ‘Balanced’ Subtypes

Tab-2. Top 10 Threshold Cell Type DEG

Tab-3. Cell Type DEG Enrichment Results by ORA

Tab-4. Cell Type DEG Enrichment Results by GSEA

#### Excel Table S3. Lineage.DEG

Tab-1. Single Subset Data: Association and Start-End Test Lineage DEG

Tab-2. Integrated Subset Data: Association and Condition Test Lineage DEG

Tab-3. Integrated Subset Data: Association and Condition Test TOP DEG

Tab-4. Integrated Subset Data: Lineage DEG Enrichment Results by ORA

Tab-5. Integrated Subset Data: Lineage DEG Enrichment Results by GSEA

#### Excel Table S4. AllelicExpression

Tab-1. Allelic Expression Results for 10 Autosomal SNV including *LMNA*

Tab-2. Allelic Expression Results for 12 X-Linked SNV with XCI Status

## DECLARATIONS

### Author Contributions

MVZ, TB, HPW, MM, AG, and QN designed the study. HPW, MM, TB, and MVZ performed cell culture and differentiations. TB conducted WB experiments and analysis. HPW and MM conducted cell collections for scRNA-seq. MVZ, TB, and HPW interpreted results. HPW and MVZ conducted initial computational analyses of scRNA-seq data with assistance from ZC, YS, and QN. MVZ conducted final computational analysis and data visualization of scRNA-seq data. MVZ wrote and prepared manuscript draft. HPW wrote draft of scRNA-seq methods. TB wrote and prepared drafts for WB sections, graphical abstract, and supplementary figures. MVZ, AG, and QN acquired financial support. All authors read and approved the final manuscript.

### Data Availability Statement

Raw and processed data for single-cell RNA-sequencing will be available in a publicly accessible repository.

## Acknowledgements

We are grateful to the study family for participating in this research, to Nancy Oropeza and Amanda Sherman in UCI Department of Pediatrics for administrative support, to Melanie Oakes, PhD at the UCI Genomics High-Throughput Facility for technical and analytic support, and to Christina Tu and Allia Fawaz at the UCI Sue & Bill Gross Stem Cell Research Center for technical training and support.

## Institutional Review Board Statement

We conducted all procedures in accordance with the Declaration of Helsinki and the ethical standards of the University of California Irvine under protocols approved by the Institutional Review Board (2011-8030 and 2014-1253) and Human Stem Cell Research Oversight Committee (2014-1043).

## Informed Consent Statement

The six study family members gave their informed written consent before participation.

## TABLE

**Table.**
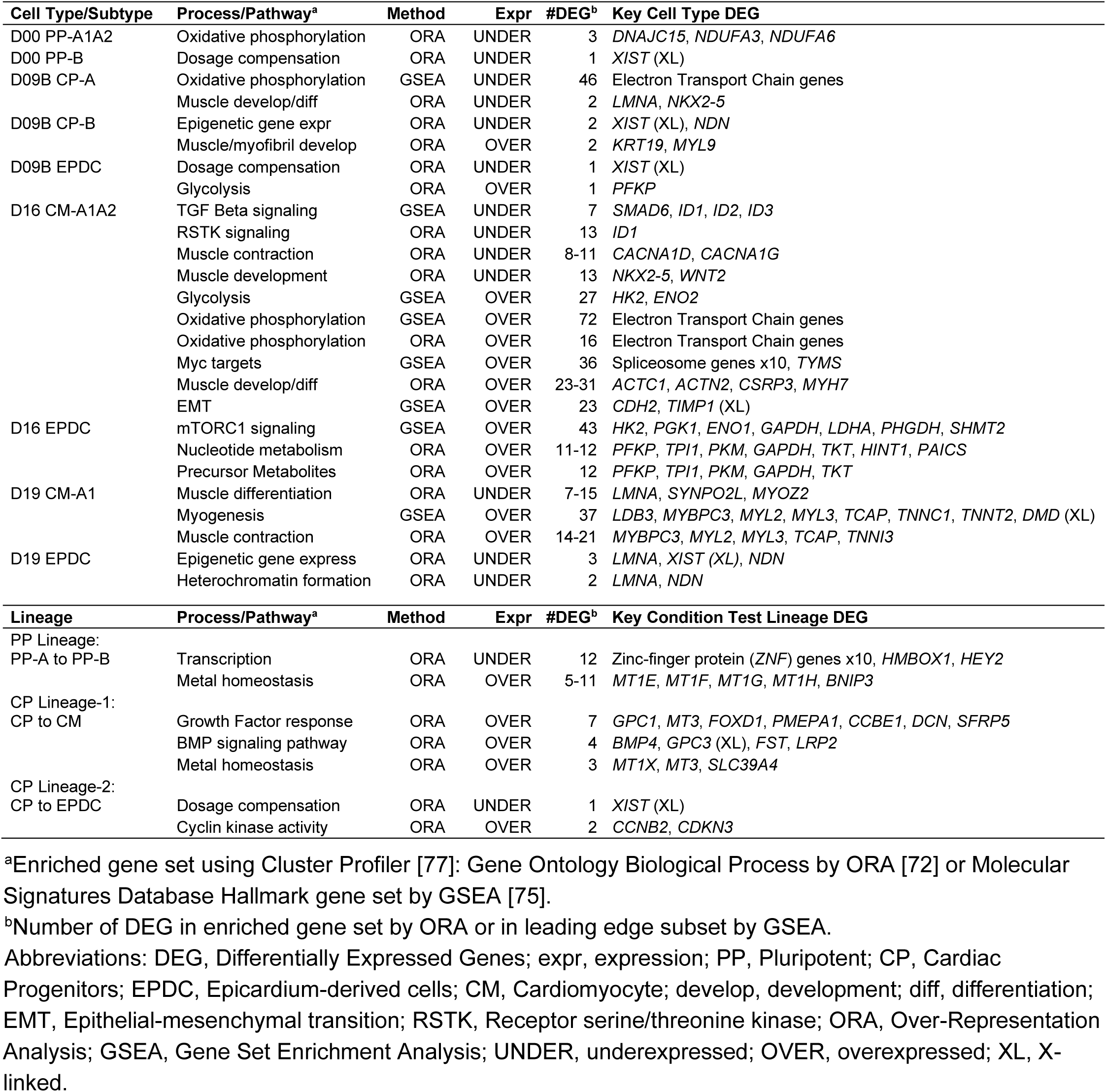
Enrichment Results for Top Cell Type and Lineage DEG.

