## Supplemental Figures for "*LMNA*-Related Dilated Cardiomyopathy: Single-Cell Transcriptomics during Patient-derived iPSC Differentiation Support Cell type and Lineage-specific Dysregulation of Gene Expression and Development for Cardiomyocytes and Epicardium-Derived Cells with Lamin A/C Haploinsufficiency"

**Supplemental Figures** **Page**

**Figure S1.** **Cardiomyocyte Differentiation Protocols A & B** **3**
**Figure S2.** **iPSC Validation Results: Differentiation Capability by ICC Staining** **4**
**Figure S3.** **Allelic Expression Using Coding Single Nucleotide Variants (SNVs)** **5 - 9**
**A. Autosomal Genes including *LMNA***
**B. X-Linked Genes with X-Chromosome Inactivation (XCI) Status and Results Summary**
**Figure S4.** **Workflow Step-I: Data Processing Results** **10 - 17**
**A. Summary: Sequential QC Processing of Raw-Corrected-Filtered-Singlet Data**
**B. Merged Data for All Single Sample Samples: QC Covariate Plots**
**C. Merged Data for All Single Sample Samples: QC Covariate Plots- Control vs. Patient**
**D. Merged Data for All Single Sample Samples: QC Covariate Plots by Cluster**
**E. Background RNA: Removal Using SoupX**
**F. Cell Quality: Removal of Low-Quality Cells Using Seurat**
**G. Doublets: Identification and Removal Using DoubletFinder (DF)**
**Figure S5.** **Workflow Step-II: Clustering and Annotation** **18 - 19**
**A. Summary: Single Sample Data for Cell Annotation**
**B. Summary: Merged Single Sample Data for Cell Annotation (110,521 Total Cells)**
**Figure S6.** **Workflow Step-II: Single Sample Data Results- Individual Analyses of Singlet Data for Main Cell Types** **20 - 32**
**A. Summary: Raw Data to Annotated Clusters**
**B. Single Sample Data Analyses: Control Samples (n=8)**
**C. Single Sample Data Analyses: Patient Samples (n=4)**
**Figure S7.** **Workflow Step-II: Single Sample Data Results- Subcluster Analyses of Subset Data for Possible Cell Subtypes** **33 - 45**
**A. Summary: Annotated Clusters to Annotated Subsets**
**B. Subcluster Analyses: Control Samples (n=8) to Annotated Subset Data (n=21)**
**C. Subcluster Analyses: Patient Samples (n=4) to Annotated Subset Data (n=11)**

Figure S8. Workflow Step-III: Data Combining & Comparative Analyses 46 - 47
A. Summary: Combined Data for Paired Sample Data
B. Summary: Combined Data for 'Balanced' Paired Subsets (n=6 Prs: 75,330 Total Cells)

Figure S9. Workflow Step-III: Paired Sample Data Results- Individual Analyses of Combined Singlet Data for Shared Cell Types 48 - 53
A. Summary: Merged vs. Integrated Singlet Data- Clusters and Imbalance
B. Summary: Cell Annotation of Integrated Singlet Data (n=4 Prs: 89,269 Total Cells)
C. Paired Sample Data Analyses: Integrated Singlet Data- Patient vs. Control (n= 4 Prs)

Figure S10. Workflow Step-III: Paired Sample Data Results- Individual Subcluster Analyses of Combined Subset Data for Possible Shared Subtypes 54 - 69
A. Summary: Merged vs. Integrated Subset Data- Clusters and Imbalance
B. Summary: Cell Annotation of Integrated Subset Data (n=11 Prs: 88,420 Total Cells)
C. Paired Sample Data Analyses: Integrated Subset Data- Patient vs. Control (n= 11 Prs)

Figure S11. Workflow Step-III: Paired Sample Data Results- Comparative Analyses for Cell Type-Specific DE 70 - 94
A. Summary: Cell Type Differentially Expressed Genes (Cell Type DEG)
B. Summary: Volcano Plots and Cell Type DEG (n=14 Cell Subtypes: 71,541 Total Cells)
C. Individual Analyses of Integrated Subsets: Cell Type DEG and Enrichment (n=14)
D. Cell Type DEG: *LMNA*, X-Linked Genes, and Imprinted Genes Across 14 Subtypes
E. Cell Type DEG Enrichment: Module Scoring of GSEA Significant Gene Sets

Figure S12. Workflow Step-III: Single Subset Data Results- Trajectory Analyses for Lineage-Specific DE and Enrichment 95 - 109
A. Summary: Annotated Subset Data to Cell Lineages
B. Single Subset Data: Trajectory, Lineage DEG, and Enrichment Analyses (n=7)

Figure S13. Workflow Step-III: Paired Subset Data Results- Trajectory Analyses for Lineage-Specific DE and Enrichment 110 - 121
A. Summary: Lineage Differentially Expressed Genes (Lineage DEG)
B. Summary: UMAP Plots and Cell Lineages (n=2: 62,488 Total Cells)
C. Paired Subset Data Analyses: Pluripotent Cell Lineage (19,346 Total Cells)
D. Paired Subset Data Analyses: Cardiac Progenitor Lineages (43,142 Total Cells)

Figure S14. Western blots (n=3) 122 - 124

**Fig. S1 CM Protocol**

### Cardiomyocyte Differentiation Protocols A & B

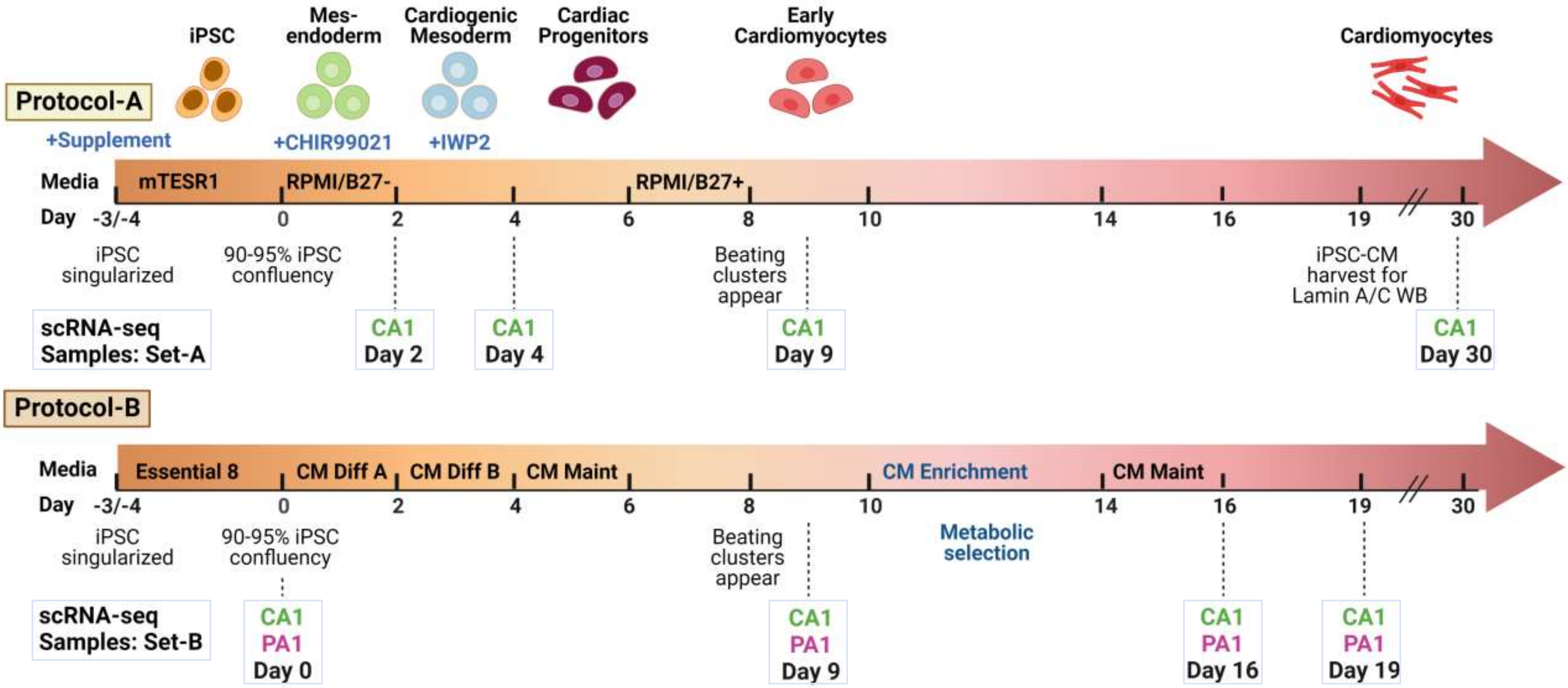

**Fig. S2 EB ICC**

### iPSC Validation Results: Differentiation Capability by Embryoid Body (EB) Immunocytochemistry (ICC) Staining

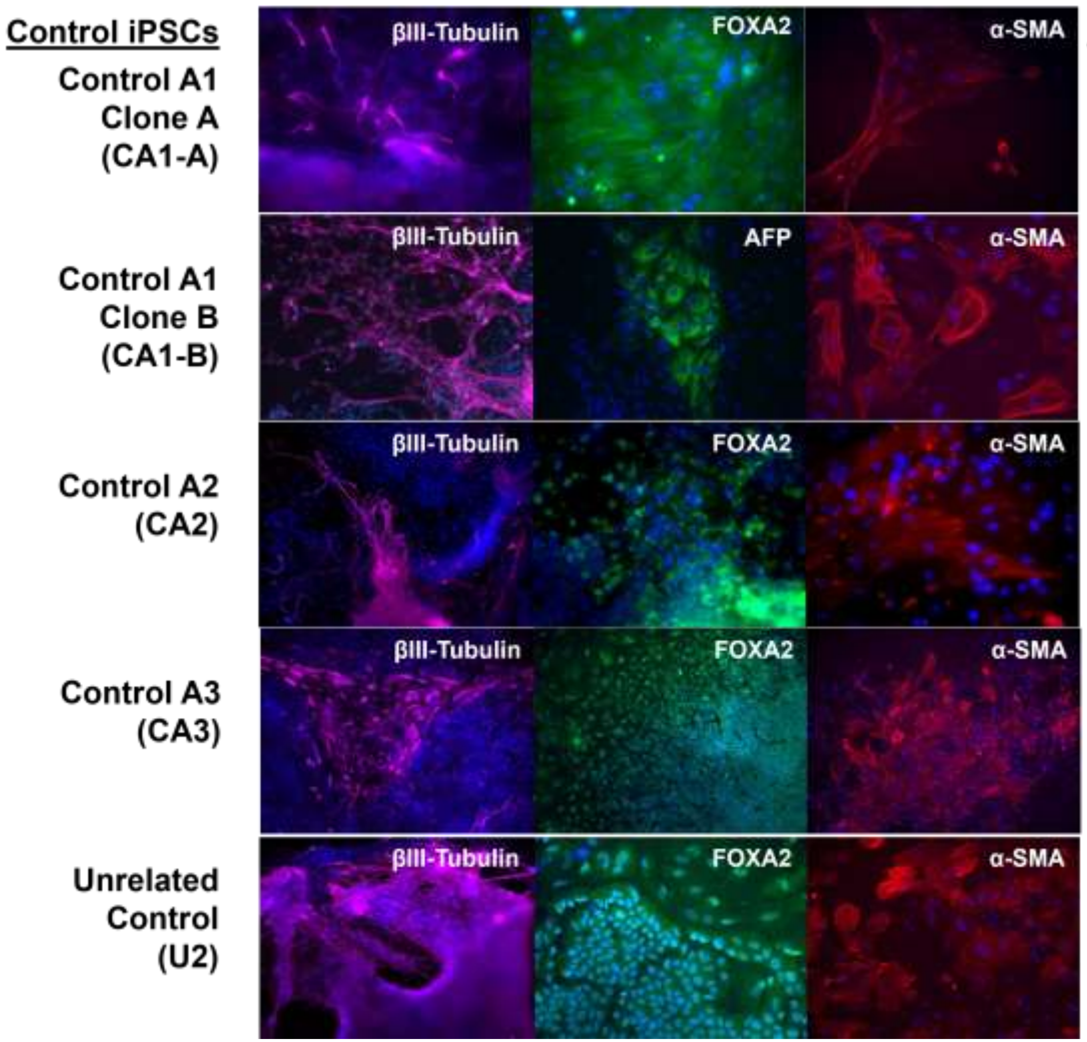


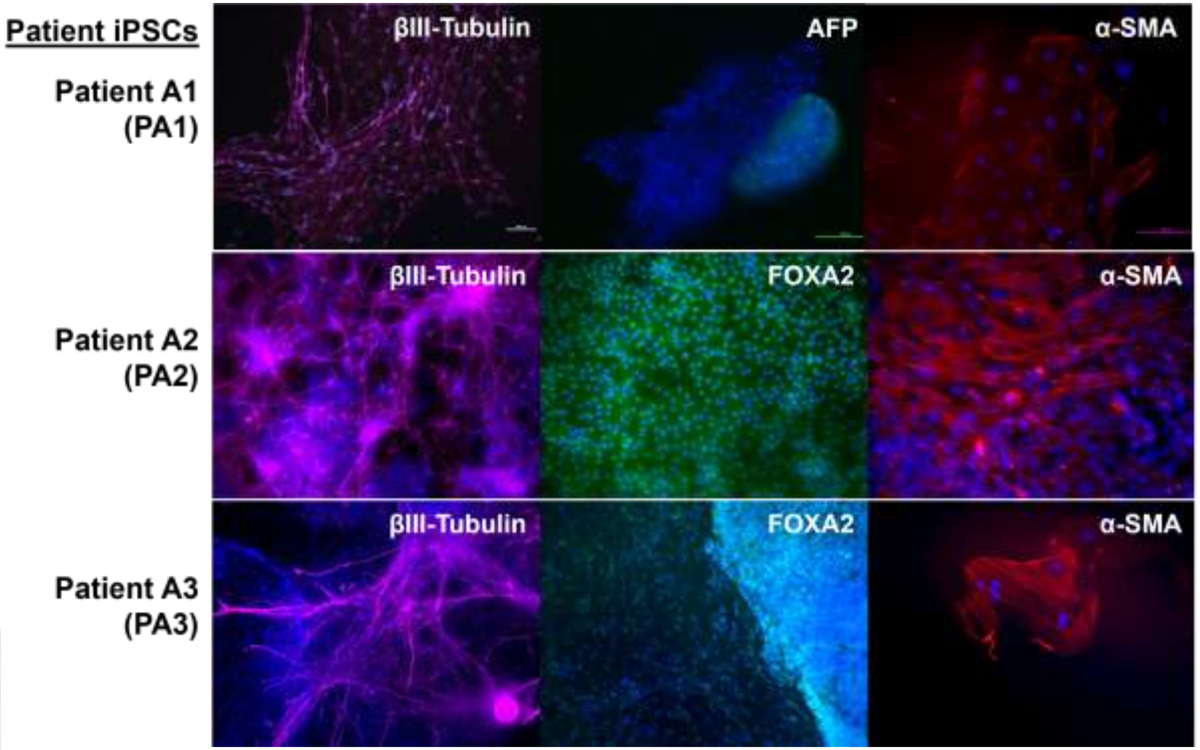


**Germ Layer Markers**

**Endoderm:** Forkhead Box A2 (FOXA2)
Alpha-fetoprotein (AFP)

**Ectoderm:** Beta-III Tubulin

**Mesoderm:** Smooth Muscle Actin ($\alpha$-SMA)

**Fig. S3 Coding SNV**

### Allelic Expression Using Coding Single Nucleotide Variants (SNVs)

#### A. Autosomal Genes including *LMNA*

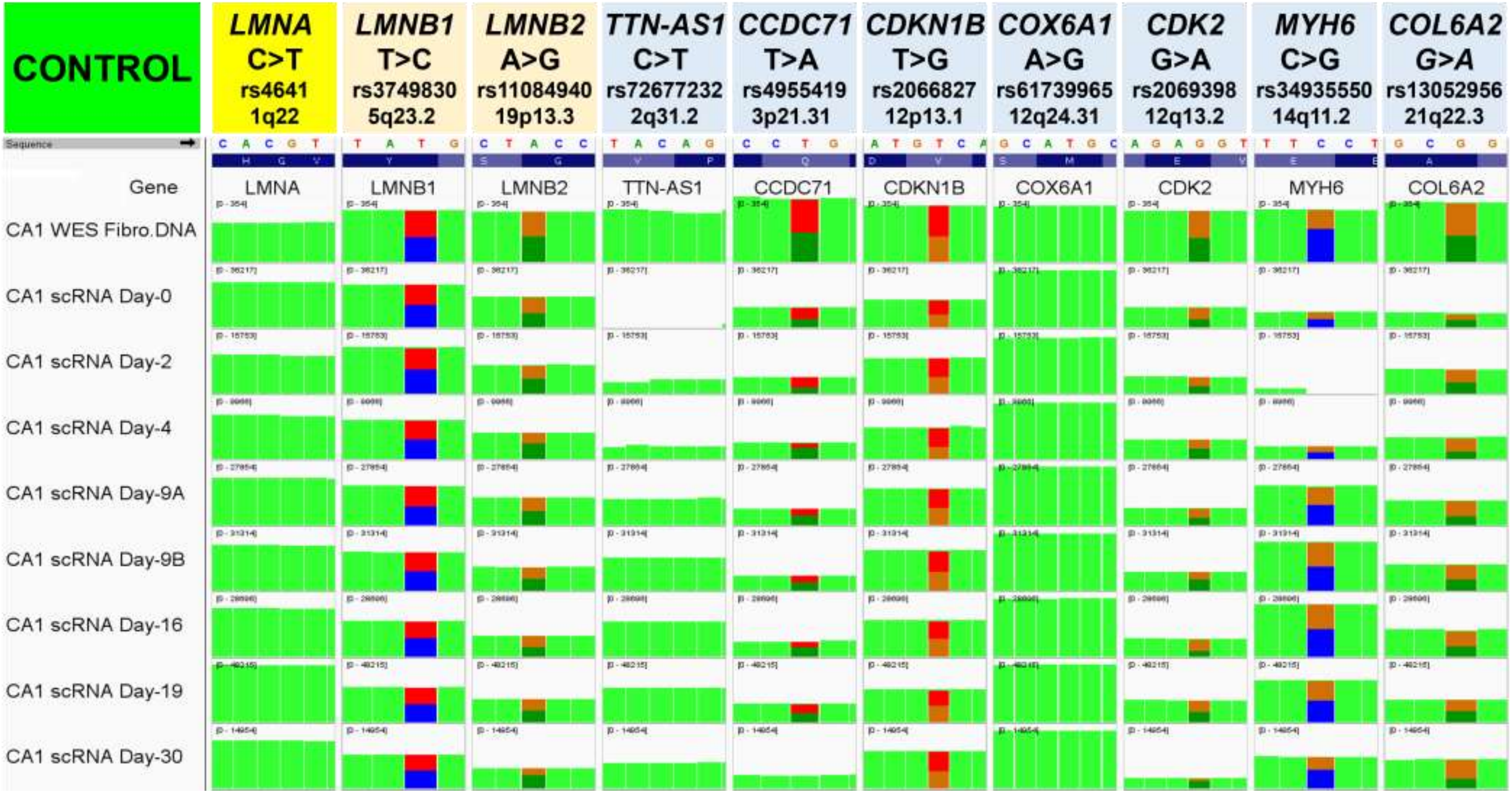


Integrative genomics viewer images of aligned reads to the human reference genome

**Fig. S3 Coding SNV**

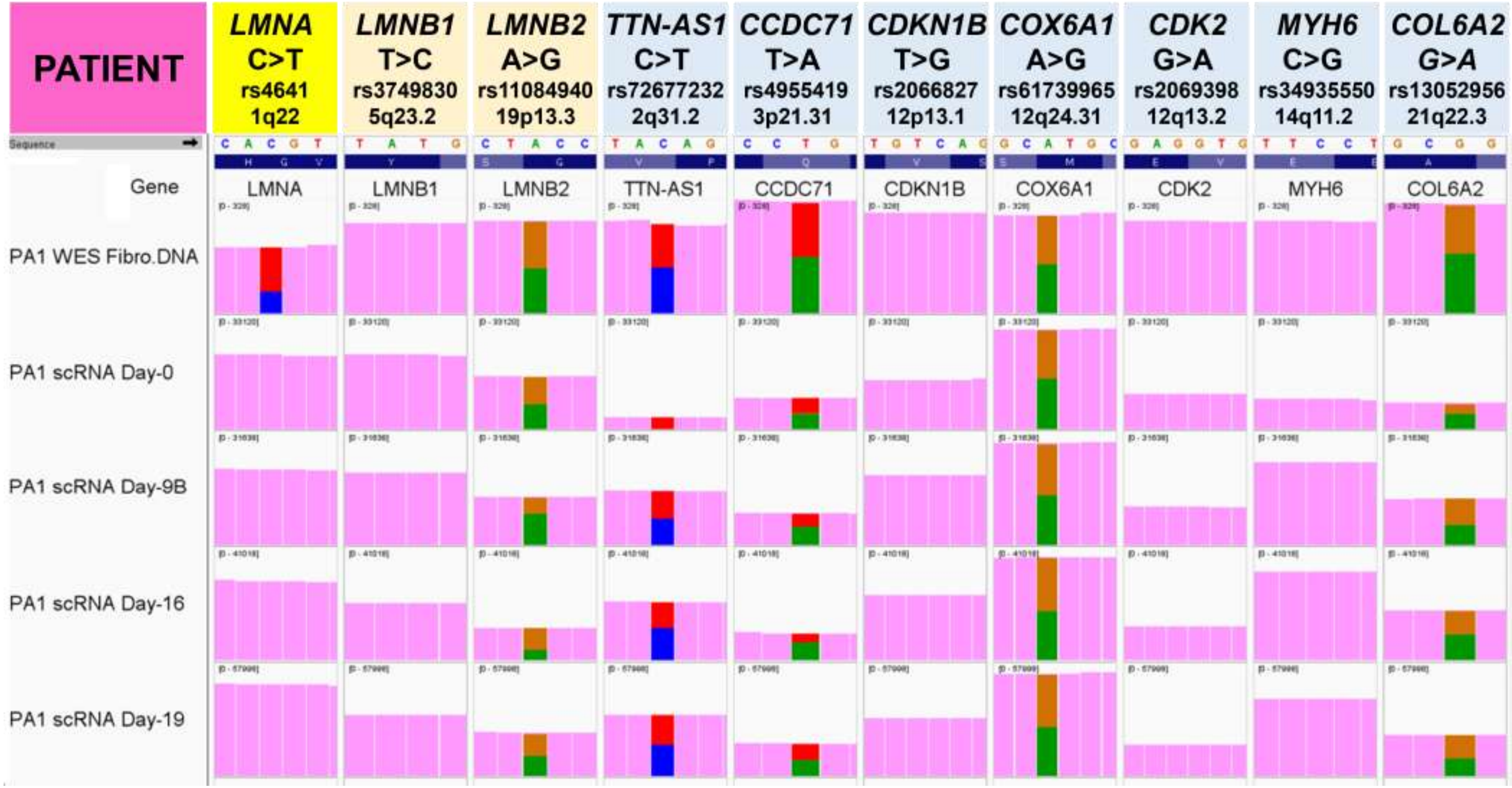

**Fig. S3 Coding SNV**

### B. X-Linked Genes with X-Chromosome Inactivation (XCI) Status*

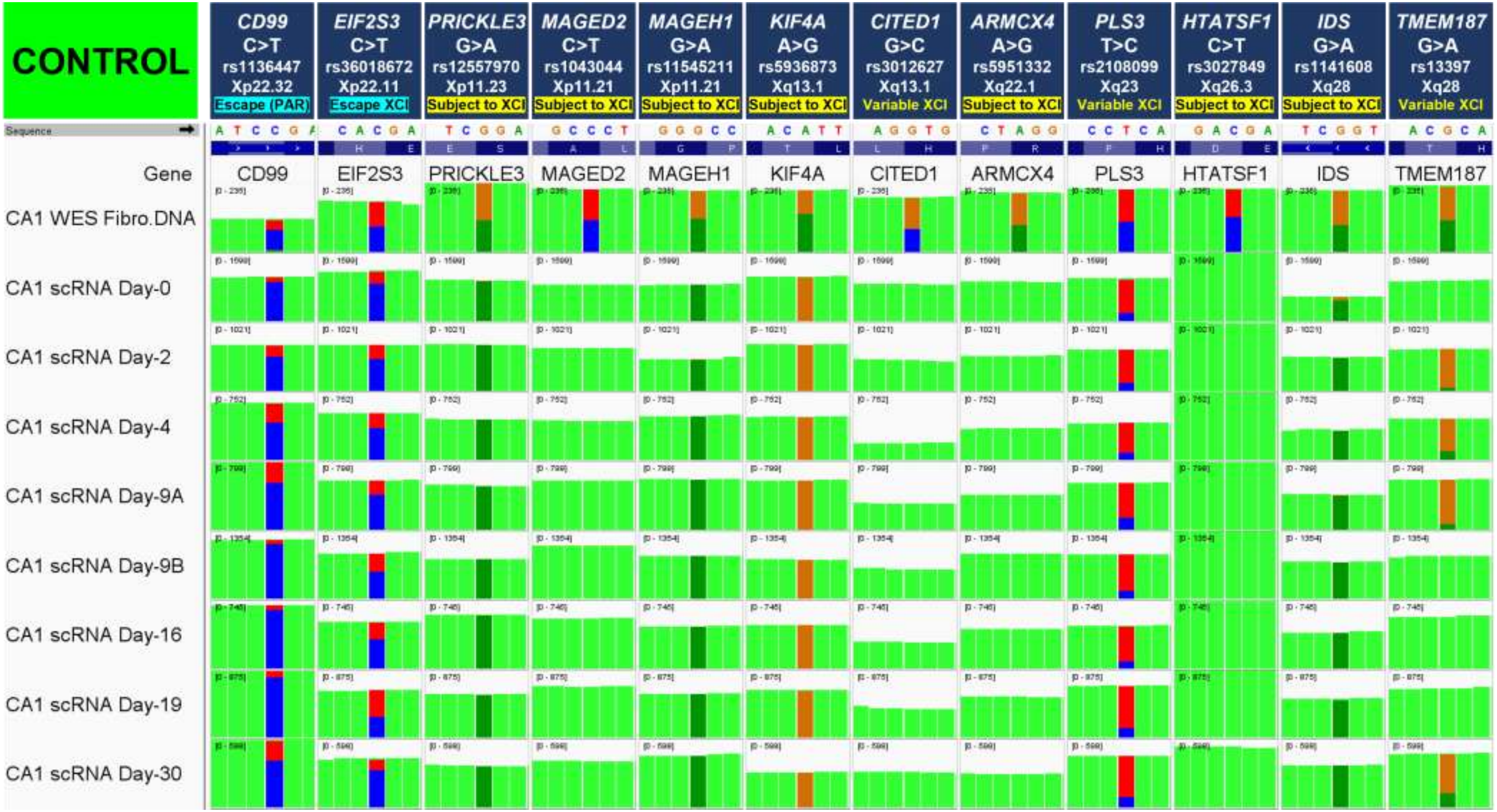

**Fig. S3 Coding SNV**

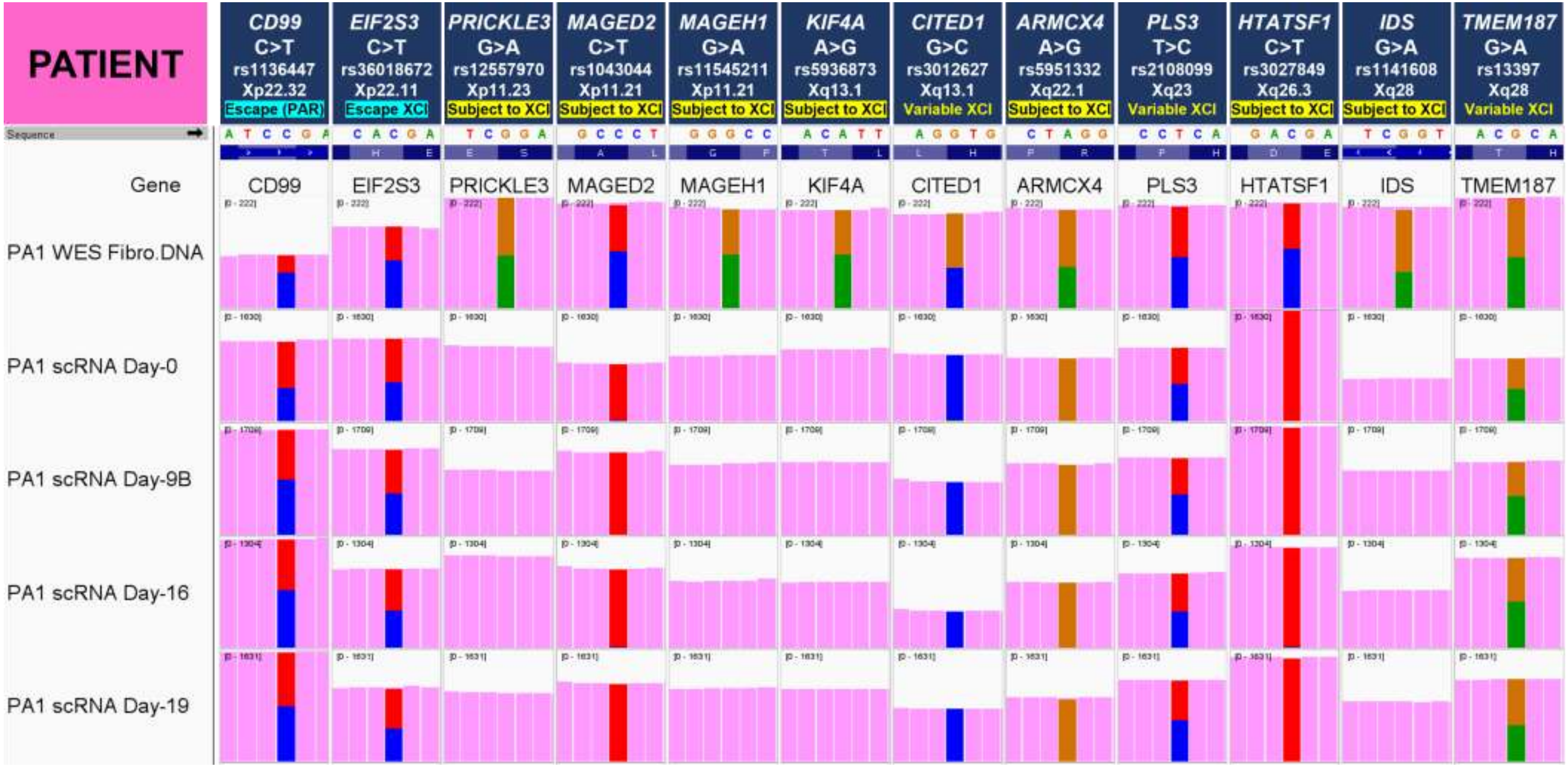


*XCI Status from Balaton *et al.* 2015

Fig. S3 Coding SNV

### Allelic Expression Using Coding Single Nucleotide Variants (SNVs)

#### RESULTS SUMMARY: X-Linked Genes with X-Chromosome Inactivation (XCI) Status

##### Total Read Count by Allele for 12 scRNA-seq Samples

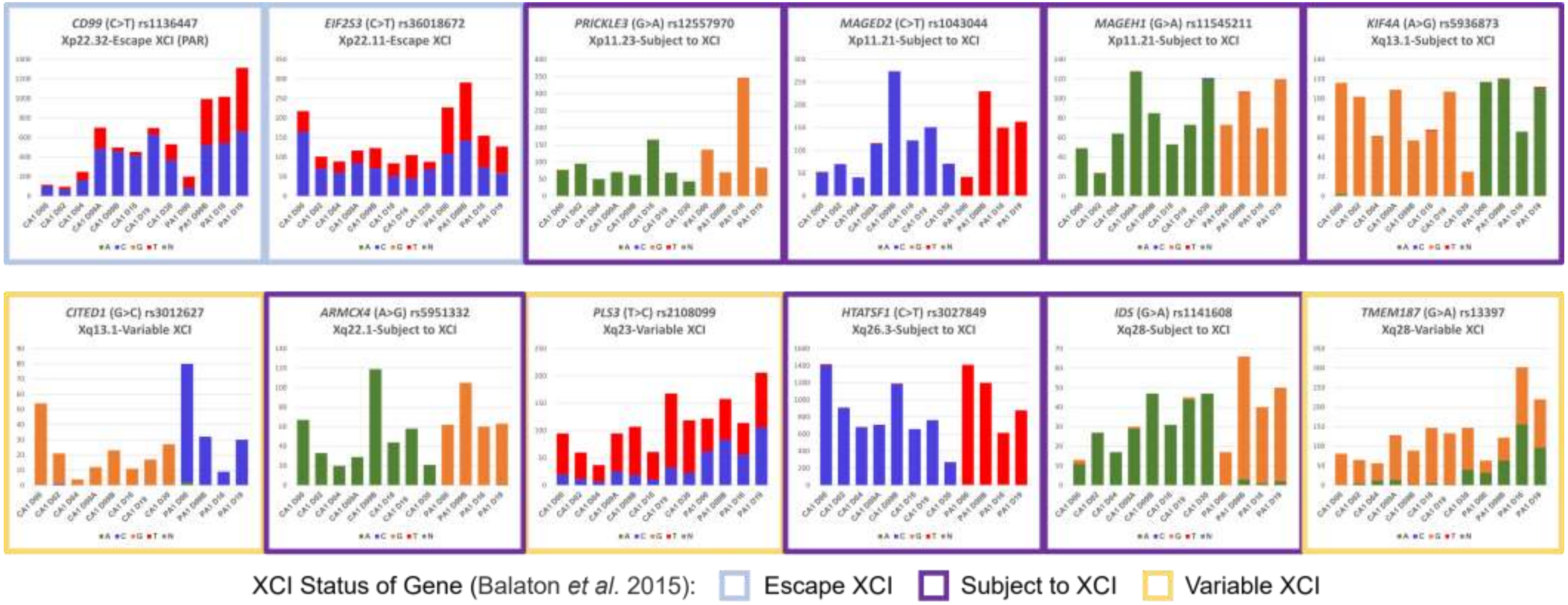

Fig. S4 Processing

### Workflow Step-I: Data Processing Results

#### A. Sequential QC Processing Summary: Raw-Corrected-Filtered-Singlet Data

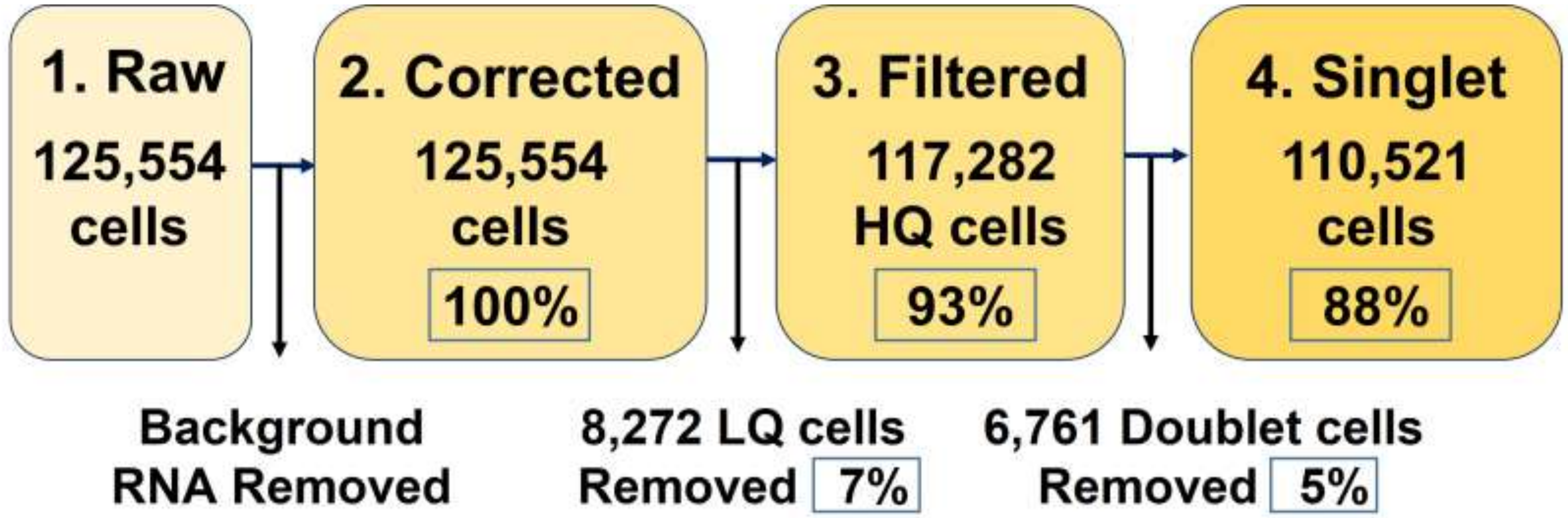


| | No. Samples | No. Cells (% Raw Total) | | | |
|---|---|---|---|---|---|
| | | 1. Raw | 2. Corrected | 3. Filtered | 4. Singlet |
| Control | 8 | 74,097 | 74,097 (100%) | 69,084 (93%) | 65,324 (88%) |
| Patient | 4 | 51,457 | 51,457 (100%) | 48,198 (94%) | 45,197 (88%) |
| All | 12 | 125,554 | 125,554 (100%) | 117,282 (93%) | 110,521 (88%) |

**Fig. S4 Processing**

#### B. Merged Data for All Single Sample Samples: QC Covariate Plots

**1. Raw Data**
**125,554**
**cells**

**2. Corrected**
**125,554**
**cells**
**100%**

**3. Filtered**
**117,282**
**HQ cells**
**93%**

**4. Singlet**
**110,521**
**Cells**
**88%**

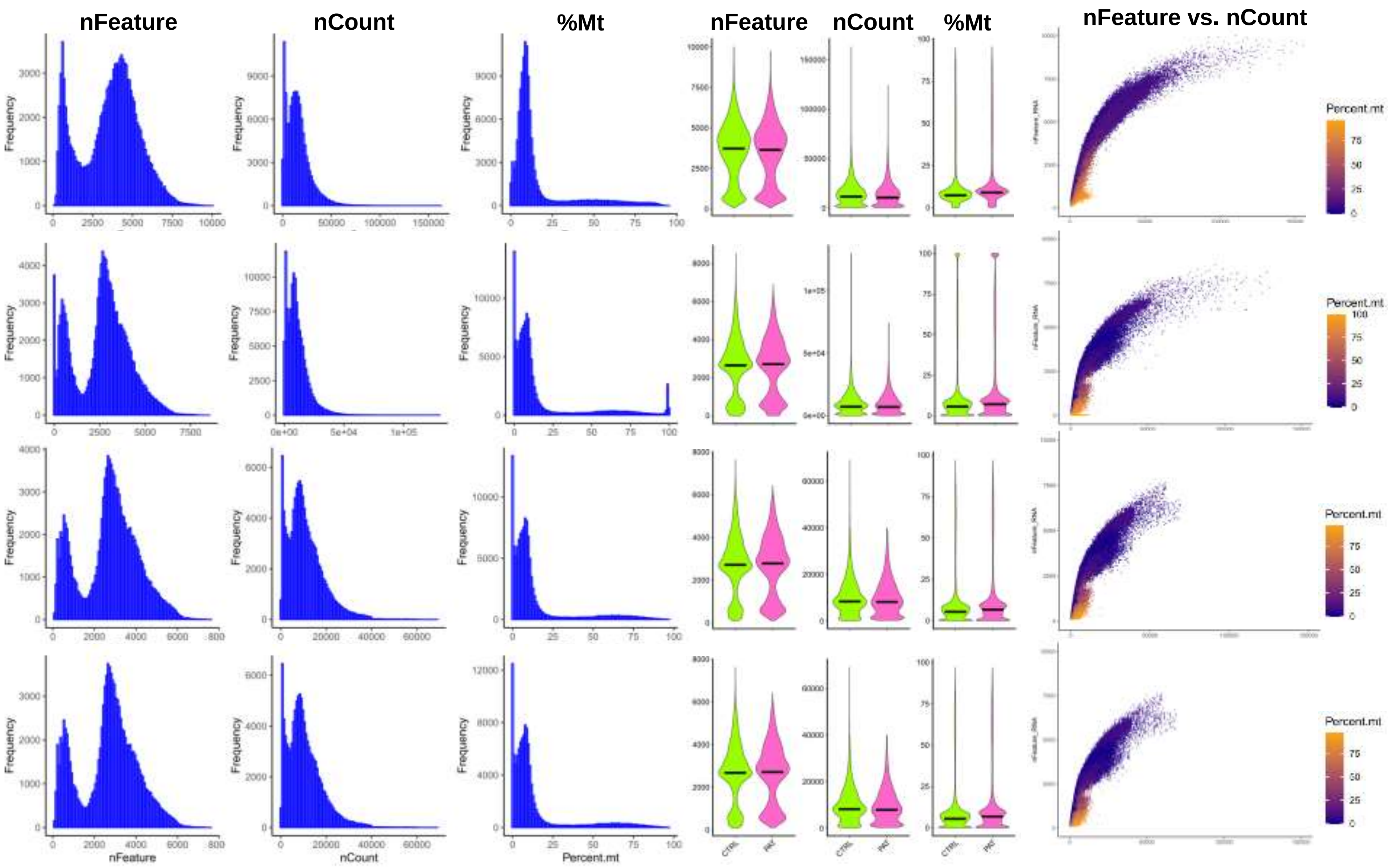

**Fig. S4 Processing**

#### C. Merged Data for All Single Sample Samples: QC Covariate Plots- Control vs. Patient

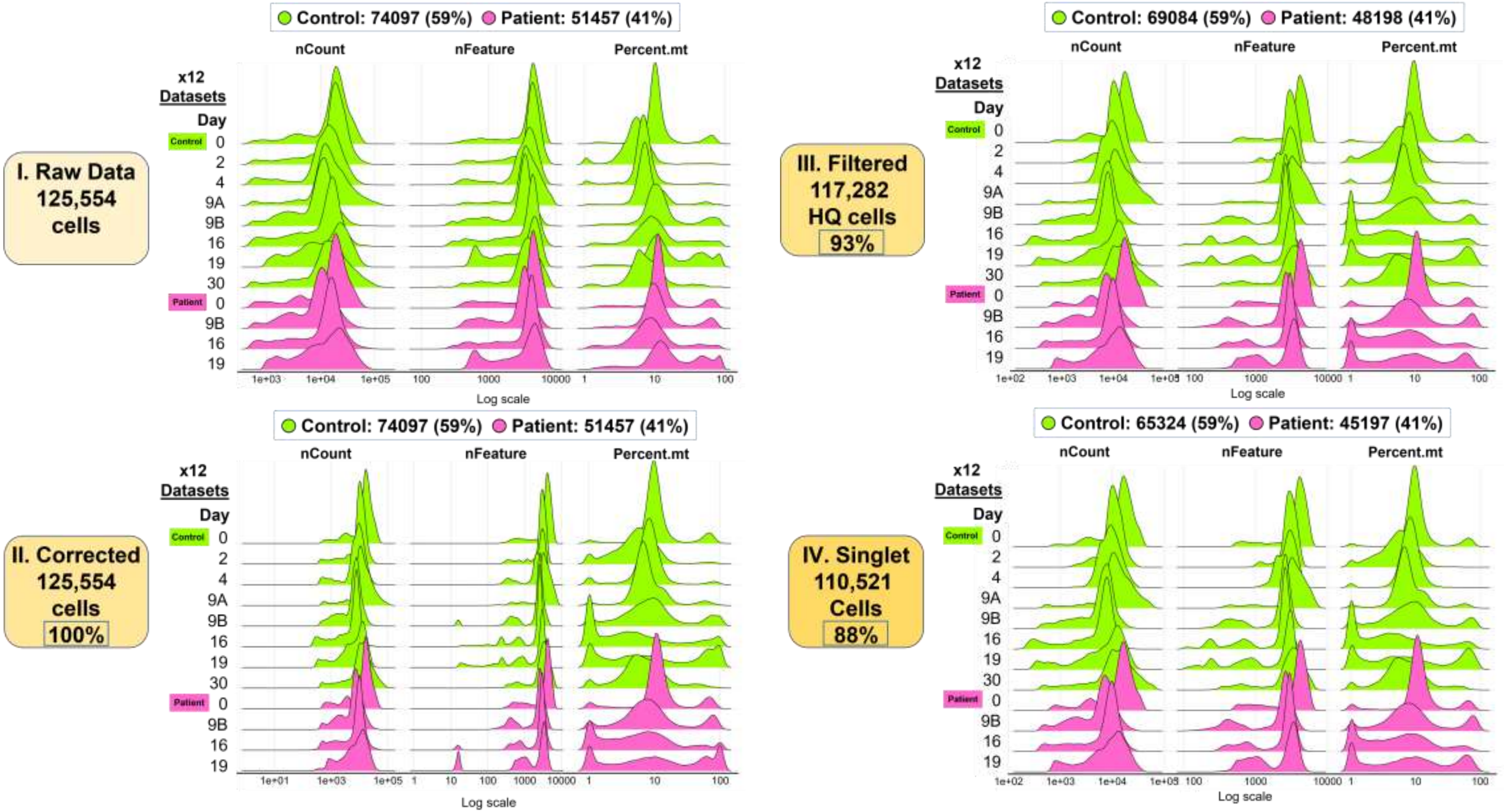

**Fig. S4 Processing**

### D. Merged Data for All Single Sample Samples: QC Covariate Plots by Cluster

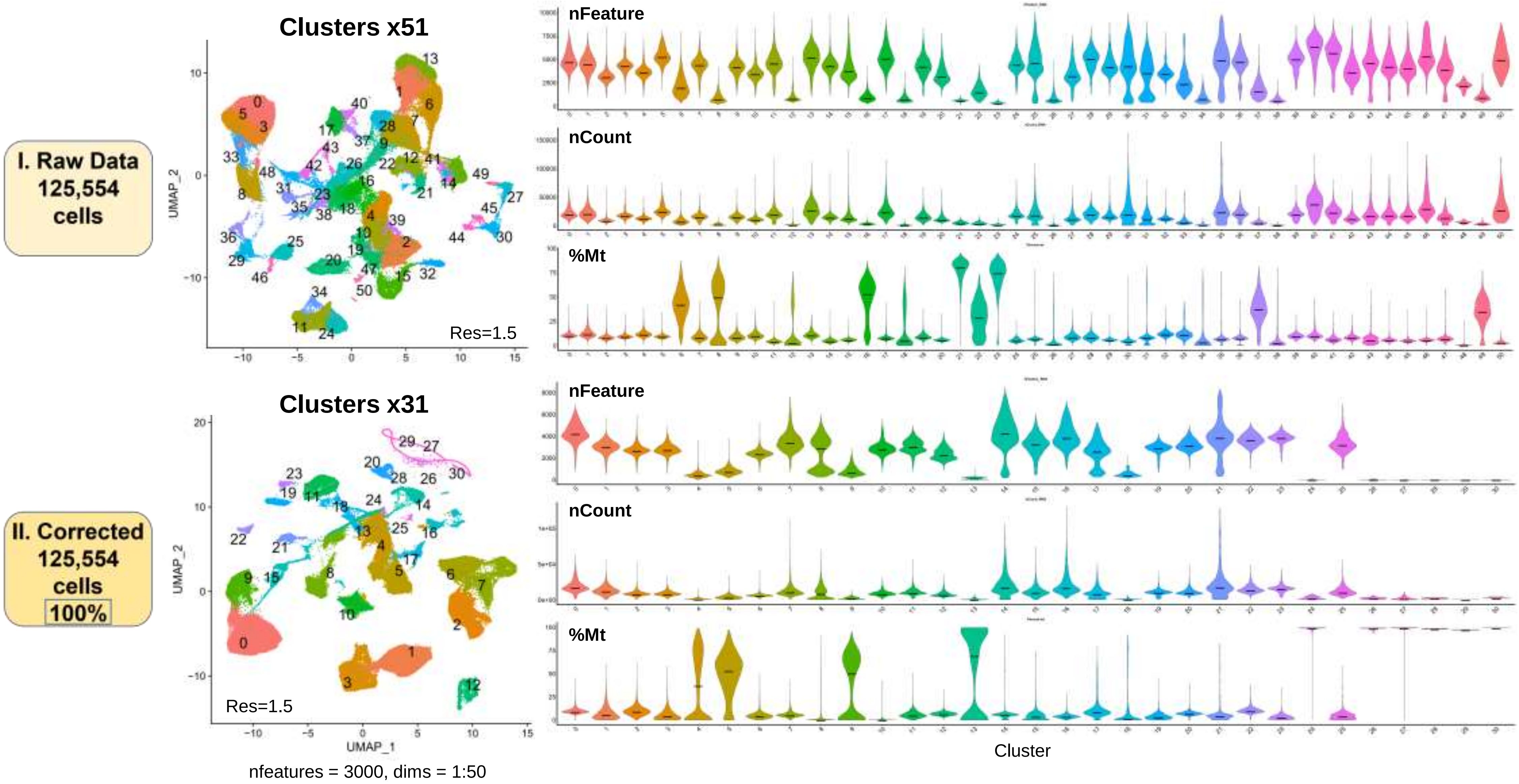

**Fig. S4 Processing**

**III. Filtered 117,282 HQ cells 93%**

**IV. Singlet 110,521 Cells 88%**

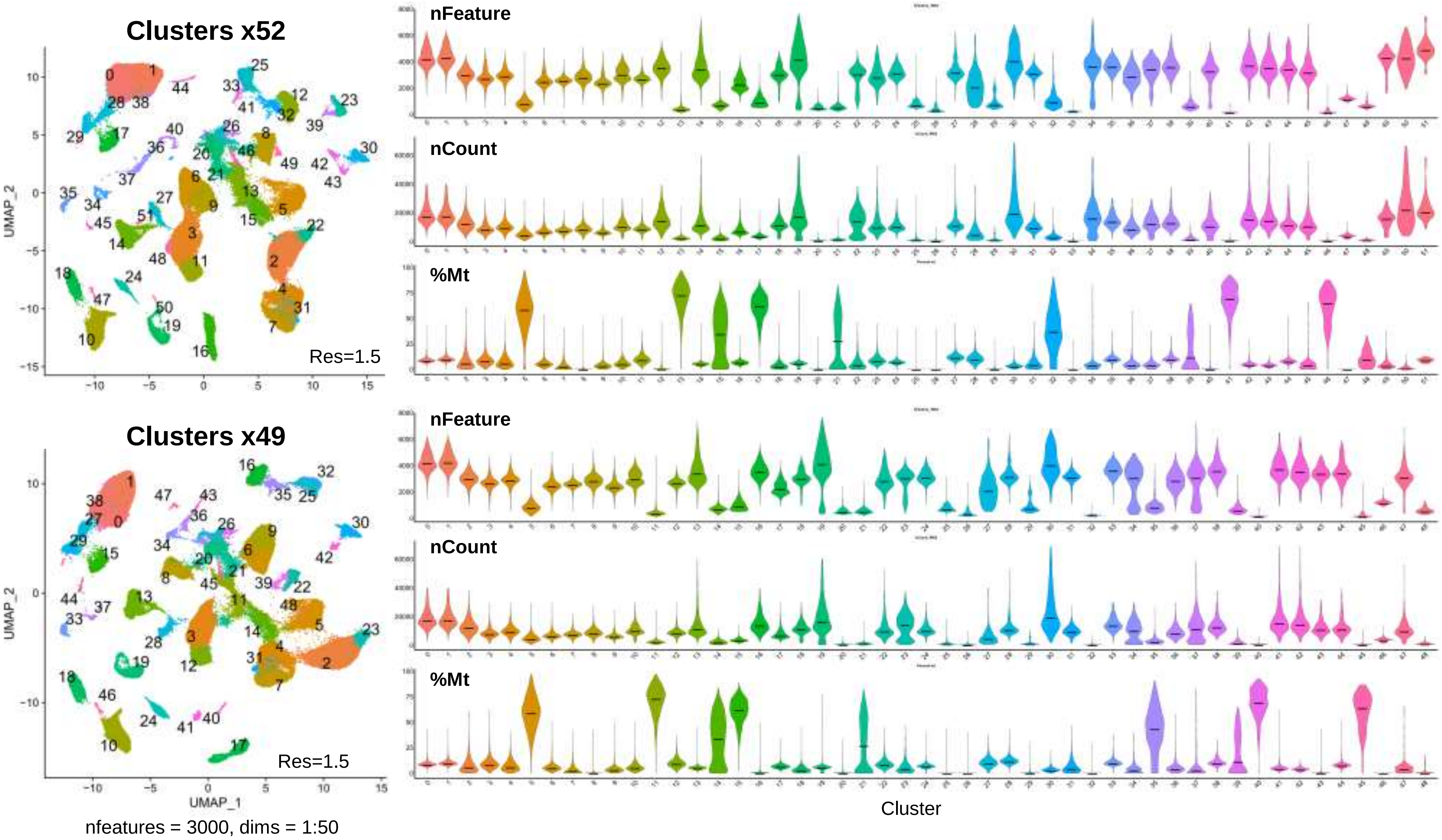

**Fig. S4 Processing**

#### E. Background RNA: Removal Using SoupX

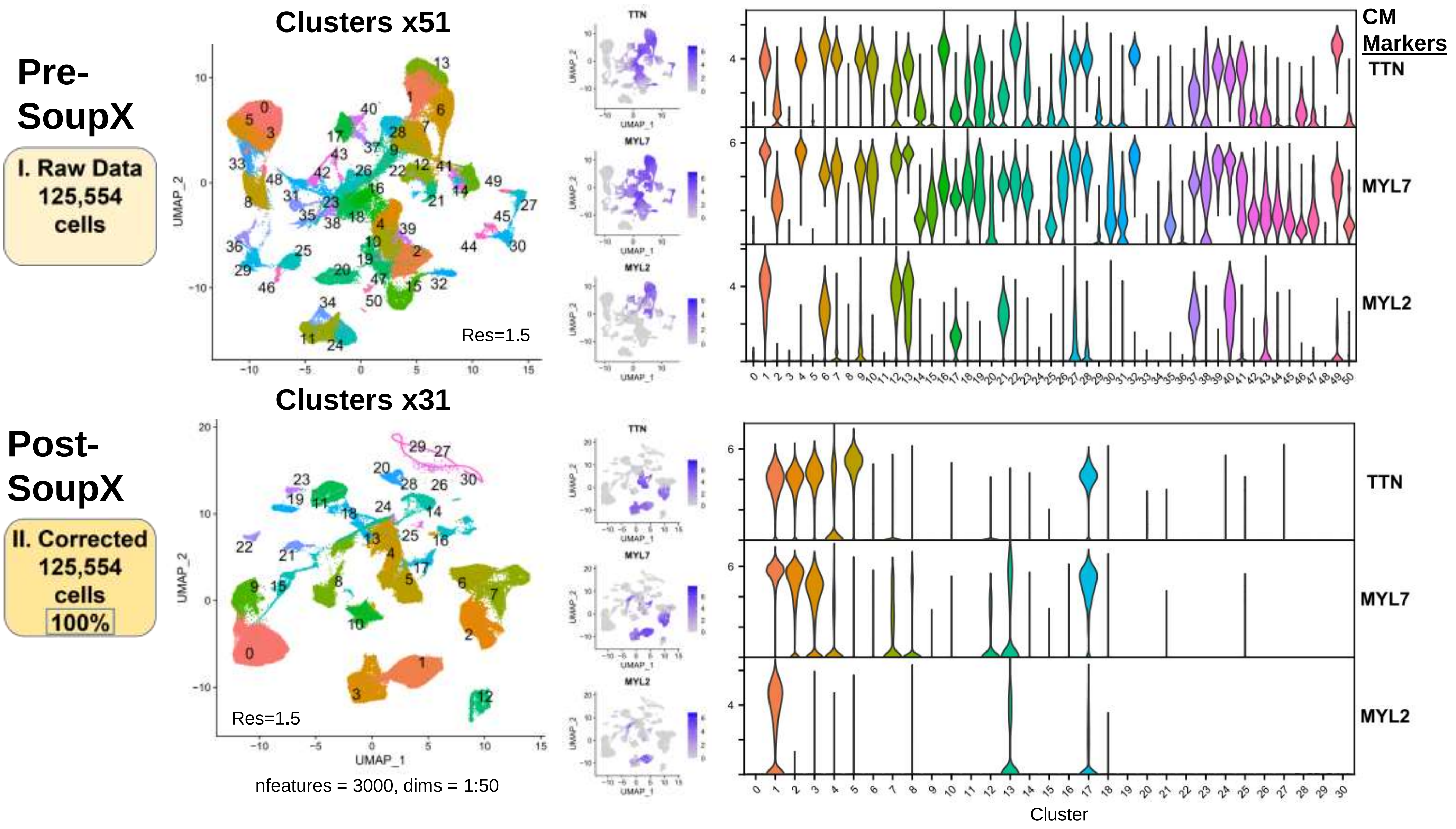

**Fig. S4 Processing**

#### F. Cell Quality: Removal of Low-Quality Cells Using Seurat

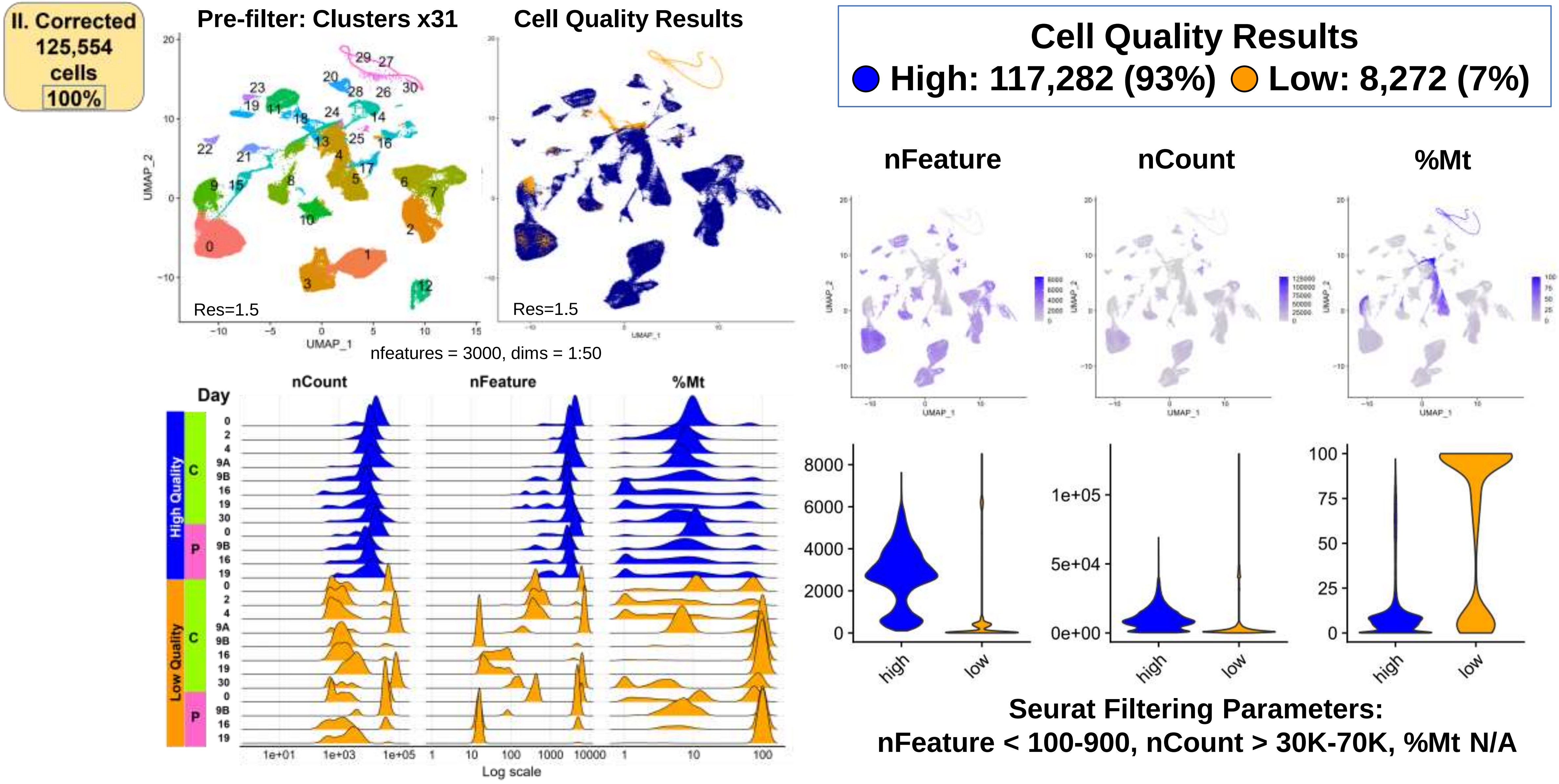

#### G. Doublets: Identification and Removal Using DoubletFinder (DF)

**III. Filtered**
**117,282**
**HQ cells**
**93%**

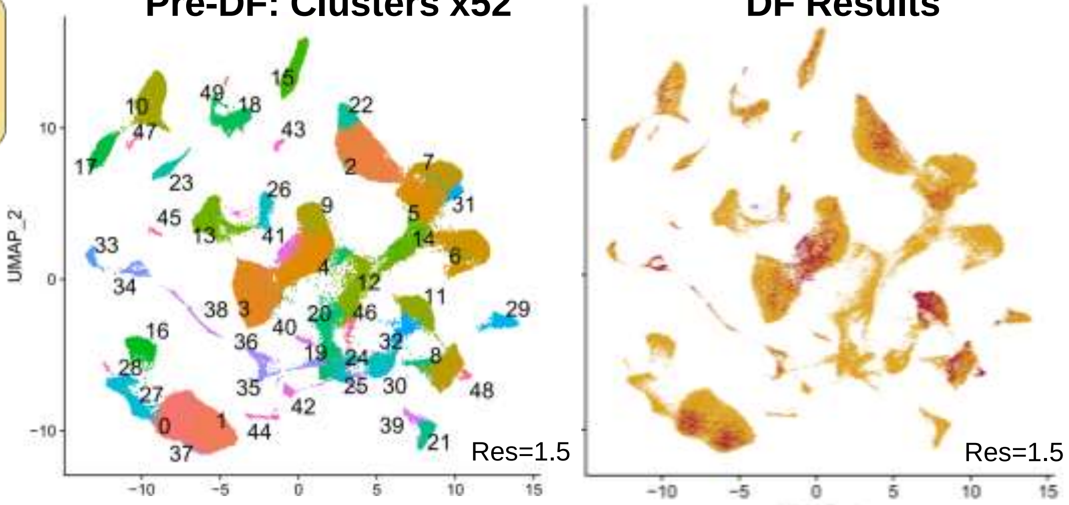


nfeatures = 3000, dims = 1:50

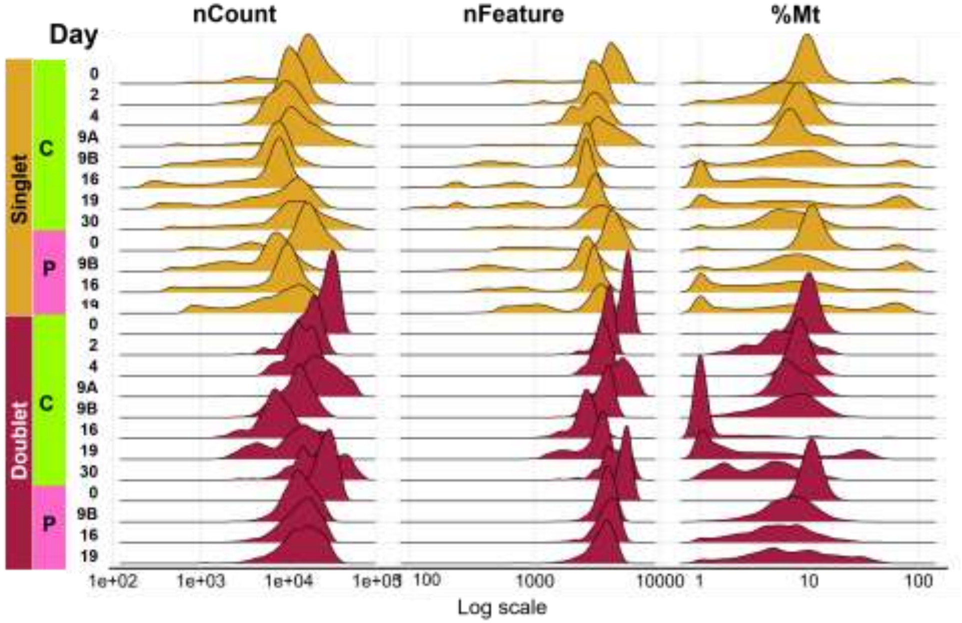


**DoubletFinder Results**
**Doublet: 6761 (6%)**
**Singlet: 110521 (94%)**

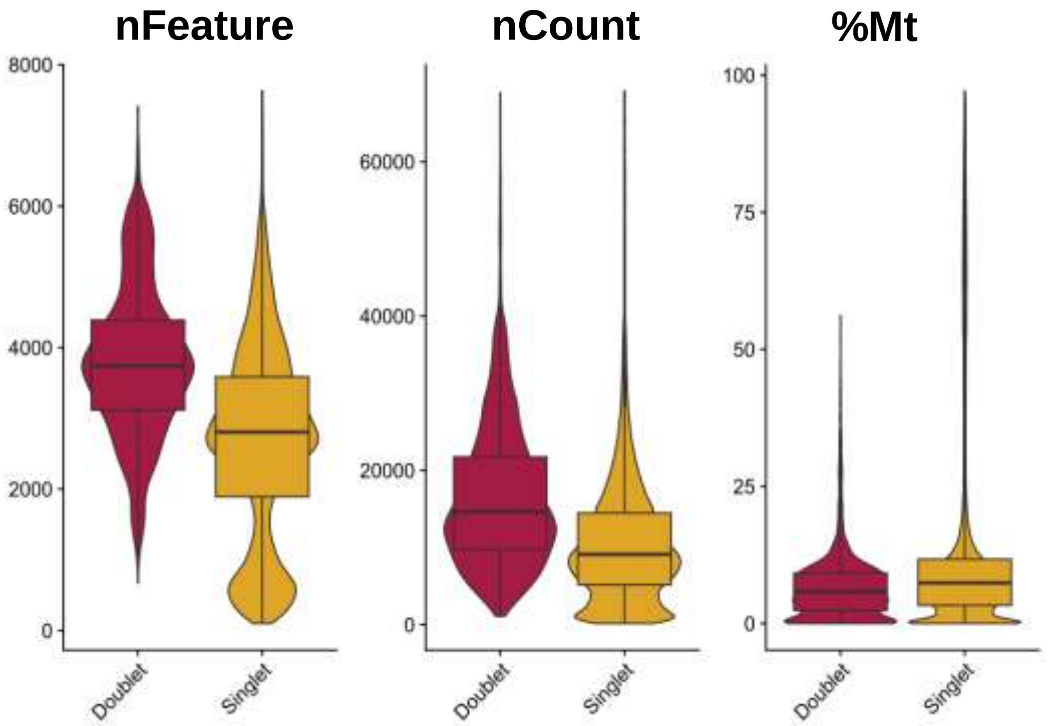


**DF Parameters:**
**pK= 0.005 - 0.30 , Exp= 0.04 - 0.10**

Fig. S5 Annotation

### Workflow Step-II: Clustering and Annotation

#### A. Summary: Single Sample Data for Cell Annotation

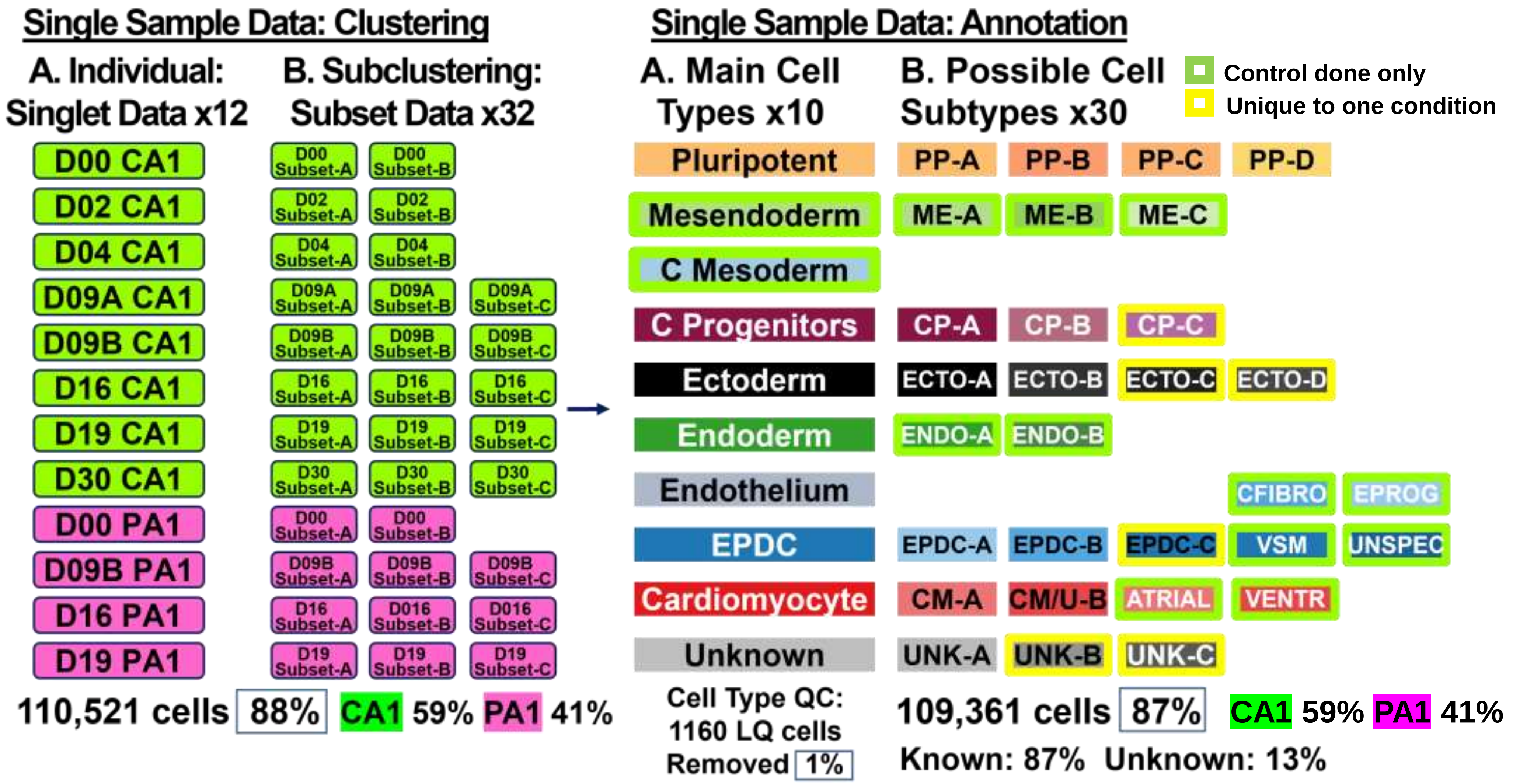

**Fig. S5 Annotation**

#### B. Summary: Merged Single Sample Data for Cell Annotation (110,521 Total Cells)

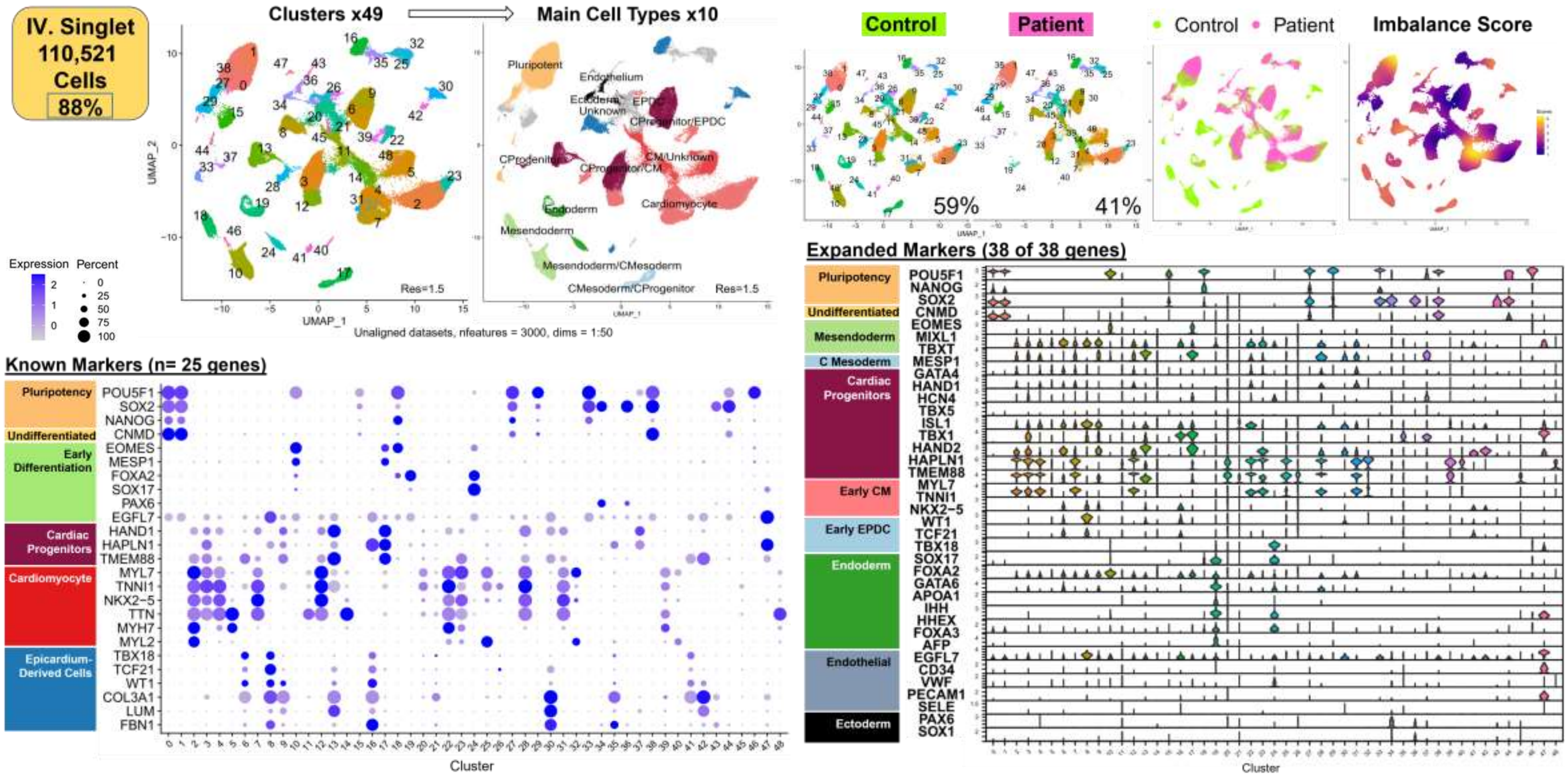

Fig. S6 Clustering

### Workflow Step-II: Single Sample Data Results

#### Individual Analyses of Singlet Data for Main Cell Types

##### A. Summary: Raw Data to Annotated Clusters

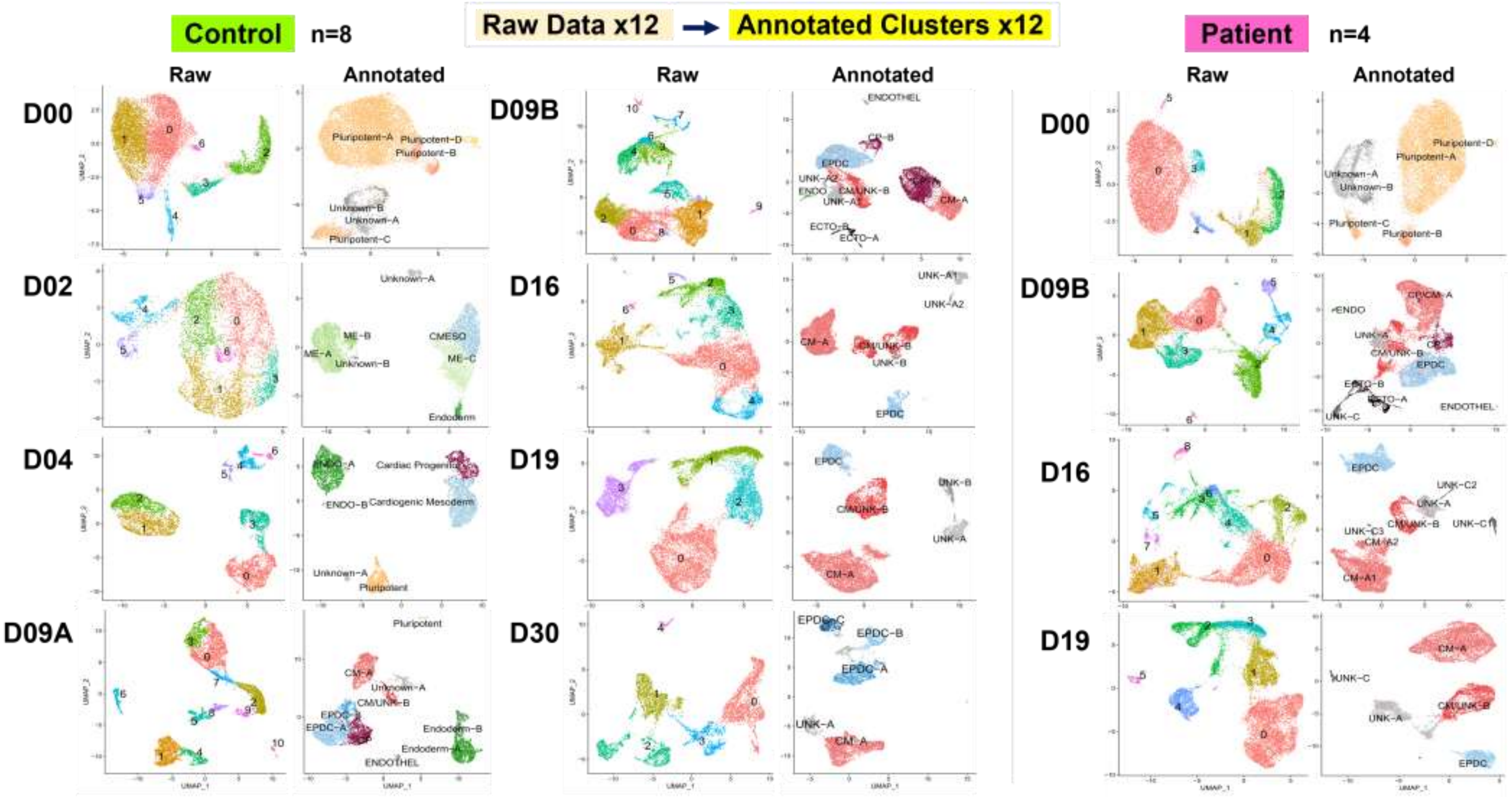

**Fig. S6 Clustering**

#### B. Single Sample Data Analyses: Control Samples (n=8)

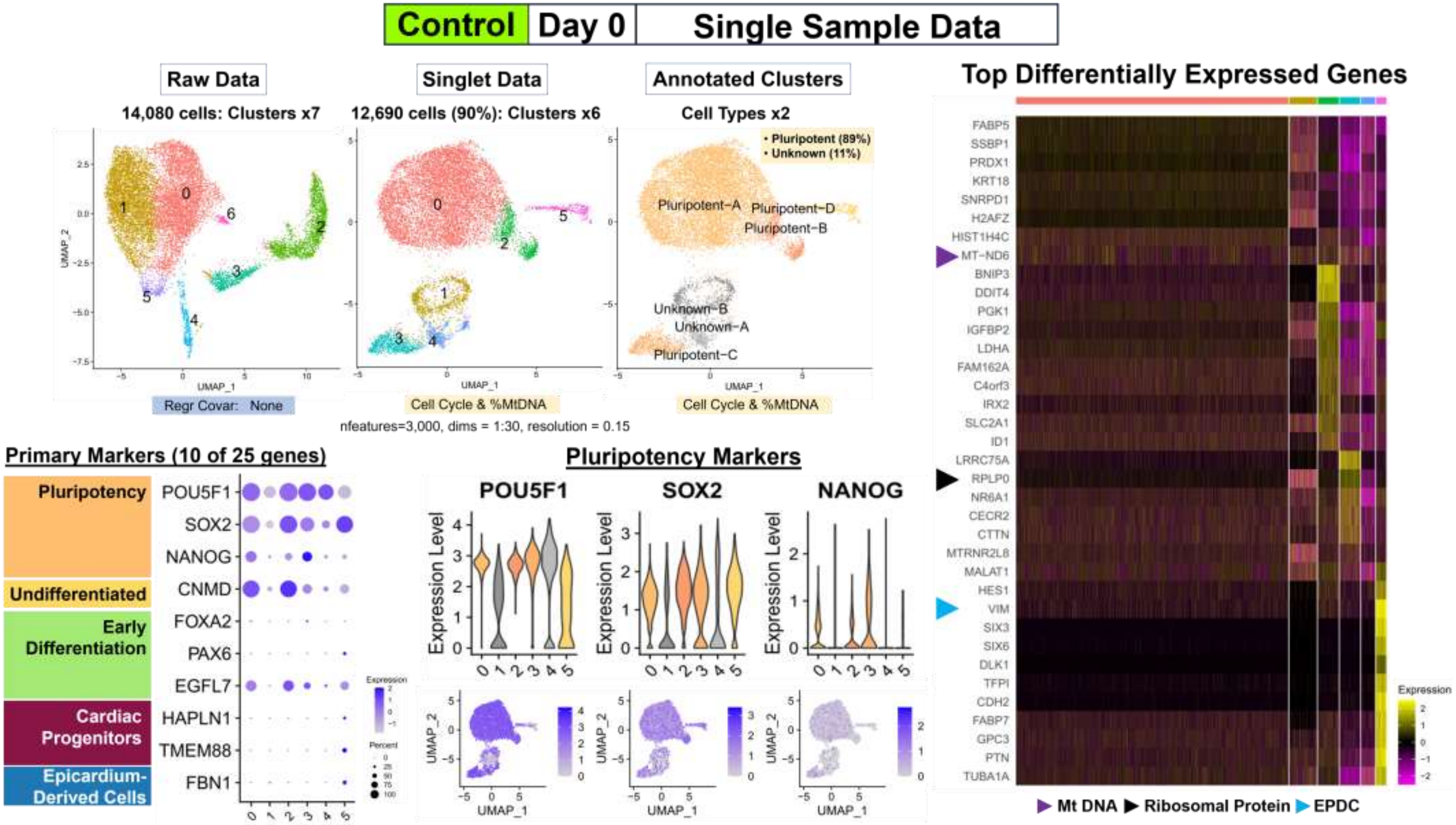

**Fig. S6 Clustering**

**Control | Day 2 | Single Sample Data**

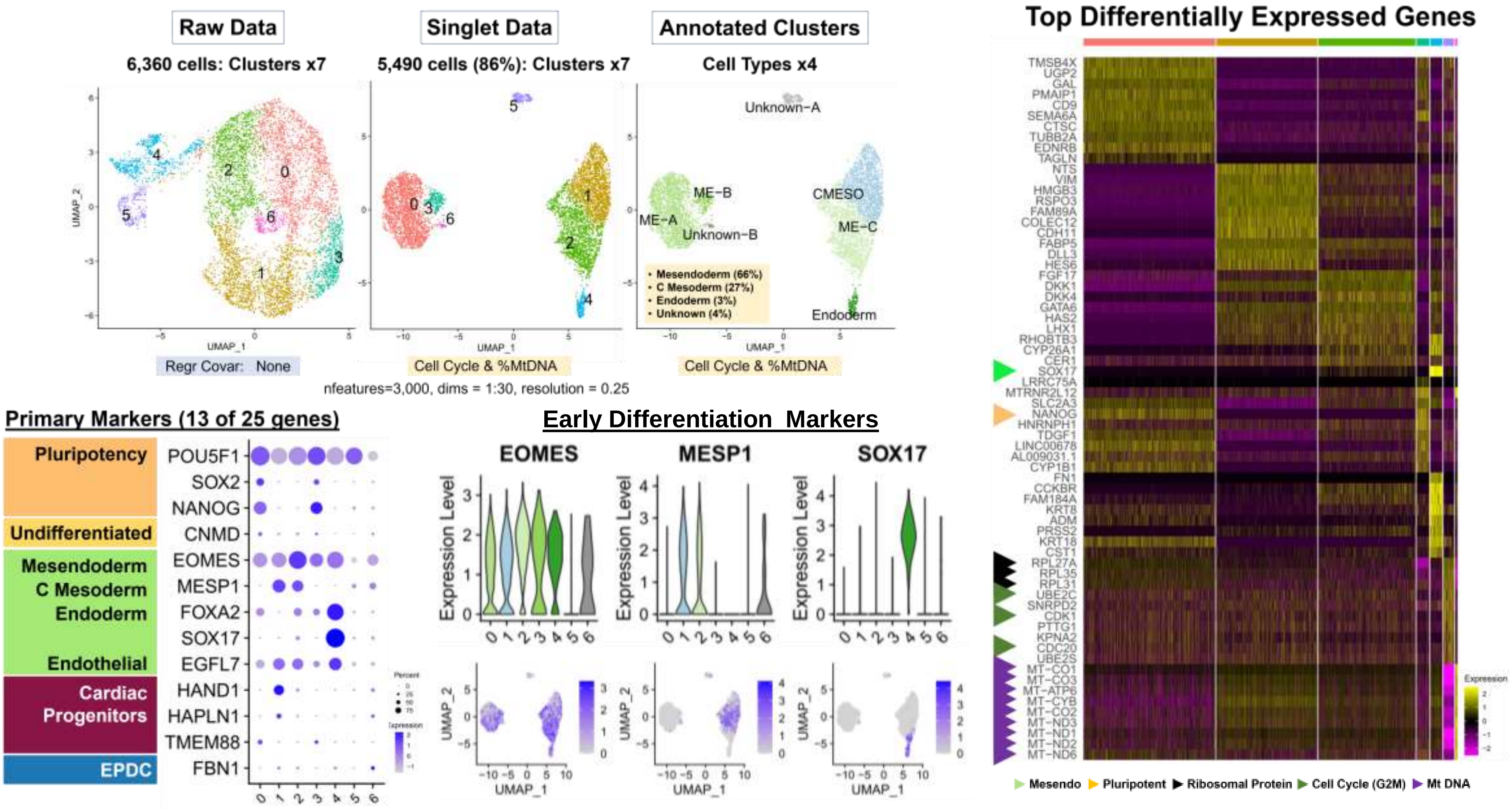

**Fig. S6 Clustering**

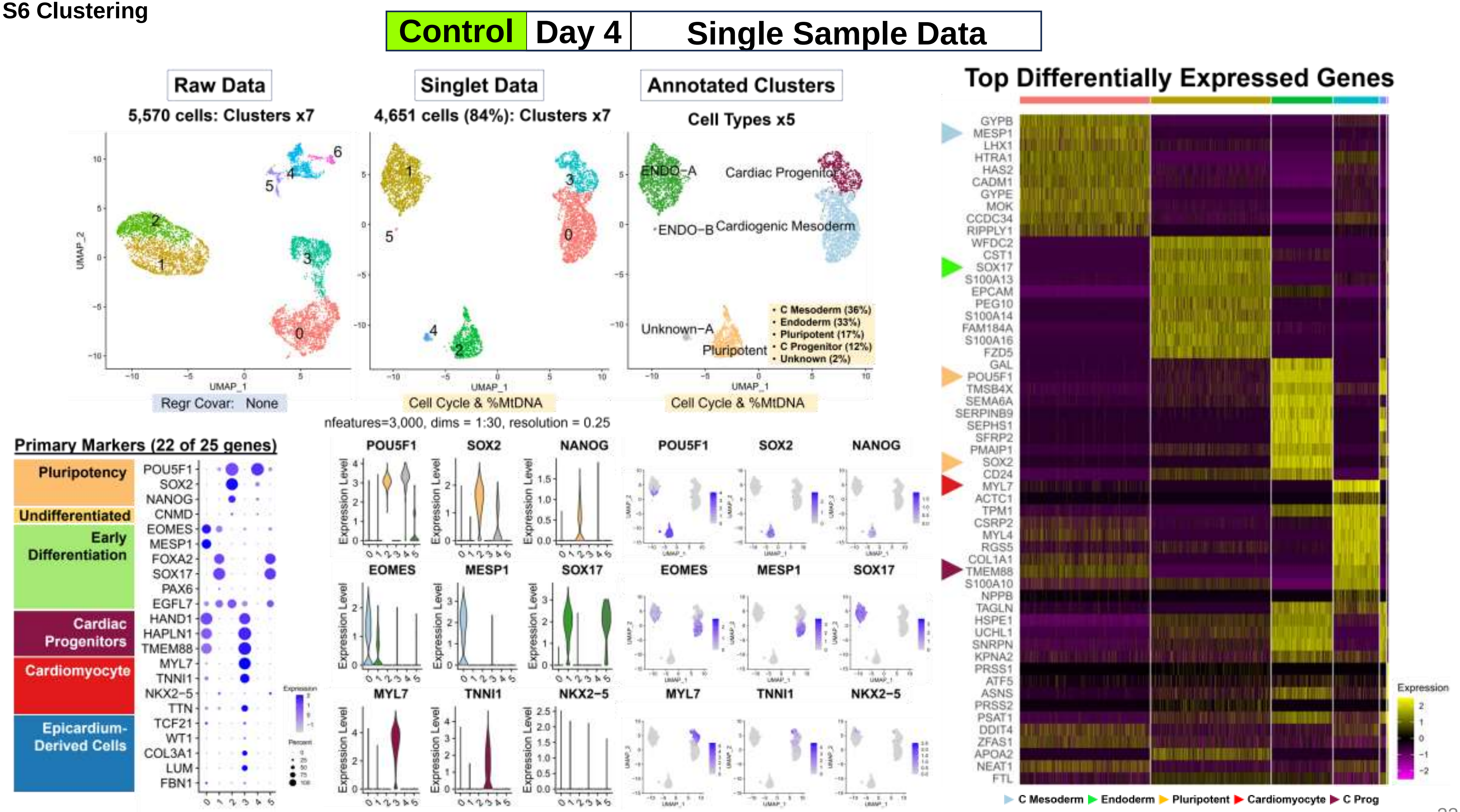

**Fig. S6 Clustering**

**Control | Day 9A | Single Sample Data**

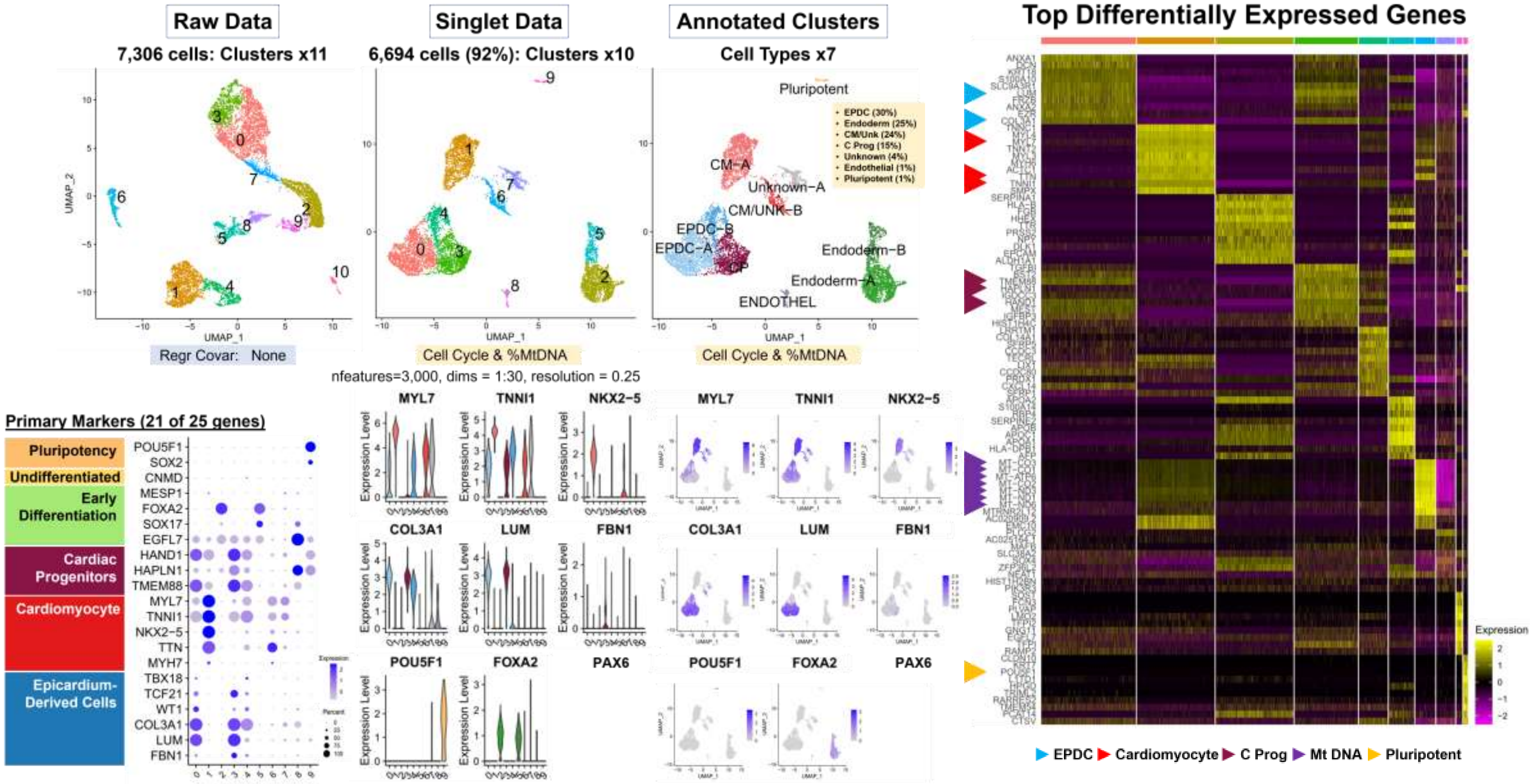

**Fig. S6 Clustering**

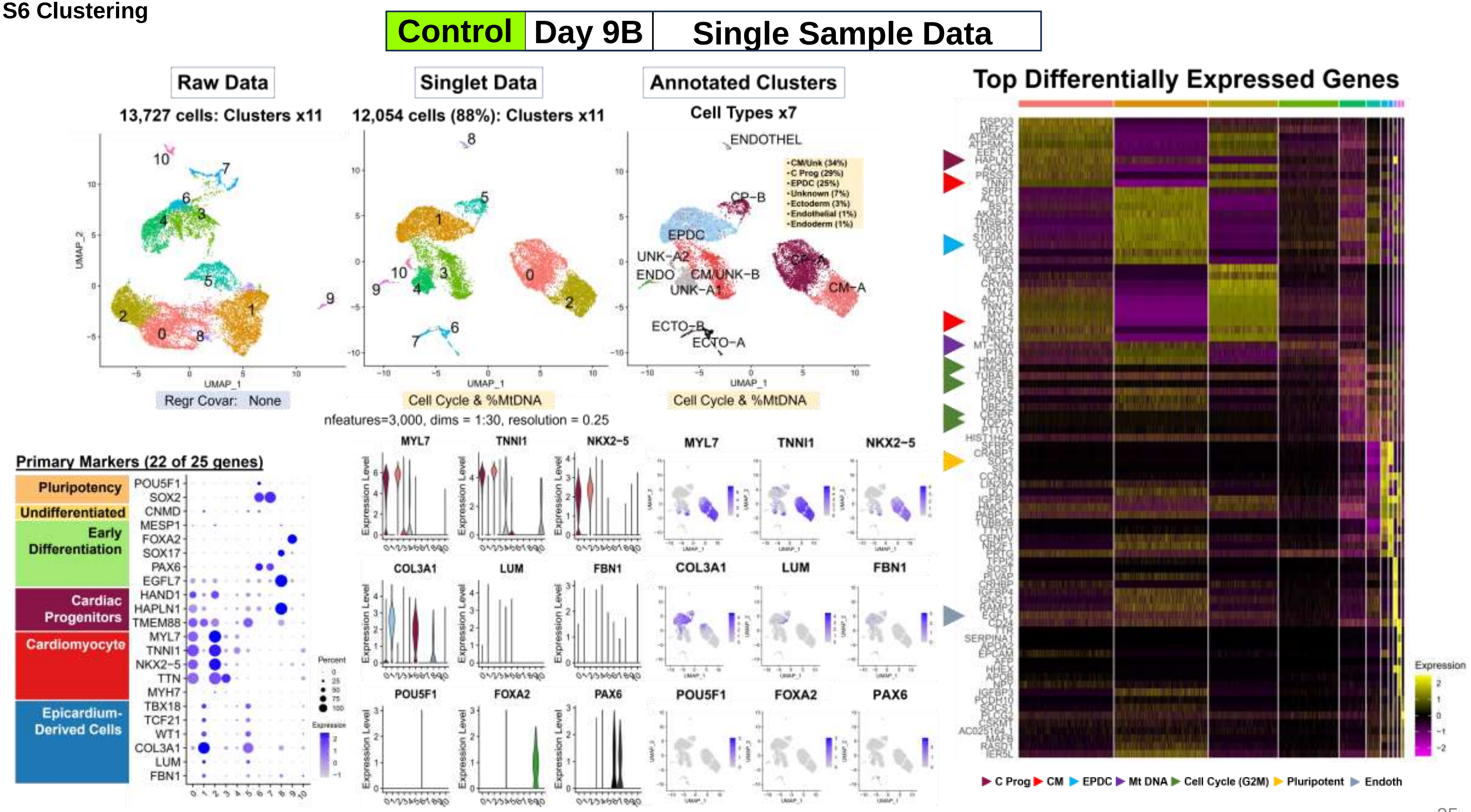

Fig. S6 Clustering

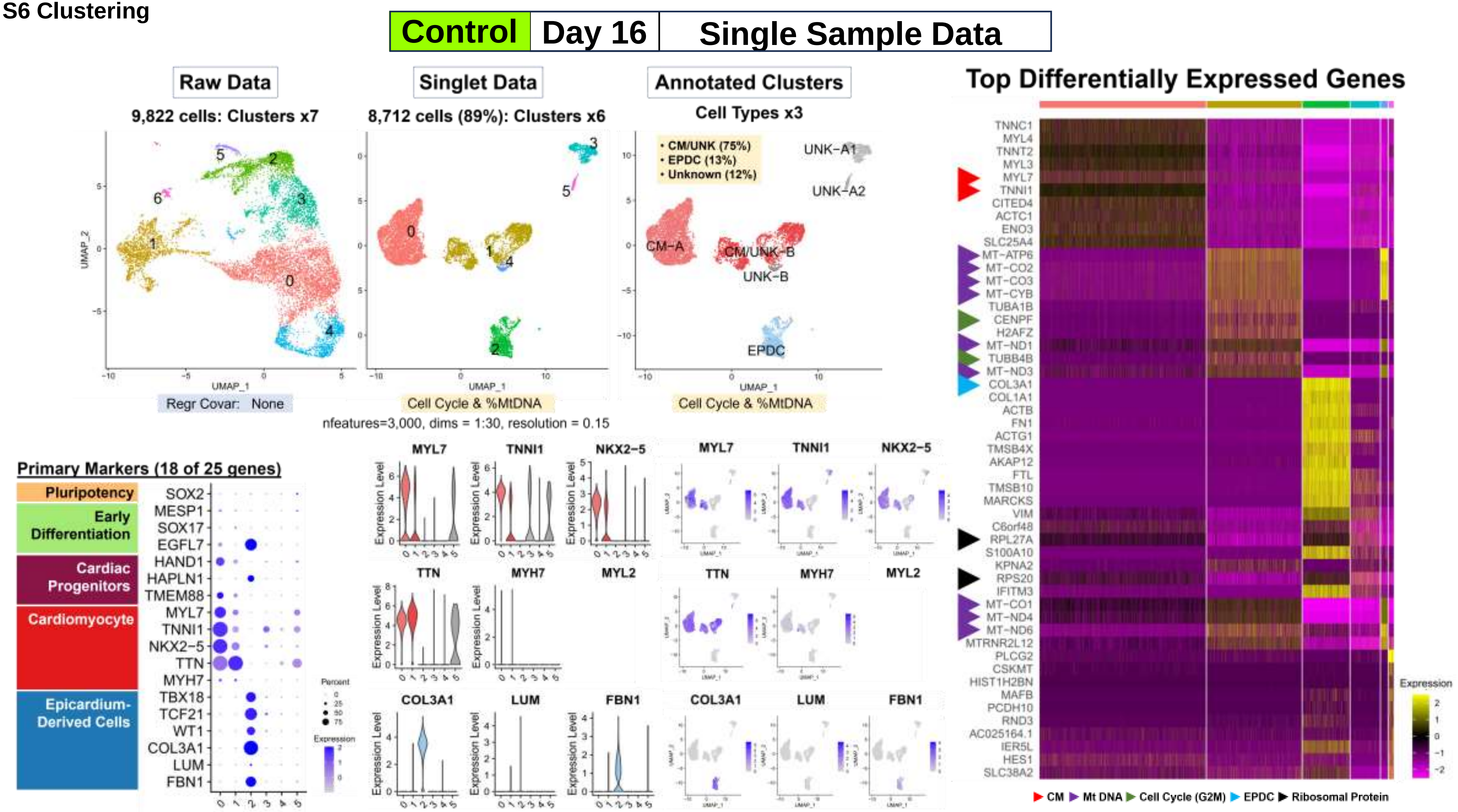

Fig. S6 Clustering

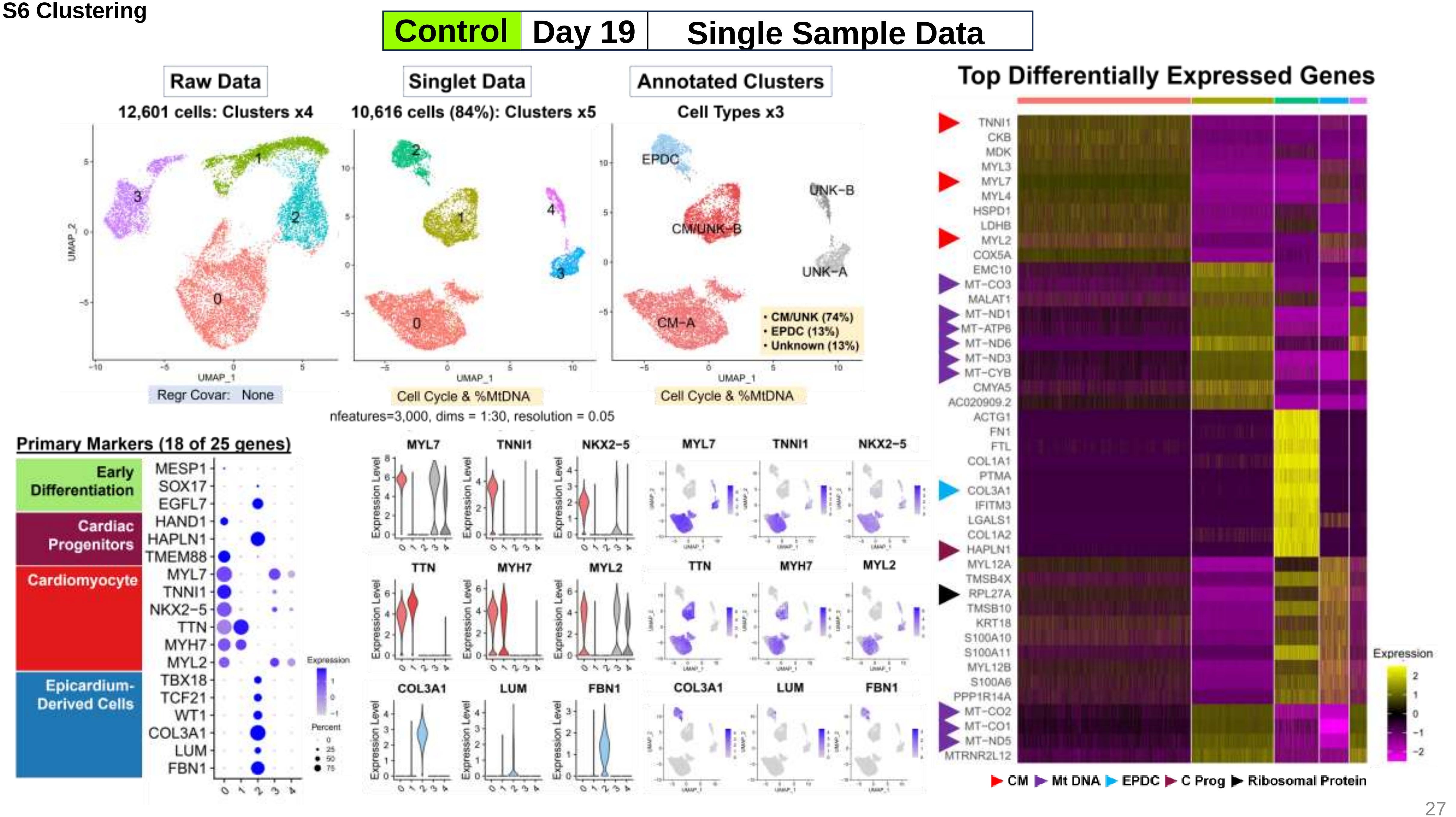

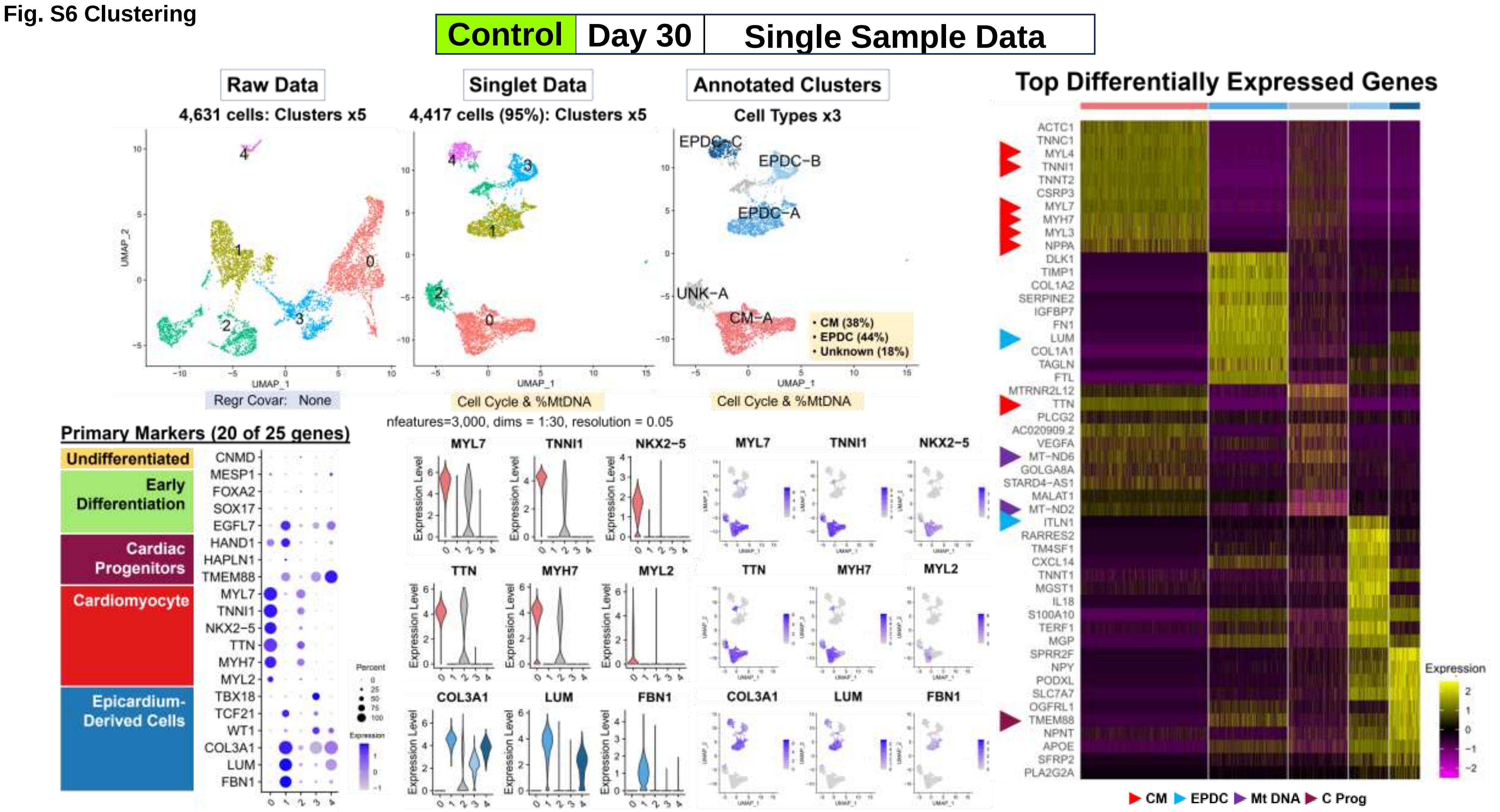
Fig. S6 Clustering
Control
Day 30
Single Sample Data
Raw Data
4,631 cells: Clusters x5
Singlet Data
4,417 cells (95%): Clusters x5
Annotated Clusters
Cell Types x3
Top Differentially Expressed Genes
EPDC-C
EPDC-B
EPDC-A
UNK-A
CM-A
• CM (38%)
• EPDC (44%)
• Unknown (18%)
Regr Covar: None
Cell Cycle & %MtDNA
Cell Cycle & %MtDNA
nfeatures=3,000, dims = 1:30, resolution = 0.05
Primary Markers (20 of 25 genes)
Undifferentiated
Early Differentiation
Cardiac Progenitors
Cardiomyocyte
Epicardium-Derived Cells
CNMD
MESP1
FOXA2
SOX17
EGFL7
HAND1
HAPLN1
TMEM88
MYL7
TNNI1
NKX2-5
TTN
MYH7
MYL2
TBX18
TCF21
WT1
COL3A1
LUM
FBN1
Percent
Expression
MYL7
TNNI1
NKX2-5
TTN
MYH7
MYL2
COL3A1
LUM
FBN1
Expression Level
ACTC1
TNNC1
MYL4
TNNI1
TNNT2
CSRP3
MYL7
MYH7
MYL3
NPPA
DLK1
TIMP1
COL1A2
SERPINE2
IGFBP7
FN1
LUM
COL1A1
TAGLN
FTL
MTRNR2L12
TTN
PLCG2
AC020909.2
VEGFA
MT-ND6
GOLGA8A
STARD4-AS1
MALAT1
MT-ND2
ITLN1
RARRES2
TM4SF1
CXCL14
TNNT1
MGST1
IL18
S100A10
TERF1
MGP
SPRR2F
NPY
PODXL
SLC7A7
OGFRL1
TMEM88
NPNT
APOE
SFRP2
PLA2G2A
Expression
CM
EPDC
Mt DNA
C Prog

**Fig. S6 Clustering**

#### C. Single Sample Data Analyses: Patient Samples (n=4)

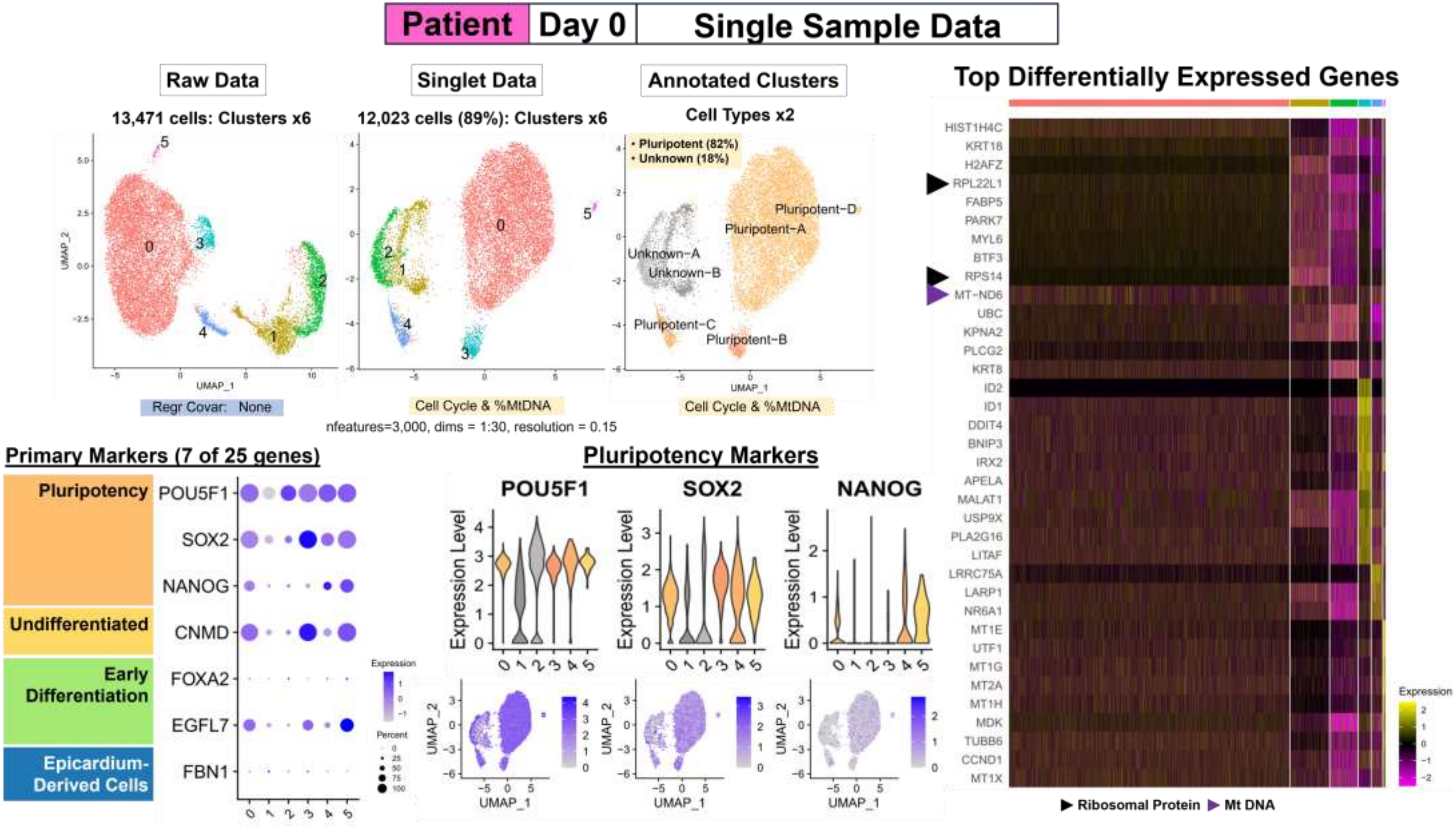

Fig. S6 Clustering

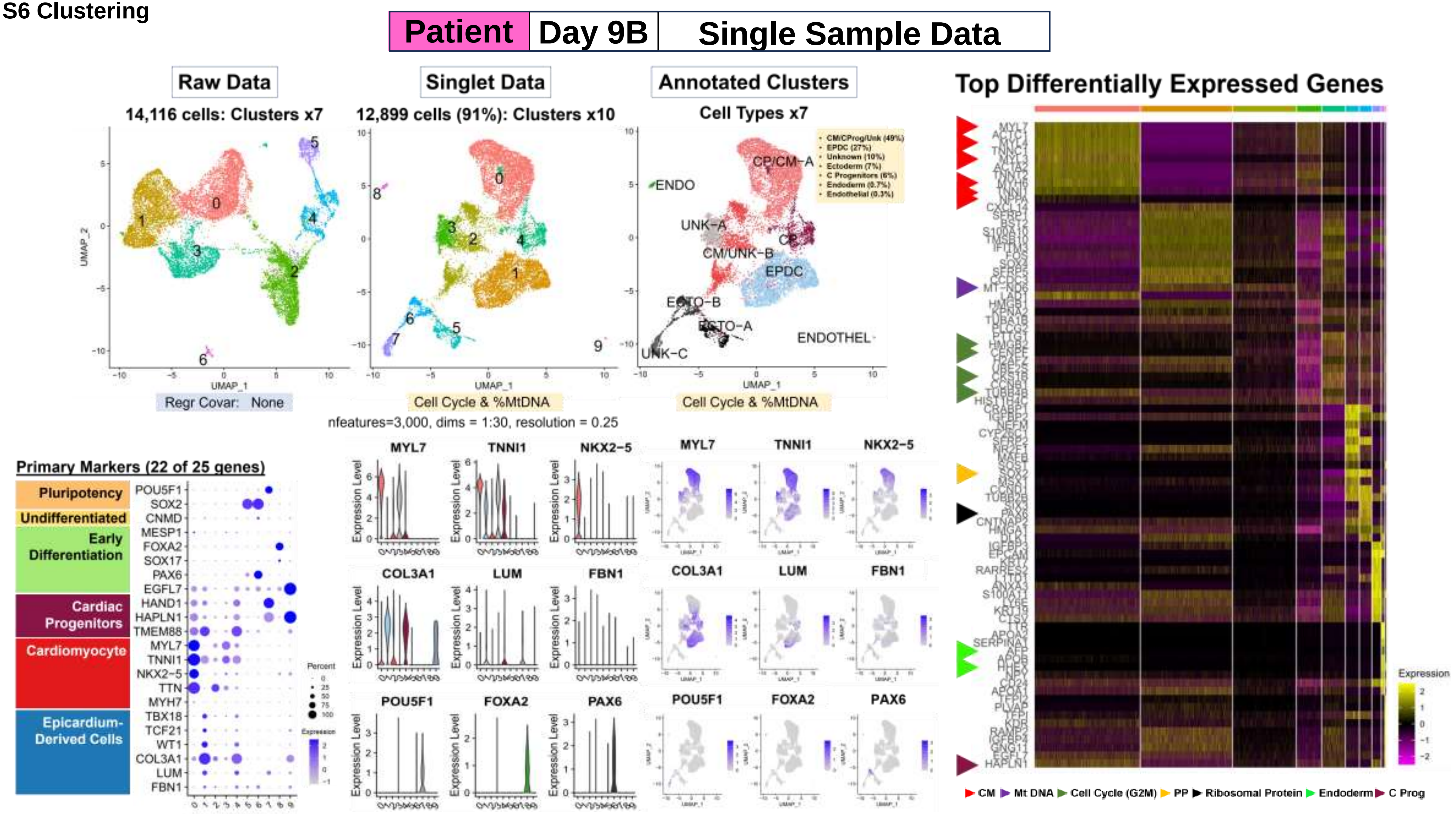

**Fig. S6 Clustering**

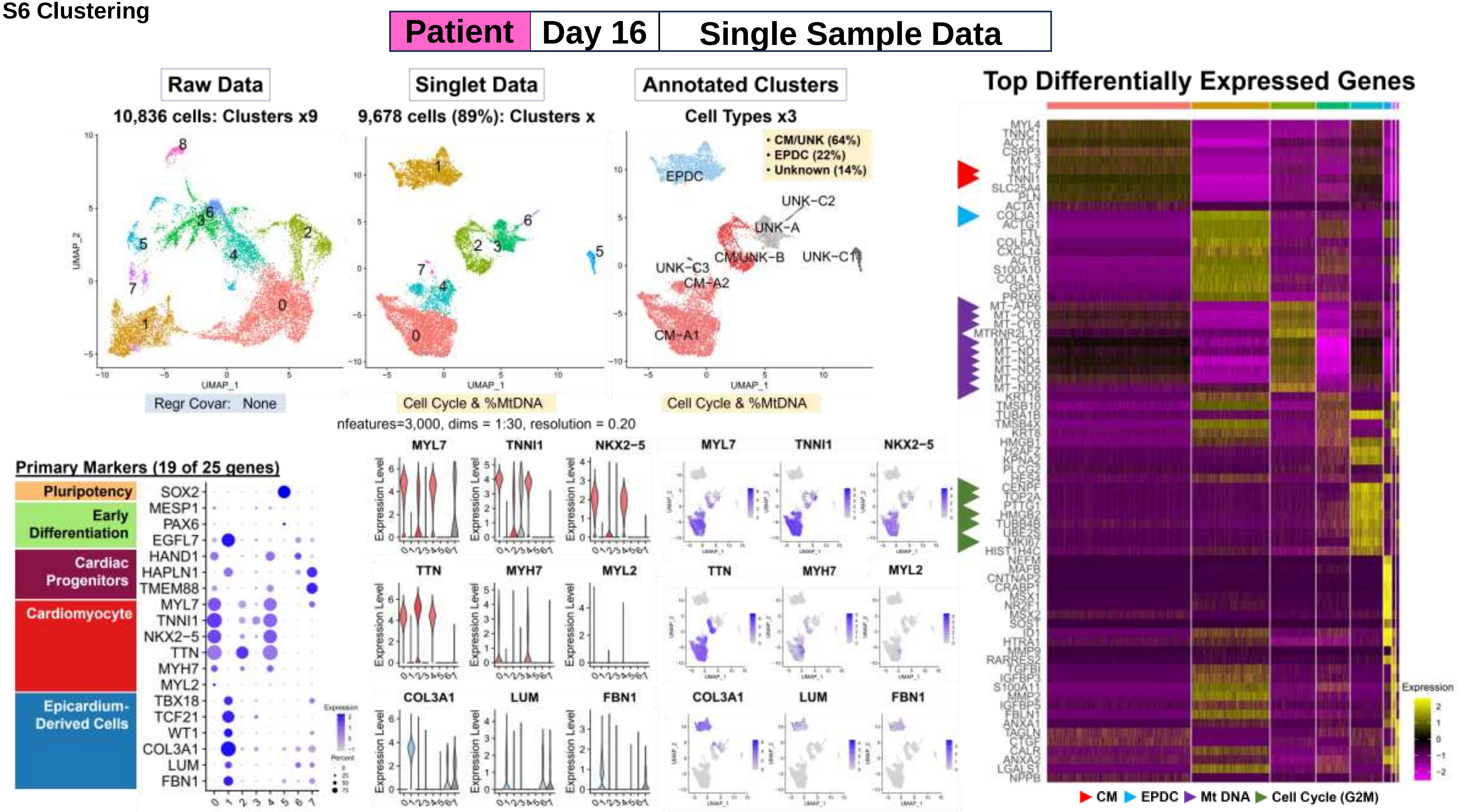

**Fig. S6 Clustering**

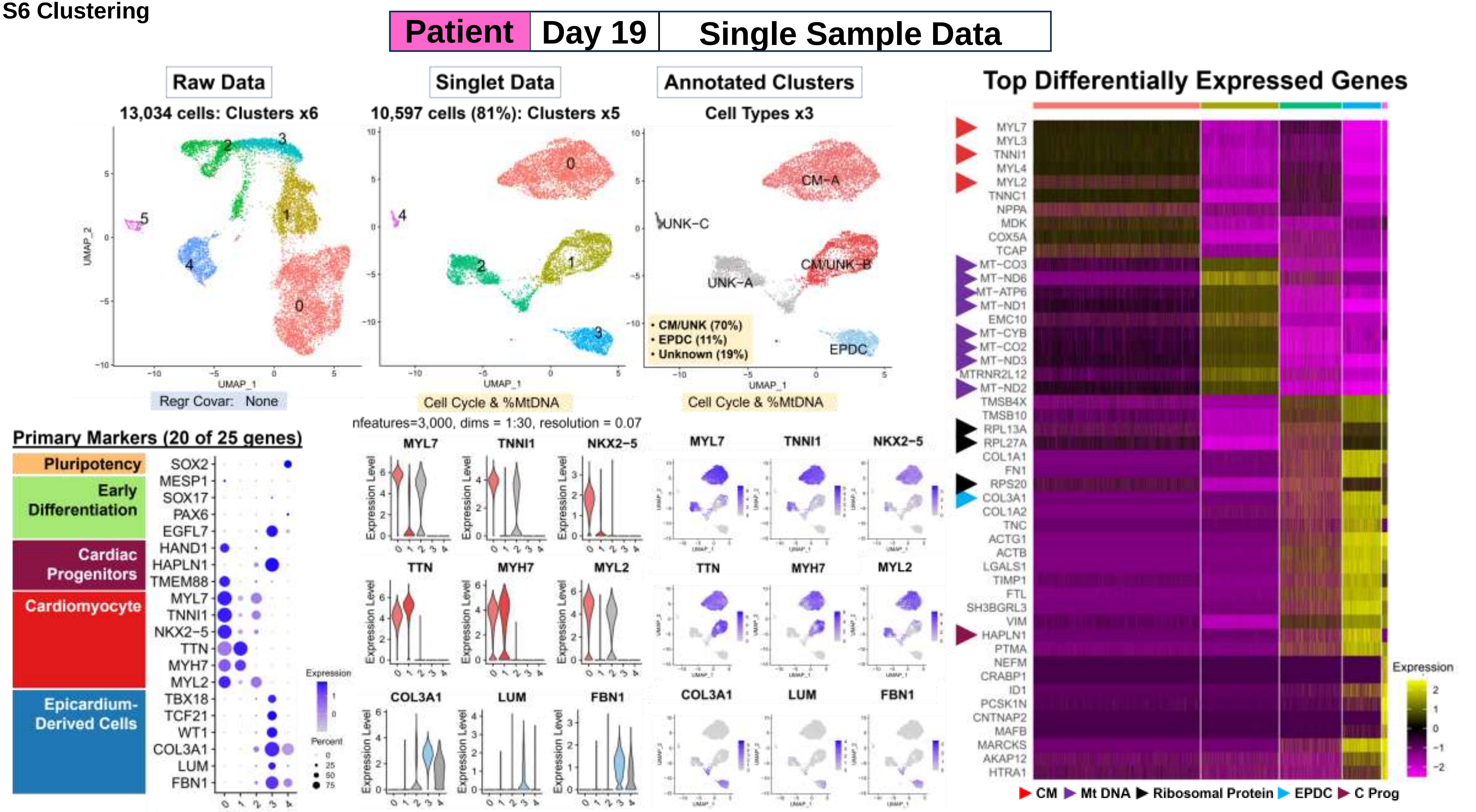

**Fig. S7 Subcluster**

### Workflow Step-II: Single Sample Data Results

#### Subcluster Analyses of Subset Data for Possible Cell Subtypes

##### A. Summary: Annotated Clusters to Annotated Subsets

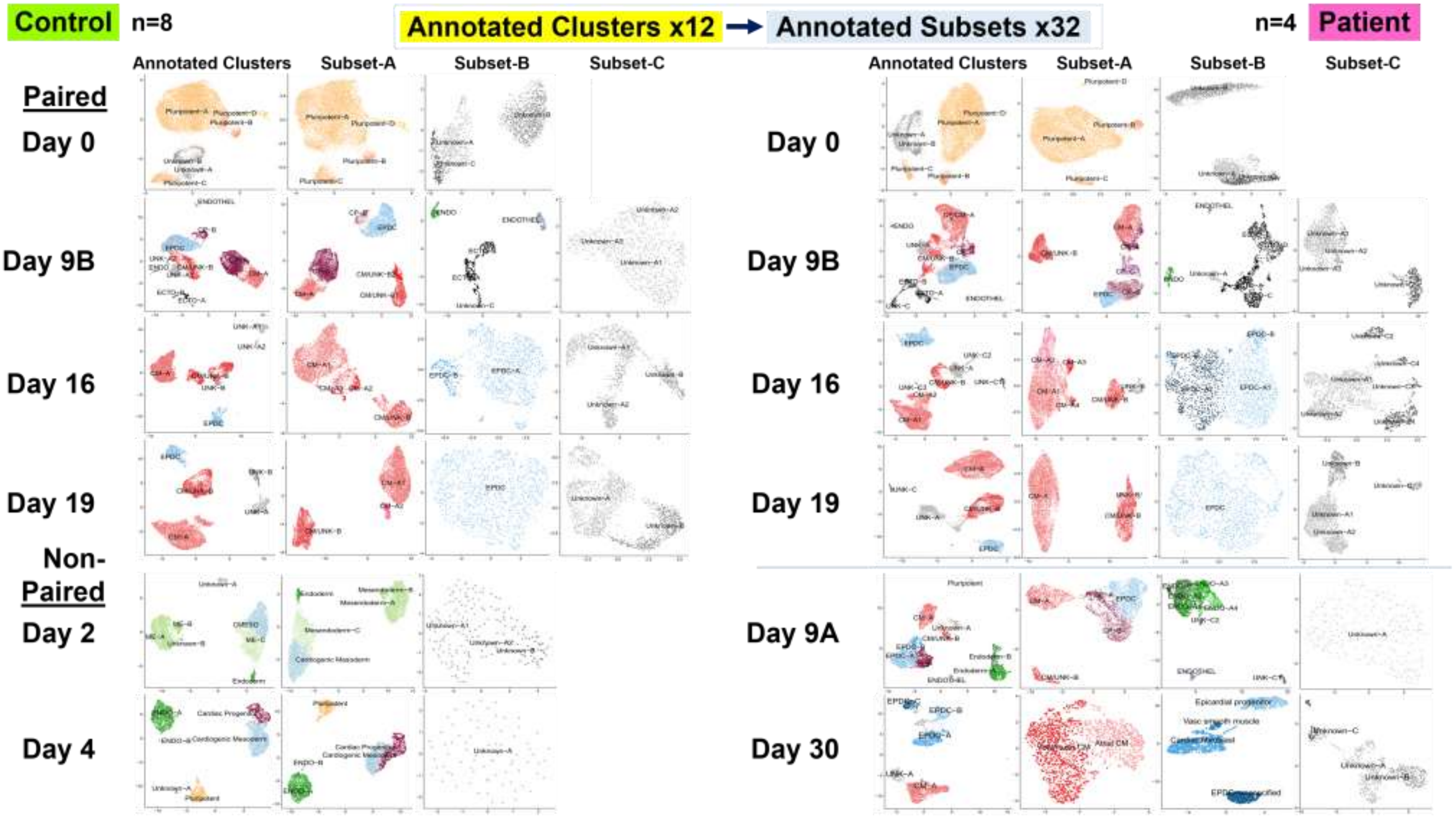

**Fig. S7 Subcluster**

### B. Subcluster Analyses: Control Samples (n=8) to Annotated Subset Data (n=21)

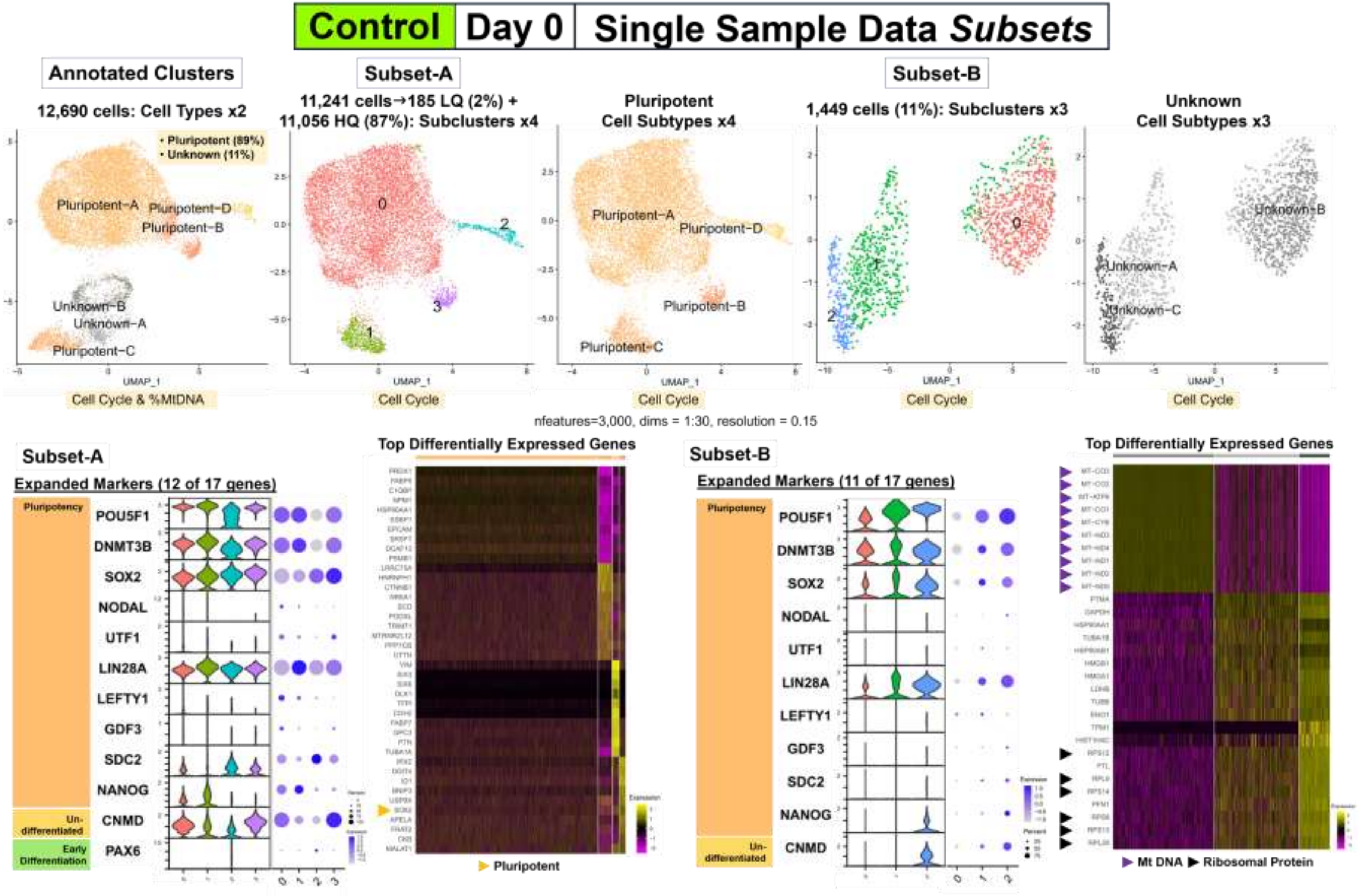

Fig. S7 Subcluster

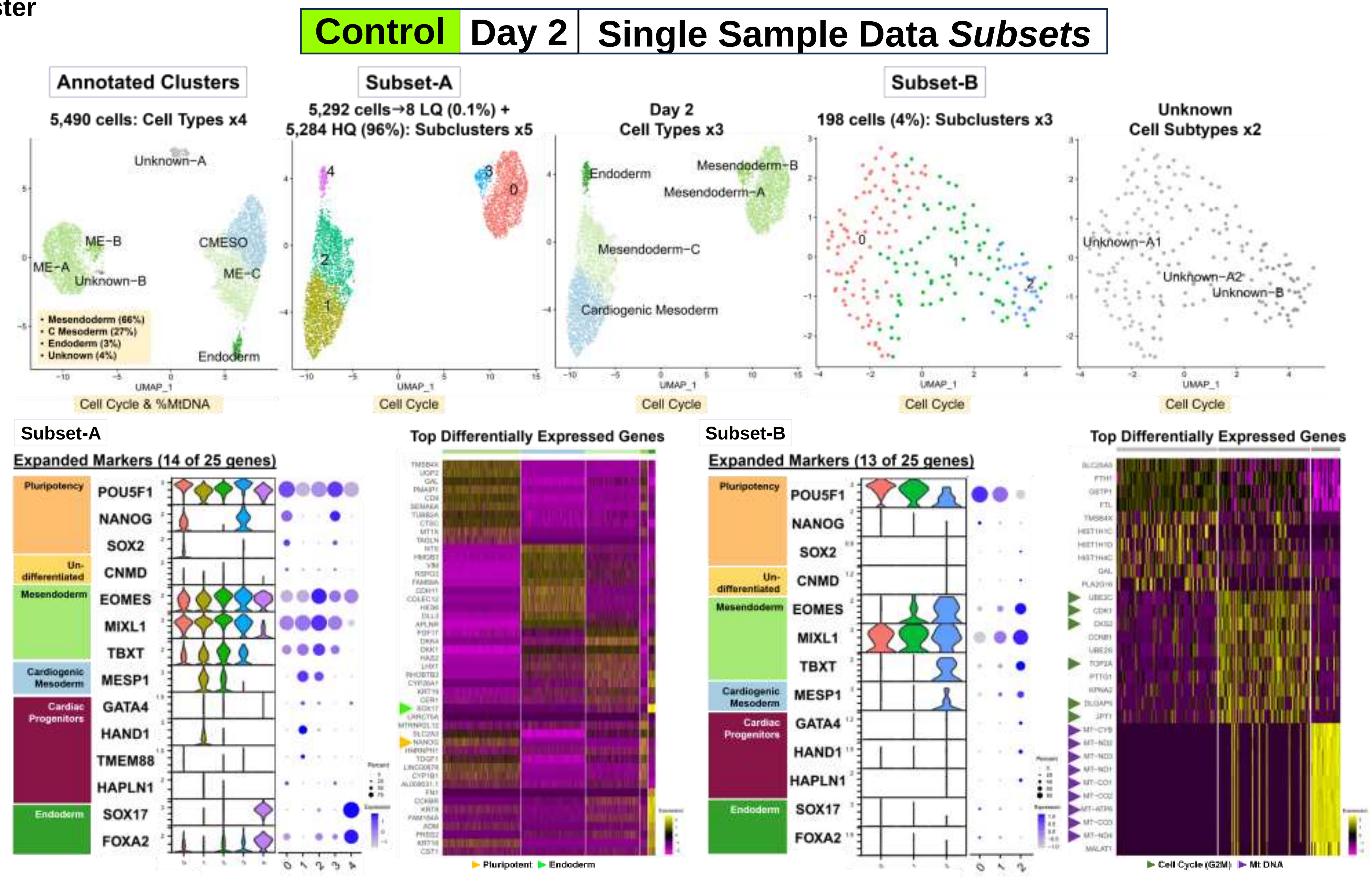

Fig. S7 Subcluster

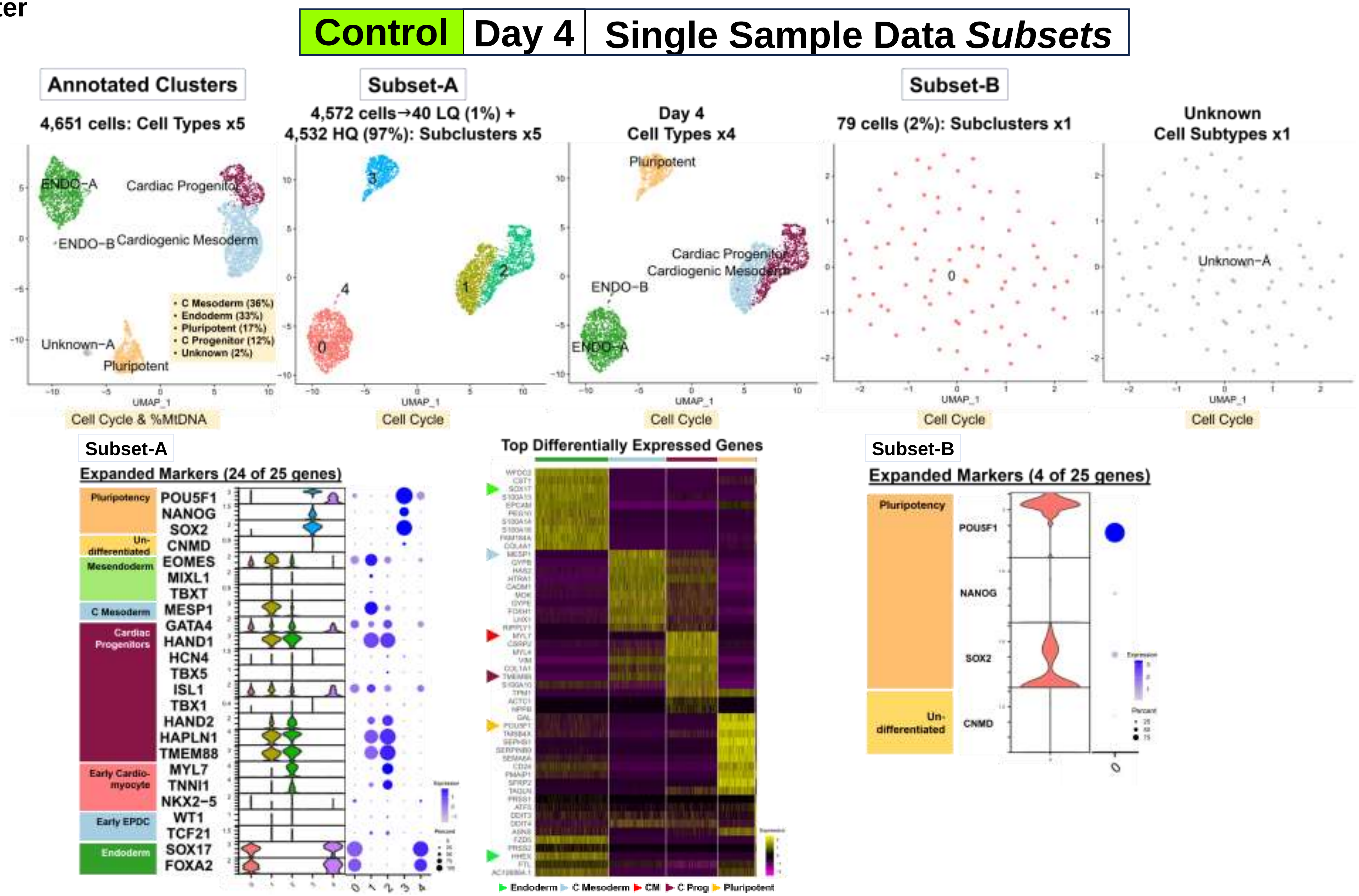

**Fig. S7 Subcluster**

| Control | Day 9A | Single Sample Data *Subsets* |
|---|---|---|

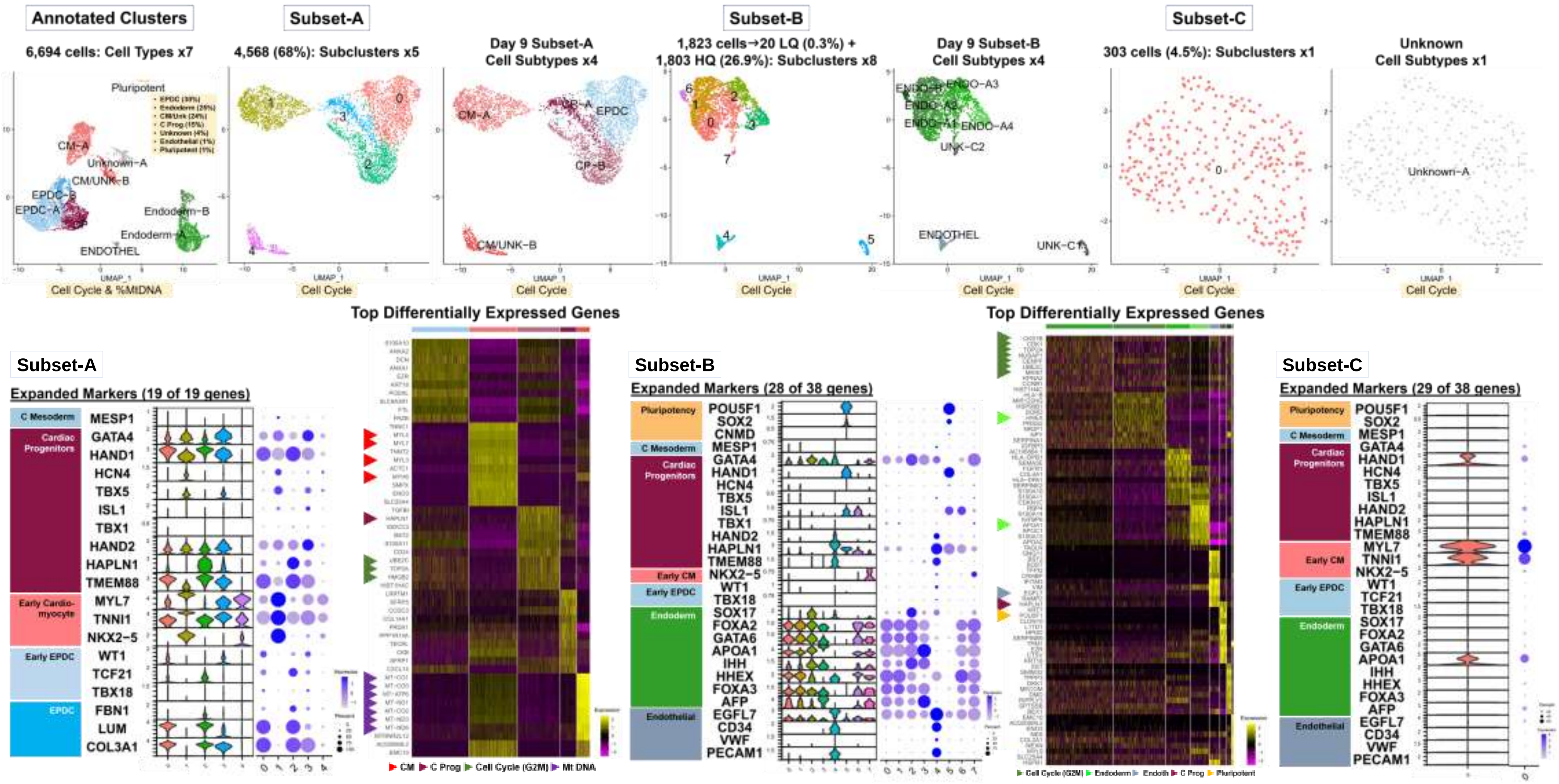

**Fig. S7 Subcluster**

**Control | Day 9B | Single Sample Data *Subsets***

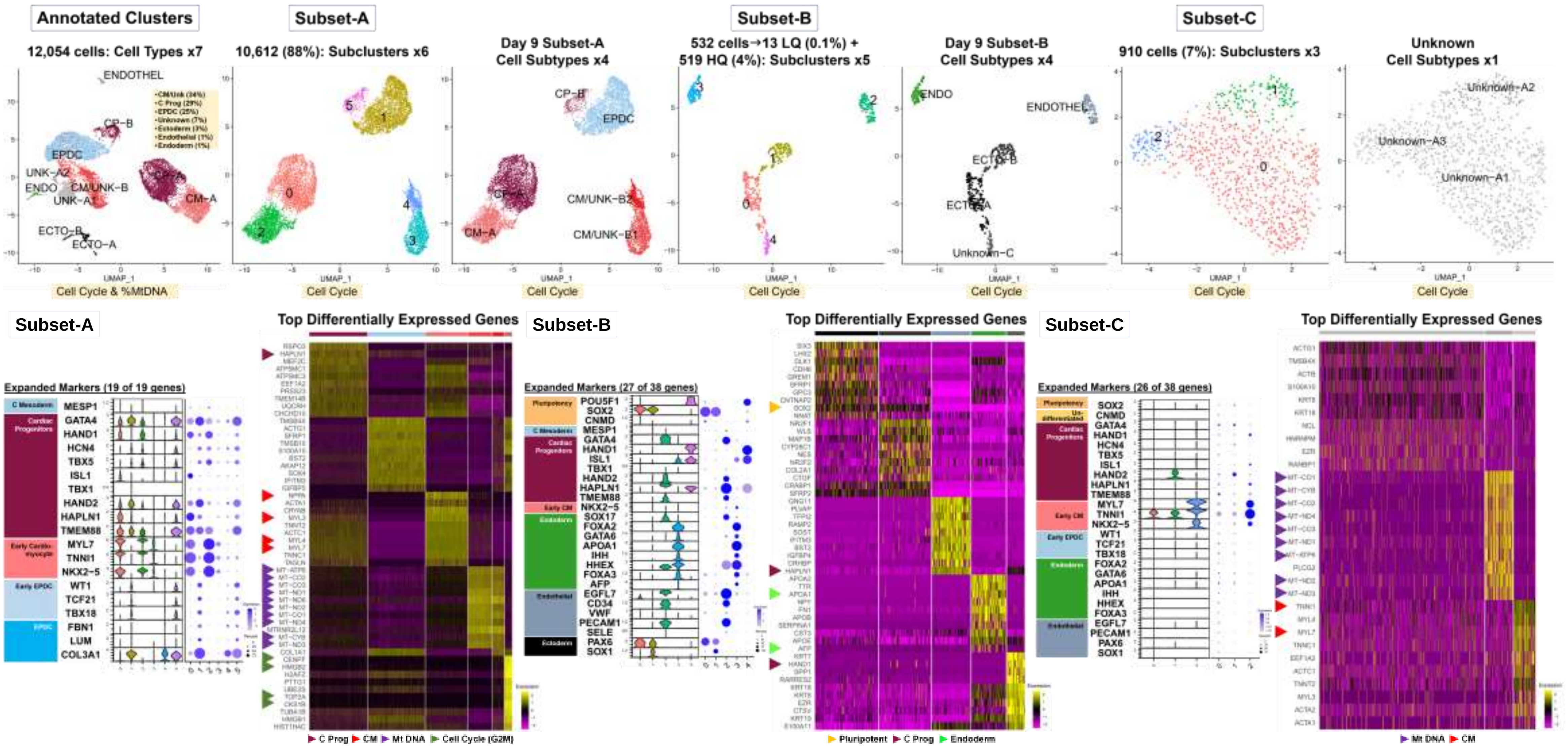

Fig. S7 Subcluster

| Control | Day 16 | Single Sample Data *Subsets* |
| --- | --- | --- |

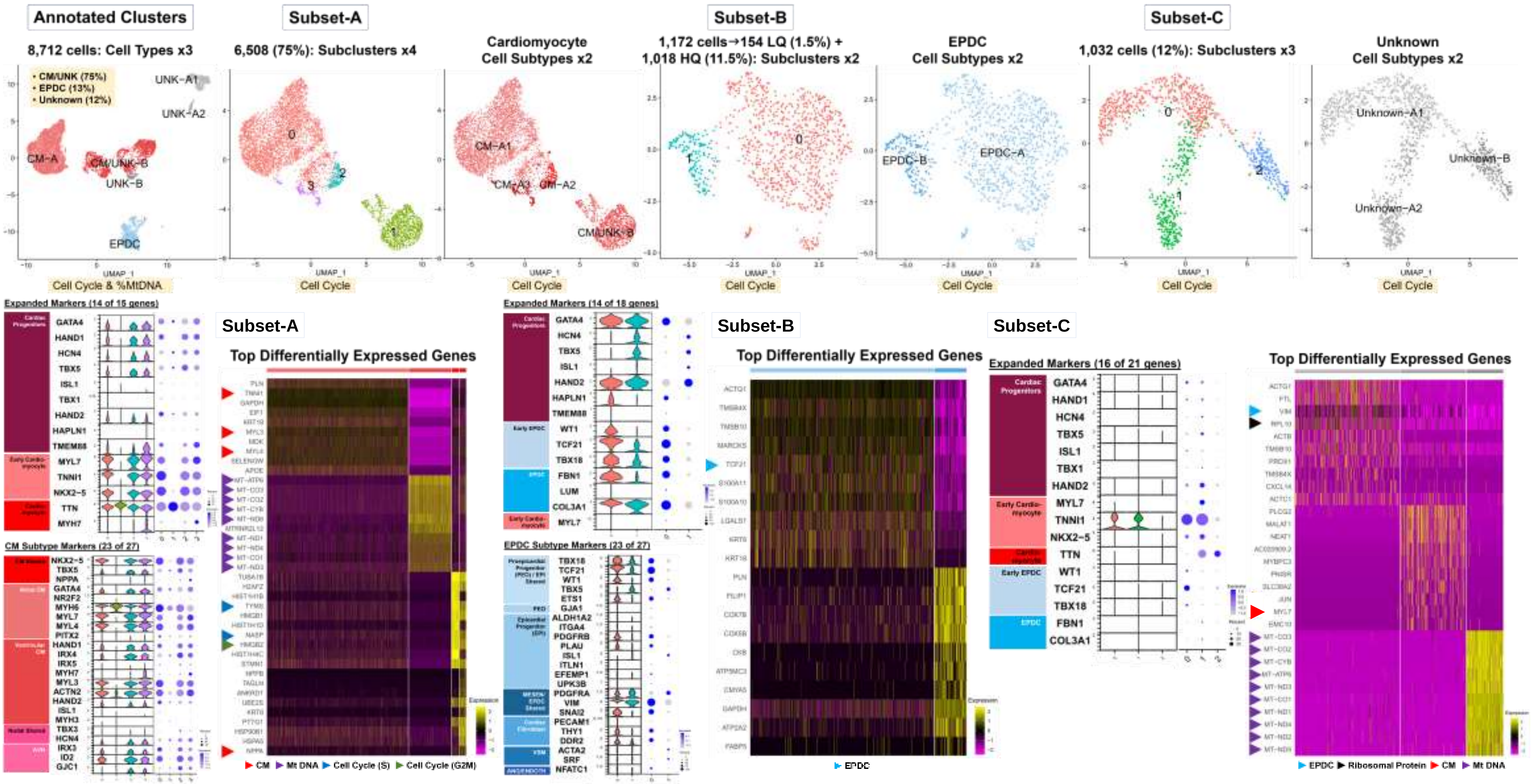

Fig. S7 Subcluster

### Control | Day 19 | Single Sample Data *Subsets*

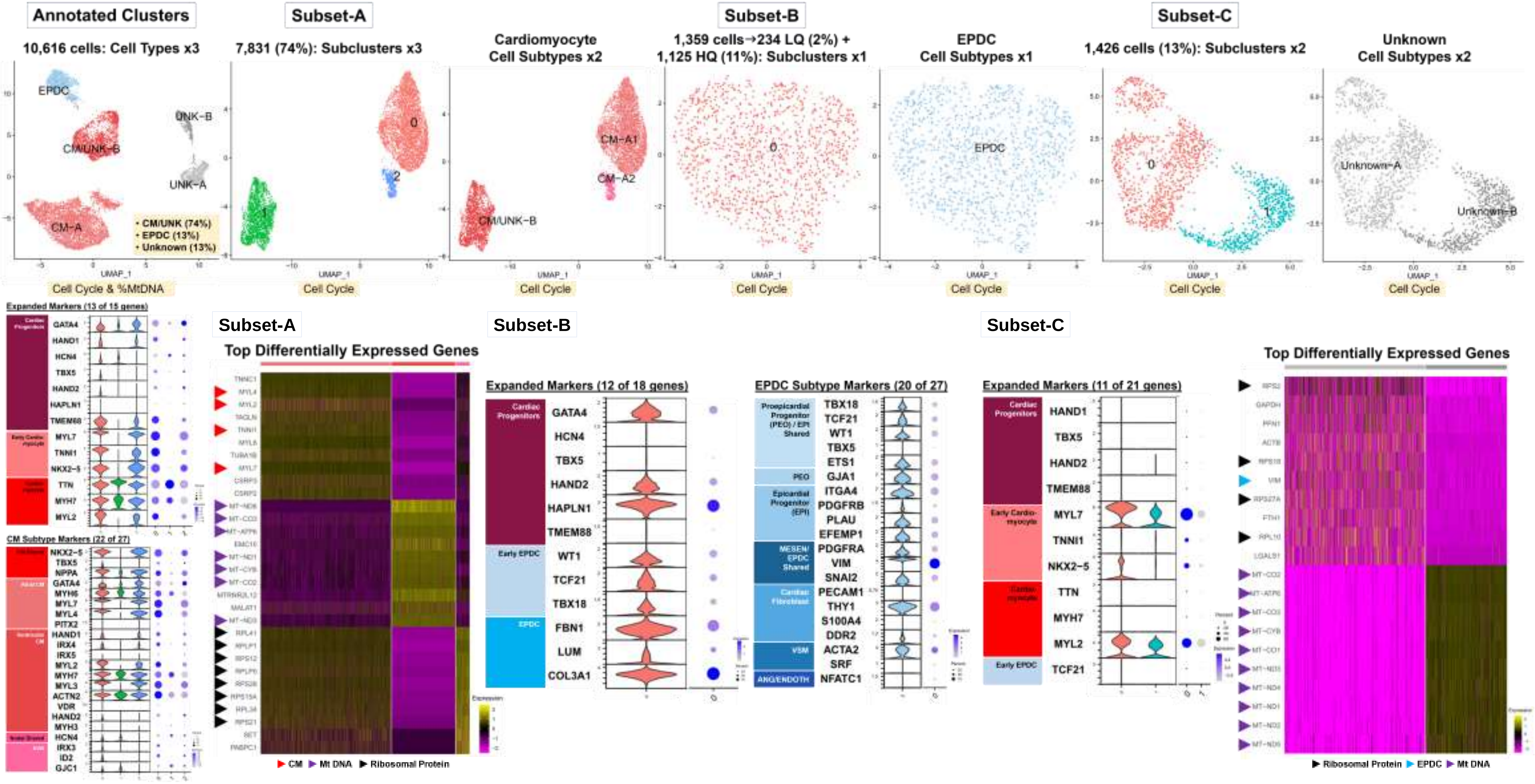

**Fig. S7 Subcluster**

| Control | Day 30 | Single Sample Data *Subsets* |
|---|---|---|

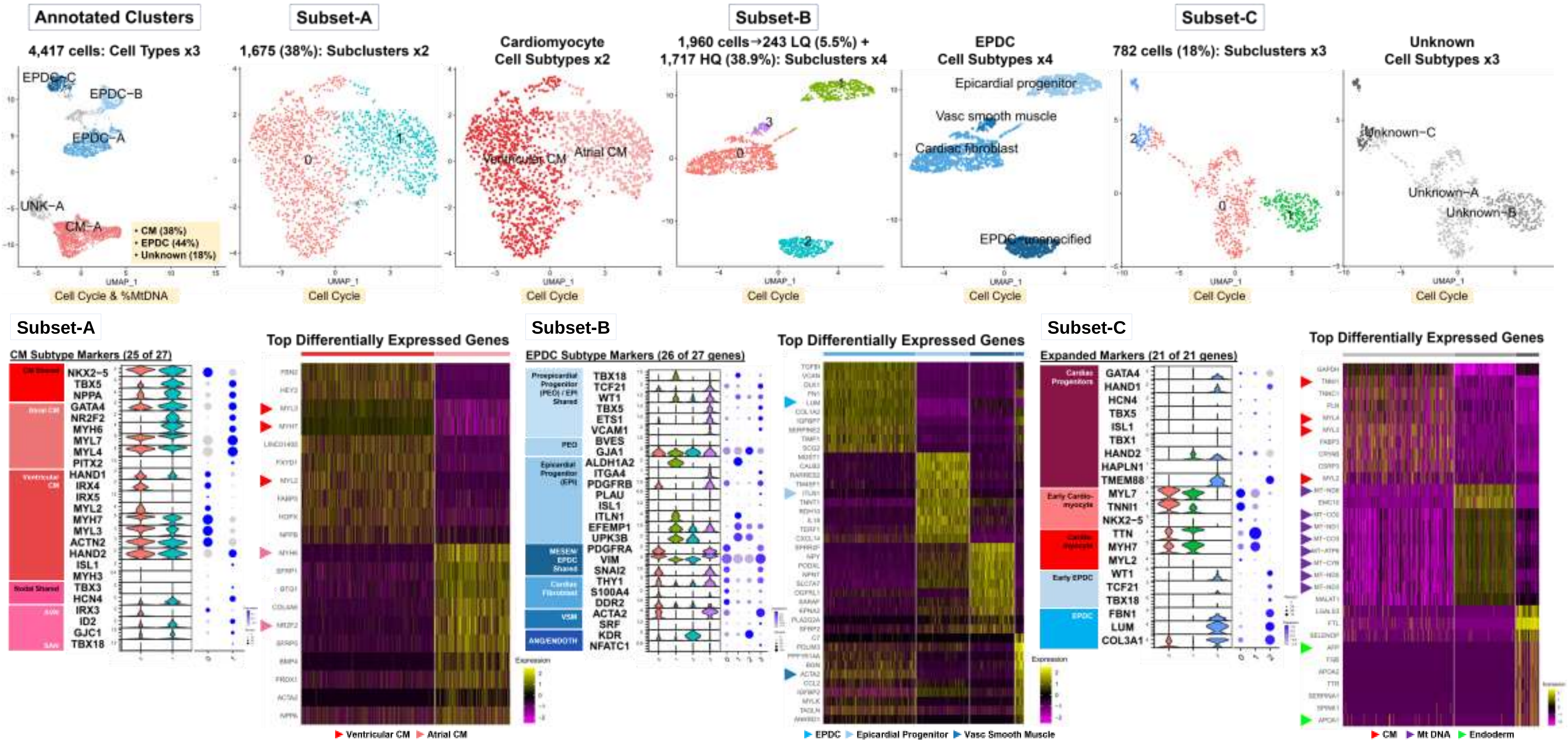

#### Fig. S7 Subcluster

### C. Subcluster Analyses: Patient Samples (n=4) to Annotated Subset Data (n=11)

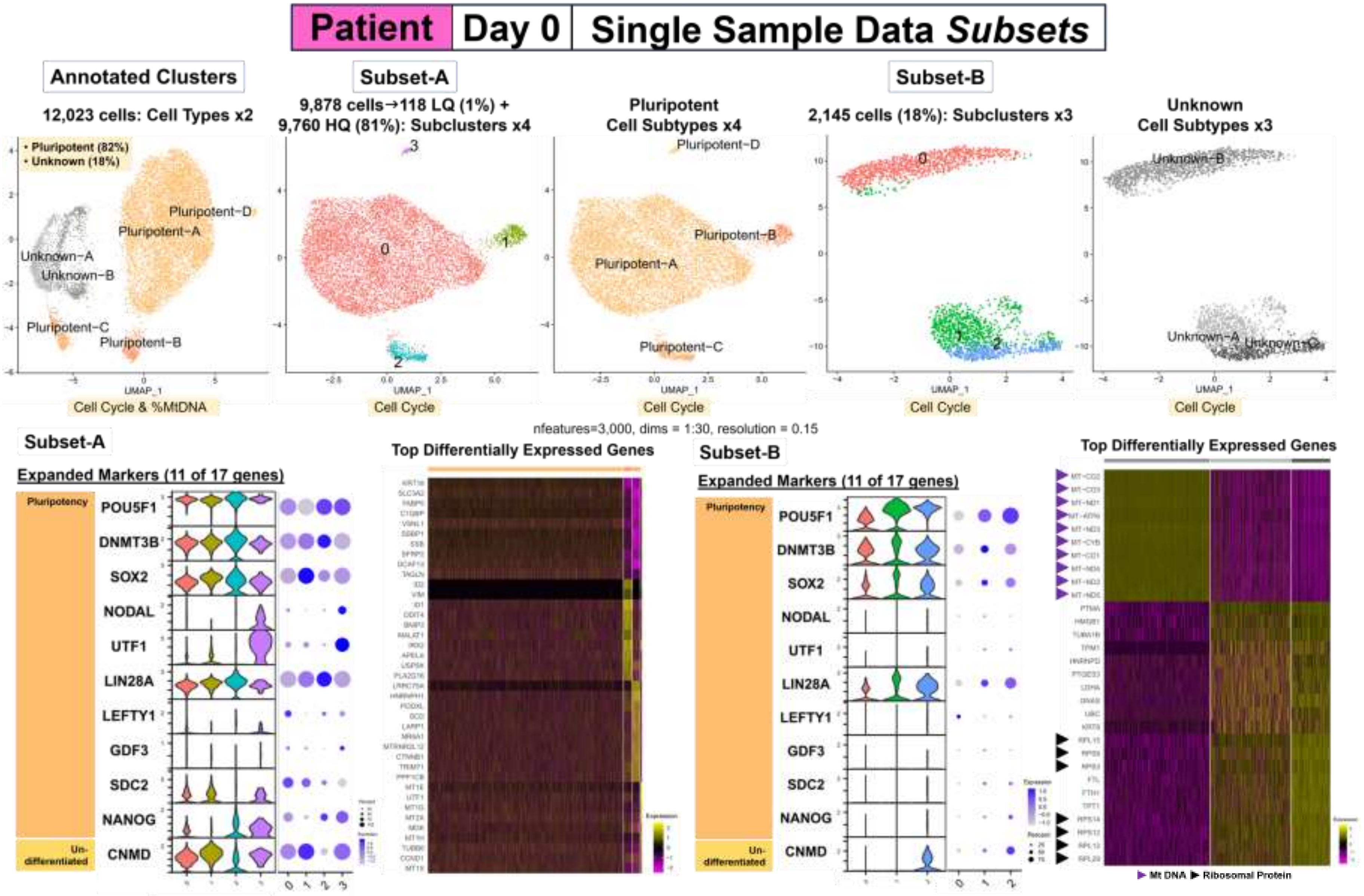

Fig. S7 Subcluster

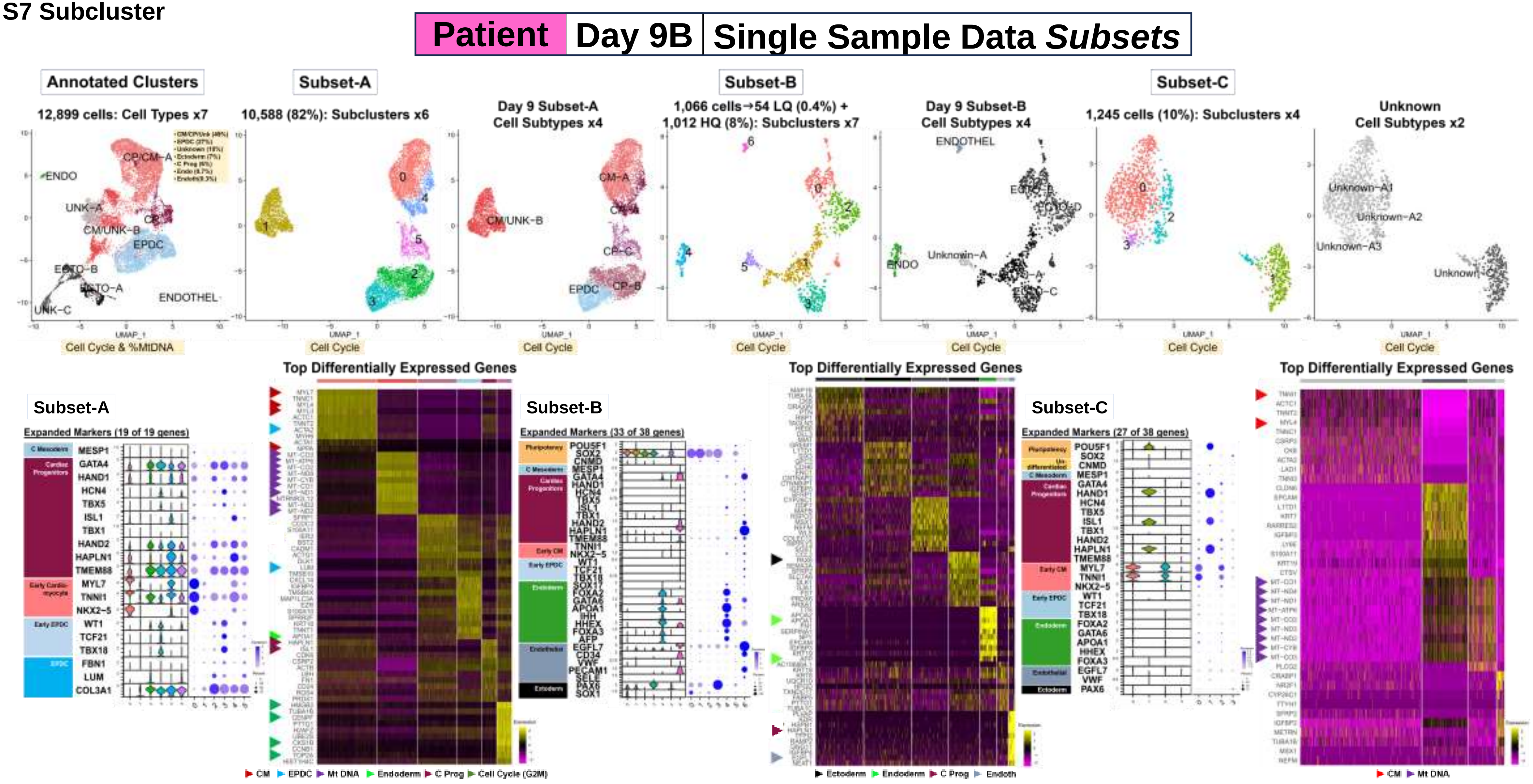

**Fig. S7 Subcluster**

| Patient | Day 16 | Single Sample Data *Subsets* |
| --- | --- | --- |

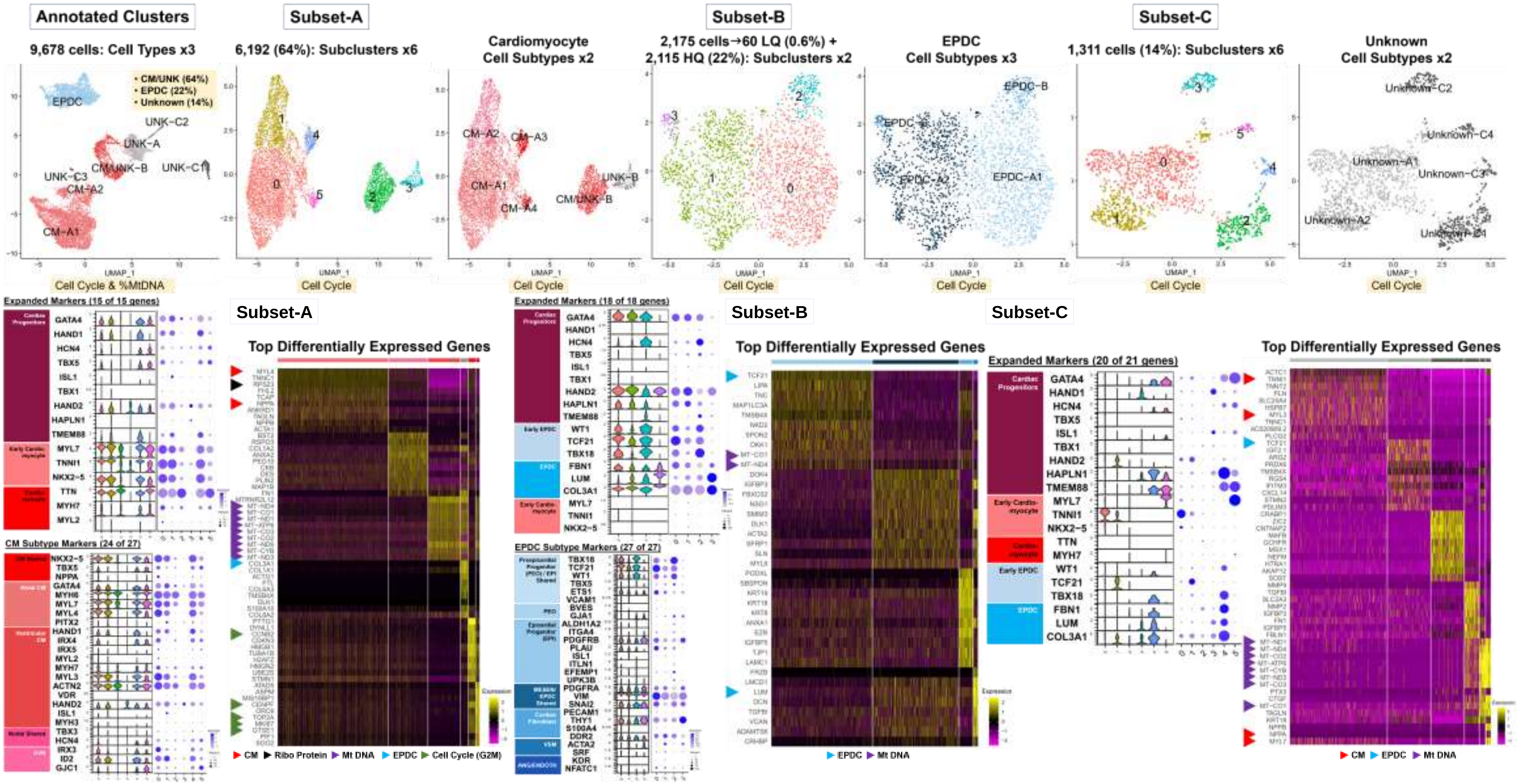

**Fig. S7 Subcluster**

**Patient | Day 19 | Single Sample Data *Subsets***

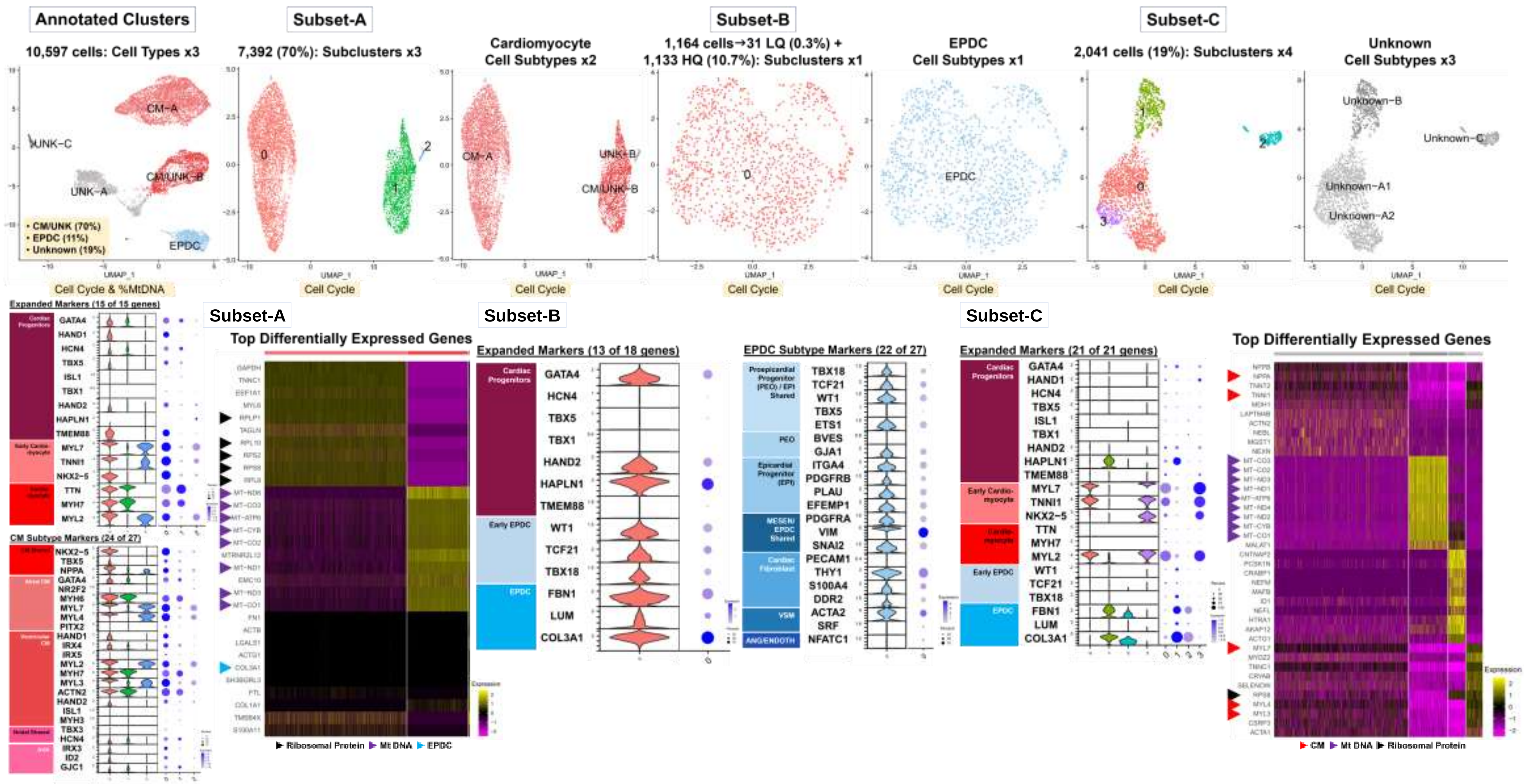

Fig. S8 Combo

### Workflow Step-III: Data Combining & Comparative Analyses

#### A. Summary: Combined Data for Paired Sample Data

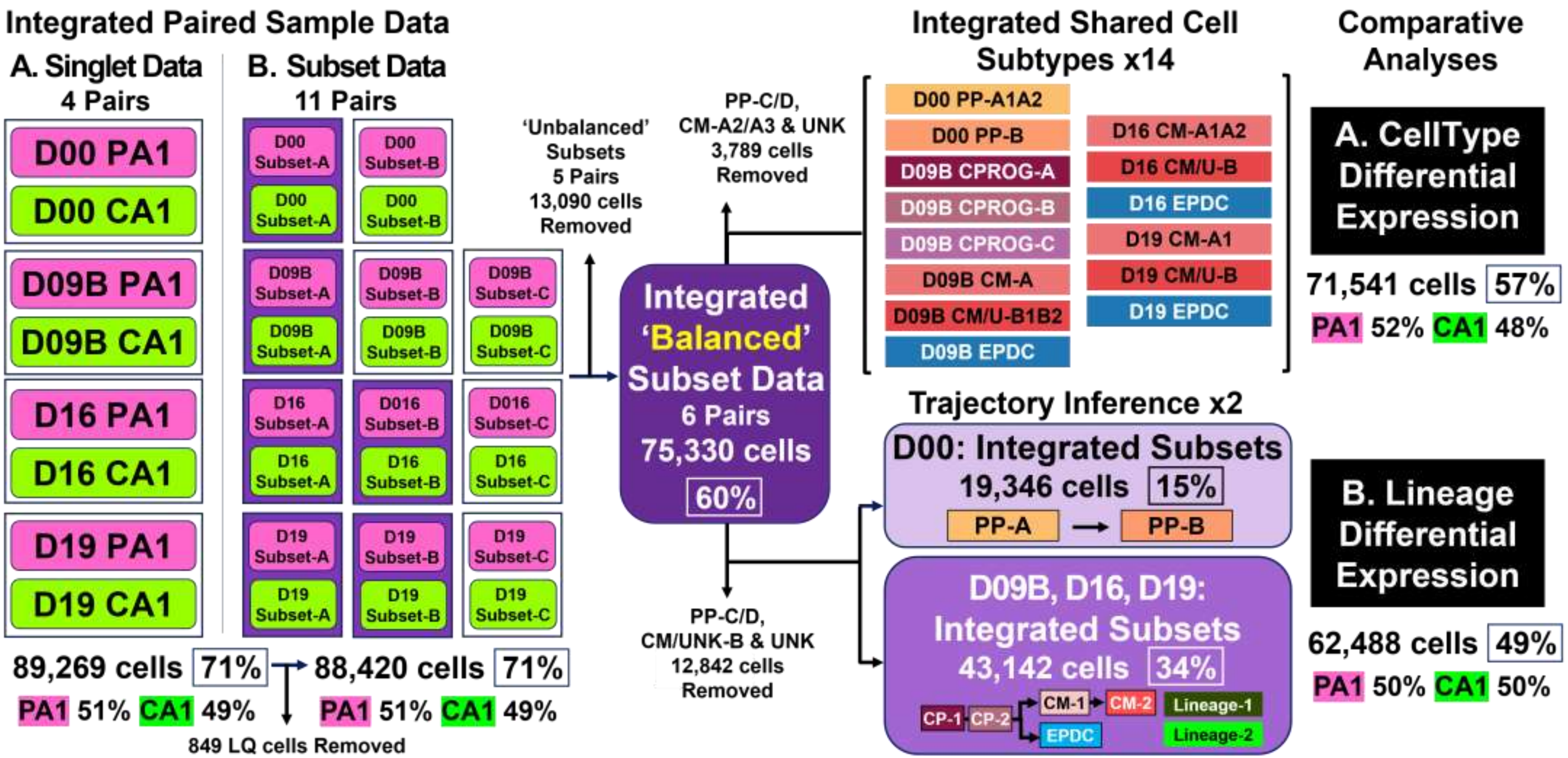

Fig. S8 Combo

#### B. Summary: Combined Data for ‘Balanced’ Paired Subsets (n=6 Prs: 75,330 Total Cells)

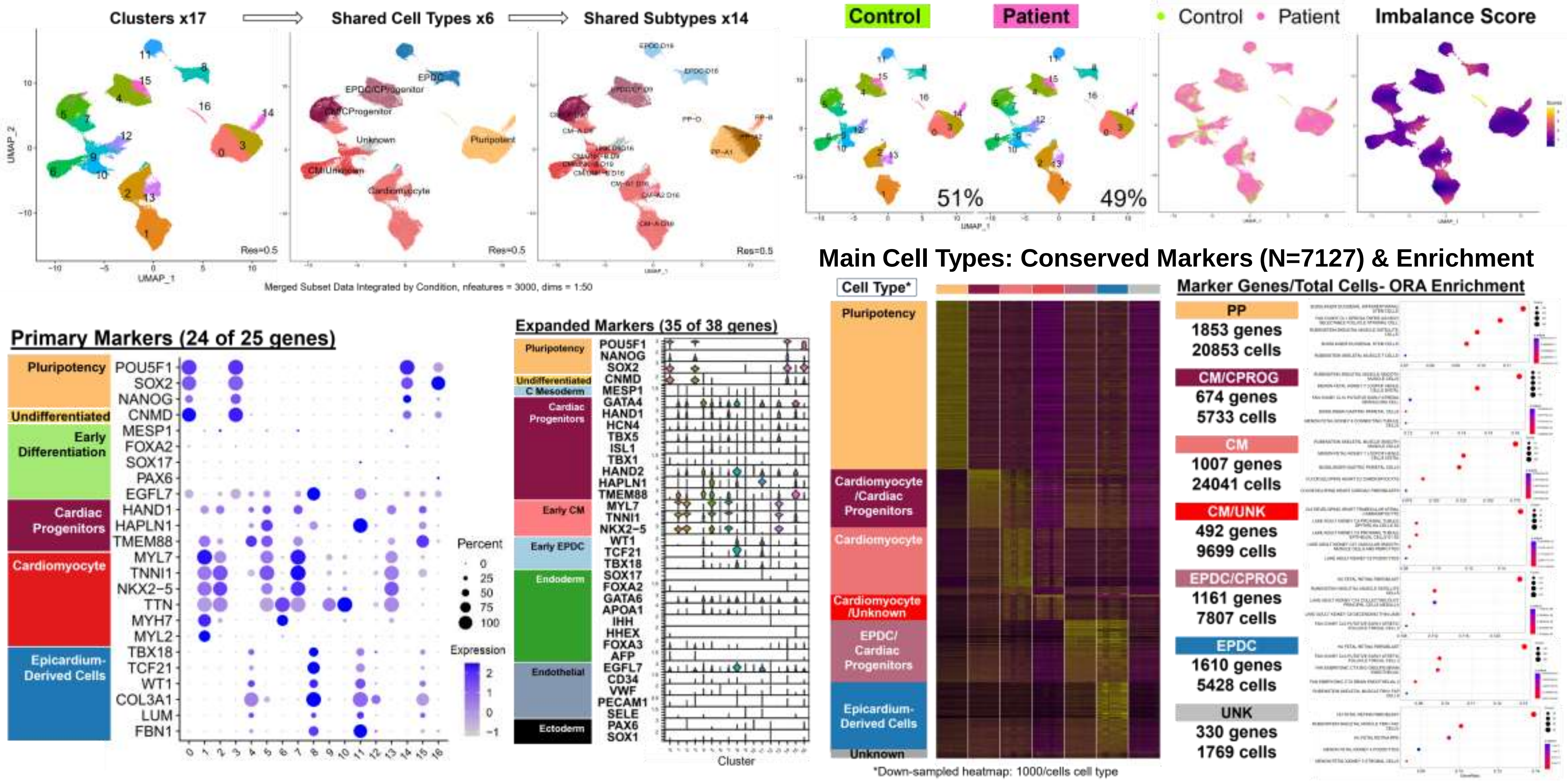

Fig. S9 Pr Cluster

### Workflow Step-III: Paired Sample Data Results

#### Individual Analyses of Combined Singlet Data for Shared Cell Types

##### A. Summary: Merged vs. Integrated Singlet Data- Clusters and Imbalance

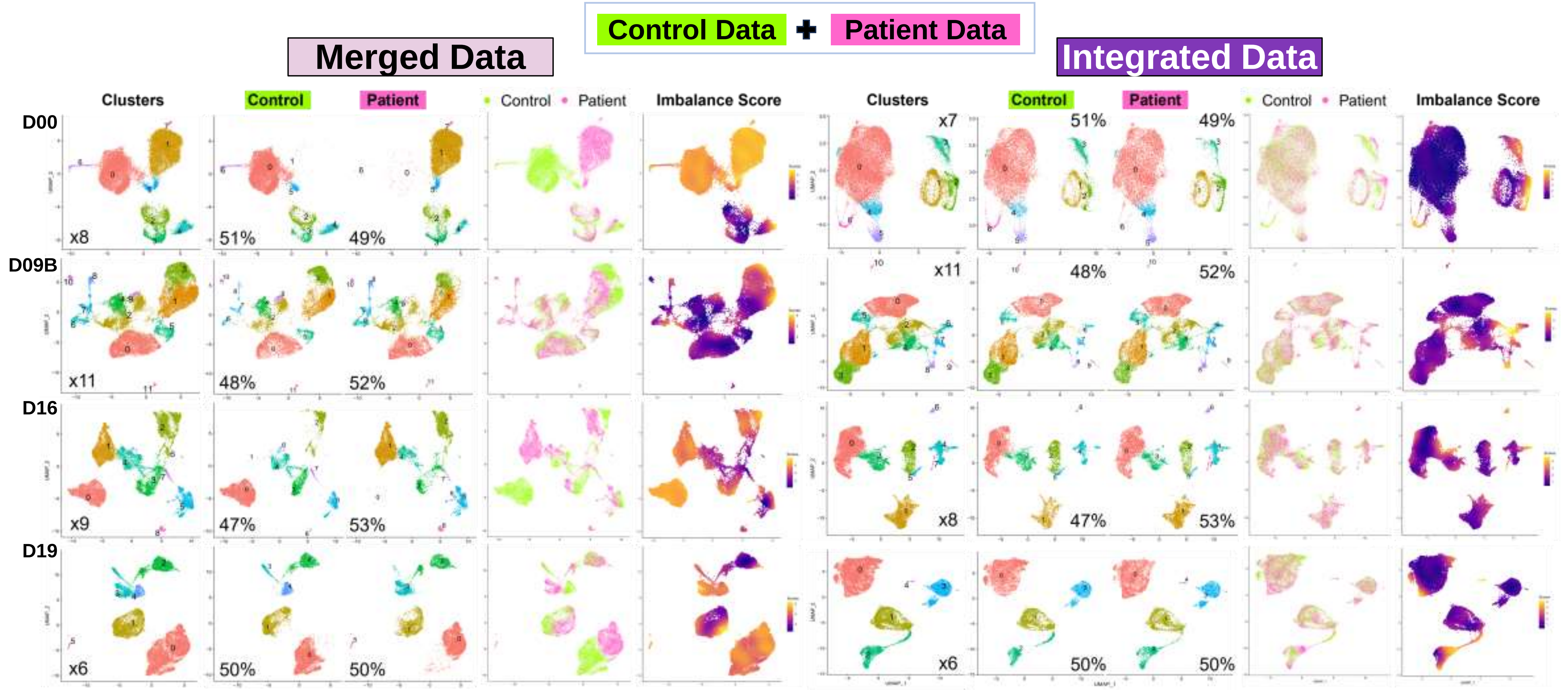

Fig. S9 Pr Cluster

#### B. Summary: Cell Annotation of Integrated Singlet Data (n=4 Prs: 89,269 Total Cells)

Control Data + Patient Data → Integrated Data

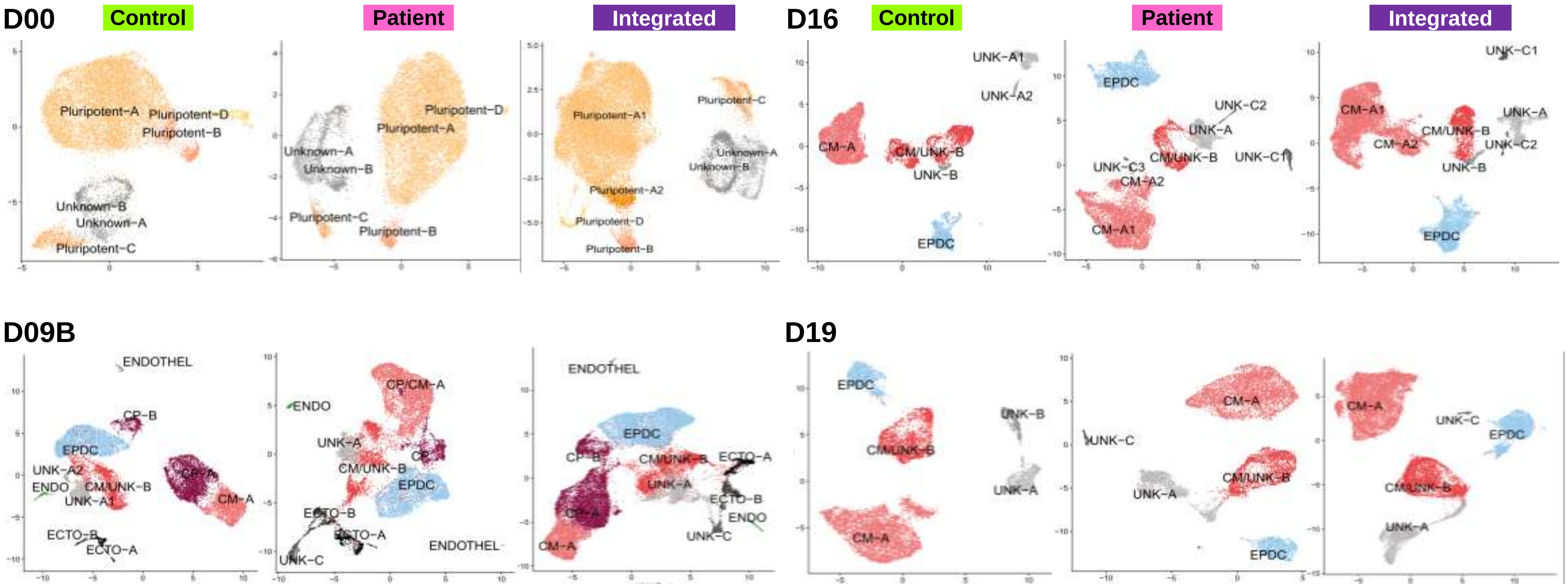

Fig. S9 Pr Cluster

#### C. Paired Sample Data Analyses: Integrated Singlet Data- Patient vs. Control (n= 4 Prs)

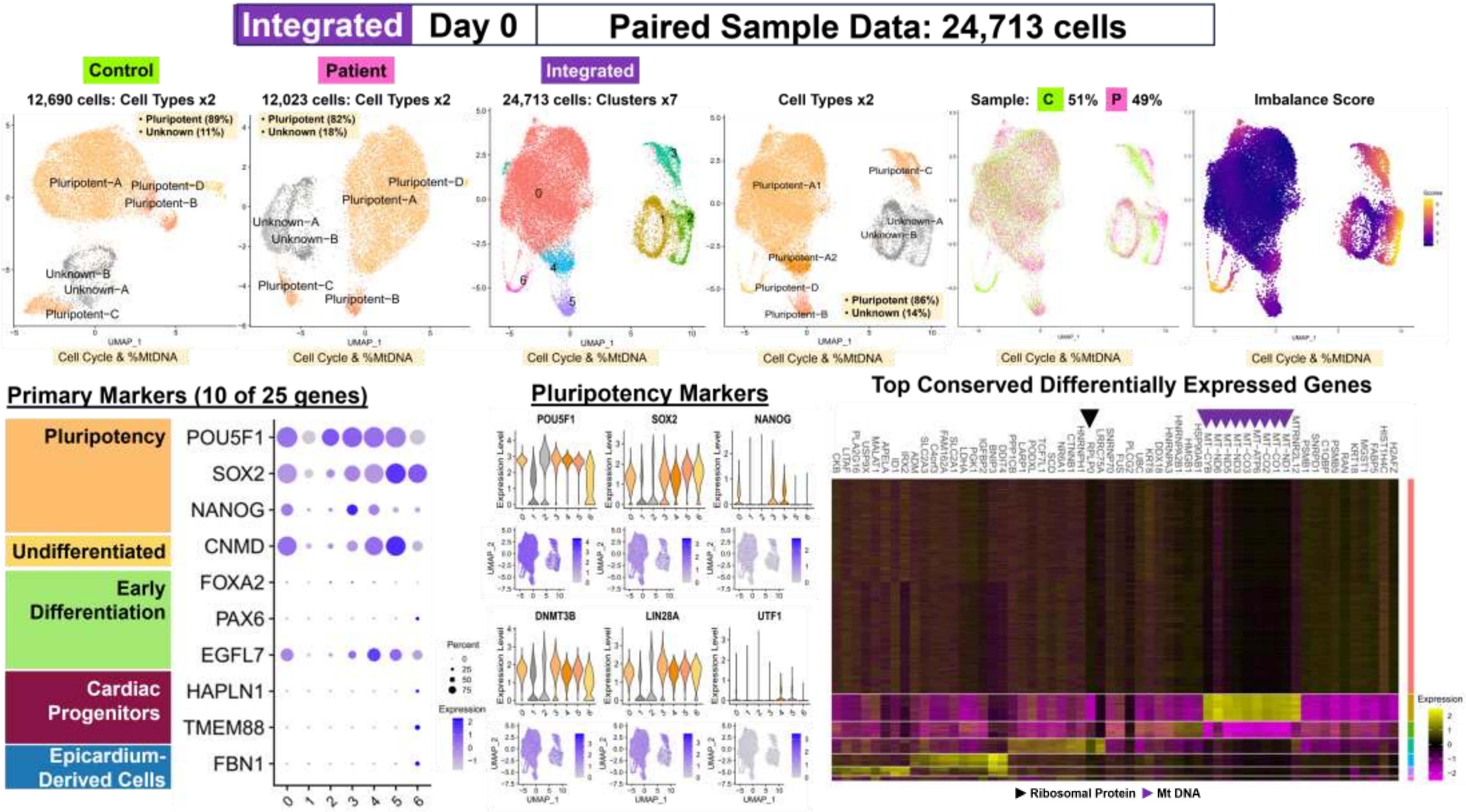

Fig. S9 Pr Cluster

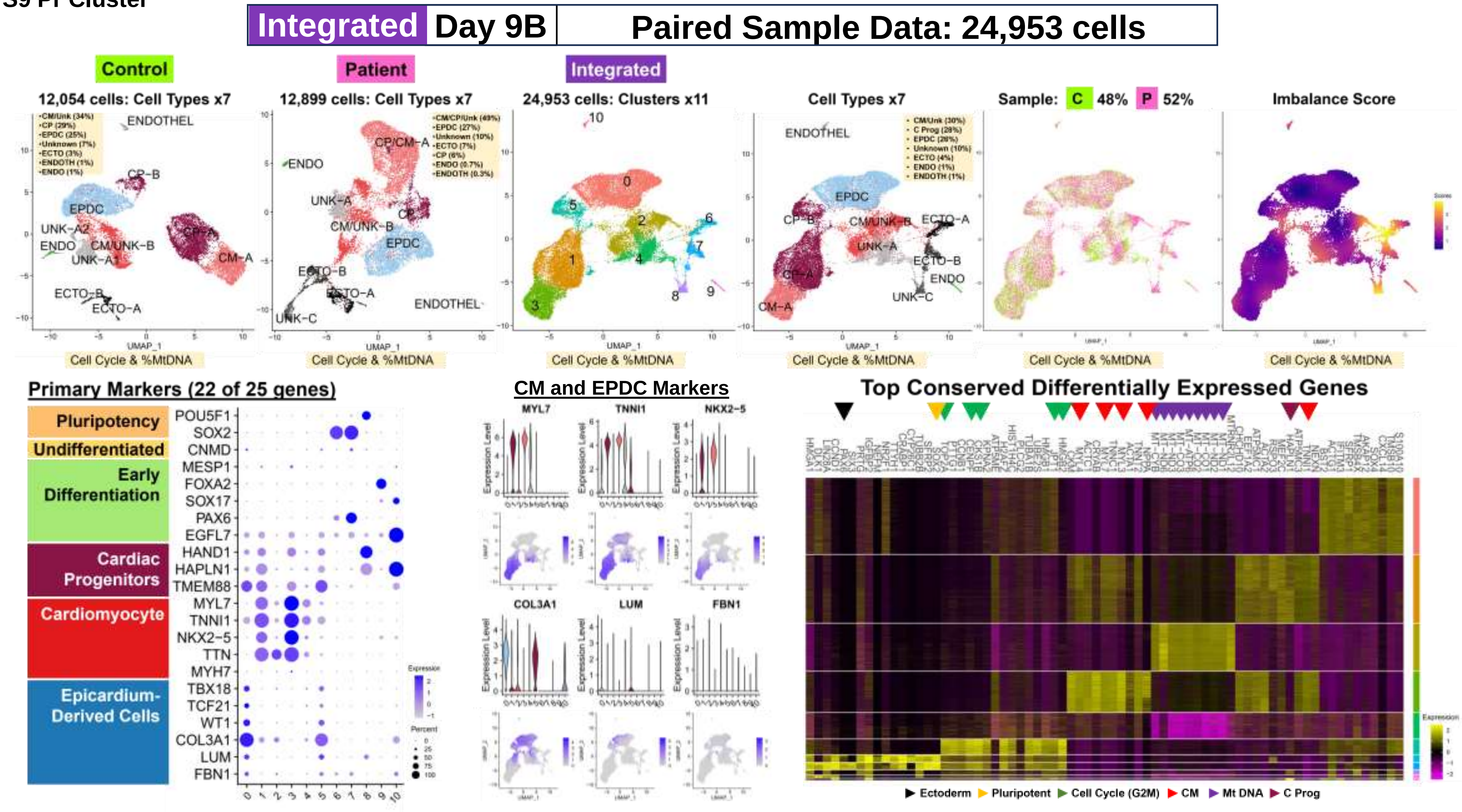

**Fig. S9 Pr Cluster**

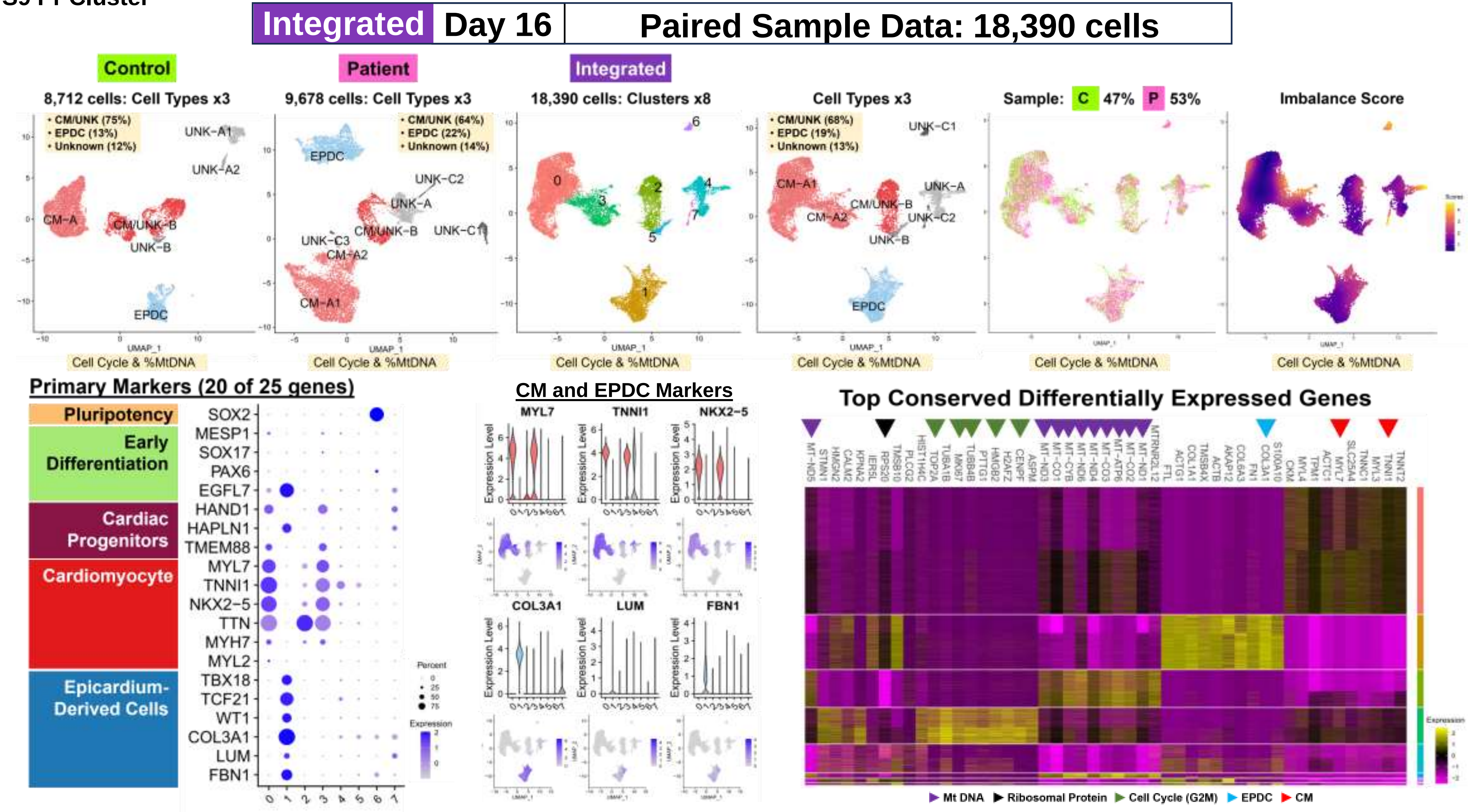

**Fig. S9 Pr Cluster**

**Integrated | Day 19 | Paired Sample Data: 21,213 cells**

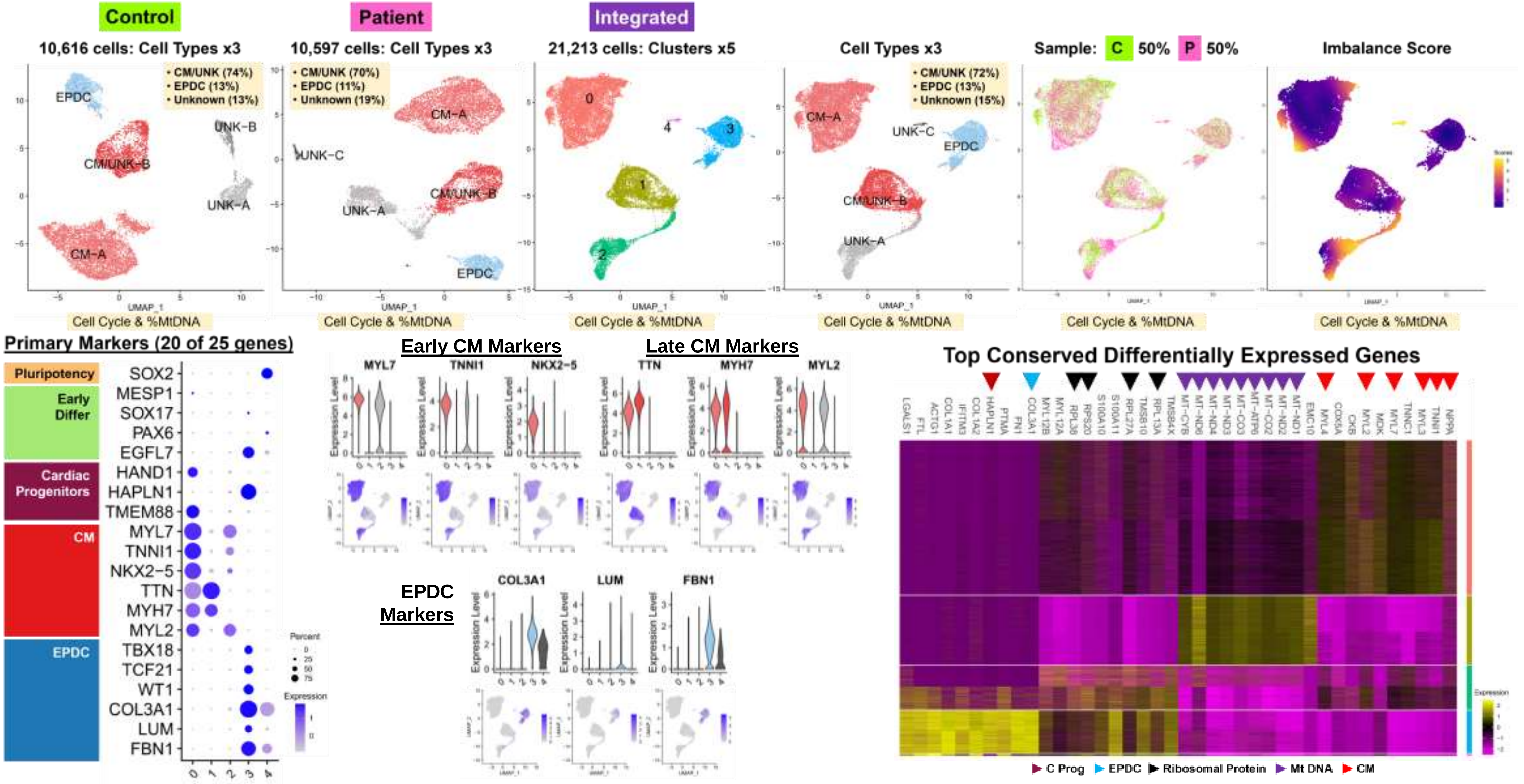

Fig. S10 Pr Subcluster

### Workflow Step-III: Paired Sample Data Results

#### Individual Subcluster Analyses of Combined Subset Data for Possible Shared Subtypes

##### A. Summary: Merged vs. Integrated Subset Data- Clusters and Imbalance

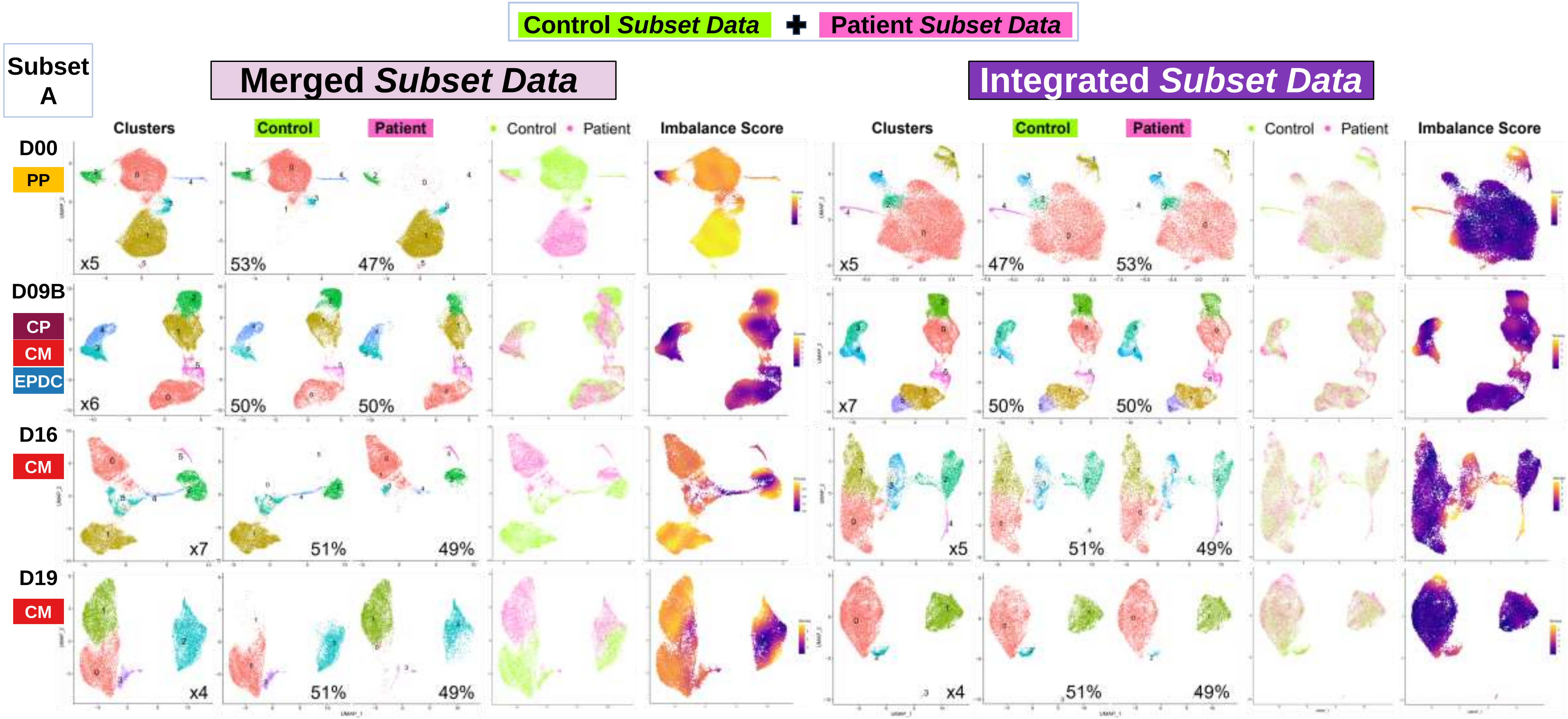

#### Fig. S10 Pr Subcluster

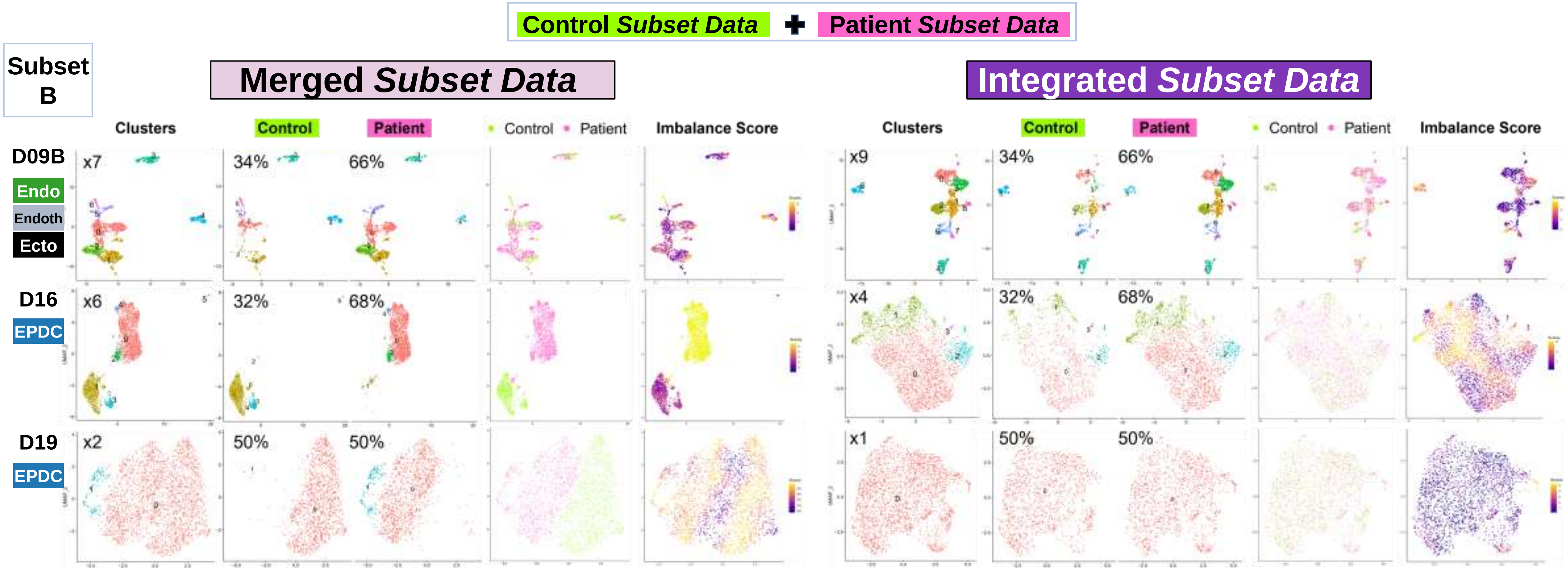

#### Fig. S10 Pr Subcluster

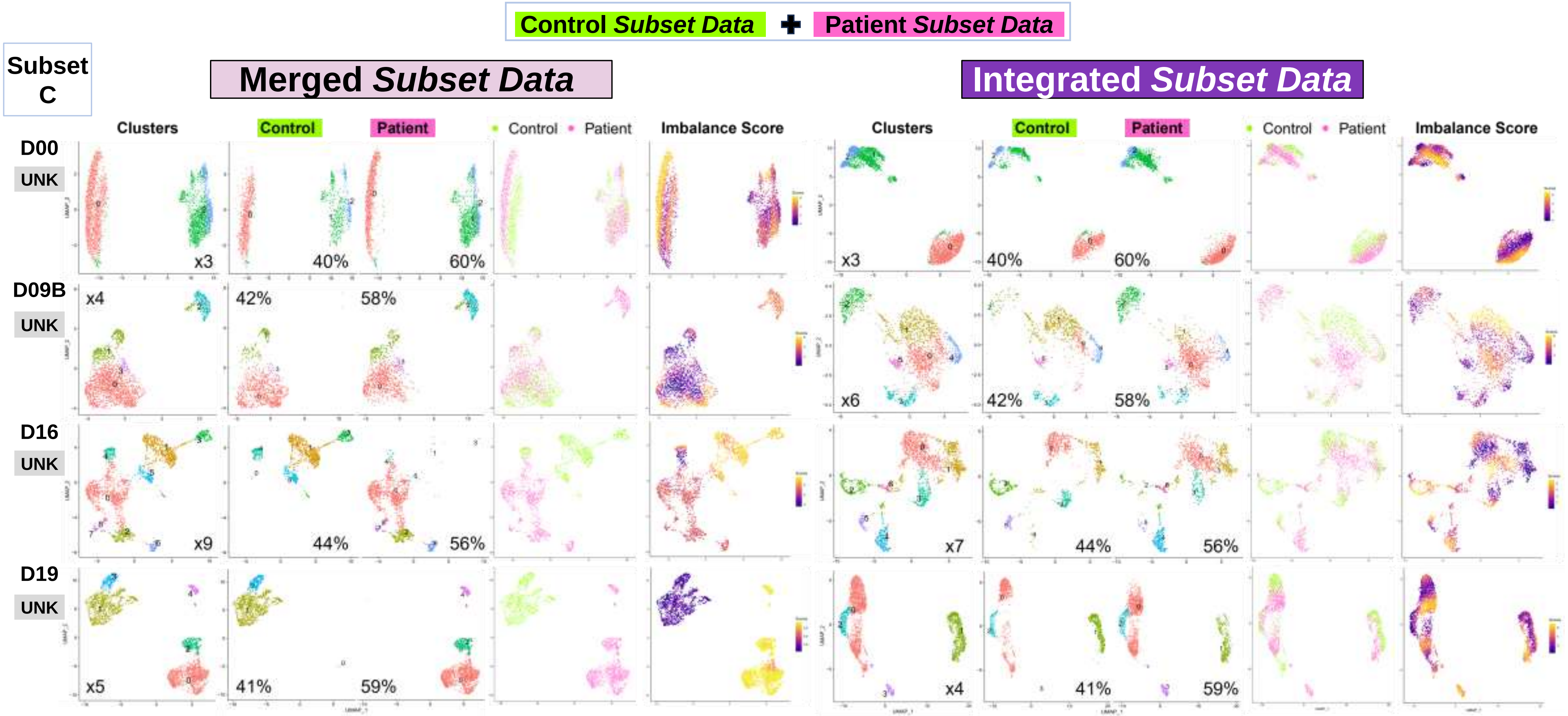

**Fig. S10 Pr Subcluster**

#### B. Summary: Cell Annotation of Integrated Subset Data (n=11 Prs: 88,420 Total Cells)

Control *Subset Data* + Patient *Subset Data* → Integrated *Subset Data*

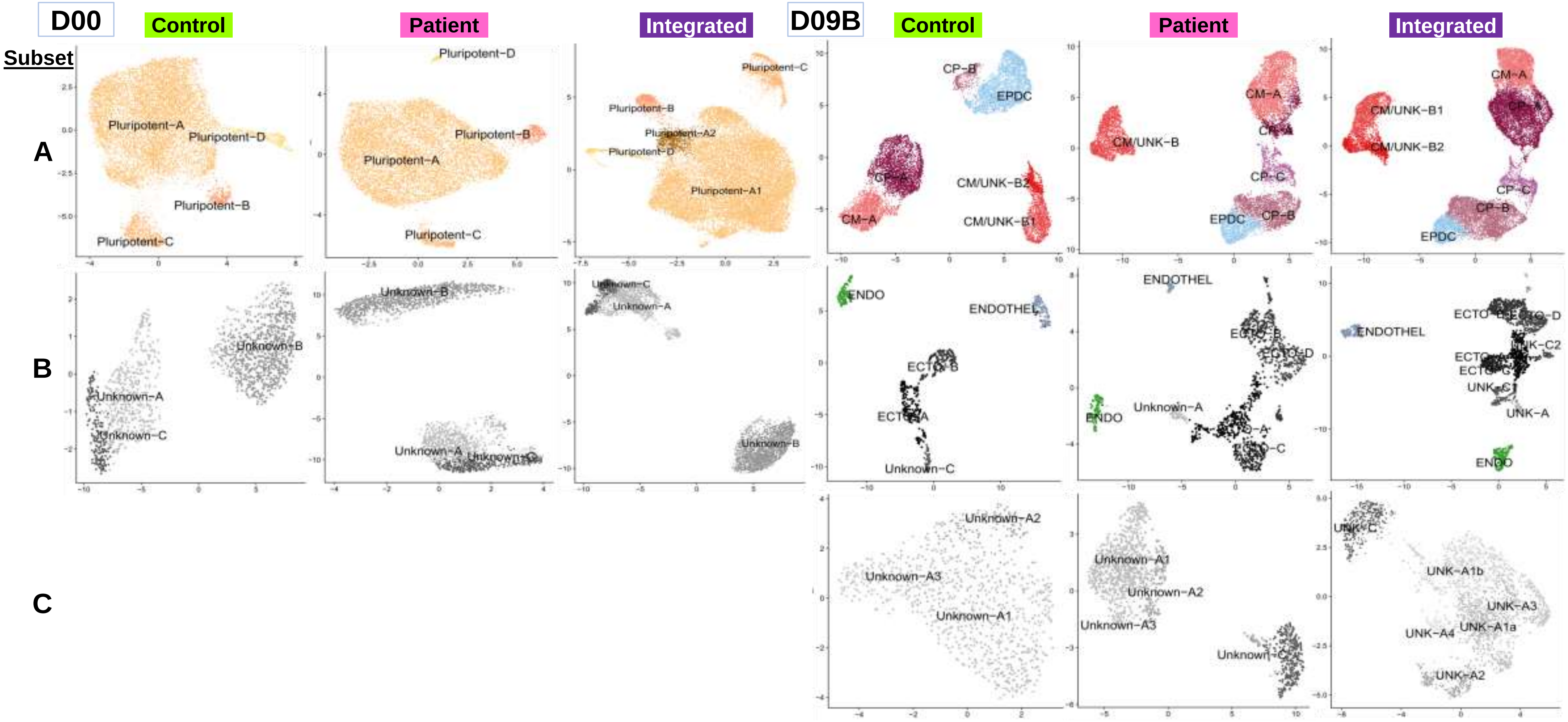

Fig. S10 Pr Subcluster

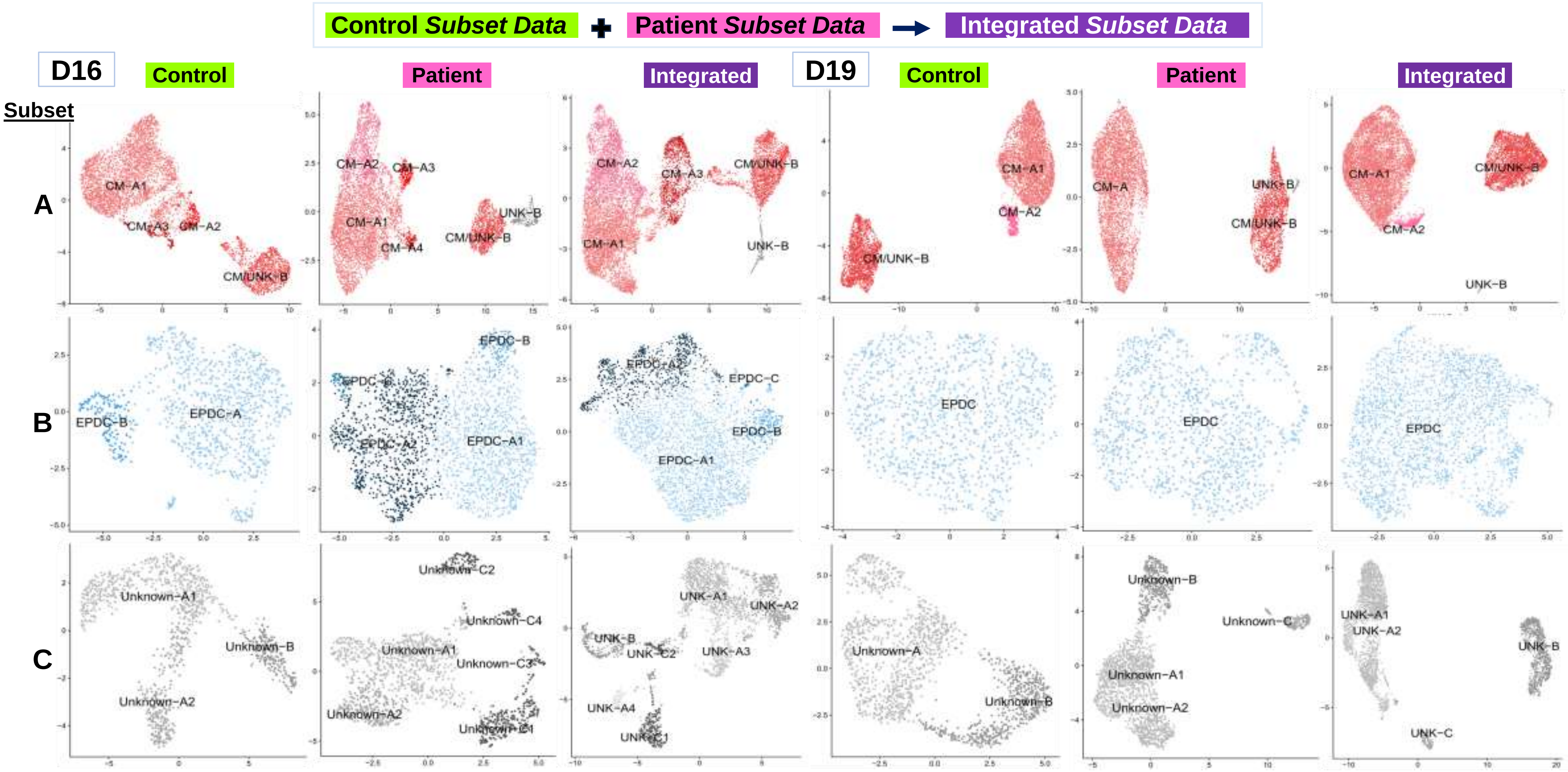

Fig. S10 Pr Subcluster

#### C. Paired Sample Data Analyses: Integrated Subset Data- Patient vs. Control (n= 11 Prs)

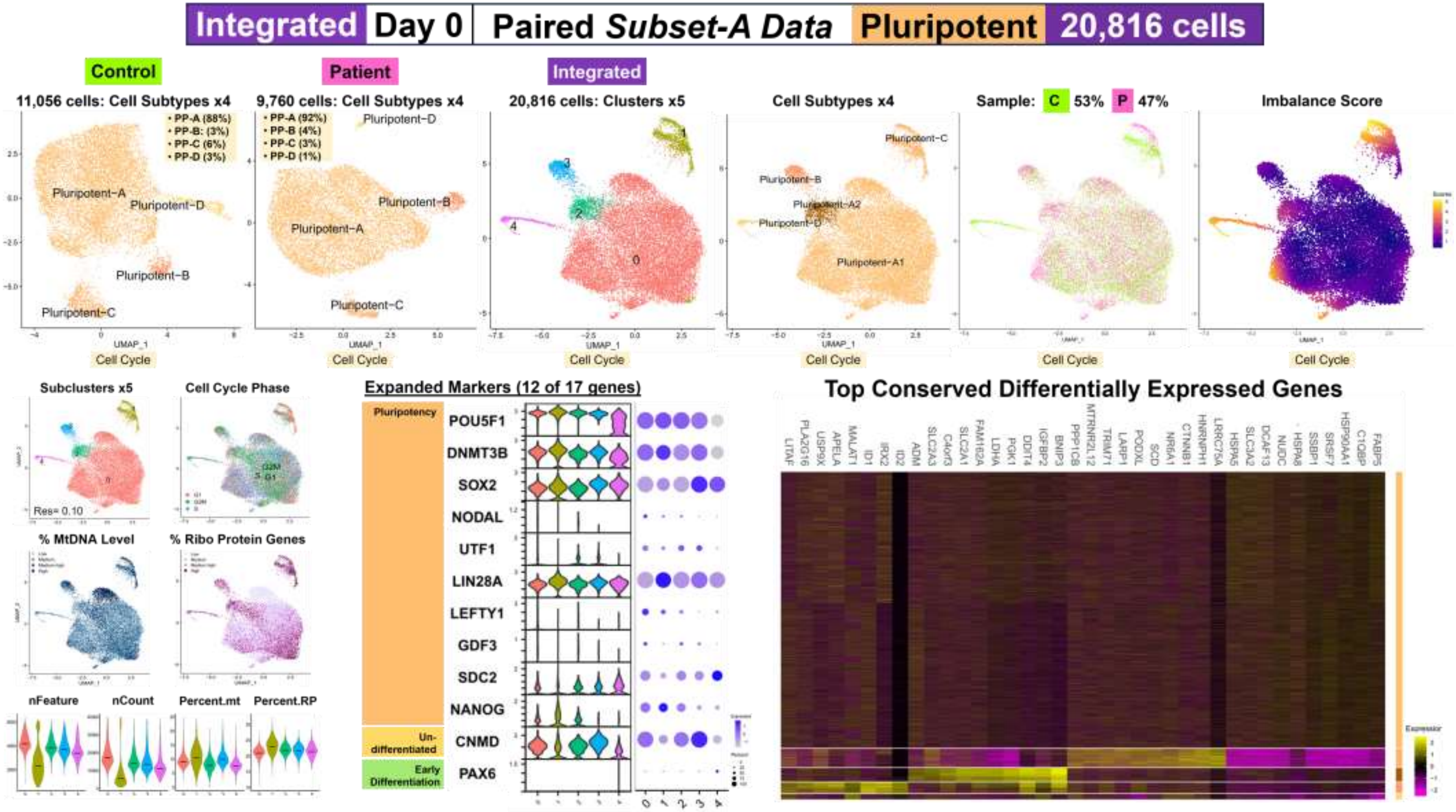

**Fig. S10 Pr Subcluster**

**Integrated | Day 0 | Paired *Subset-B Data* | Unknown | 3,594 cells**

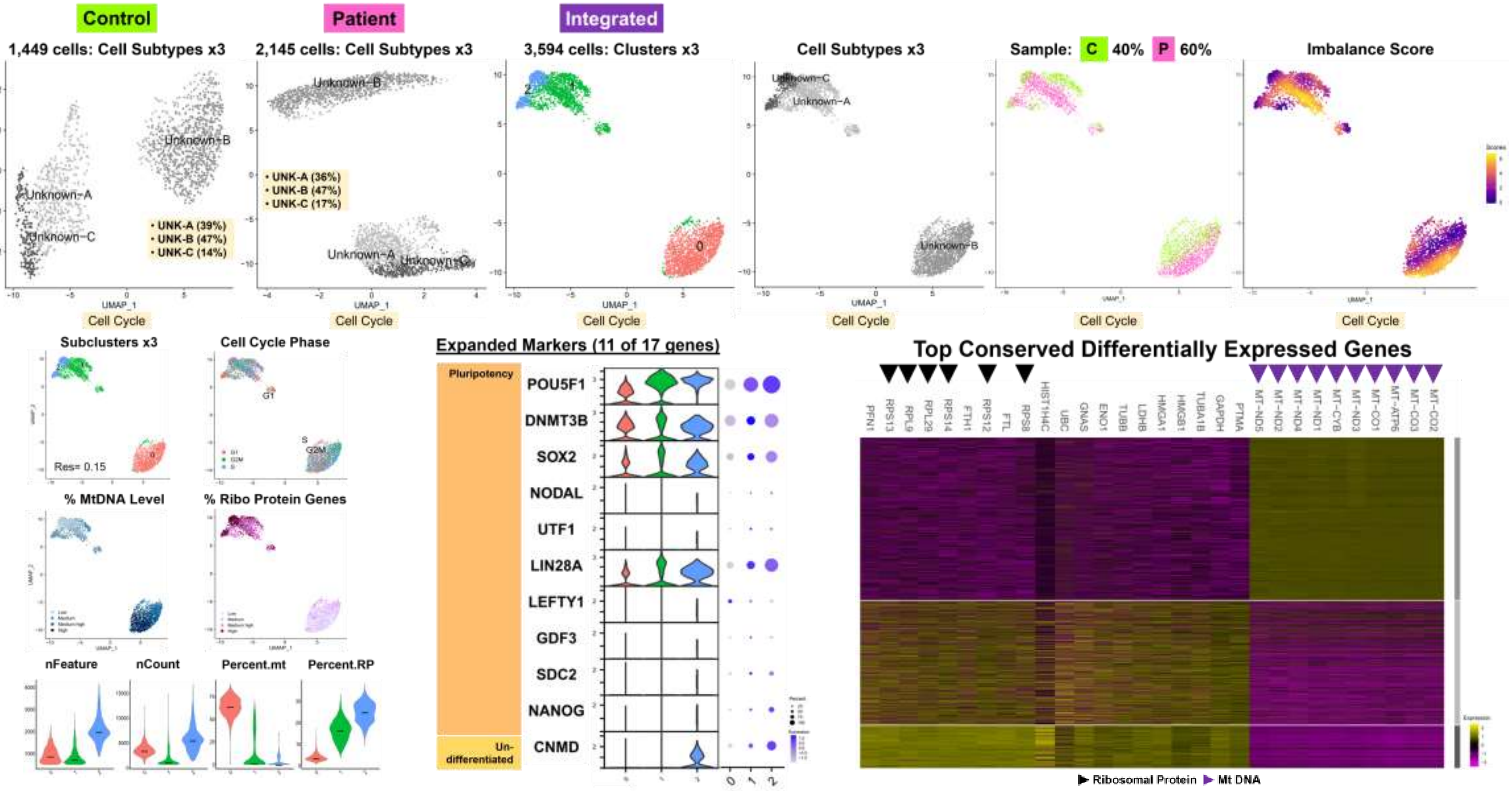

Fig. S10 Pr Subcluster

Integrated | Day 9B | Paired *Subset-A Data* CP-CM-EPDC 21,200 cells

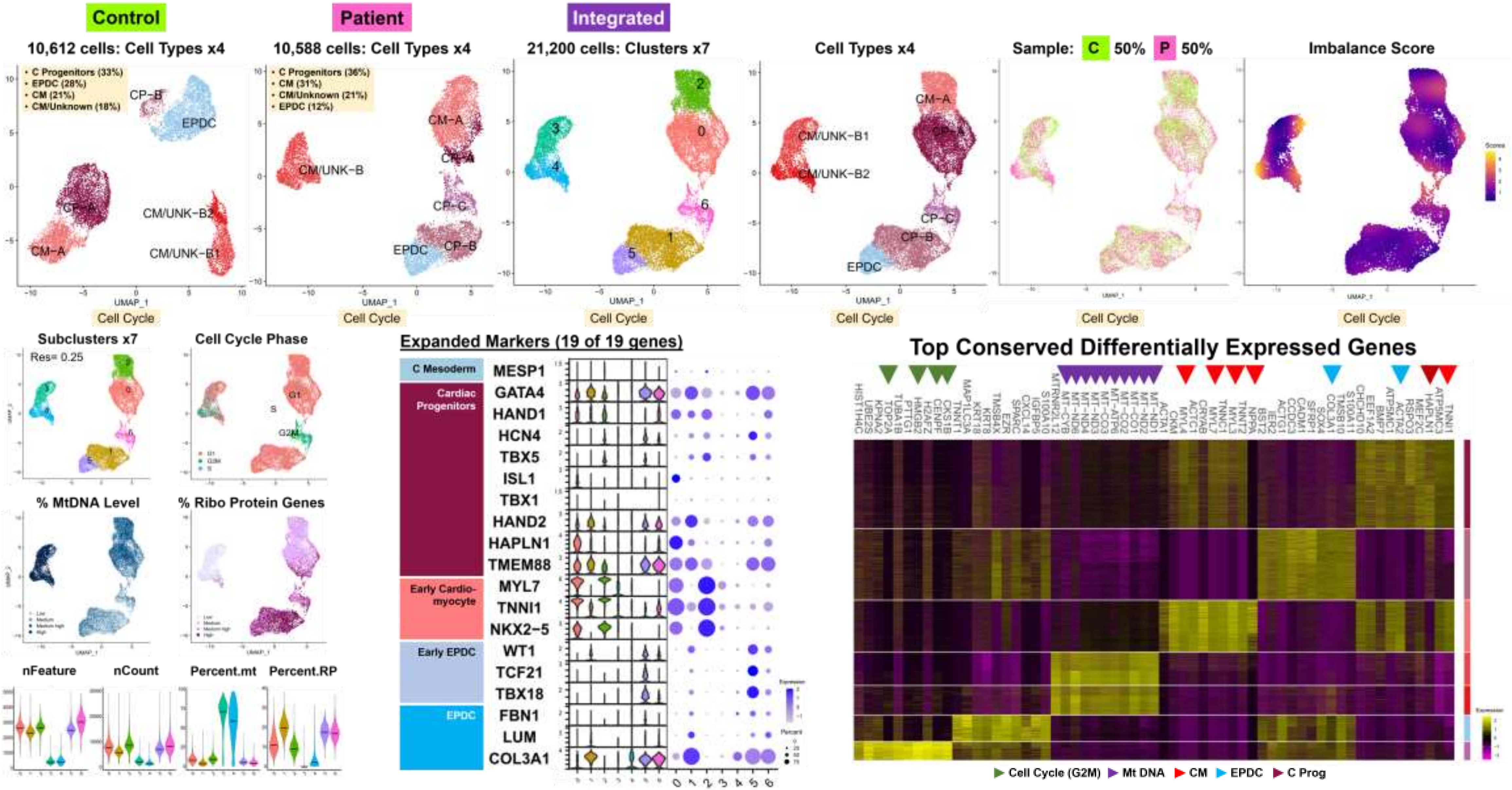

Fig. S10 Pr Subcluster

**Integrated | Day 9B | Paired *Subset-B Data* | Endo-Ecto-Endoth | 1,531 cells**

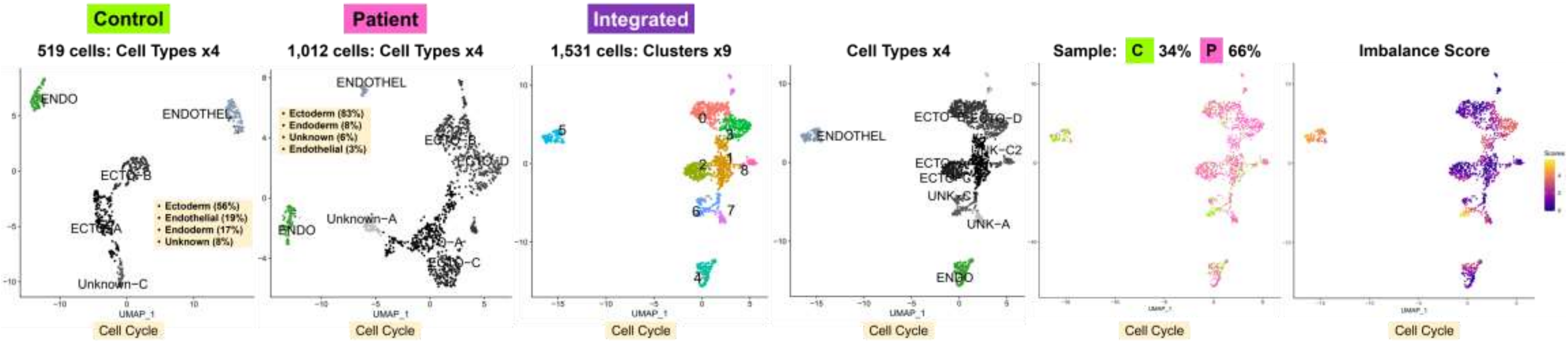


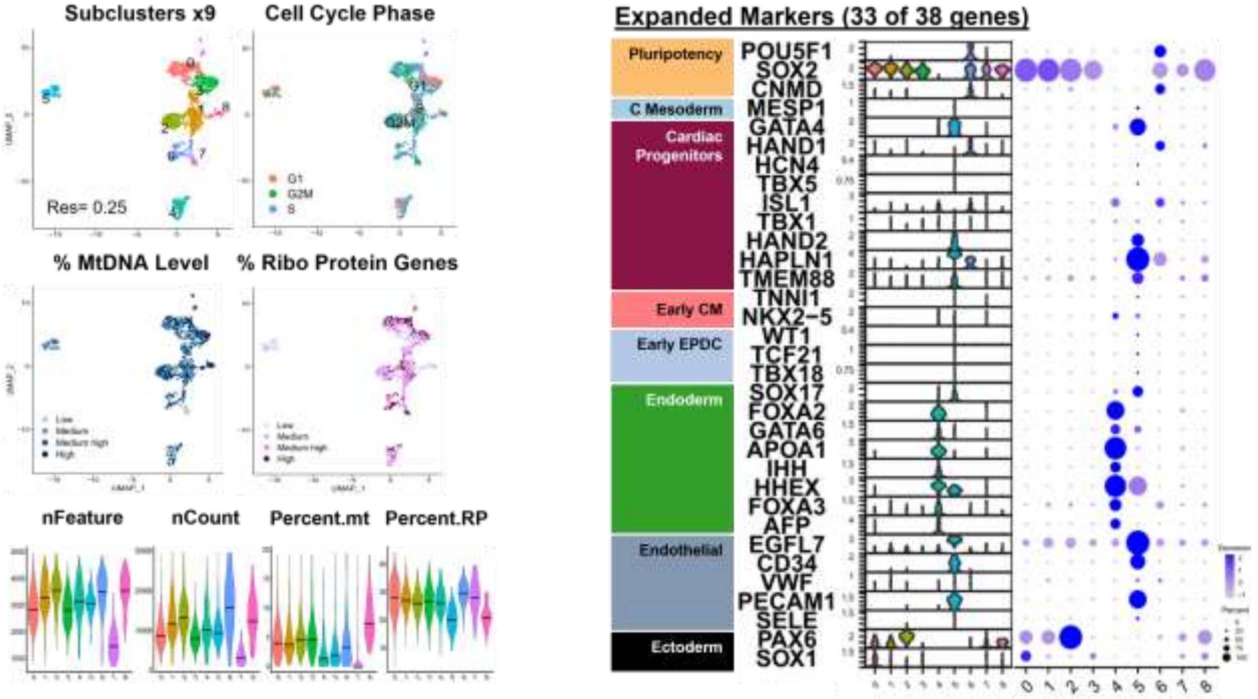


**Hmap Conserved DEG:**
**Not done since <100 cells each for all Control and Patient clusters**

**Fig. S10 Pr Subcluster**

| Integrated | Day 9B | Paired *Subset-C Data* | Unknown | 2,155 cells |
|---|---|---|---|---|

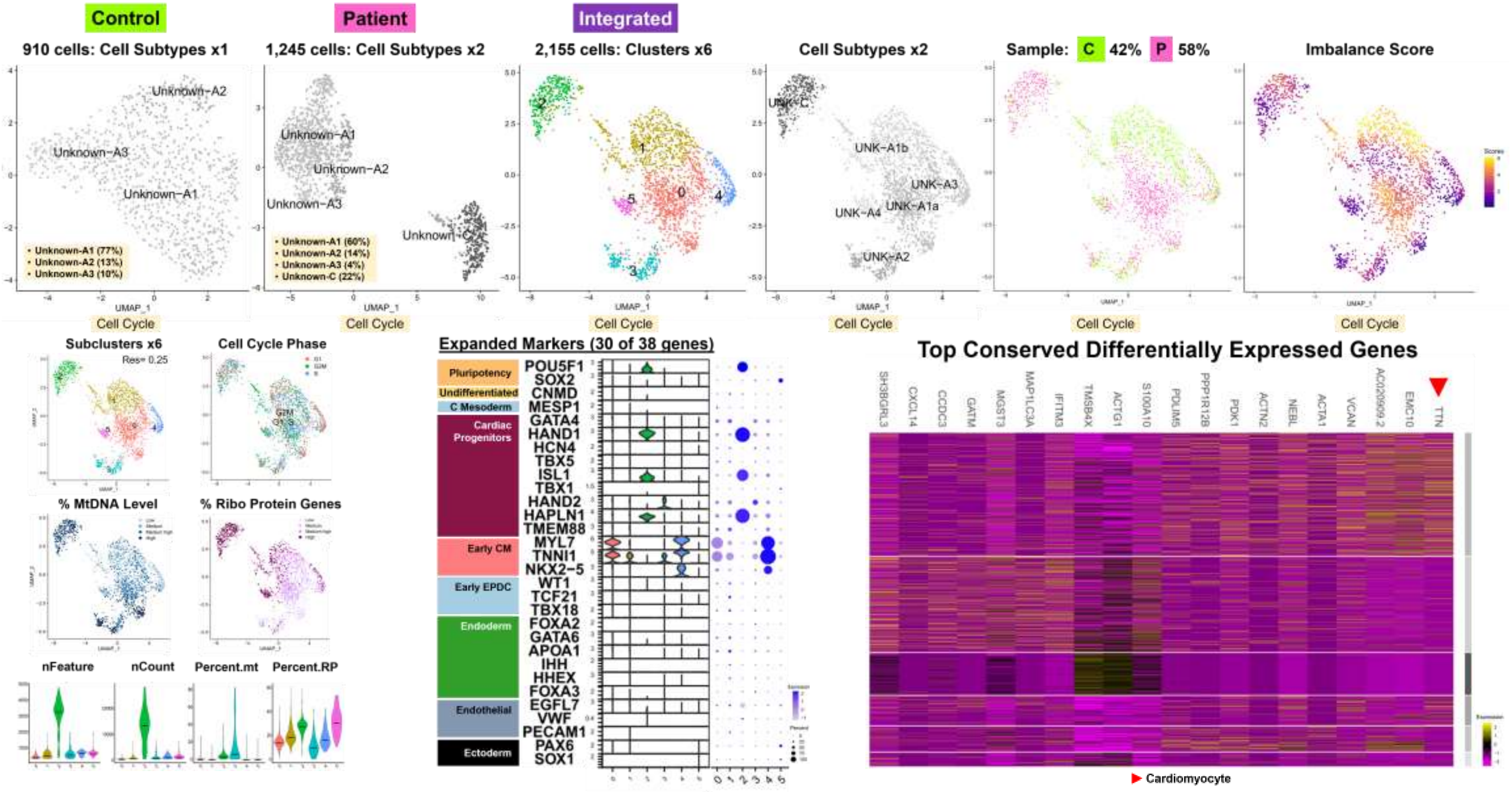

Fig. S10 Pr Subcluster

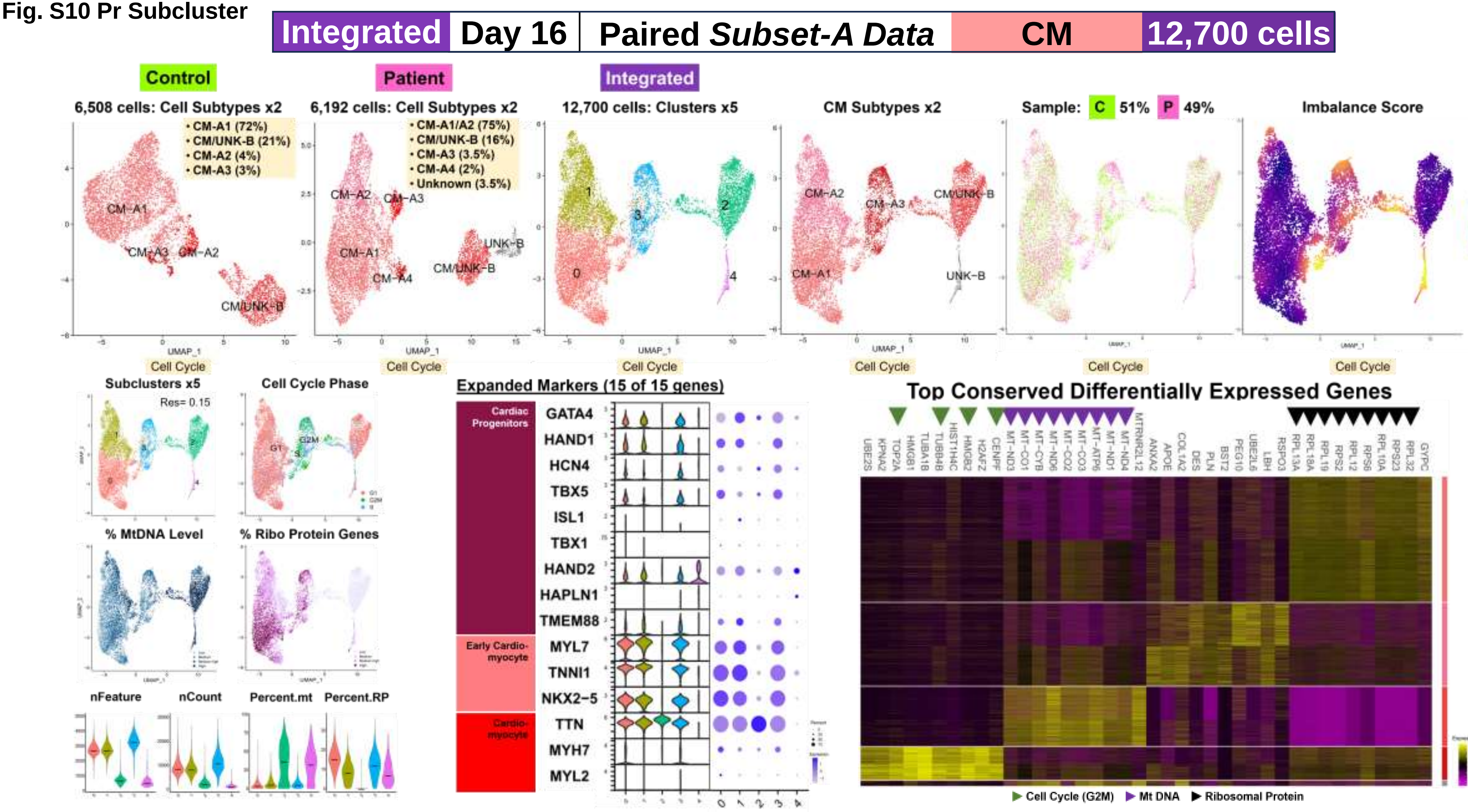

Fig. S10 Pr Subcluster

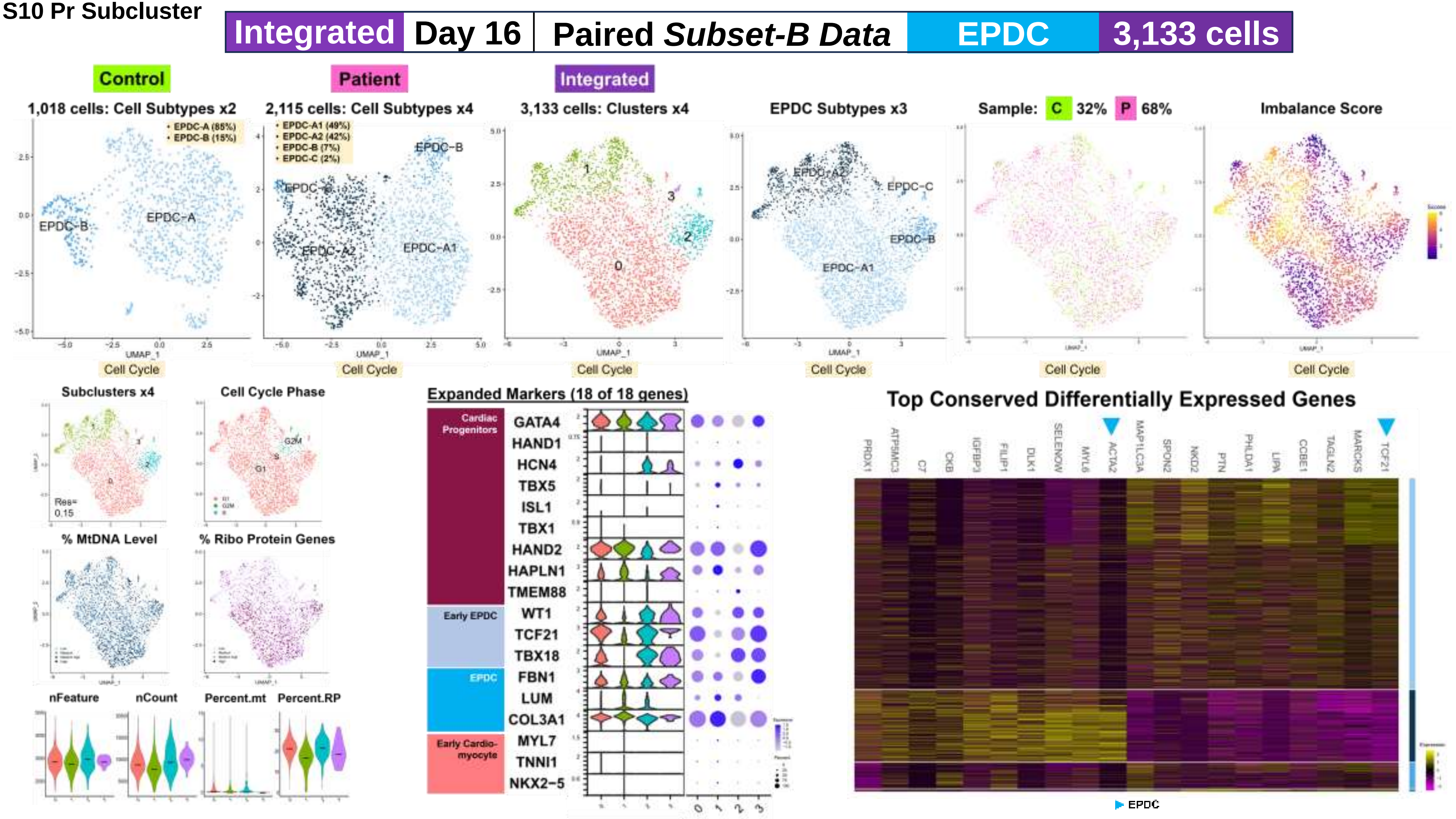

**Fig. S10 Pr Subcluster**

**Integrated | Day 16 | Paired *Subset-C Data* | Unknown | 2,343 cells**

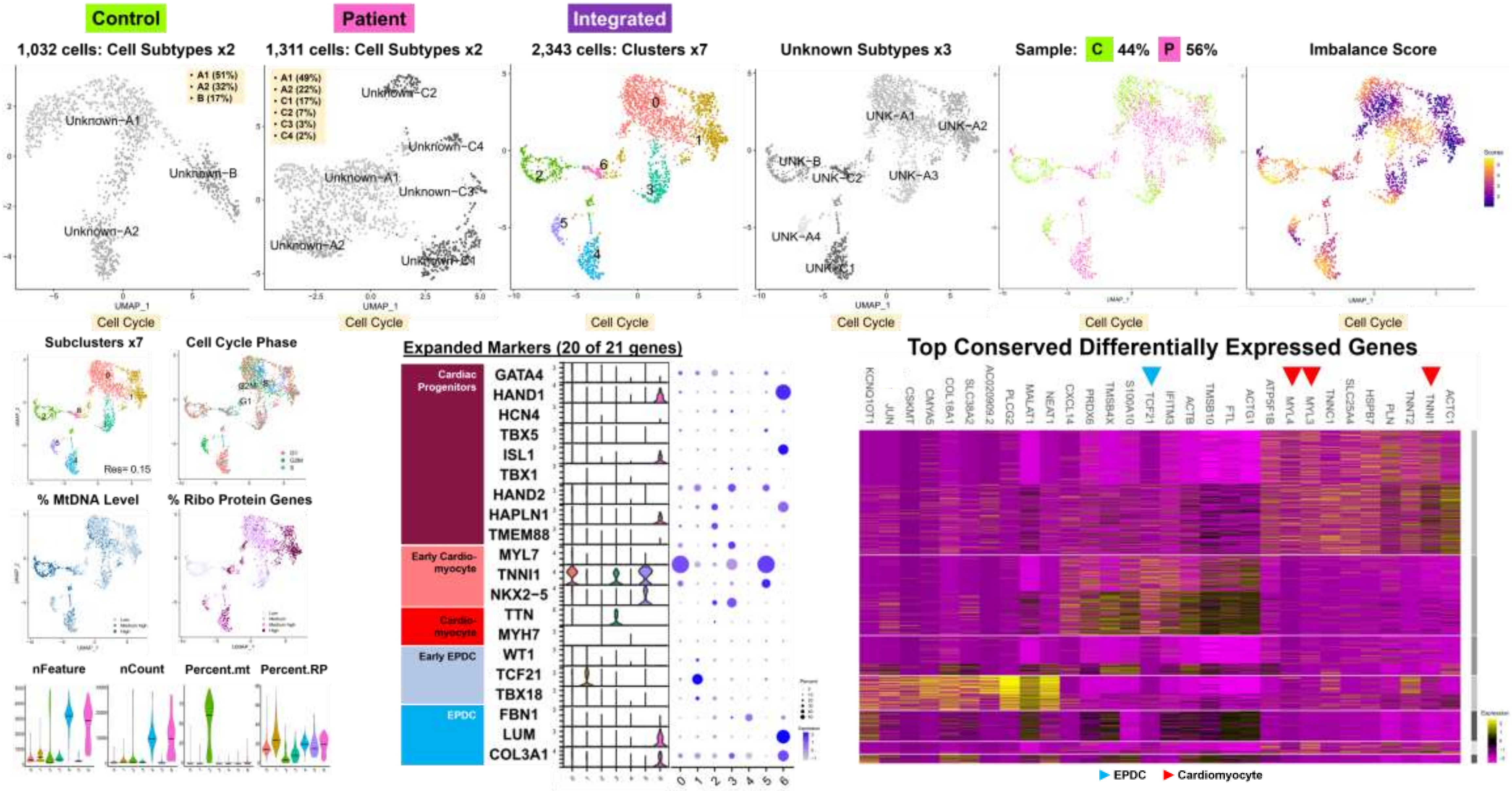

**Fig. S10 Pr Subcluster**

| Integrated | Day 19 | Paired *Subset-A Data* | CM | 15,223 cells |
|---|---|---|---|---|

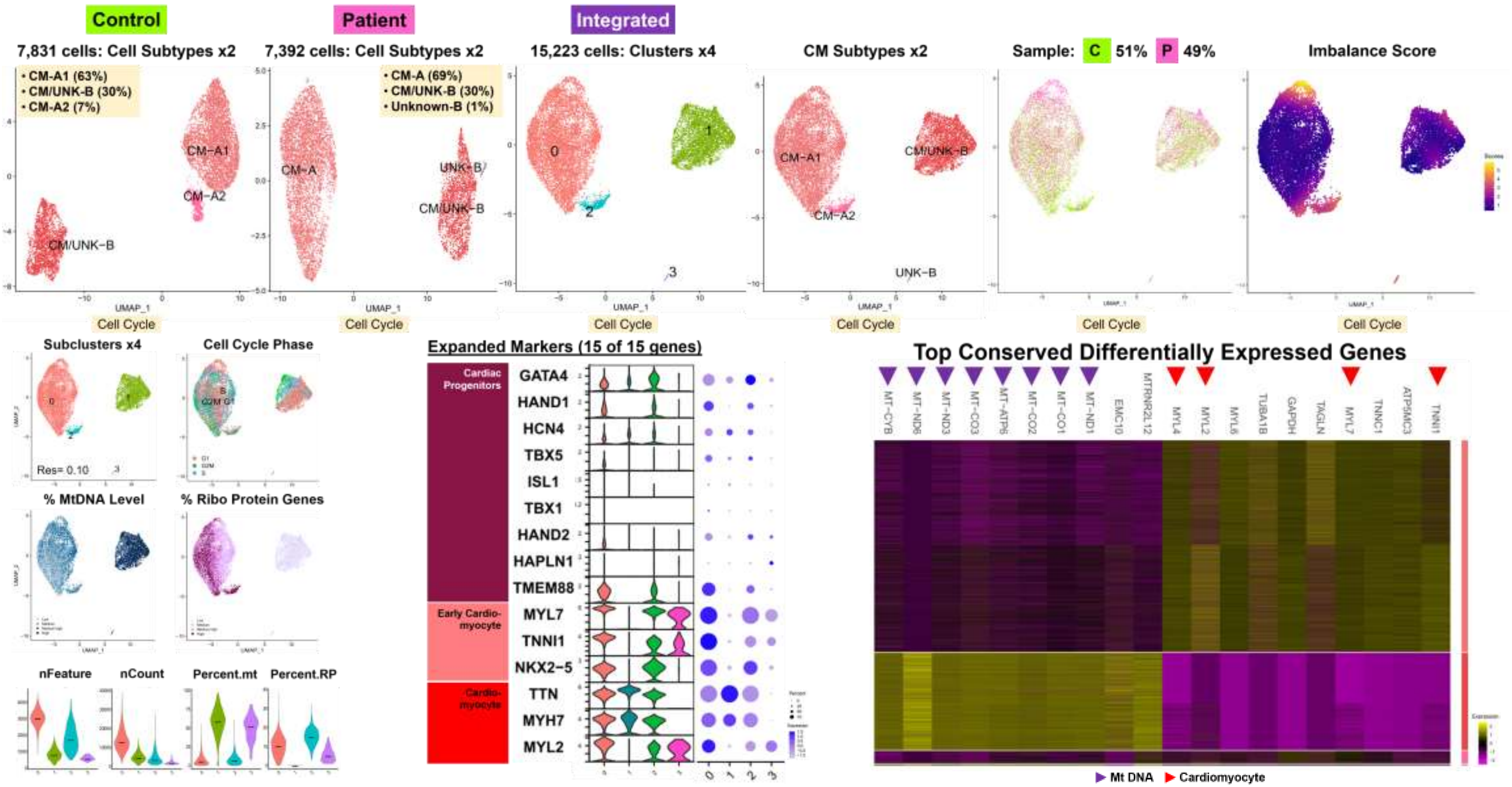

**Fig. S10 Pr Subcluster**

| Integrated | Day 19 | Paired *Subset-B Data* | EPDC | 2,258 cells |
|---|---|---|---|---|

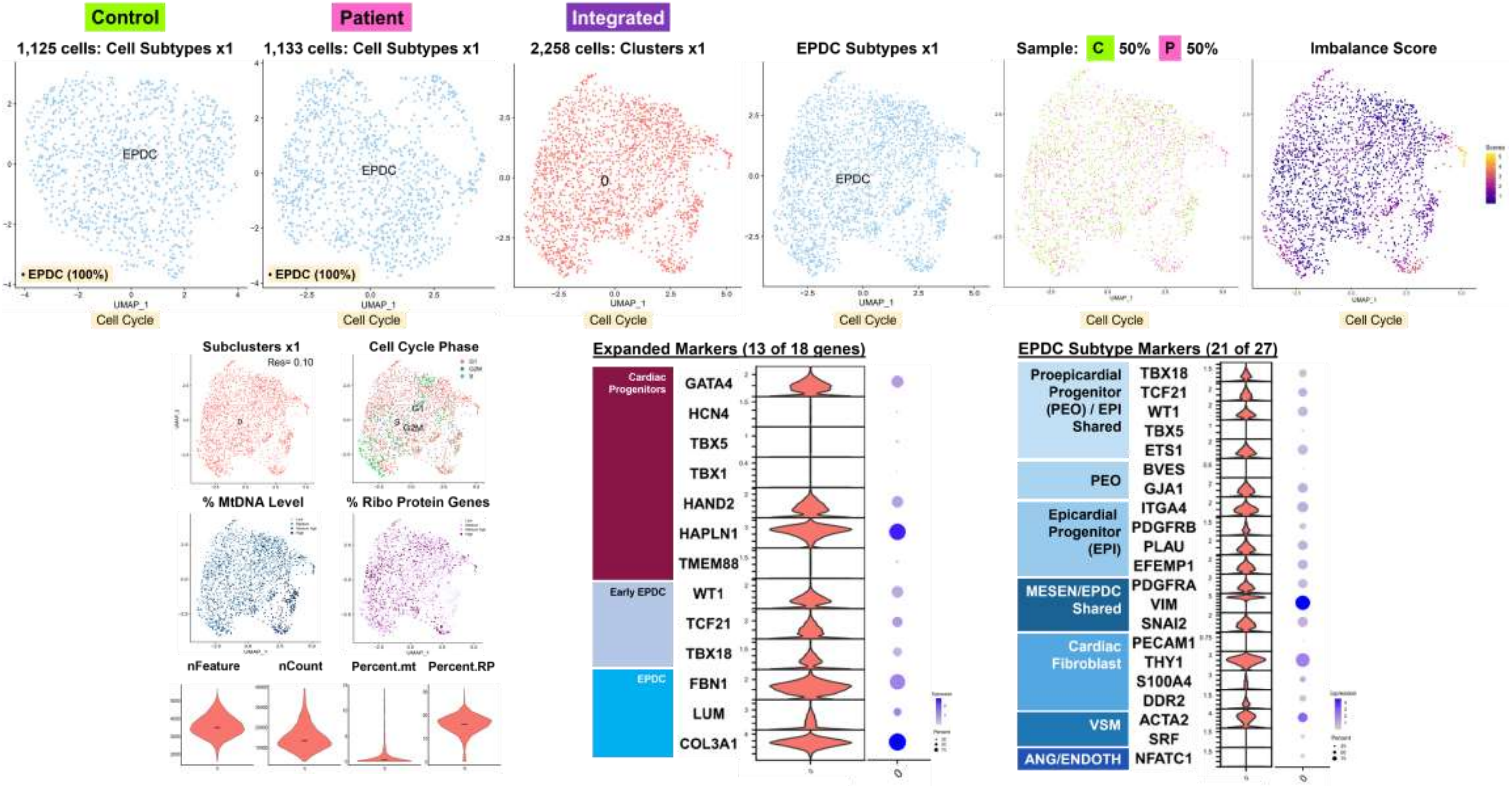

Fig. S10 Pr Subcluster

| Integrated | Day 19 | Paired *Subset-C Data* | Unknown | 3,467 cells |
|---|---|---|---|---|

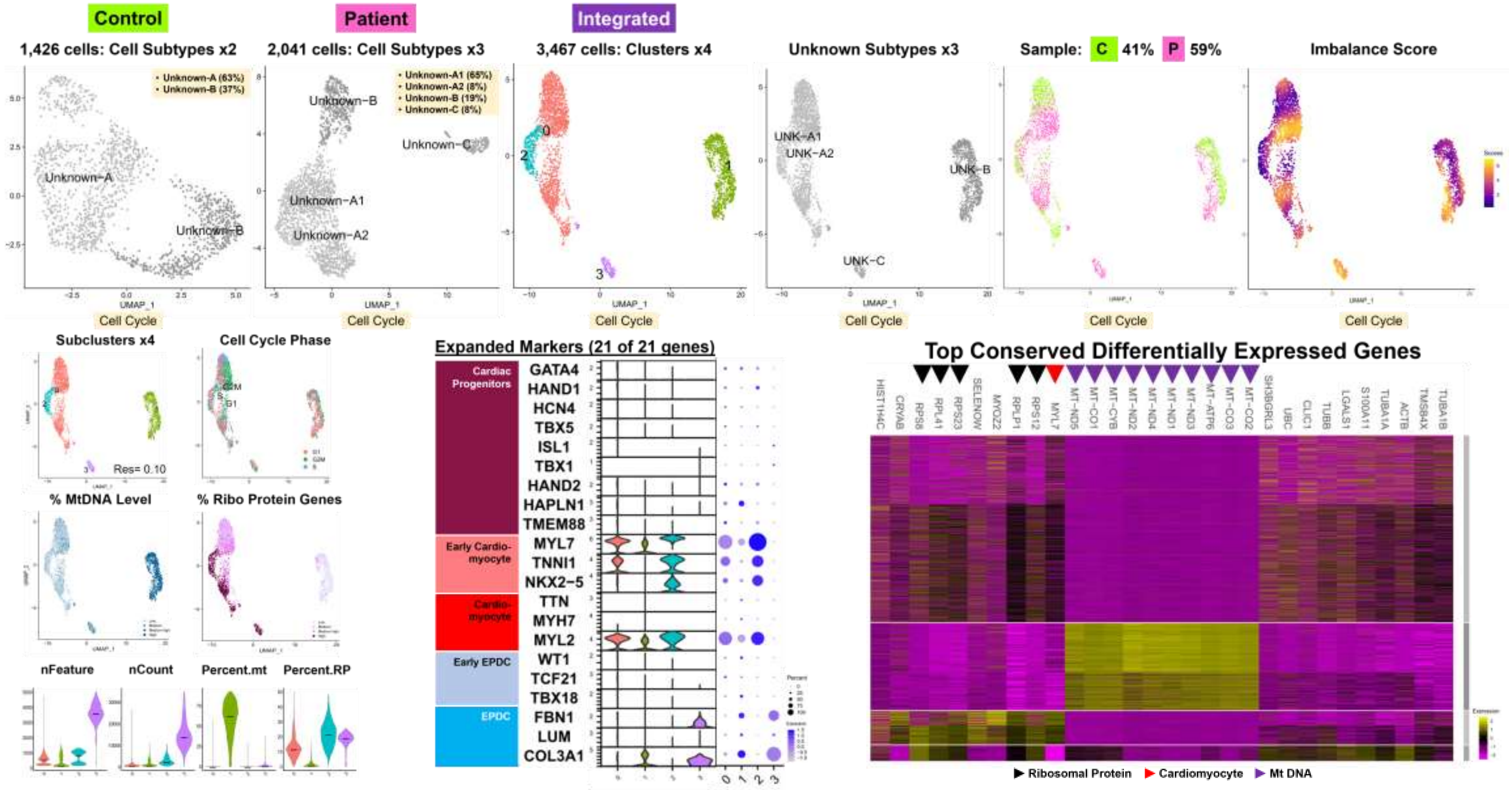

Fig. S11 Cell Type DEG

### Workflow Step-III: Paired Sample Data Results

#### Comparative Analyses for Cell Type-Specific Differential Expression

##### A. Summary: Cell Type Differentially Expressed Genes (Cell Type DEG)

###### Integrated Subset Data: Individual Analysis of ‘Balanced’ Cell Subtypes (n=14: 71,541 Total Cells)

| Day | INTEGR Subset[a] | Cells | PA1 | CA1 | Threshold DEGs[b] Total | Threshold DEGs[b] Under | Threshold DEGs[b] Over | ORA[c] GS Under | ORA[c] GS Over | GO Biological Processes: Top Enriched GS[c] | | DEGs-NoThreshold Total | DEGs-NoThreshold Under | DEGs-NoThreshold Over | GSEA[d] GS | GSEA MSigDB Hallmark GS: Enriched GS[d] |
|---|---|---|---|---|---|---|---|---|---|---|---|---|---|---|---|---|
| 0 | PP-A1A2 | 18,554 | 48% | 52% | 75 | 22 | 53 | 6 | 0 | Ox Phos | | 6,395 | 3,022 | 3,373 | 0 | |
| | PP-B | 716 | 54% | 46% | 3 | 3 | 0 | 4 | 0 | RNA Splicing/Dosage Comp | | 5,942 | 2,934 | 3,008 | 0 | |
| 9B | CPROG-A | 5,946 | 49% | 51% | 82 | 20 | 62 | 2 | 33 | Musc Dev/Diff | Ribosome Process | 3,913 | 2,154 | 1,759 | 1 | OX PHOS |
| | CPROG-B | 4,750 | 53% | 47% | 15 | 6 | 9 | 2 | 2 | Cell Recog/Epigen | Musc Dev/Myofibril | 3,259 | 1,779 | 1,480 | 0 | |
| | CPROG-C | 1,216 | 64% | 36% | 2 | 2 | 0 | - | 0 | | | 4,697 | 2,511 | 2,186 | 0 | |
| | CM-A | 3,433 | 35% | 65% | 26 | 12 | 14 | 0 | 0 | | | 4,018 | 2,205 | 1,813 | 0 | |
| | CM/U-B1B2 | 4,127 | 54% | 46% | 5 | 3 | 2 | 19 | - | Gene Silencing/Dosage Comp | | 167 | 120 | 47 | 0 | |
| | EPDC | 1,728 | 56% | 44% | 7 | 3 | 4 | 26 | 45 | Dosage Comp | Glycolysis | 3,674 | 1,879 | 1,795 | 0 | |
| 16 | CM-A1A2 | 8,639 | 49% | 51% | 475 | 136 | 339 | 32 | 131 | Musc Dev/Signal | Musc Dev/Ox Phos | 4,506 | 2,773 | 1,733 | 5 | TGFB SIGNAL/GLY/OXPHOS/EMT/MYC |
| | CM/U-B | 2,472 | 44% | 56% | 48 | 3 | 45 | 3 | 132 | Dosage Comp | Metab/Musc Dev | 619 | 398 | 221 | 2 | GLYCOLYSIS / MYOGENESIS |
| | EPDC | 3,133 | 68% | 32% | 80 | 12 | 68 | 0 | 38 | Purine Ribonucleotide Metab | | 4,594 | 2,409 | 2,185 | 1 | MTORC1 SIGNAL |
| 19 | CM-A1 | 9,978 | 51% | 49% | 392 | 165 | 227 | 22 | 118 | Musc Diff/Myofibril | Musc Contraction | 4,953 | 2,334 | 2,619 | 1 | MYOGENESIS |
| | CM/U-B | 4,591 | 48% | 52% | 47 | 8 | 39 | 5 | 72 | Signal/Vasc Dev | Musc Contraction | 965 | 395 | 570 | 0 | |
| | EPDC | 2,258 | 50% | 50% | 22 | 5 | 17 | 34 | 0 | Epigen / Heterochromatin | | 5,846 | 3,084 | 2,762 | 0 | |
| | Total | 71,541 | | | 1,279 | 400 | 879 | 155 | 571 | | Total | 53,548 | 27,997 | 25,551 | 10 | |
| | Average | 5,110 | 52% | 48% | 91 | 29 | 63 | 12 | 44 | | Average | 3,825 | 2,000 | 1,825 | 0.7 | |
| | Median | 4,127 | | | 47 | 8 | 39 | 5 | 36 | | Median | 4,506 | 2,334 | 1,813 | 0 | |

[a]Integrated (INTEGR) Data Subsets with primarily ‘balanced’ shared cell subtypes (n=14) used for Cell Type Differential Expression Analyses

[b]DEG Threshold: Adjusted p-value < 10e-50 & I Avg Log2 FC I > 0.25, Fold Change = 1.2x

[c]ORA Parameters: min # genes= 3, Adj p-value < 0.05, q-value cutoff 0.20; Highlighted in Red= Underexpressed GS and Green=Overexpressed GS in Patient cells compared to Control cells

[d]GSEA Parameters: Adj p-value cutoff= 0.05, GS size 10-500 genes; Highlighted in Pink= Downregulated GS; Bright Green= Upregulated GS in Patient compared to Control cells

Abbreviations: PP, Pluripotent; CPROG, Cardiac Progenitors; CM, Cardiomyocyte; EPDC, Epicardium-derived cells; U, Unknown; DEG, differentially expressed gene; ORA, Over-Representation Analysis; GS, Gene Set; GSEA, Gene Set Enrichment Analysis; MSigDB, Molecular Signatures Database; Ox Phos, Oxidative Phosphorylation; Comp, Compensation; Musc, Muscle; Dev, Development; Diff, Differentiation; Recog, Recognition; Epigen, Epigenetic; Metab, Metabolism; Signal, Signaling; Vasc, Vascular; EMT, Epithelial-Mesenchymal Transition; MYC, MYC Targets; Gly, Glycolysis

**Fig. S11 Cell Type DEG**

### B. Summary: Volcano Plots and Cell Type DEG (n=14 Cell Subtypes: 71,541 Total Cells)

**Day 00: x2 Cell Subtypes**

**Day 09B: x3 Cell Subtypes**

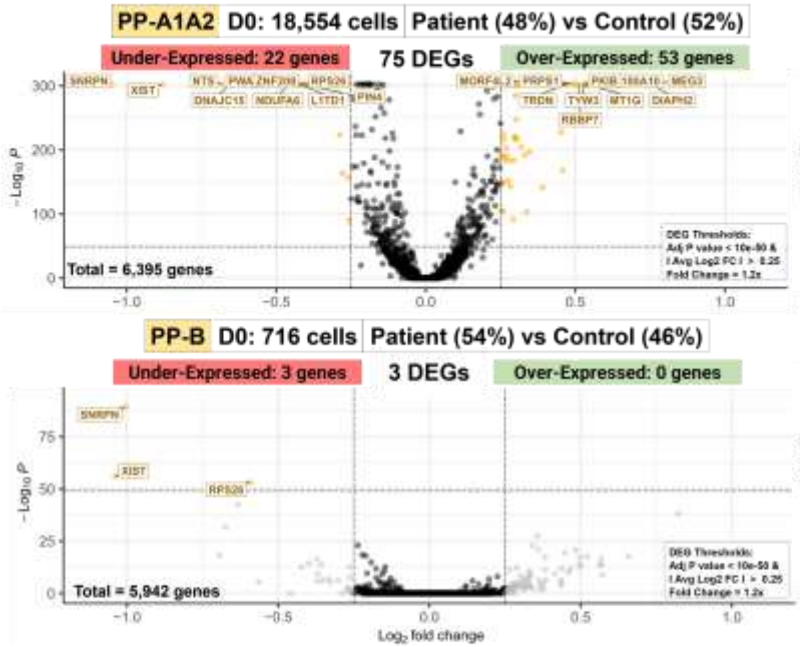


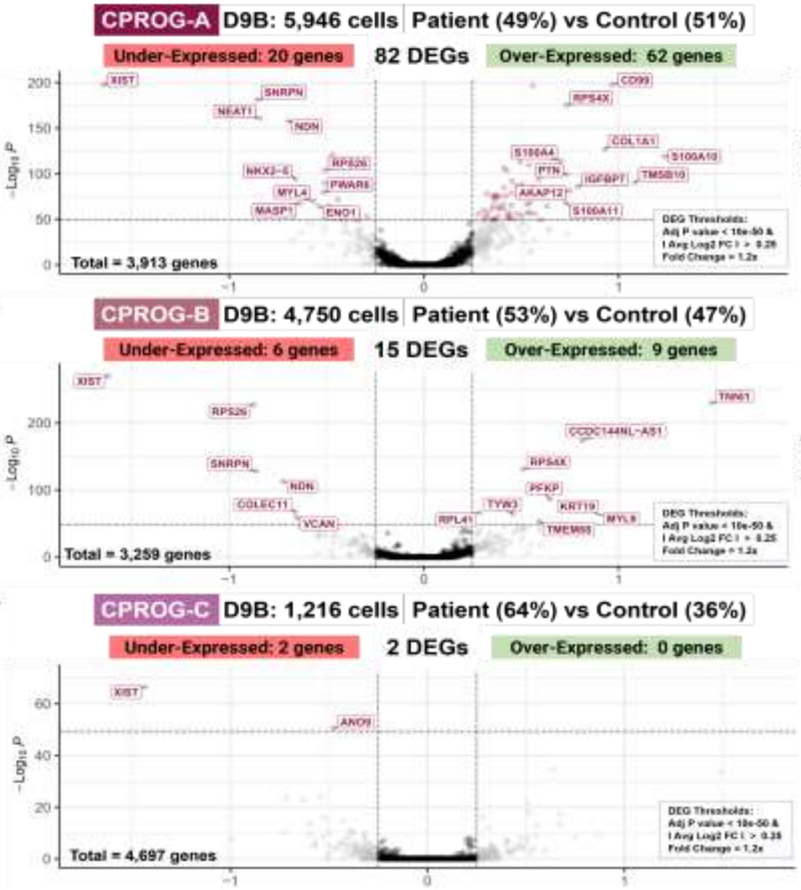


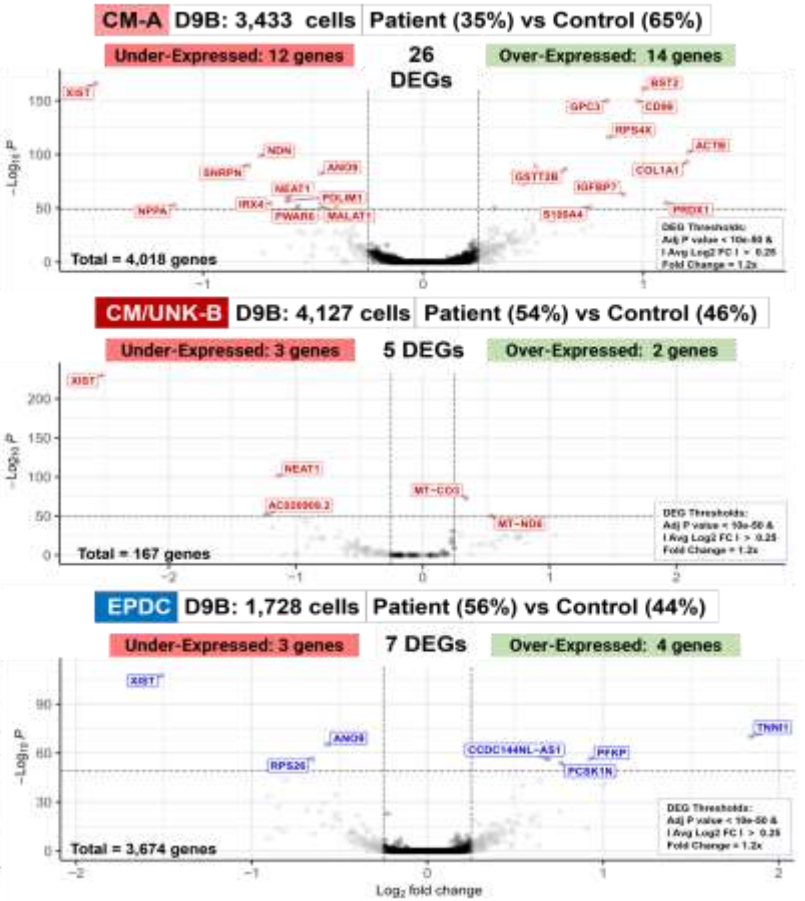


Top 10 Cell Type Threshold DEG are labeled.

#### Fig. S11 Cell Type DEG

##### Day 16: x3 Cell Subtypes

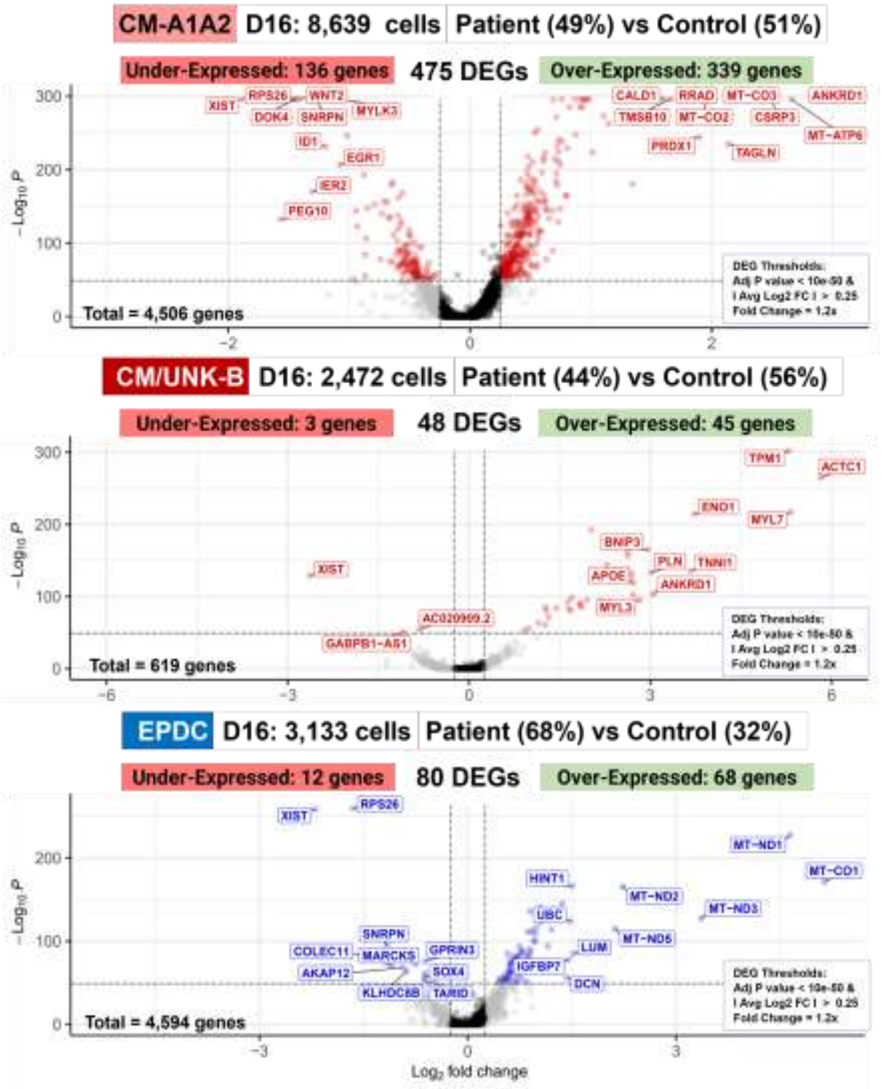


##### Day 19: x3 Cell Subtypes

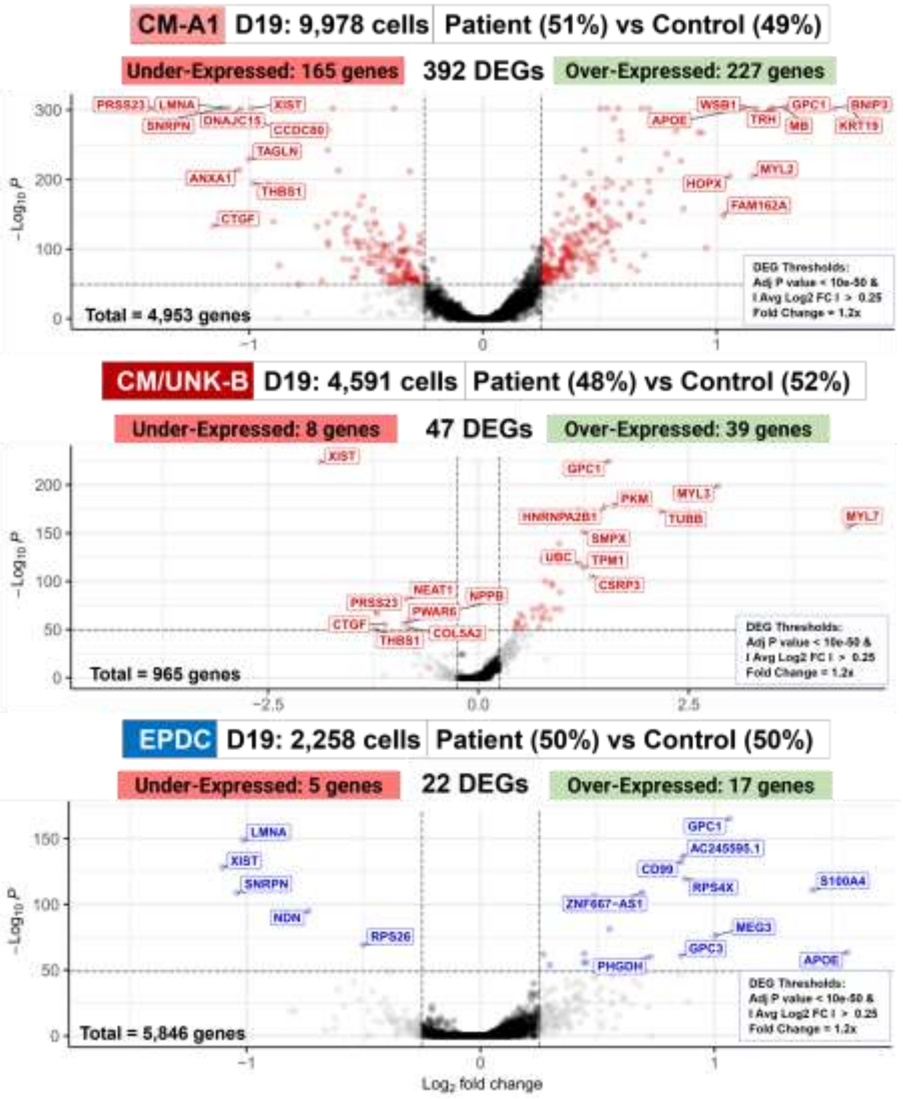

Fig. S11 Cell Type DEG

#### C. Individual Analyses of Integrated Subsets: Cell Type DE Analyses and Enrichment (n=14)

| Integrated | D00 | PP-A1A2 | 18,554 cells |
|---|---|---|---|

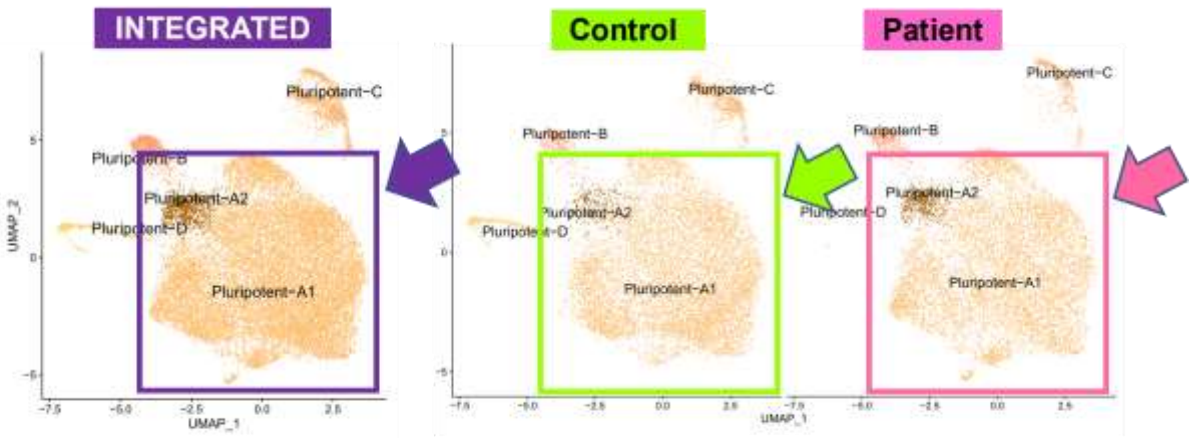


| PP-A1A2 | D0: 18,554 cells | Patient (48%) vs Control (52%) |
|---|---|---|

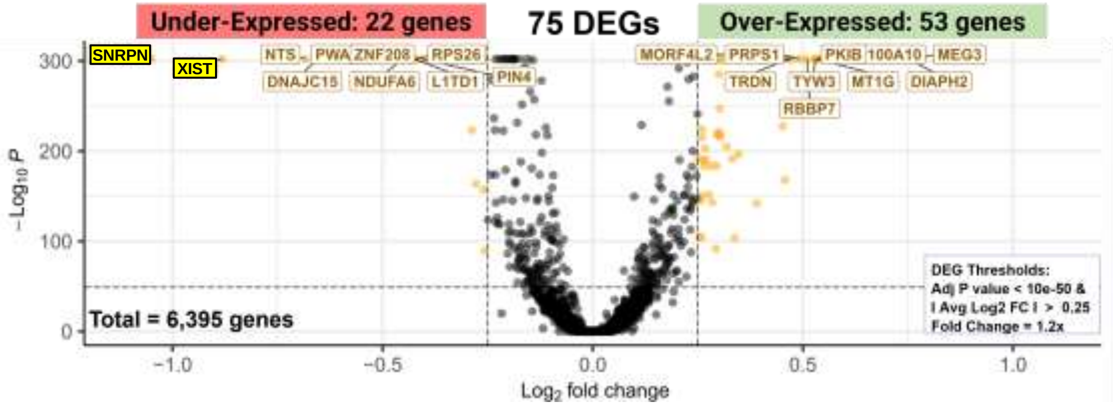


##### Over-Representation Analysis (ORA)

**Under-Expressed: 22 genes** Enriched Gene Sets = 6

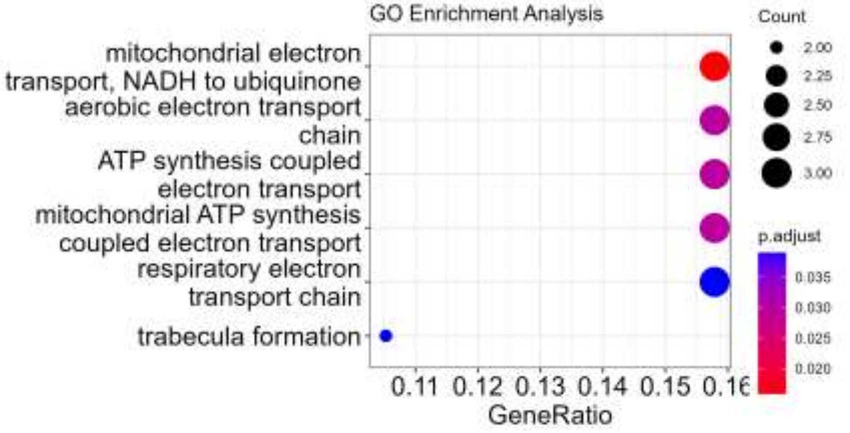


**Over- Expressed: 53 genes** Enriched Gene Sets = 0

##### Enrichment Map

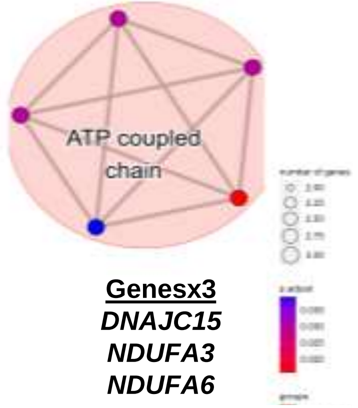


**Genesx3**
*DNAJC15*
*NDUFA3*
*NDUFA6*

**Genesx2**
*HEY2*
*SFRP1*

trabecula formation

##### Gene Set Enrichment Analysis (GSEA)

**All 6395 DEGs**
**Under-expressed: 3022 genes**
**Over-expressed: 3373 genes**

**No Enriched MSigDB Hallmark Gene Sets**

#### Fig. S11 Cell Type DEG

| Integrated | D00 | PP-B | 716 cells |
|---|---|---|---|

INTEGRATED | Control | Patient

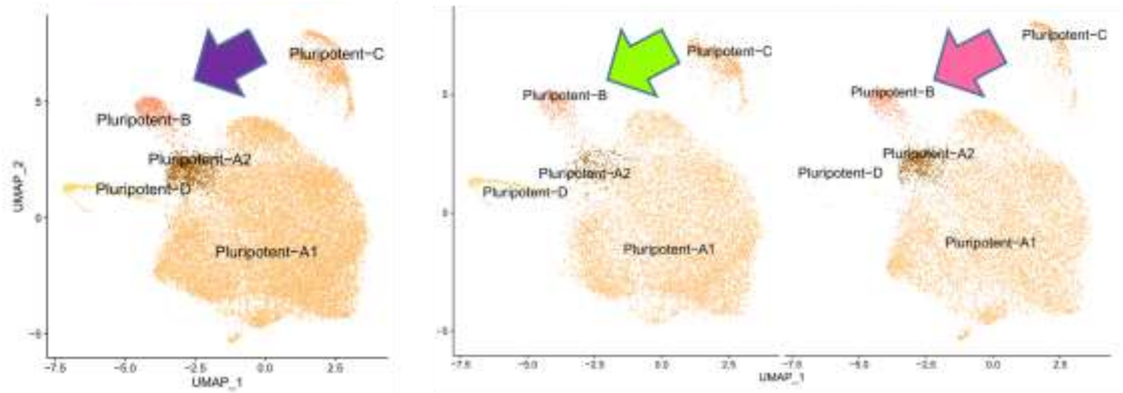


PP-B | D0: 716 cells | Patient (54%) vs Control (46%)

Under-Expressed: 3 genes | 3 DEGs | Over-Expressed: 0 genes

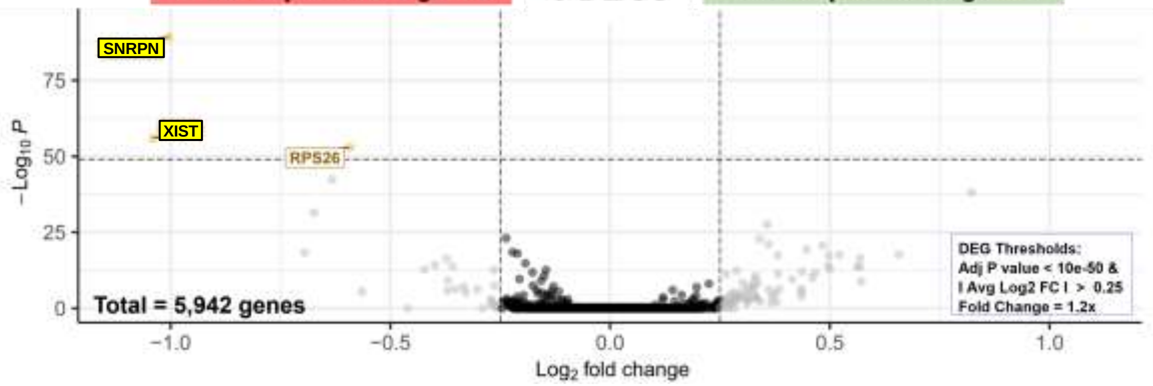


##### Over-Representation Analysis (ORA)

**Under-Expressed: 3 genes** Enriched Gene Sets = 4

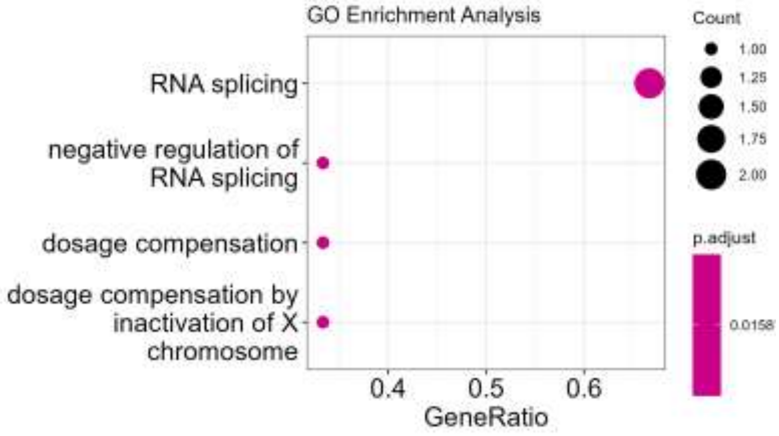


**Enrichment Map**

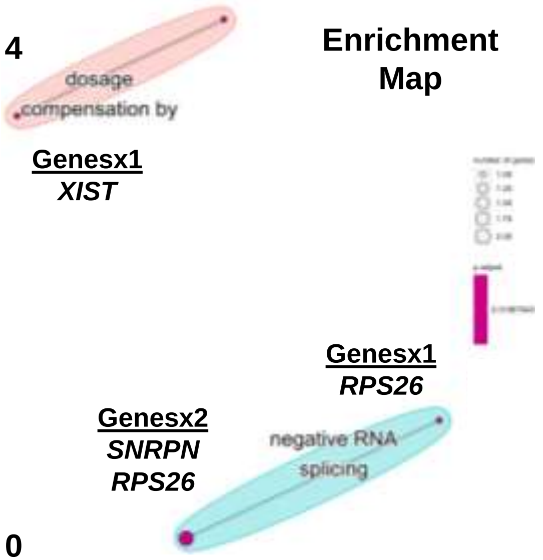


**Over- Expressed: 0 genes** Enriched Gene Sets = 0

##### Gene Set Enrichment Analysis (GSEA)

**All 5942 DEGs**
**Under-expressed: 2934 genes**
**Over-expressed: 3008 genes**

**No Enriched MSigDB Hallmark Gene Sets**

### Fig. S11 Cell Type DEG

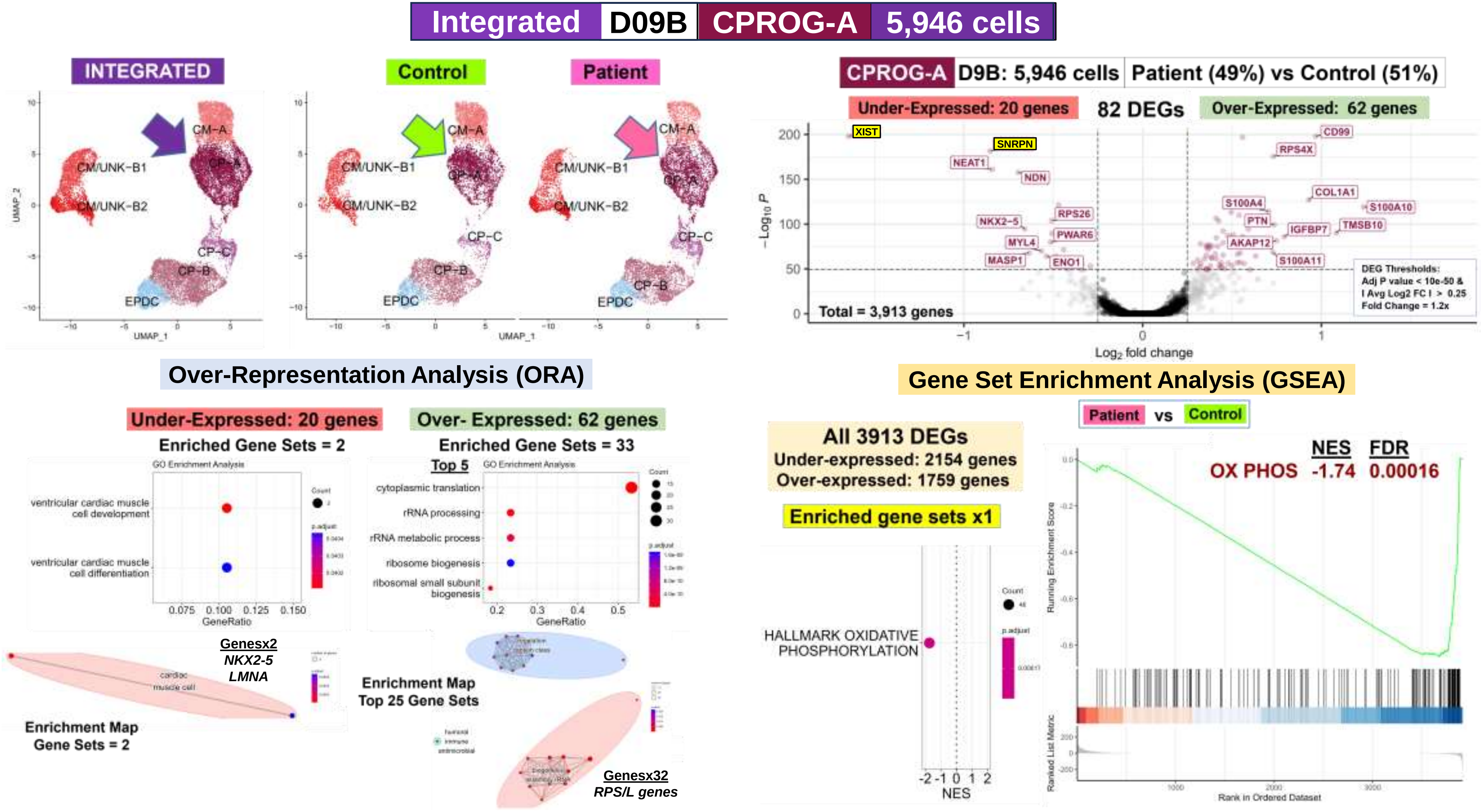

### Fig. S11 Cell Type DEG

**Integrated | D09B | CPROG-B | 4,750 cells**

INTEGRATED | Control | Patient

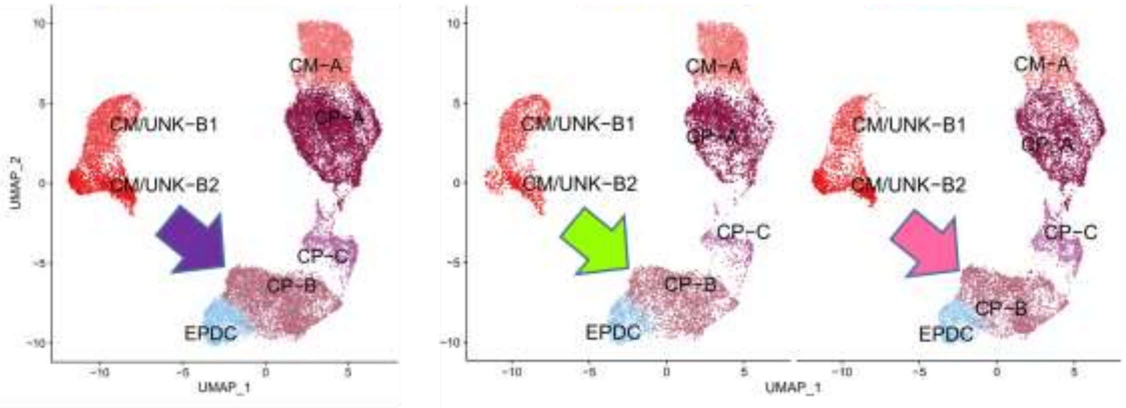


CPROG-B | D9B: 4,750 cells | Patient (53%) vs Control (47%)

Under-Expressed: 6 genes | 15 DEGs | Over-Expressed: 9 genes

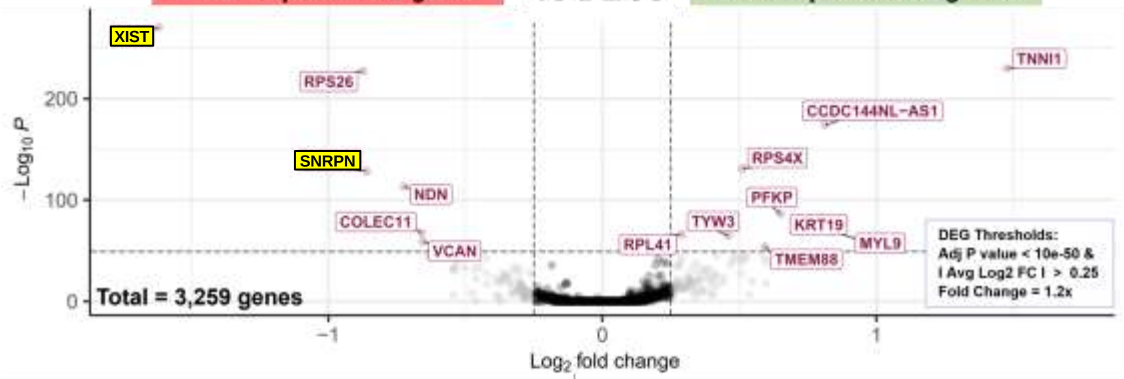


#### Over-Representation Analysis (ORA)

**Under-Expressed: 6 genes**

Enriched Gene Sets = 2

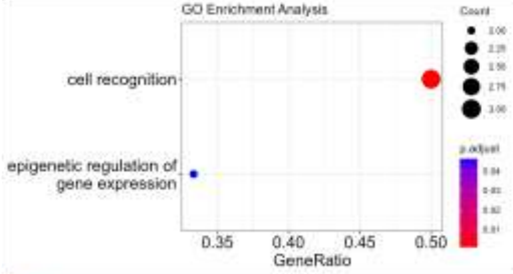


**Over- Expressed: 9 genes**

Enriched Gene Sets = 2

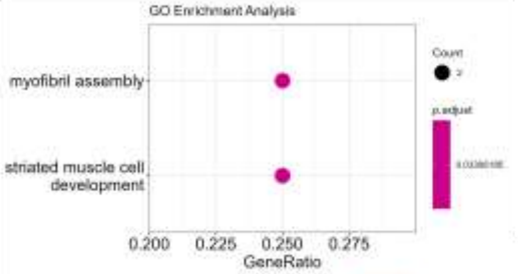


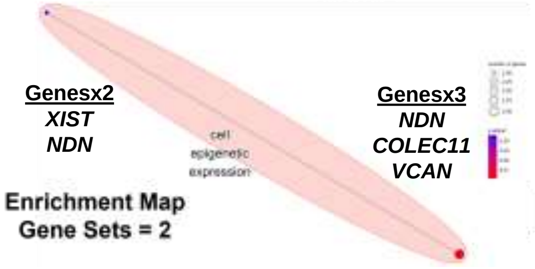


Enrichment Map
Gene Sets = 2

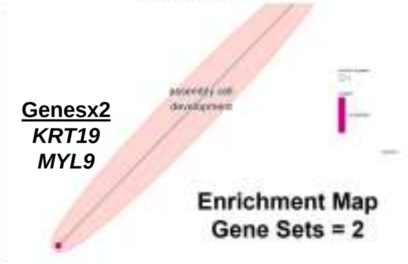


Enrichment Map
Gene Sets = 2

#### Gene Set Enrichment Analysis (GSEA)

**All 3259 DEGs**
**Under-expressed: 1779 genes**
**Over-expressed: 1480 genes**

**No Enriched MSigDB Hallmark Gene Sets**

#### Fig. S11 Cell Type DEG

**Integrated | D09B | CPROG-C | 1,216 cells**

| INTEGRATED | Control | Patient |
|---|---|---|

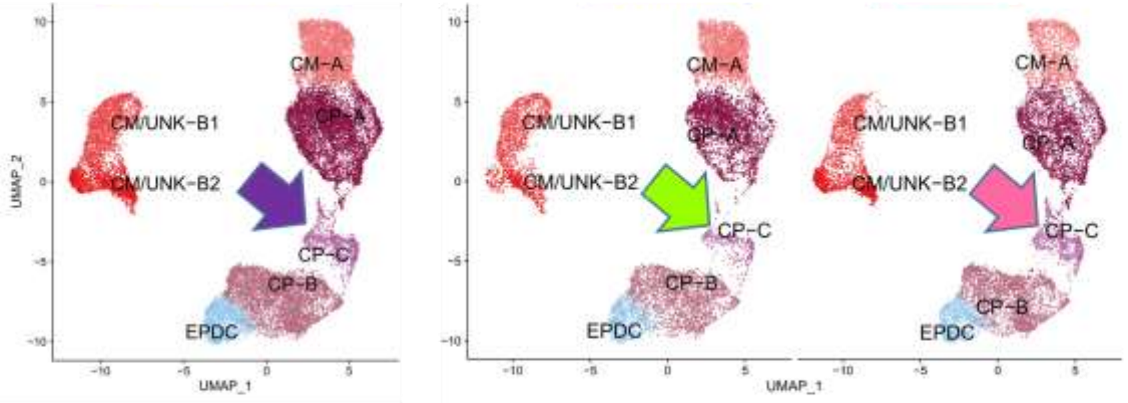


**CPROG-C | D9B: 1,216 cells | Patient (64%) vs Control (36%)**

**Under-Expressed: 2 genes | 2 DEGs | Over-Expressed: 0 genes**

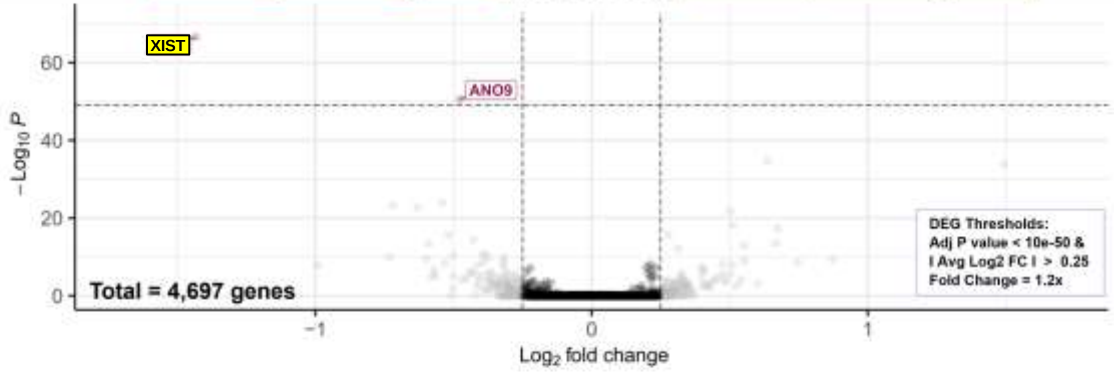


##### Over-Representation Analysis (ORA)

**Under-Expressed: 2 genes**

NOT DONE

**Over- Expressed: 0 genes**

Enriched Gene Sets = 0

##### Gene Set Enrichment Analysis (GSEA)

**All 4697 DEGs**
**Under-expressed: 2511 genes**
**Over-expressed: 2186 genes**

**No Enriched MSigDB Hallmark Gene Sets**

**Fig. S11 Cell Type DEG**

| Integrated | D09B | CM-A | 3,433 cells |
|---|---|---|---|

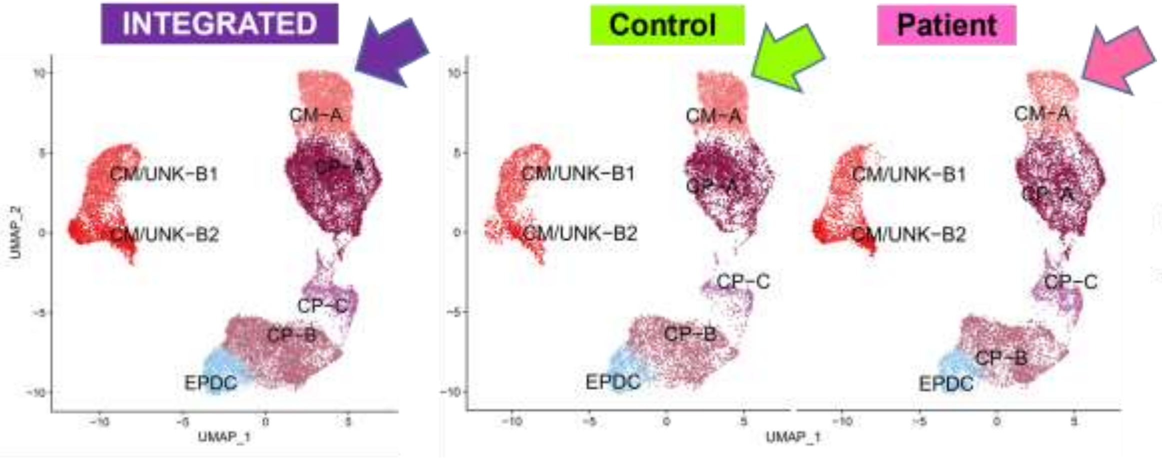


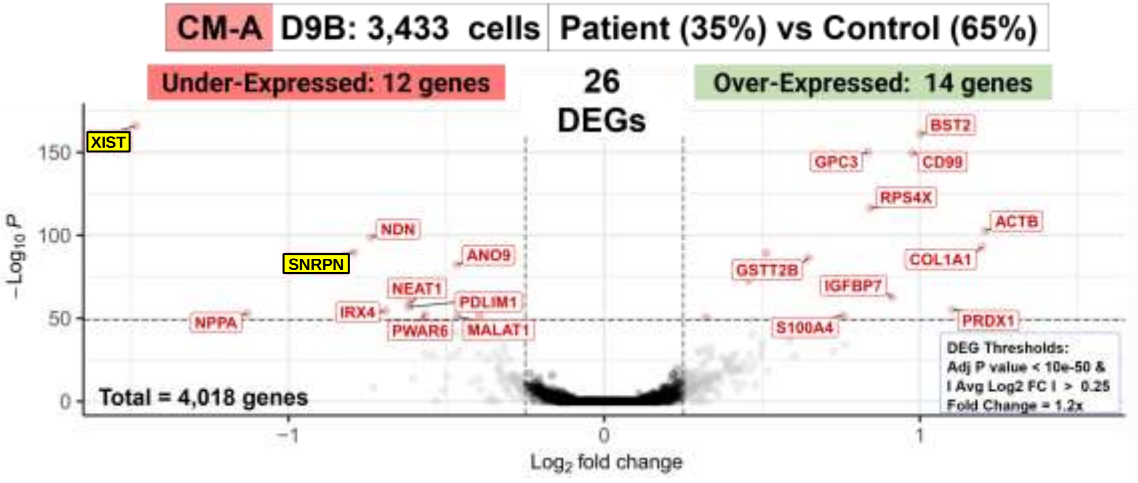


#### Over-Representation Analysis (ORA)

| Under-Expressed: 12 genes | No Enriched GO Biological Process Gene Sets |
|---|---|
| Over-Expressed: 14 genes | No Enriched GO Biological Process Gene Sets |

#### Gene Set Enrichment Analysis (GSEA)

**All 4018 DEGs**
**Under-expressed: 2205 genes**
**Over-expressed: 1813 genes**

**No Enriched MSigDB Hallmark Gene Sets**

#### Fig. S11 Cell Type DEG

**Integrated | D09B | CM/UNK-B | 4,127 cells**

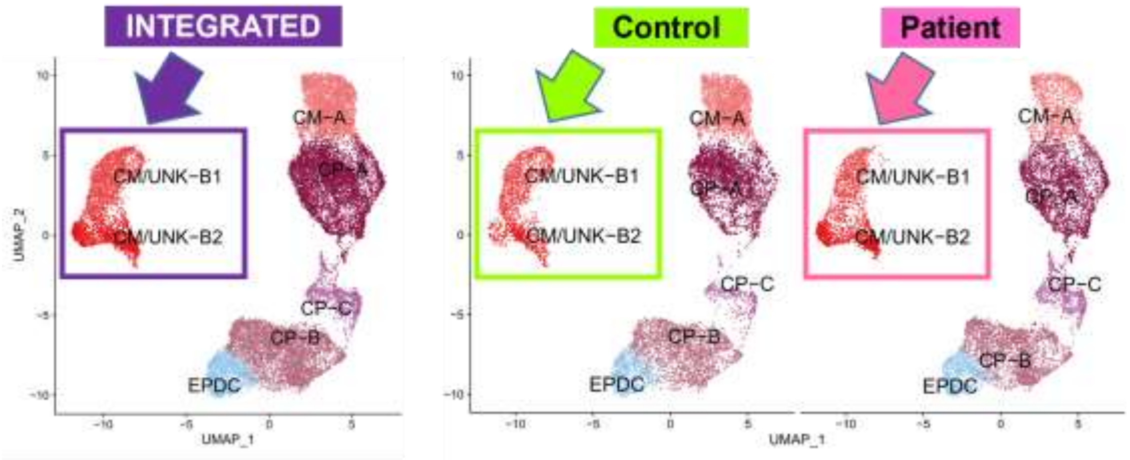


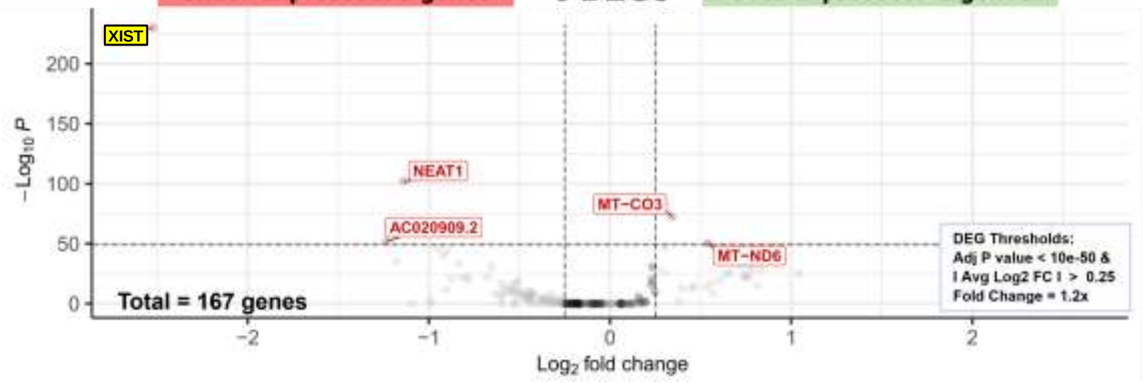


**Over-Representation Analysis (ORA)**

**Under-Expressed: 3 genes**

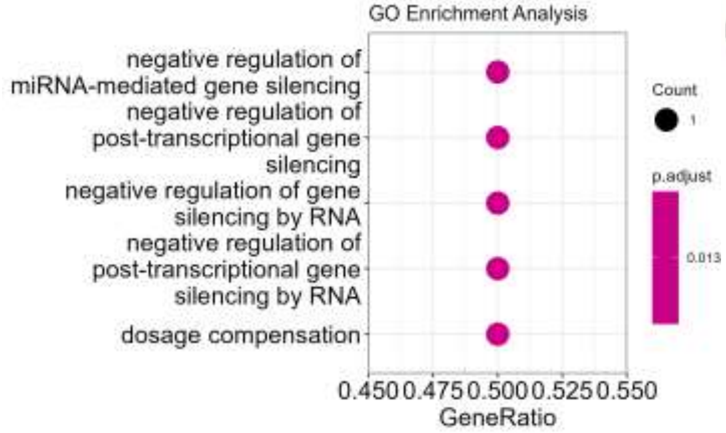


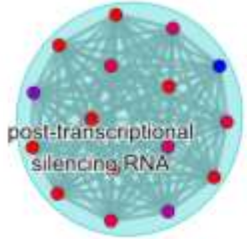


Genesx1
*NEAT1*

**Enrichment Map**

**Enriched Gene Sets = 19**

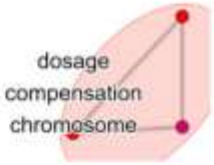


Genesx1
*XIST*

**Over- Expressed: 2 genes** NOT DONE

**Gene Set Enrichment Analysis (GSEA)**

**All 167 DEGs**
**Under-expressed: 120 genes**
**Over-expressed: 47 genes**

**No Enriched MSigDB Hallmark Gene Sets**

#### Fig. S11 Cell Type DEG

| Integrated | D09B | EPDC | 1,728 cells |
|---|---|---|---|

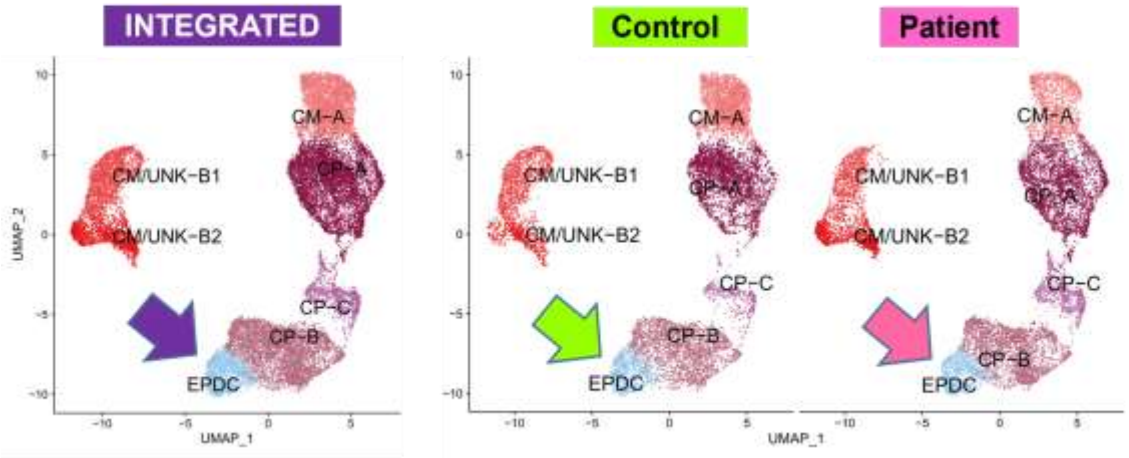


| EPDC | D9B: 1,728 cells | Patient (56%) vs Control (44%) |
|---|---|---|

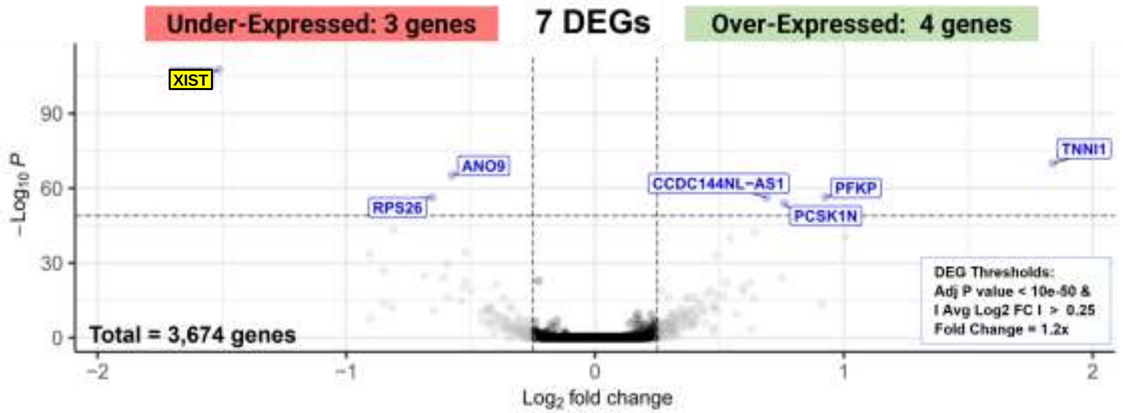


##### Over-Representation Analysis (ORA)

**Under-Expressed: 3 genes**

Enriched Gene Sets = 26

Top 5

**Over- Expressed: 4 genes**

Enriched Gene Sets = 45

Top 5

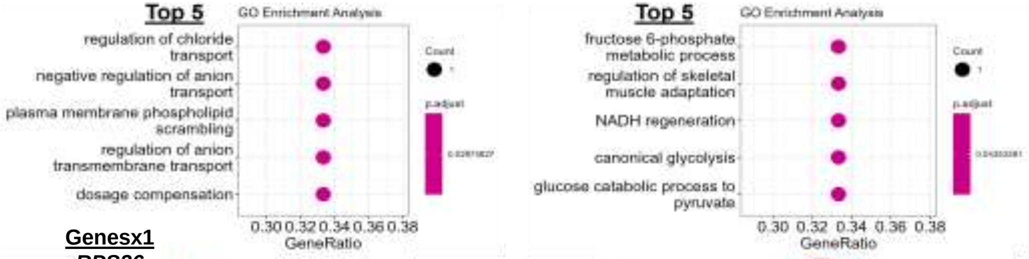


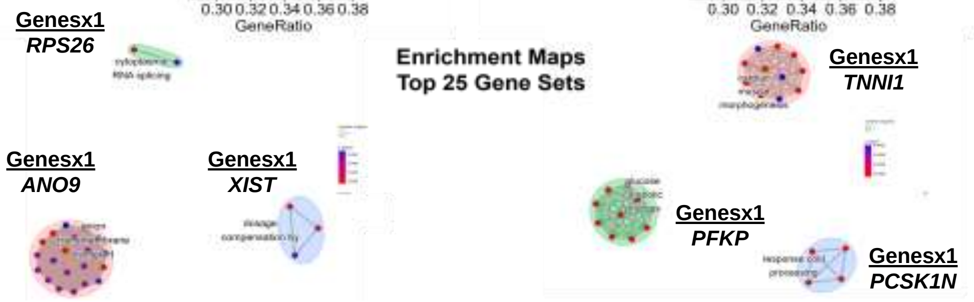


##### Gene Set Enrichment Analysis (GSEA)

**All 3674 DEGs**
**Under-expressed: 1879 genes**
**Over-expressed: 1795 genes**

**No Enriched MSigDB Hallmark Gene Sets**

#### Fig. S11 Cell Type DEG

| Integrated | D16 | CM-A1A2 | 8,639 cells |
|---|---|---|---|

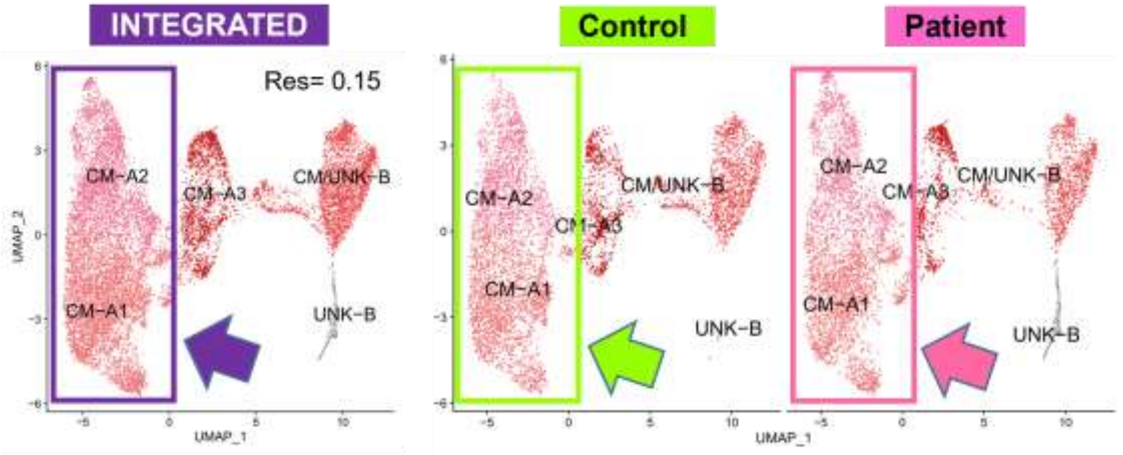


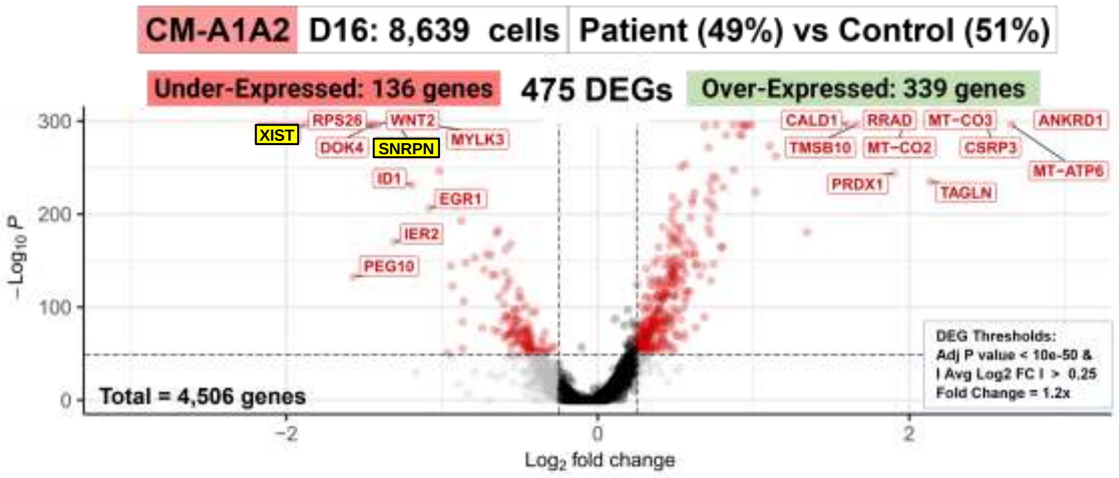


#### Over-Representation Analysis (ORA)

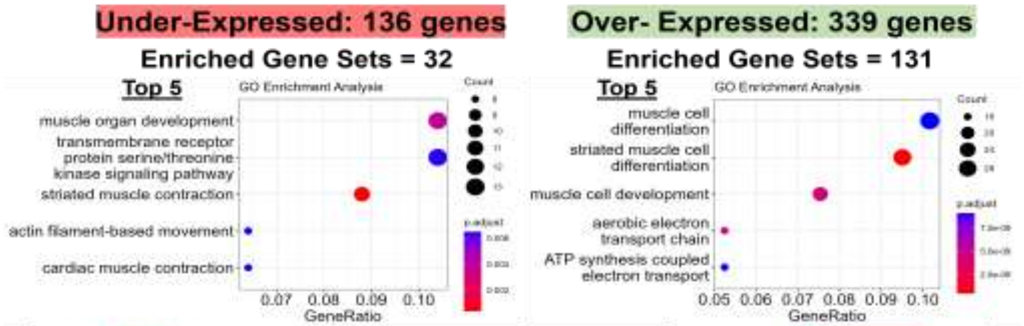


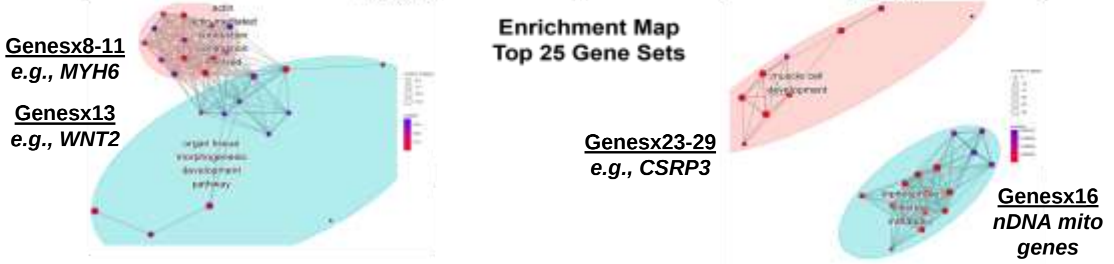


#### Gene Set Enrichment Analysis (GSEA)

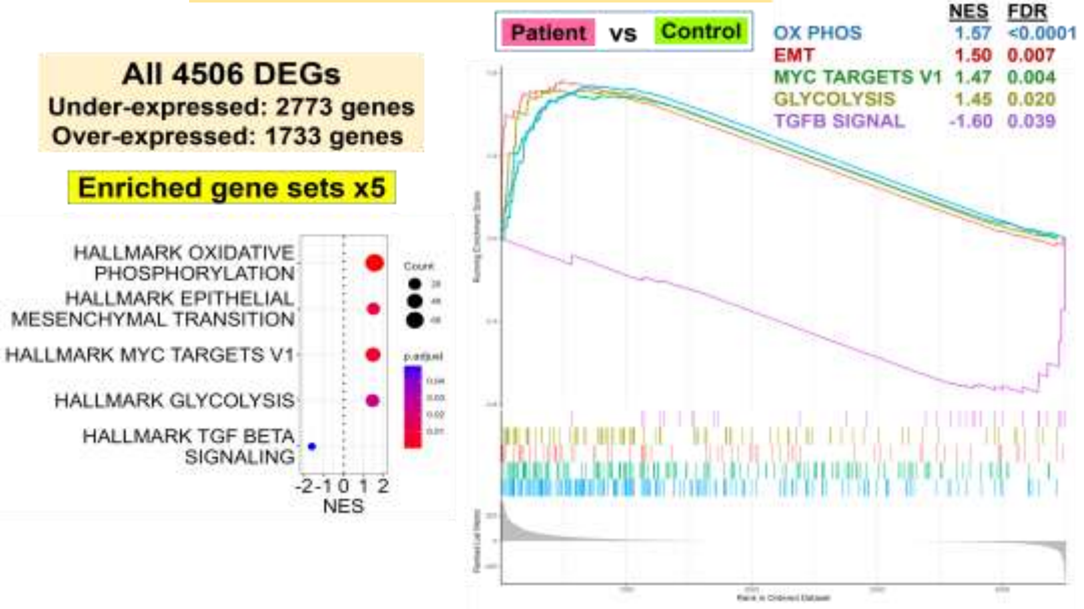

### Fig. S11 Cell Type DEG

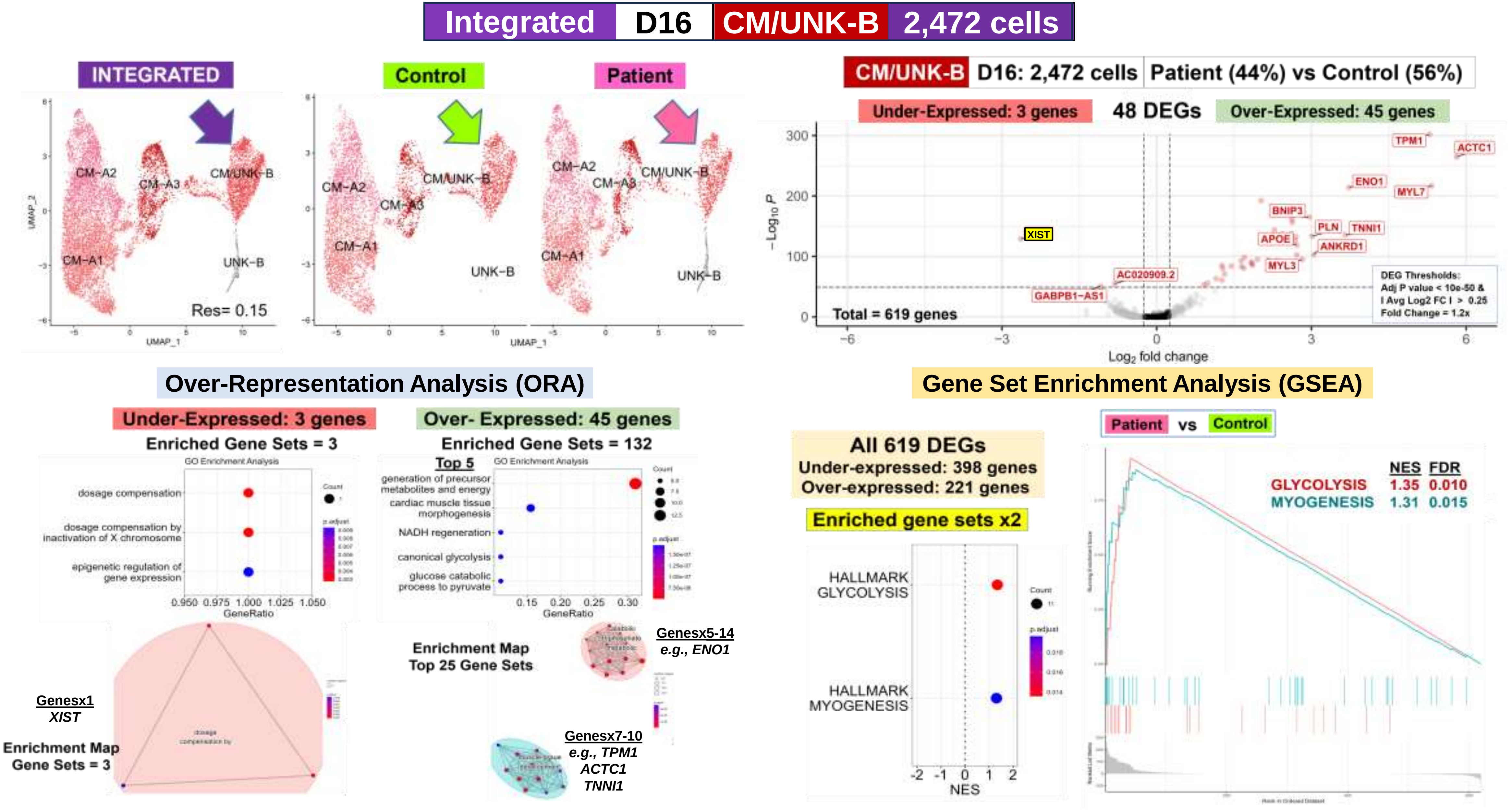

**Fig. S11 Cell Type DEG**

| Integrated | D16 | EPDC | 3,133 cells |
|---|---|---|---|

INTEGRATED | Control | Patient

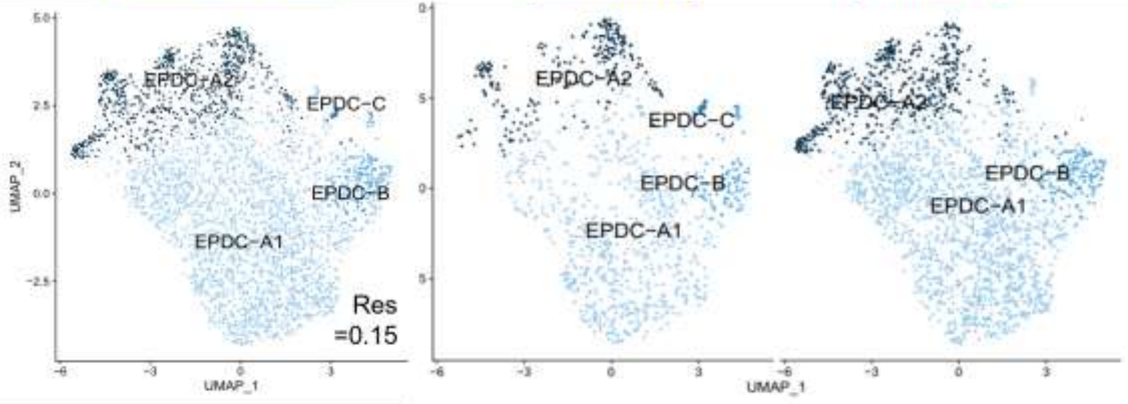


EPDC | D16: 3,133 cells | Patient (68%) vs Control (32%)

Under-Expressed: 12 genes | 80 DEGs | Over-Expressed: 68 genes

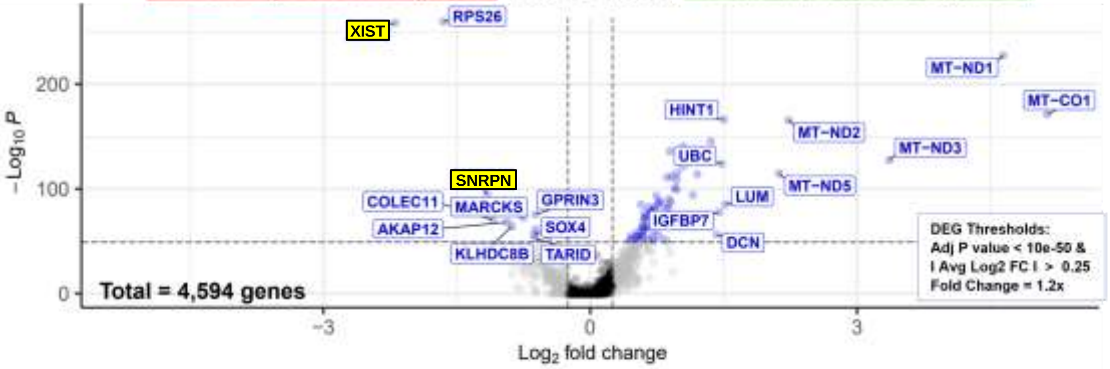


#### Over-Representation Analysis (ORA)

**Under-Expressed: 12 genes**
Enriched Gene Sets = 0

**No Enrichment: GO Biological Process Gene Sets**

**Over- Expressed: 68 genes**
Enriched Gene Sets = 38

**Top 5**

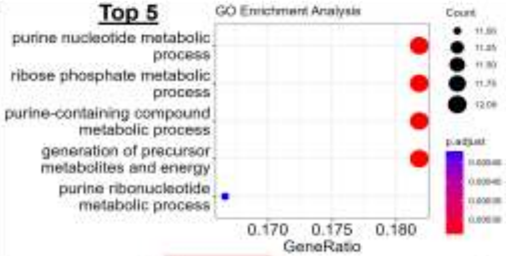


**Enrichment Map**
**Top 25 Gene Sets**

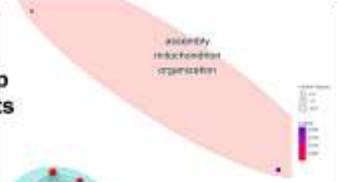

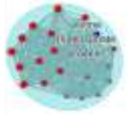

Genesx12
*e.g., HINT1*

#### Gene Set Enrichment Analysis (GSEA)

**All 4594 DEGs**
**Under-expressed: 2409 genes**
**Over-expressed: 2185 genes**

**Enriched gene sets x1**

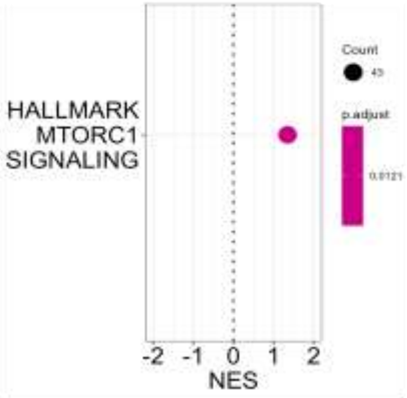


Patient vs Control

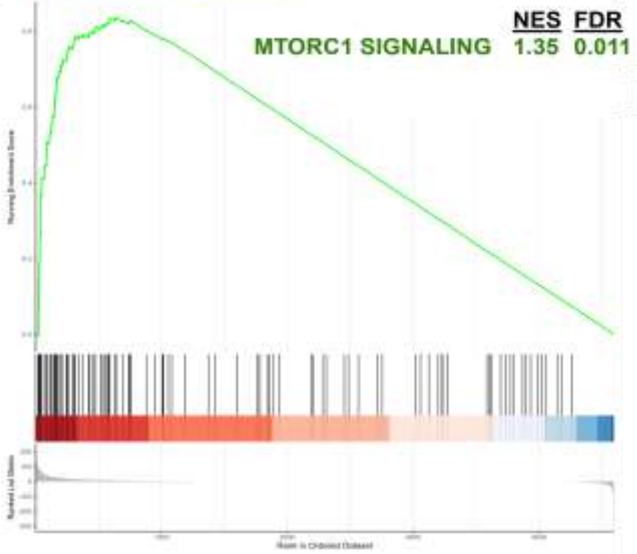

Fig. S11 Cell Type DEG

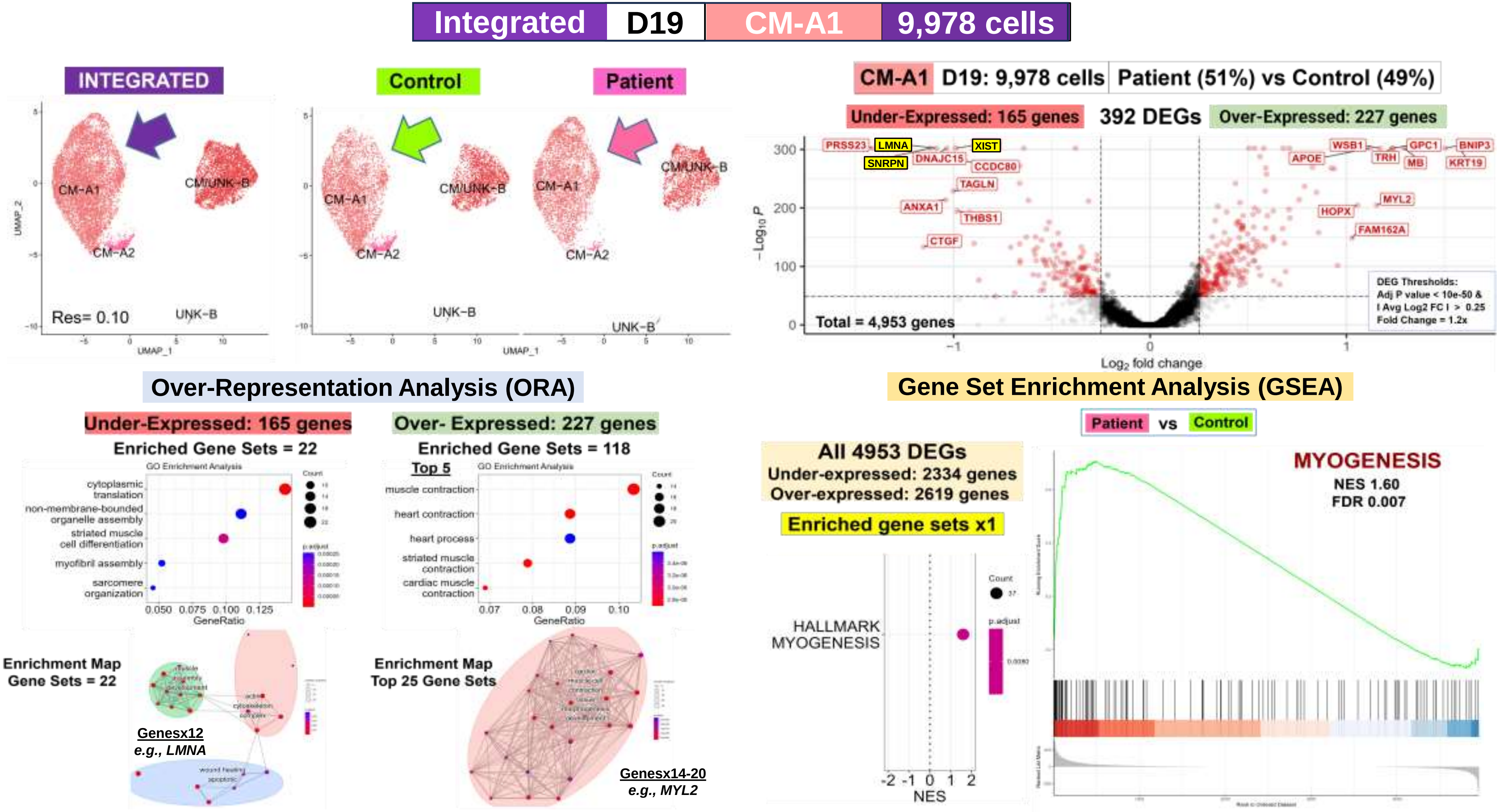

### Fig. S11 Cell Type DEG

Integrated | D19 | CM/UNK-B | 4,591 cells

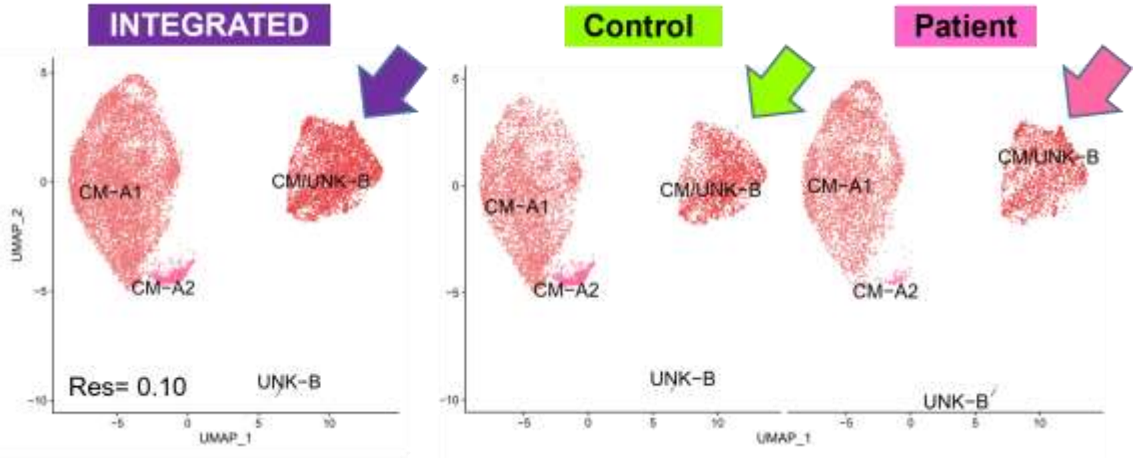


CM/UNK-B | D19: 4,591 cells | Patient (48%) vs Control (52%)

Under-Expressed: 8 genes | 47 DEGs | Over-Expressed: 39 genes

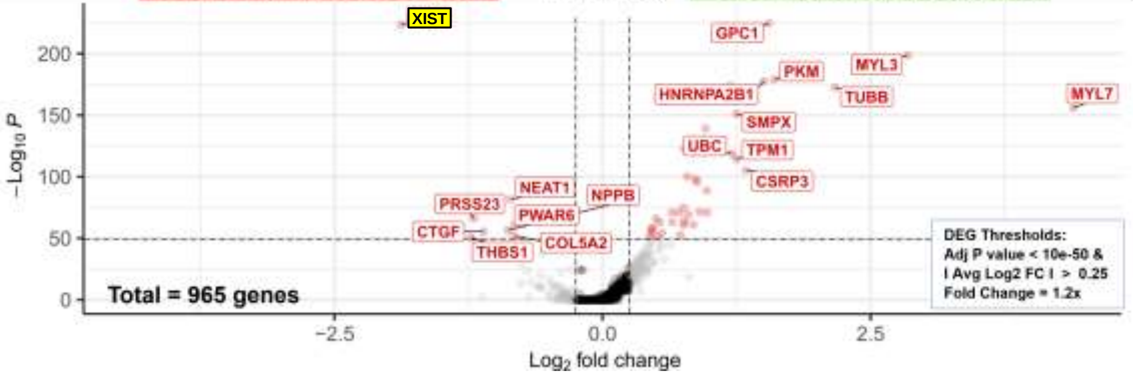


#### Over-Representation Analysis (ORA)

**Under-Expressed: 8 genes**

Enriched Gene Sets = 5

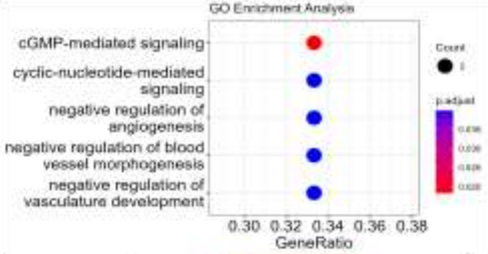


**Over- Expressed: 39 genes**

Enriched Gene Sets = 72

Top 5

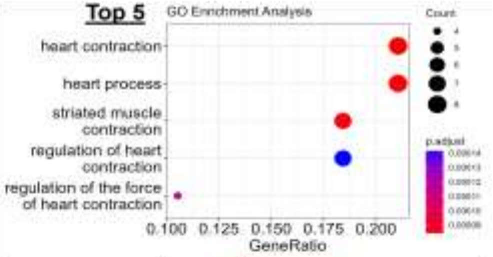


Enrichment Map
Gene Sets = 5

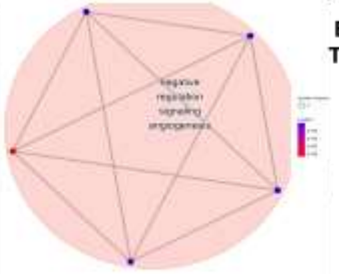

Genesx2
*NPPB*
*THBS1*

Enrichment Map
Top 25 Gene Sets

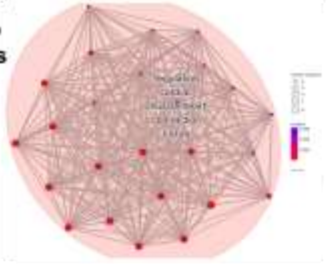

Genesx4-8
*e.g., MYL3*
*MYL7*
*TPM1*
*CSRP3*

#### Gene Set Enrichment Analysis (GSEA)

All 965 DEGs
Under-expressed: 395 genes
Over-expressed: 570 genes

**No Enriched MSigDB Hallmark Gene Sets**

#### Fig. S11 Cell Type DEG

**Integrated | D19 | EPDC | 2,258 cells**

INTEGRATED | Control | Patient

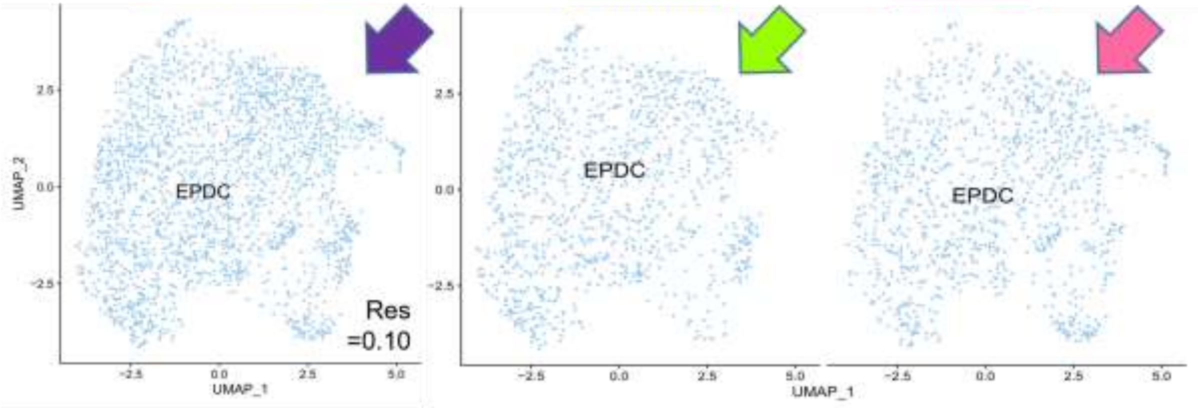


**EPDC | D19: 2,258 cells | Patient (50%) vs Control (50%)**

Under-Expressed: 5 genes | **22 DEGs** | Over-Expressed: 17 genes

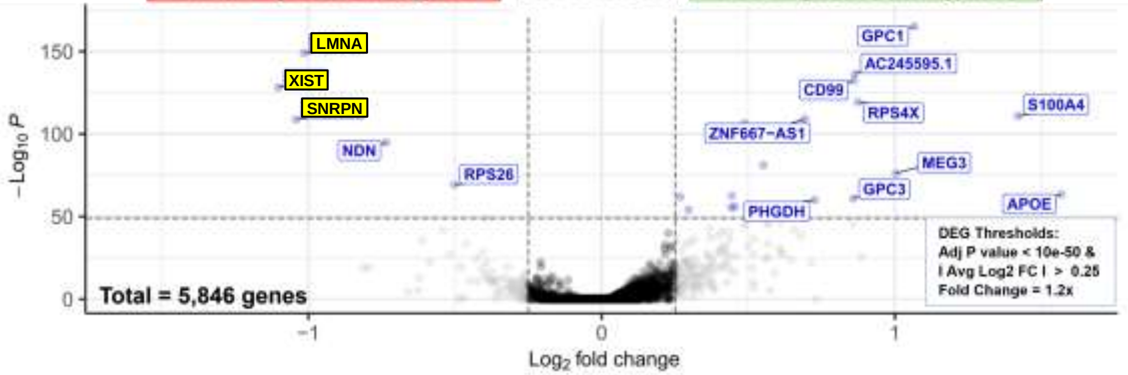


##### Over-Representation Analysis (ORA)

**Under-Expressed: 5 genes**

Enriched Gene Sets = 34

Top 5

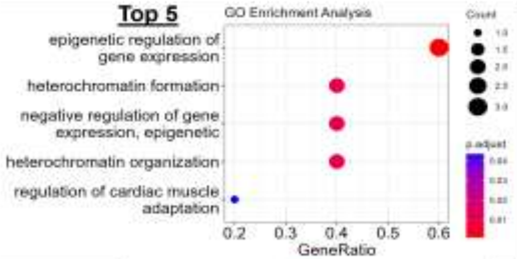


Enrichment Map
Top 25 Gene Sets

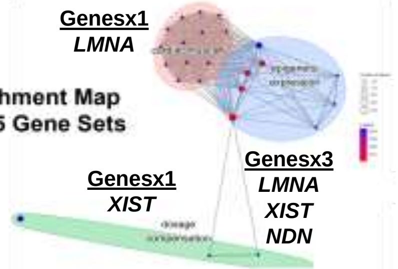


**Over- Expressed: 17 genes**

Enriched Gene Sets = 0

**No Enrichment:
GO Biological Process
Gene Sets**

##### Gene Set Enrichment Analysis (GSEA)

**All 5846 DEGs
Under-expressed: 3084 genes
Over-expressed: 2762 genes**

**No Enriched
MSigDB Hallmark
Gene Sets**

Fig. S11 Cell Type DEG

#### D. Cell Type DEG: *LMNA*, X-Linked Genes, and Imprinted Genes Across 14 Cell Subtypes

**Integrated** | **'Balanced' Paired Data Subsets (n= 6 Pairs)** | **75,330 cells**

***Lamin A/C (LMNA)***
**Underexpressed**
**Day 19 Patient Cells**

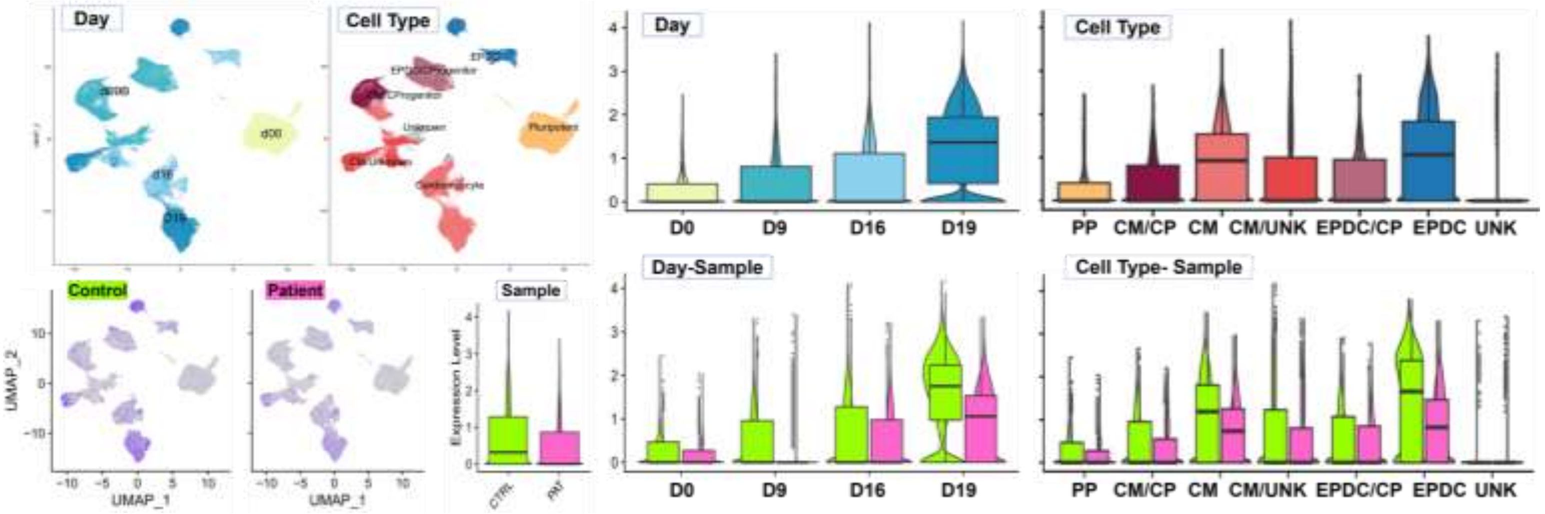


***Lamin B1***

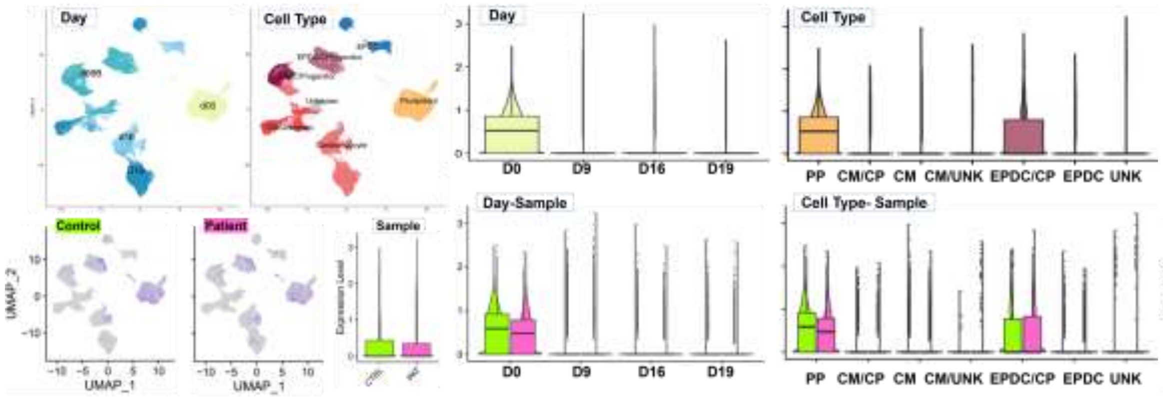


***Lamin B2***

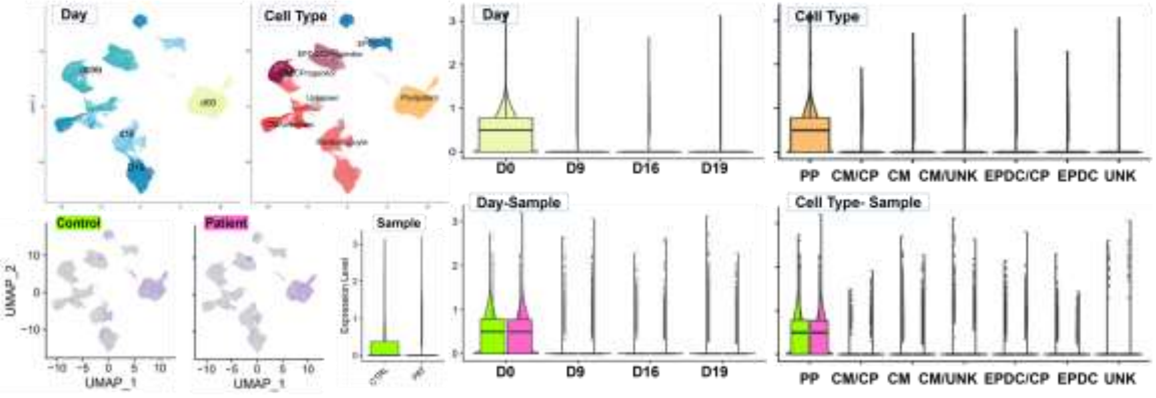

Fig. S11 Cell Type DEG

***X Inactive Specific Transcript (XIST)***
**Underexpressed**
**All Patient Cells**

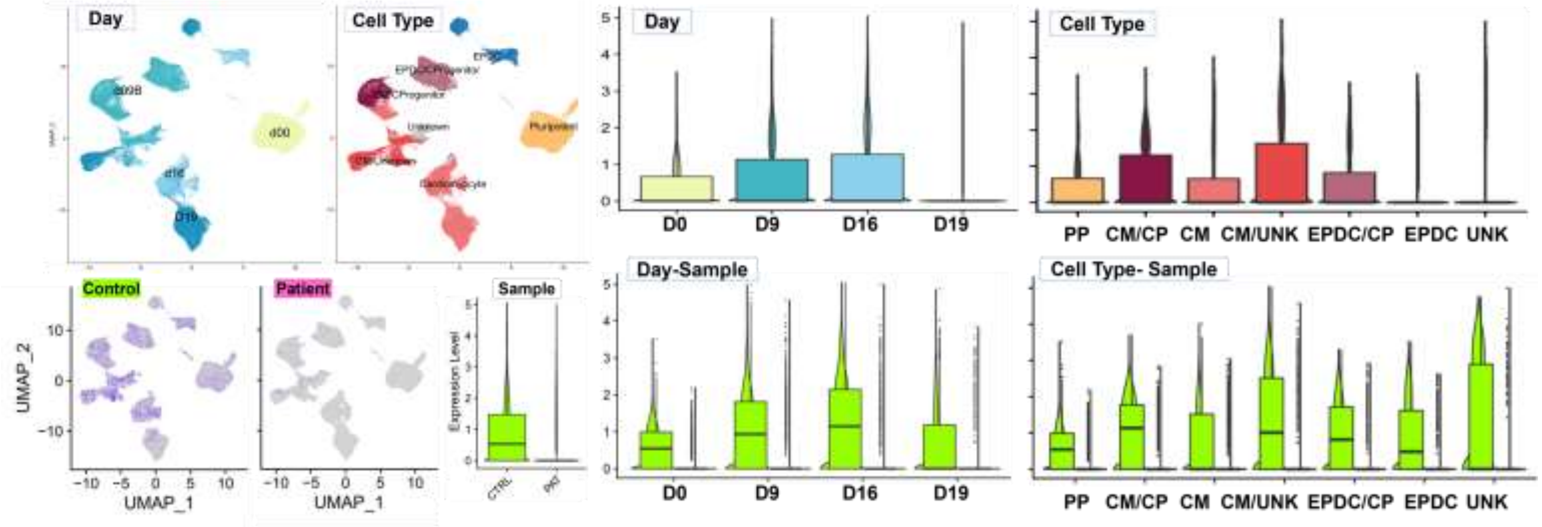


***X-Linked Glypican-3 (GPC3)***
**Overexpressed**
**Day 9 Patient Cells**

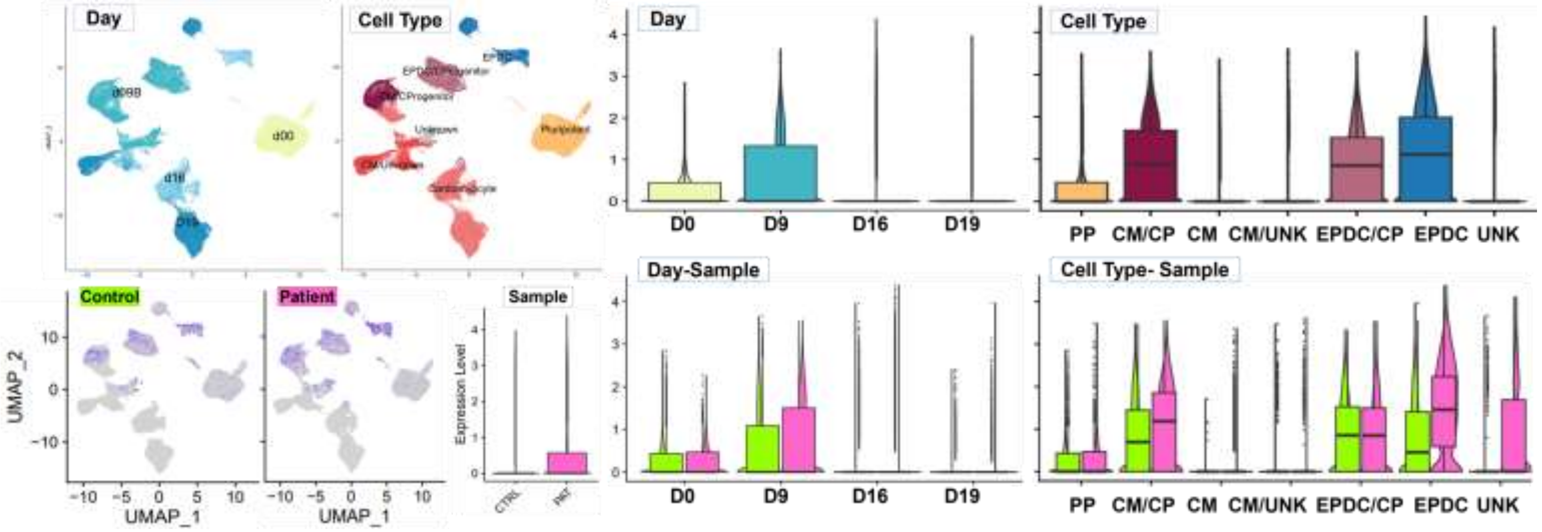

**Fig. S11 Cell Type DEG**

**Imprinted (PAT)**
***SNRPN***
**chr15q11.2**
**Underexpressed All Patient Cells**

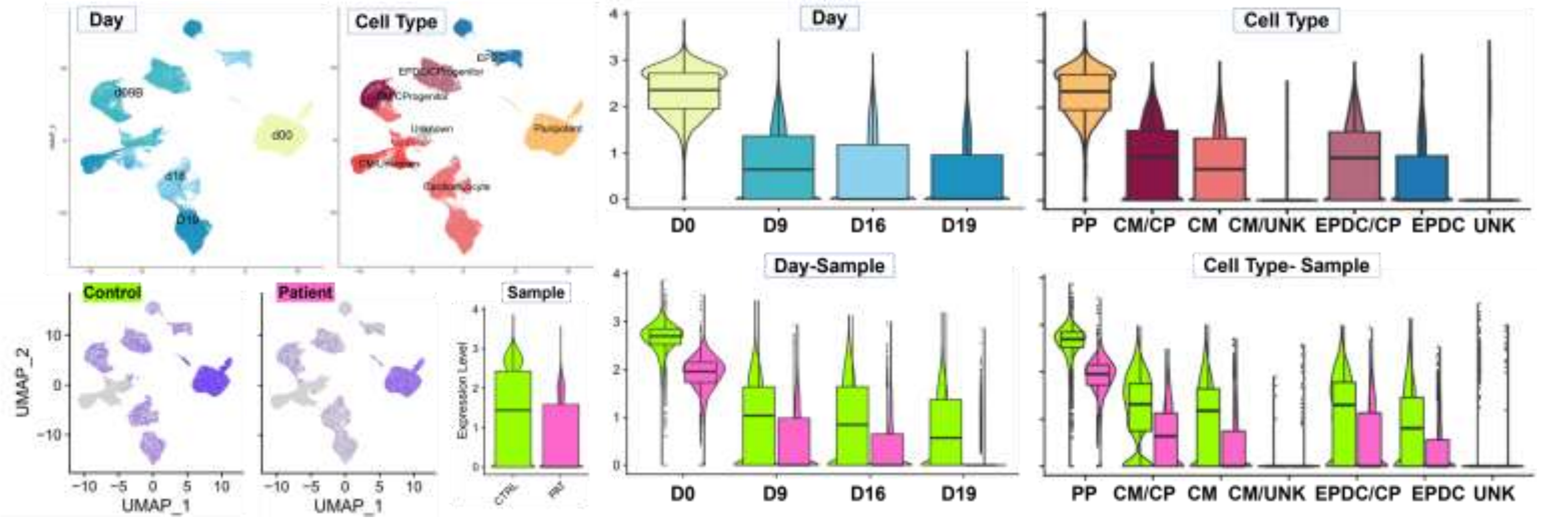


**Imprinted (MAT)**
***MEG3***
**chr14q32.2**
**Overexpressed Pluripotent (PP) & EPDC Patient Cells**

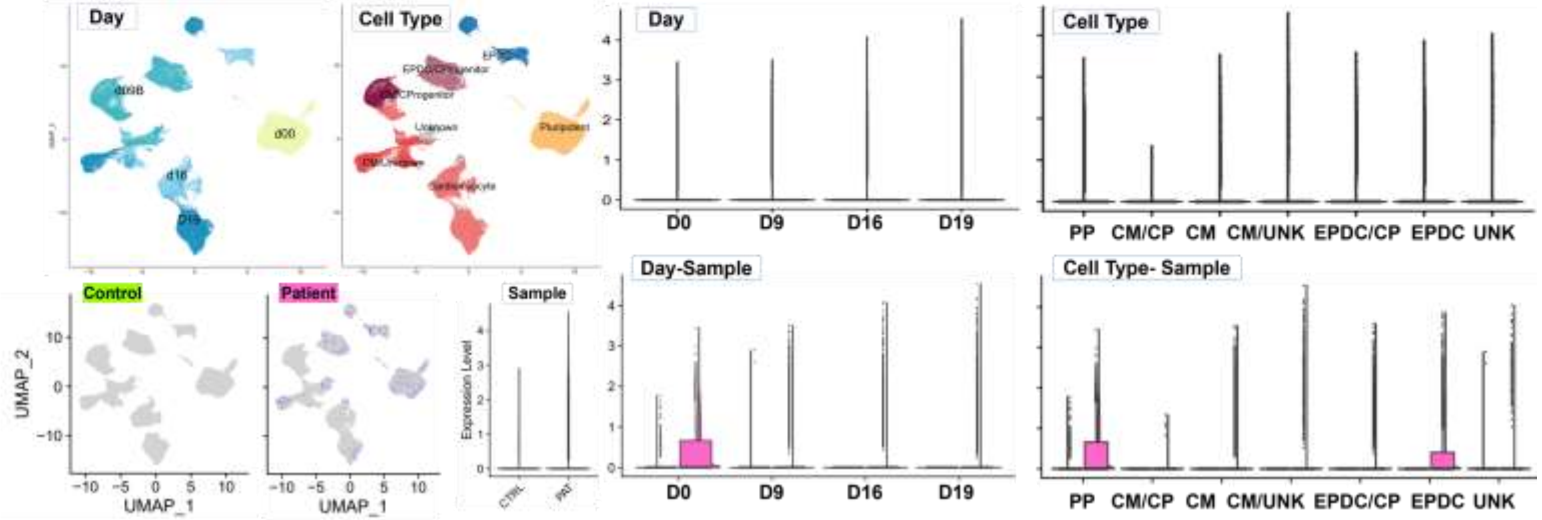

#### Fig. S11 Cell Type DEG

**X-linked *PIN4***
**Underexpressed Day 00 Patient Cells**

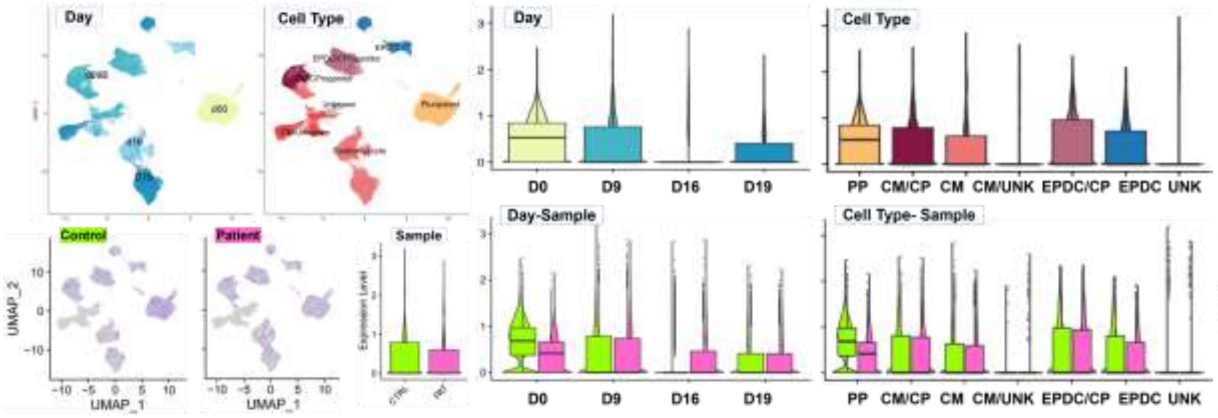


**Imprinted (PAT) *NDN* at chr15q11.2**
**Underexpressed in Patient Cells**

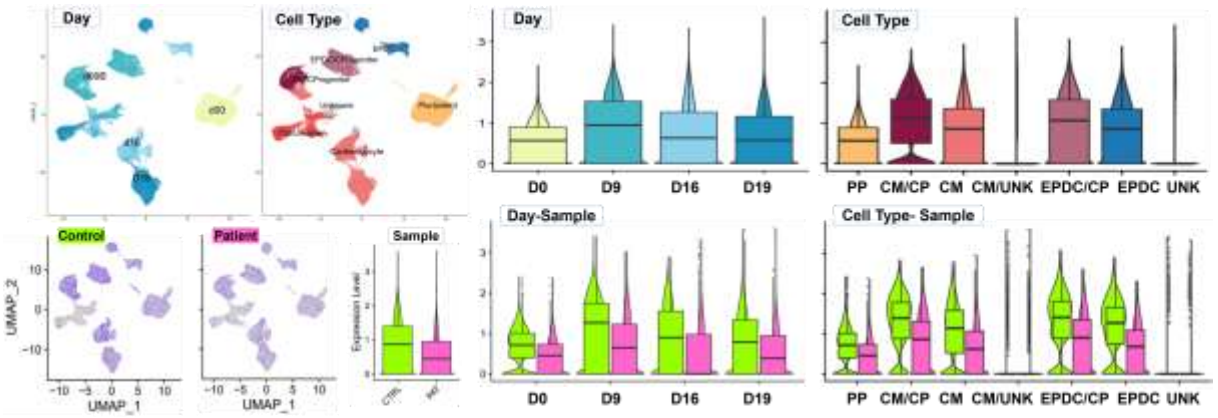


**Imprinted (PAT) *PWAR6* at chr15q11.2**
**Underexpressed in Patient Cells**

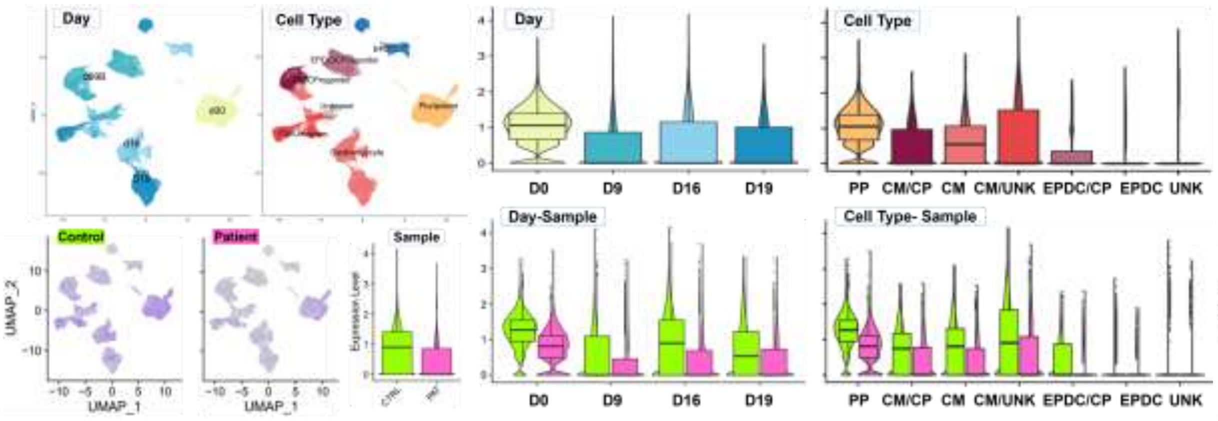


**Imprinted (PAT) *PEG10* at chr7q21.3**
**Underexpressed in Day 16 CM Patient Cells**

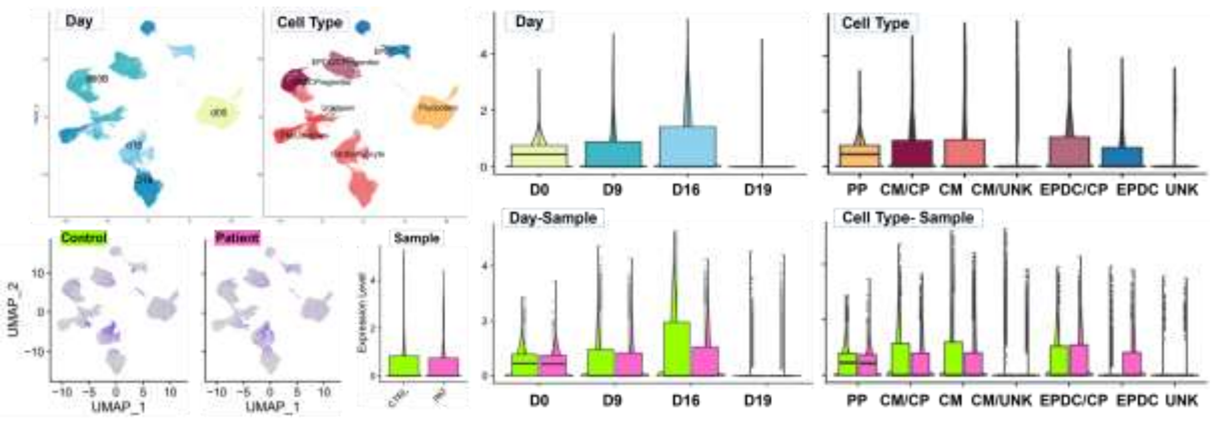

Fig. S11 Cell Type DEG

### E. Cell Type DEG Enrichment: Module Scoring of GSEA Significant Gene Sets

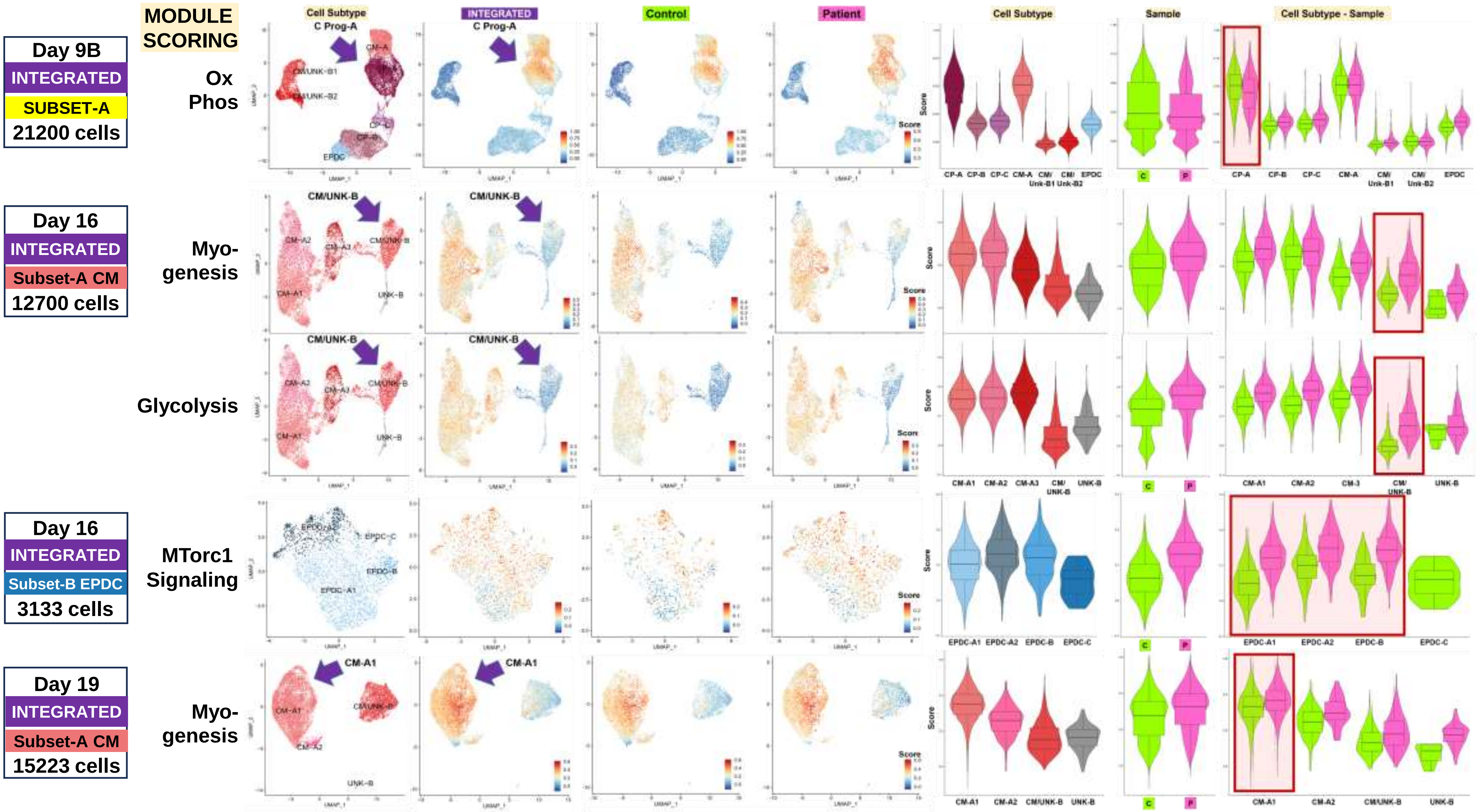

### Fig. S11 Cell Type DEG

| Day 16 |
| --- |
| INTEGRATED |
| CM Subset |
| 12700 cells |

MODULE SCORING

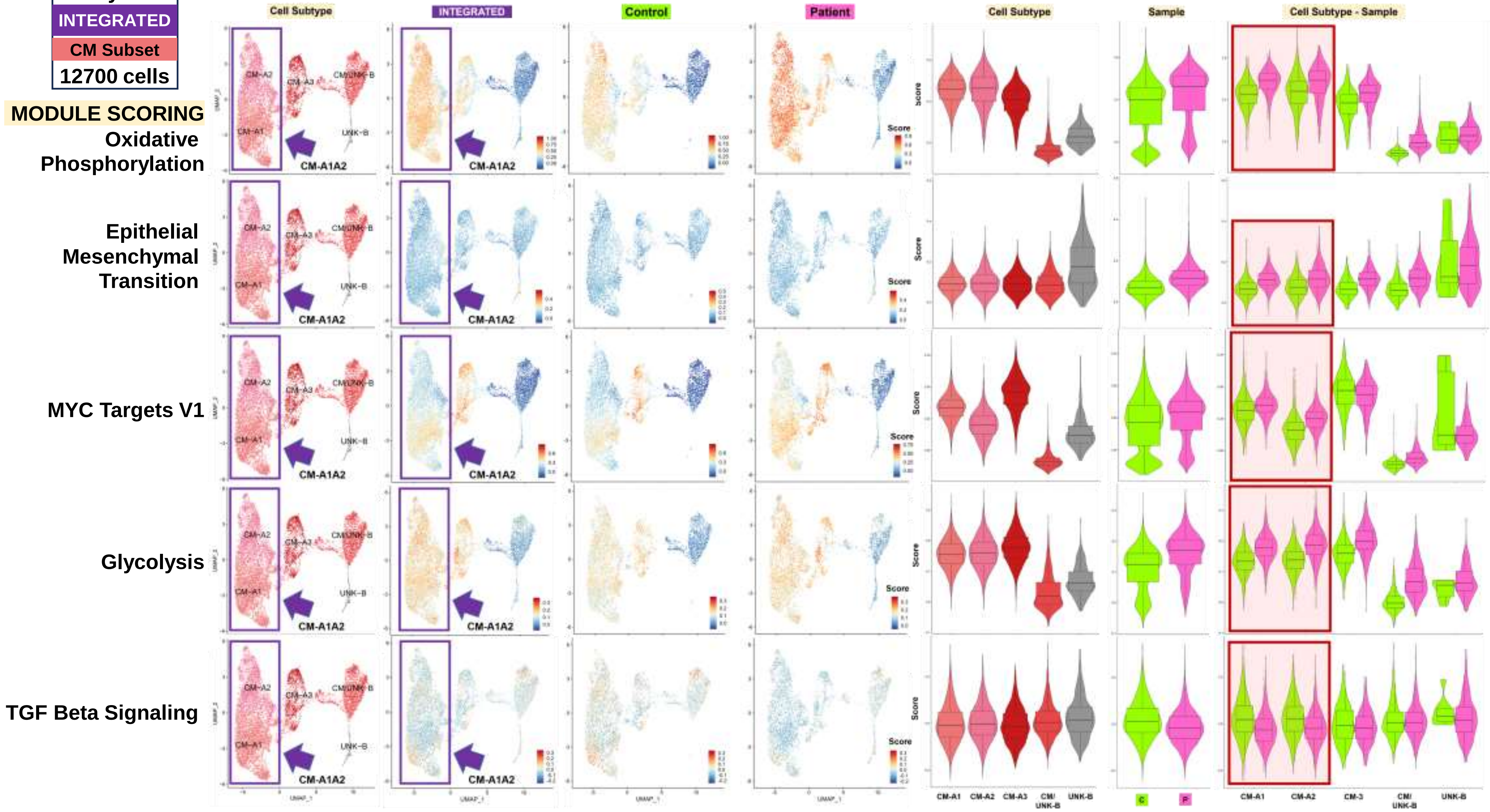

### Fig. S11 Cell Type DEG

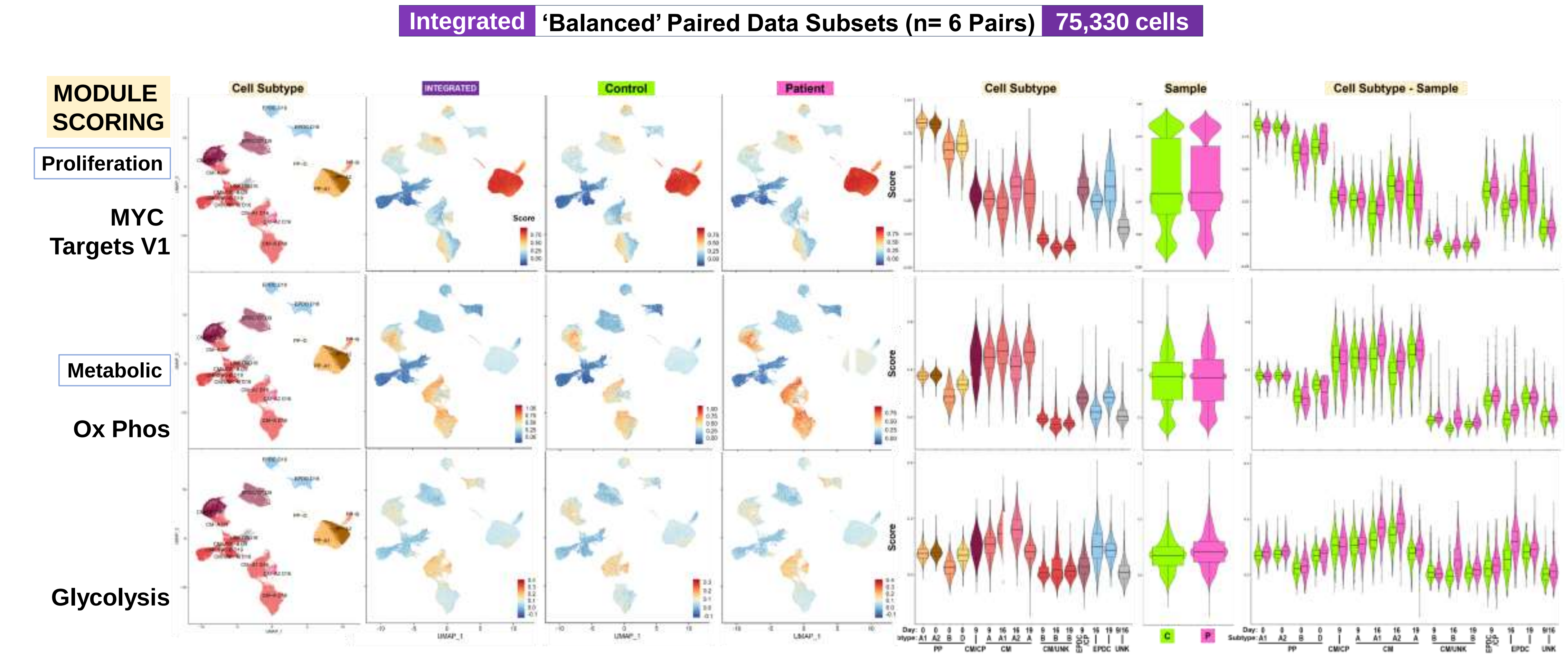

#### Fig. S11 Cell Type DEG

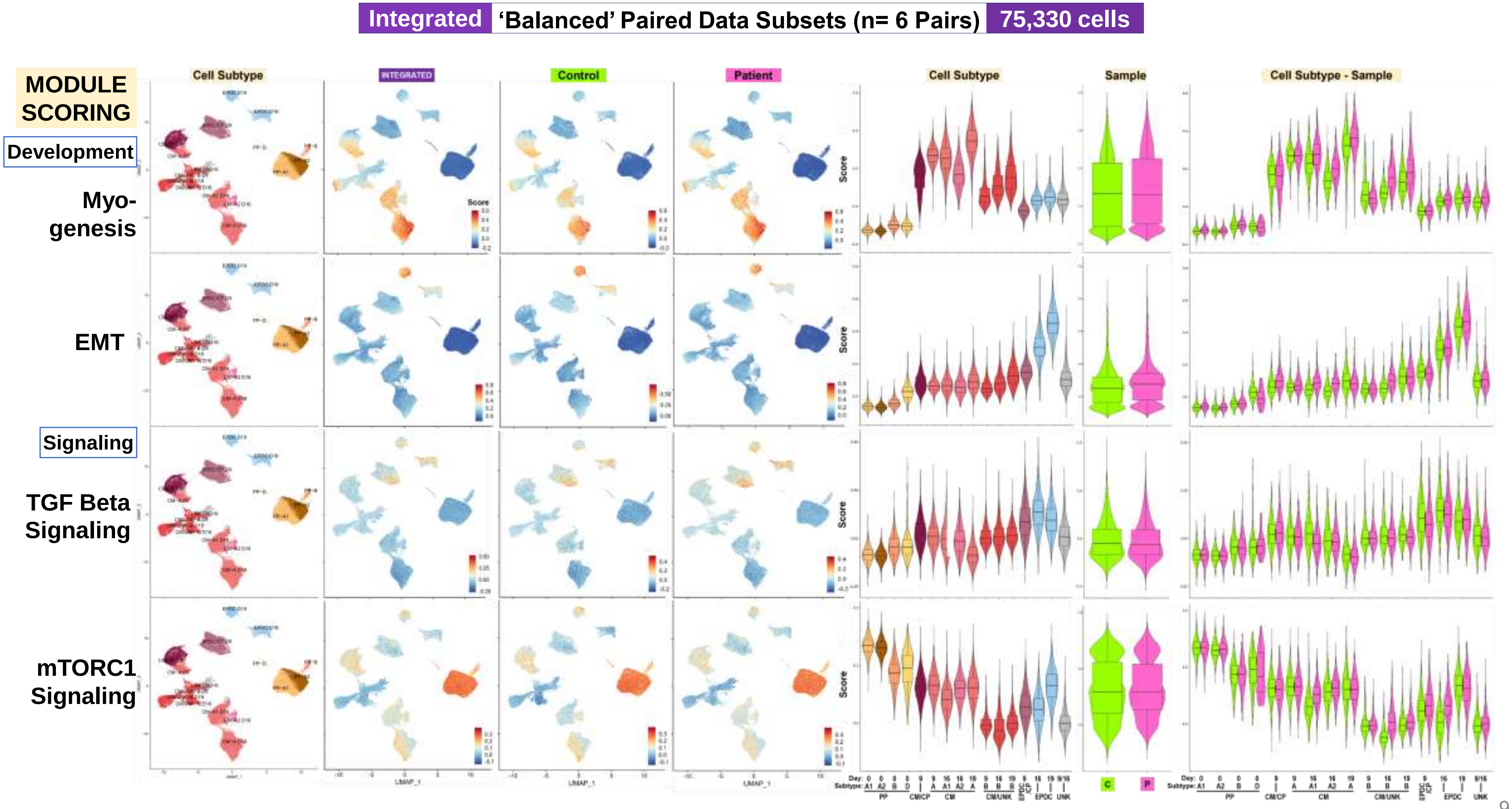

Fig. S12
Lineage DEG SS

### Workflow Step-III: Single Subset Data Results

#### Trajectory Analyses for Lineage-Specific Differential Expression and Enrichment

##### A. Summary: Annotated Subset Data to Cell Lineages

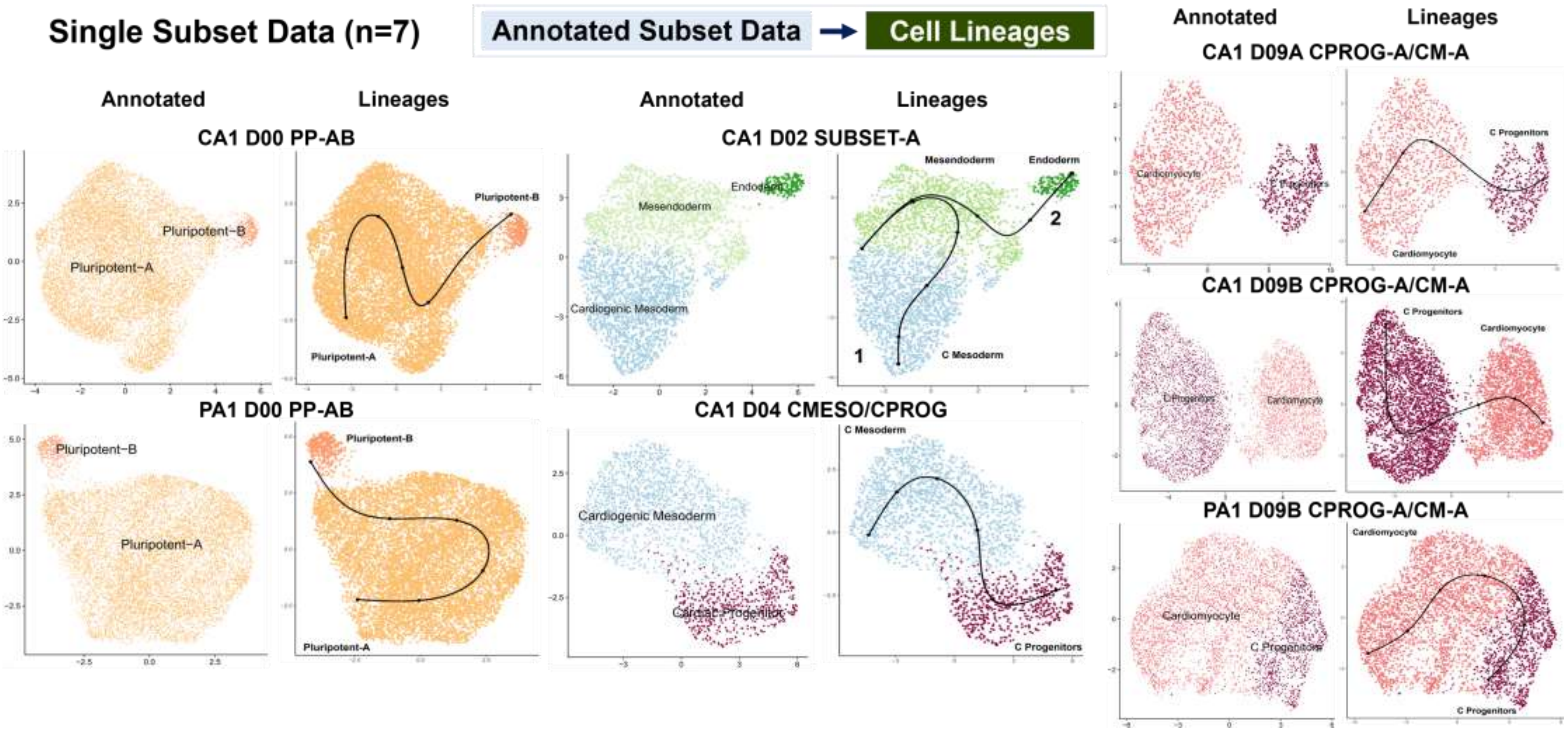

**Fig. S12**
**Lineage DEG SS**

#### B. Single Subset Data: Trajectory, Lineage DEG, and Enrichment Analyses (n=7)

| Control | D00 | PP-AB | 9,988 cells | Lineage DE Analyses & Enrichment |
|---|---|---|---|---|

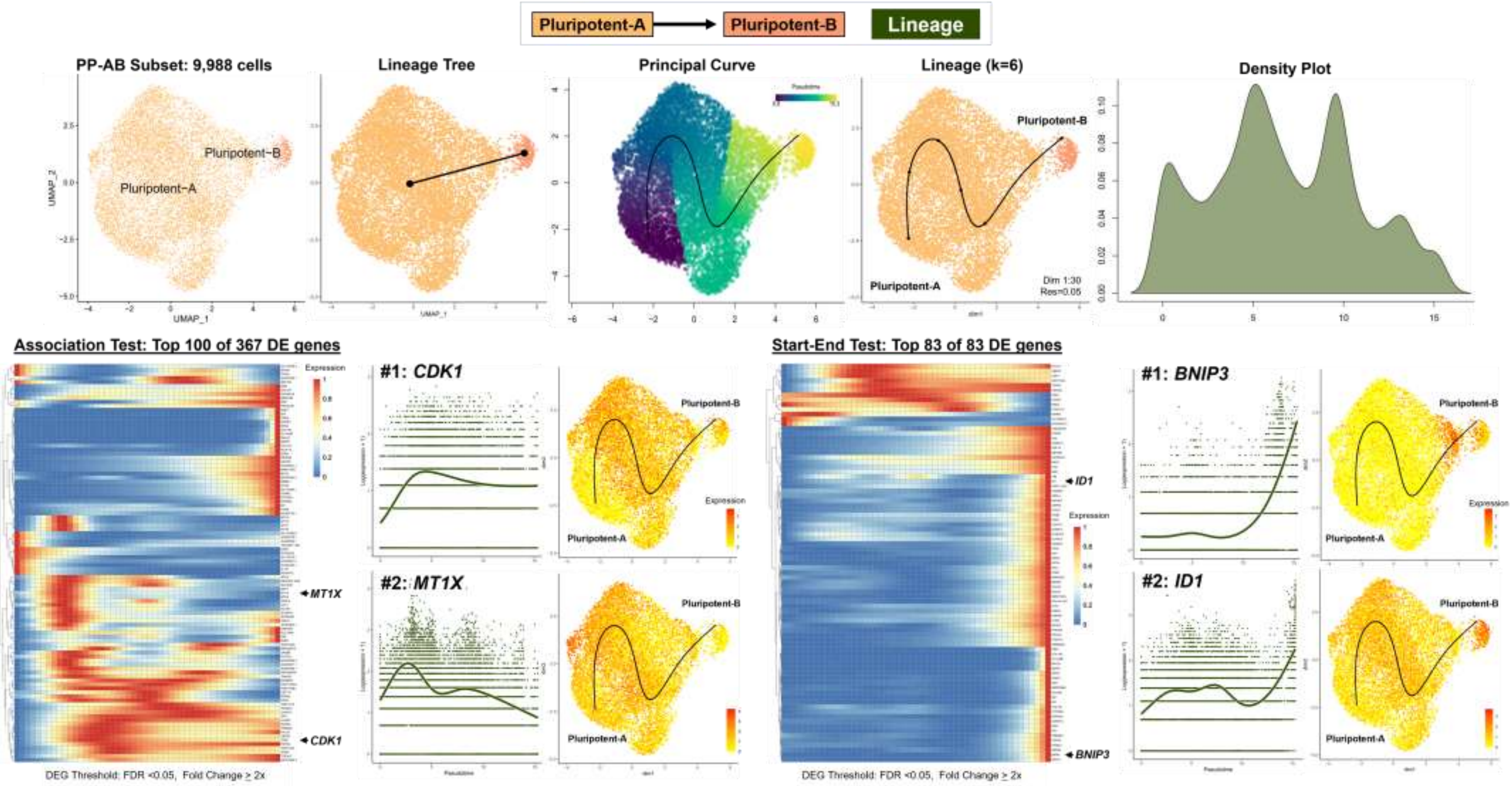

**Fig. S12**
**Lineage DEG SS**

| Control | D00 | PP-AB | 9,988 cells | Lineage DE Analyses & Enrichment |
| --- | --- | --- | --- | --- |

#### Over-Representation Analysis

Pluripotent-A → Pluripotent-B | Lineage

| Enrichment Plots: Top 5 Gene Sets | Enrichment Maps Top 25 Gene Sets | Enrichment Trees Top 25 Gene Sets |
| --- | --- | --- |

**Top 100 of 367 Association DEG**

**Enriched Gene Sets = 36**

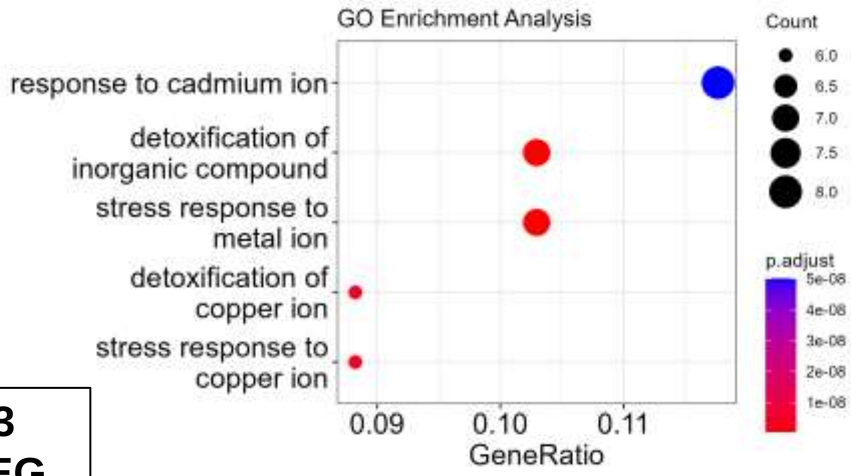


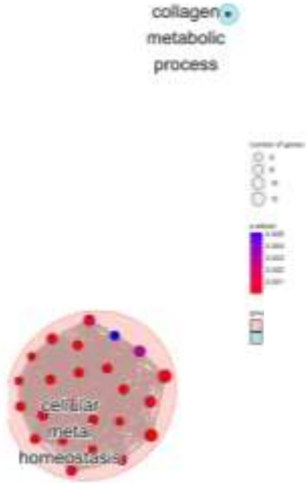


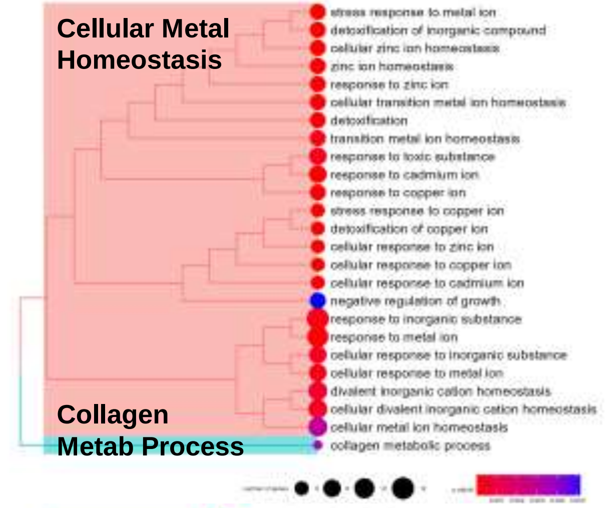


**Top 83 of 83 Start-End DEG**

**Enriched Gene Sets = 45**

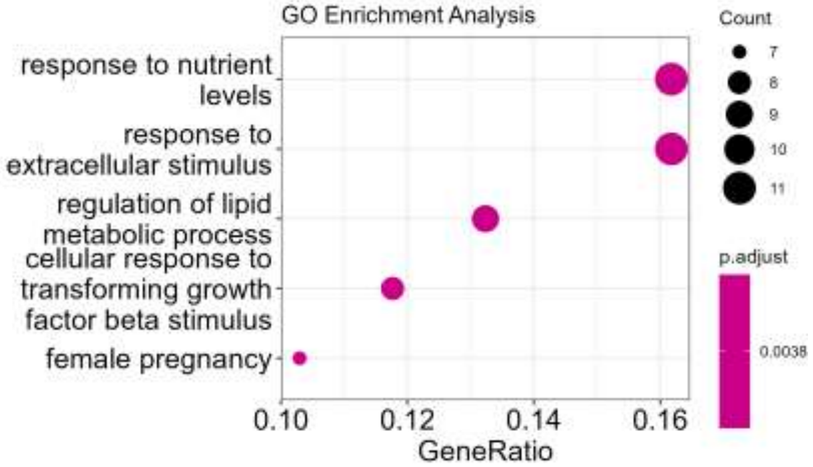


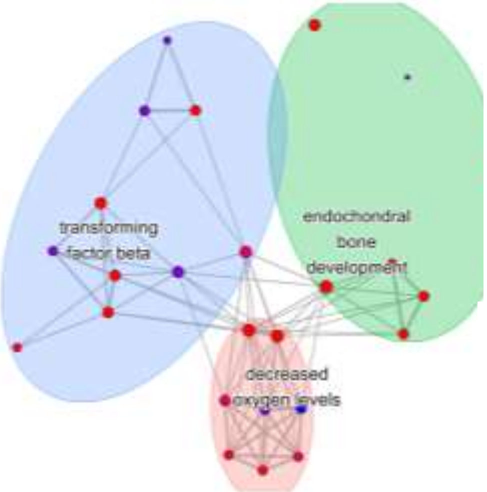


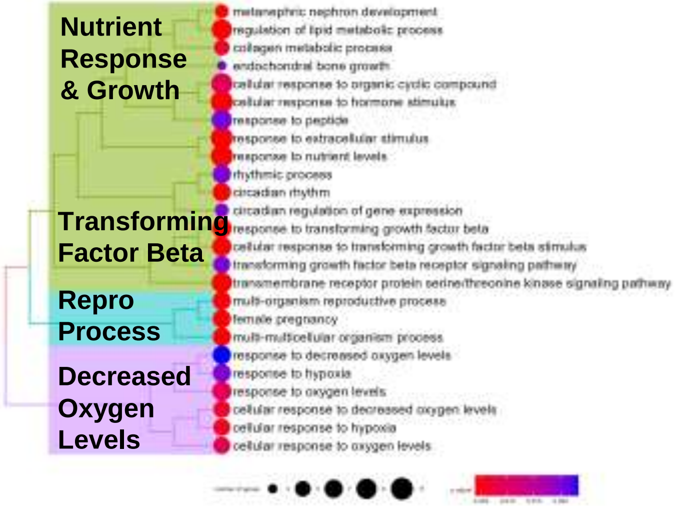

**Fig. S12**
**Lineage DEG SS**

| Patient | D00 | PP-AB | 9,358 cells | Lineage DE Analyses & Enrichment |
|---|---|---|---|---|

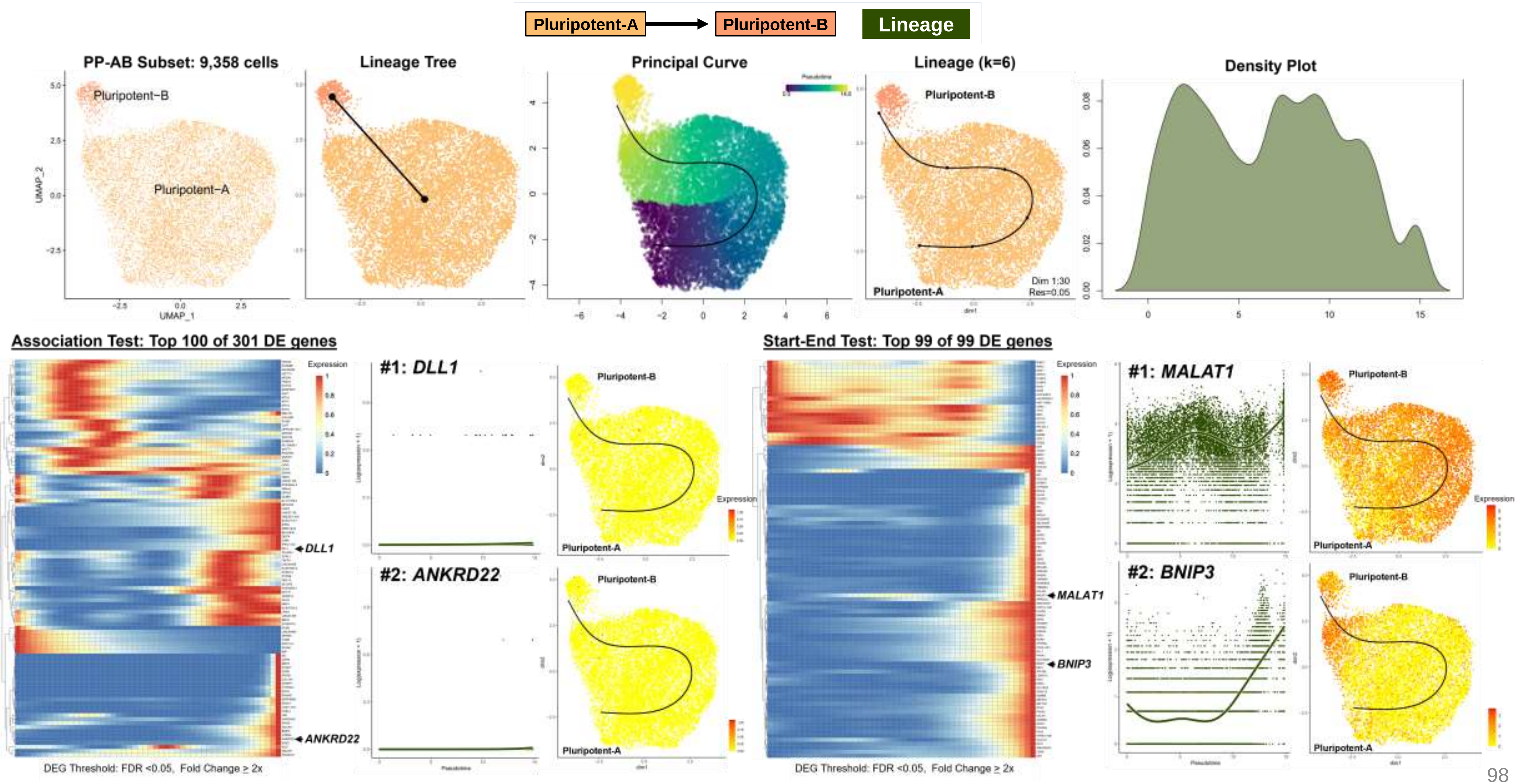

**Fig. S12**
**Lineage DEG SS**

| Patient | D00 | PP-AB | 9,358 cells | Lineage DE Analyses & Enrichment |
|---|---|---|---|---|

#### Over-Representation Analysis

Pluripotent-A → Pluripotent-B | Lineage

**Enrichment Plots: Top 5 Gene Sets** | **Enrichment Maps Top 25 Gene Sets** | **Enrichment Trees Top 25 Gene Sets**

**Top 100 of 301 Association DEG**

**Enriched Gene Sets = 174**

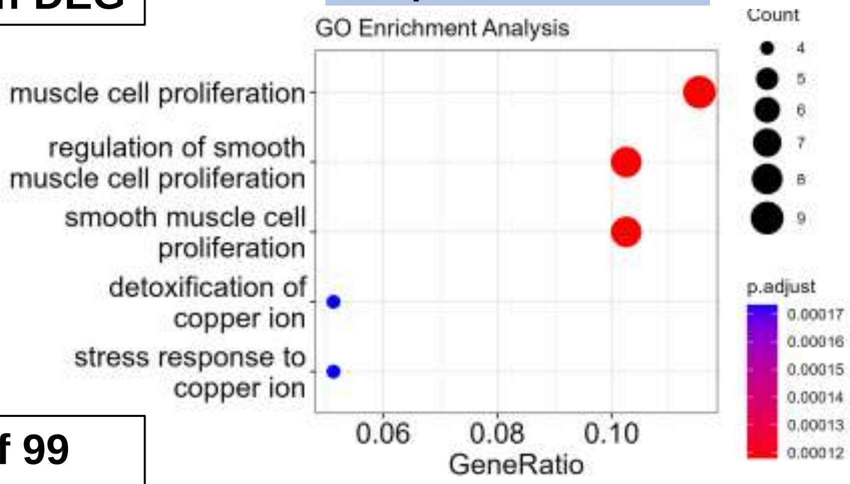


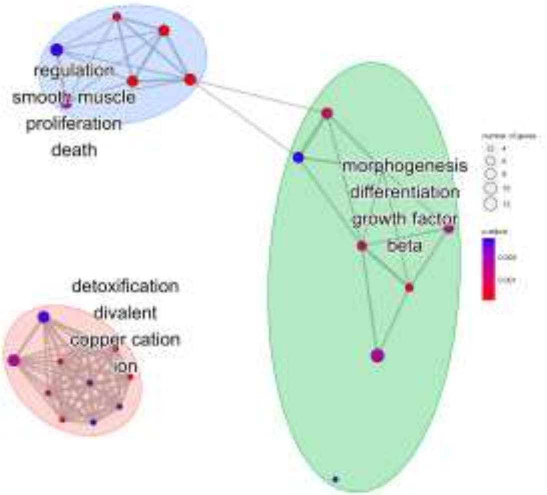


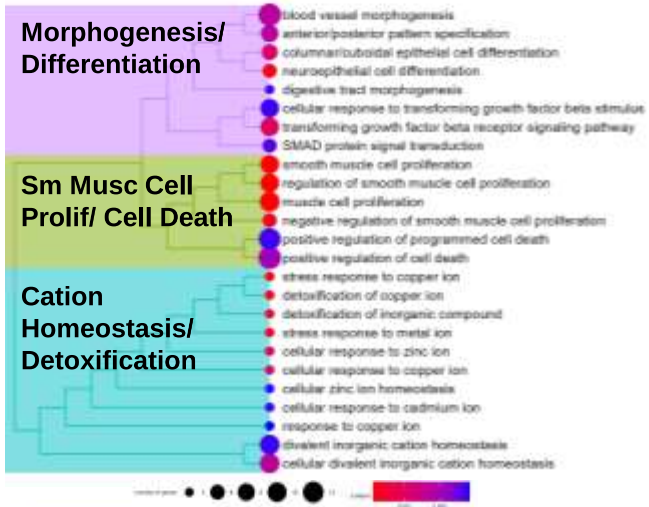


**Top 99 of 99 Start-End DEG**

**Enriched Gene Sets = 246**

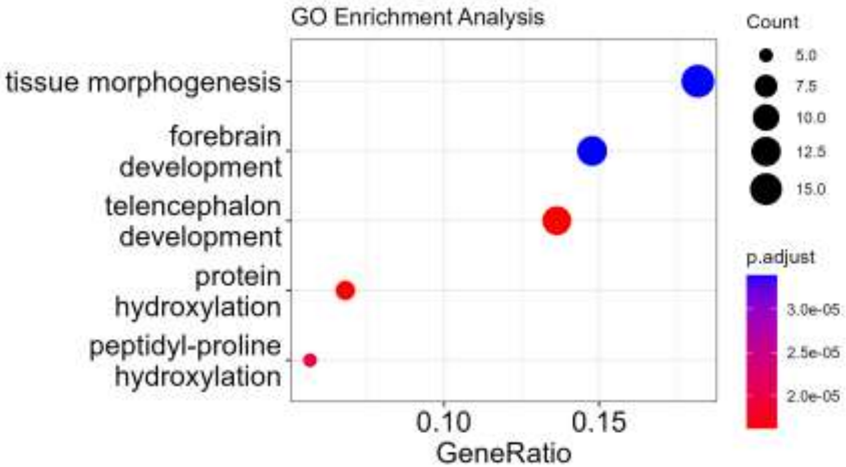


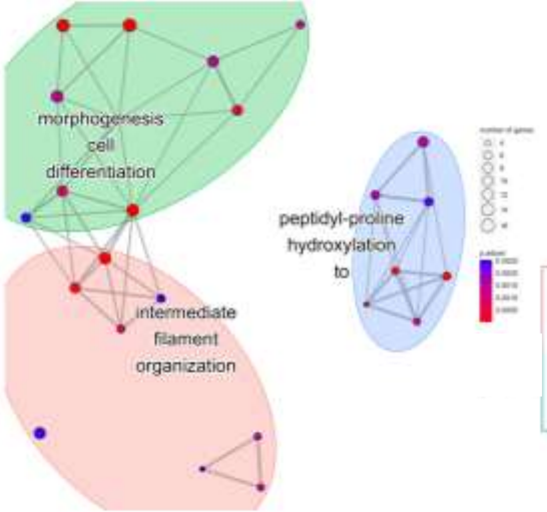


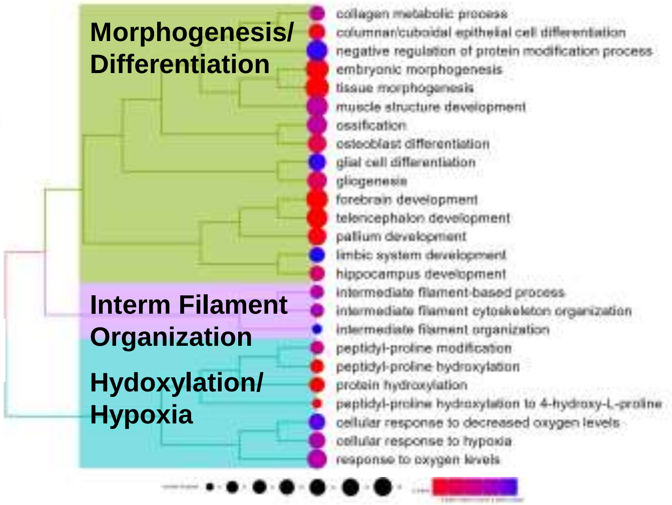

**Fig. S12**
**Lineage DEG SS**

| Control | D02 | ME/CMESO/ENDO | 3,145 cells | Lineage DE Analyses & Enrichment |
|---|---|---|---|---|

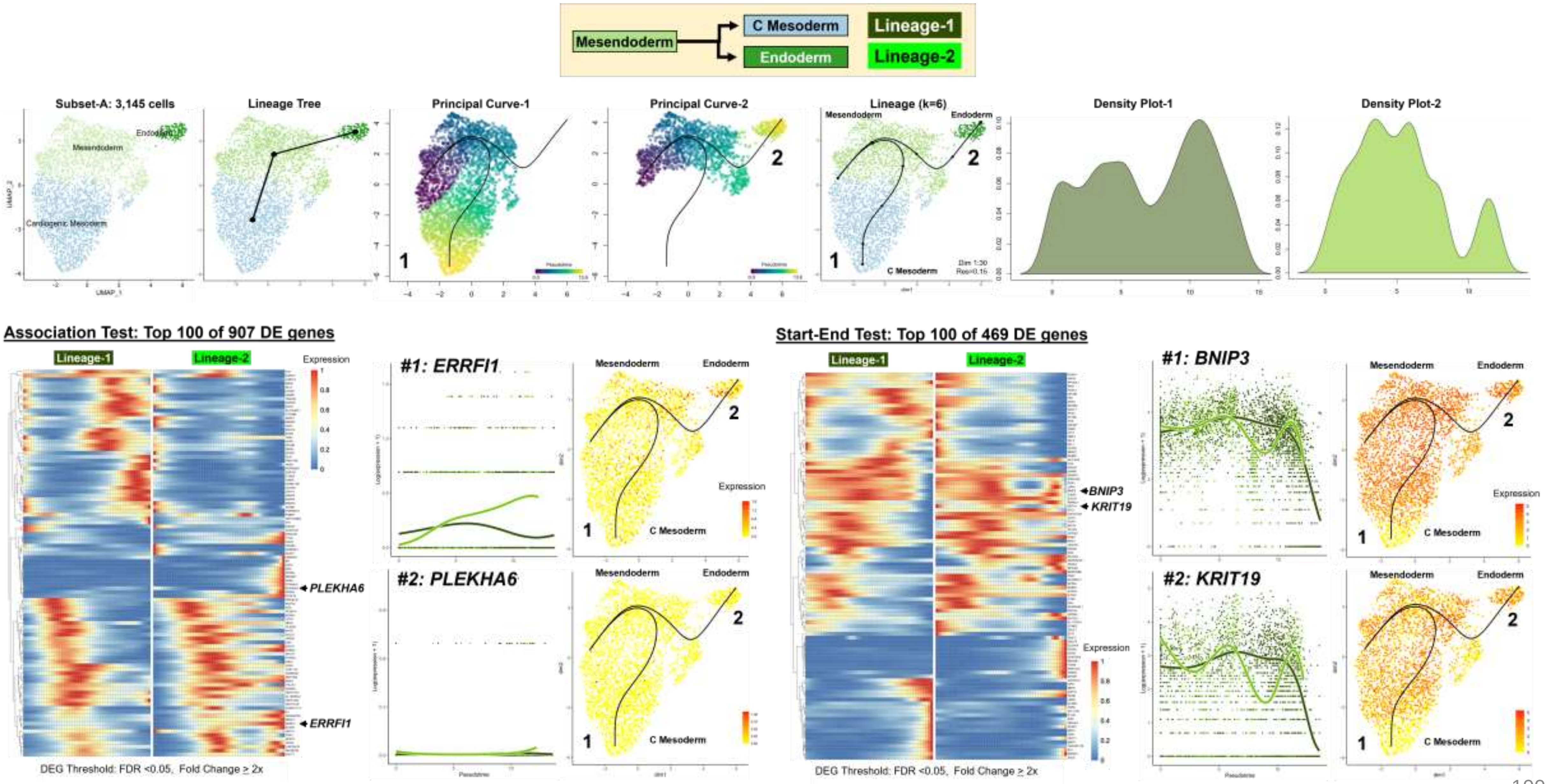

Fig. S12
Lineage DEG SS

| Control | D02 | ME/CMESO/ENDO | 3,145 cells | Lineage DE Analyses & Enrichment |
|---|---|---|---|---|

#### Over-Representation Analysis

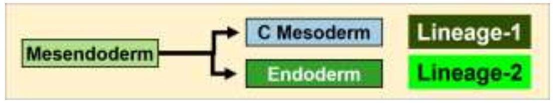


**Enrichment Plots: Top 5 Gene Sets**

**Enrichment Maps Top 25 Gene Sets**

**Enrichment Trees Top 25 Gene Sets**

**Top 100 of 907 Association DEG**

Enriched Gene Sets = 27

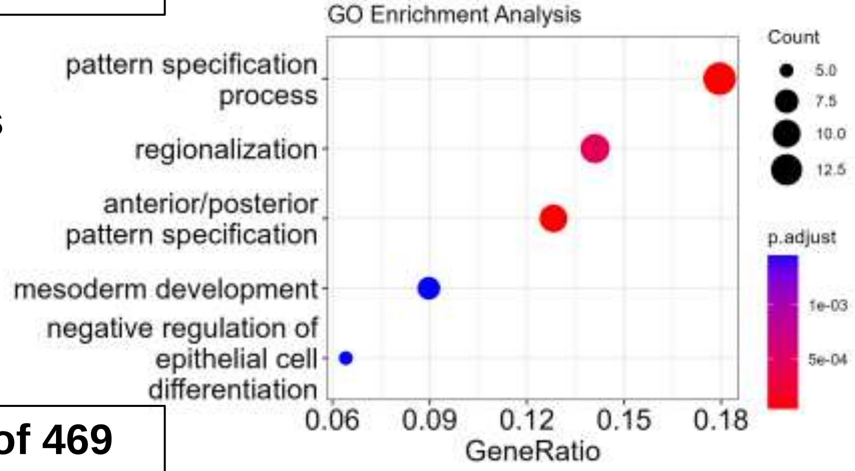


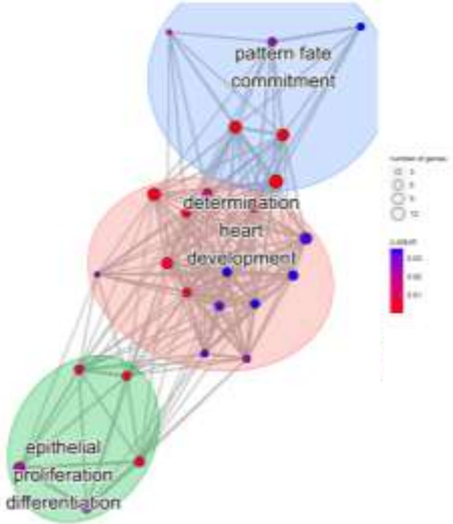


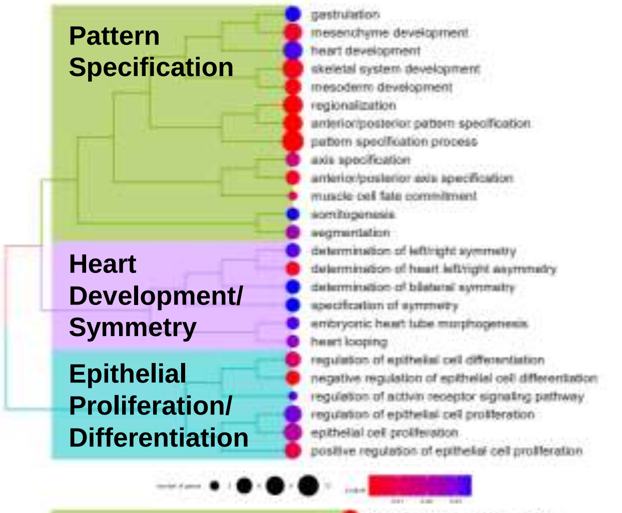


**Top 100 of 469 Start-End DEG**

Enriched Gene Sets = 187

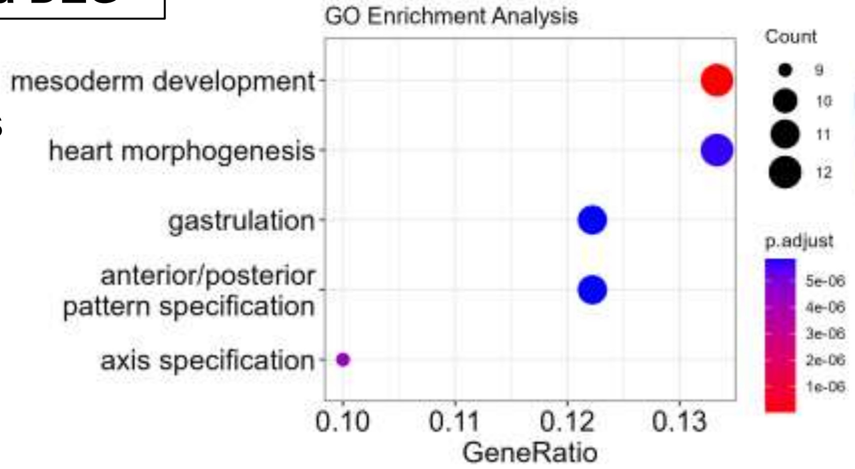


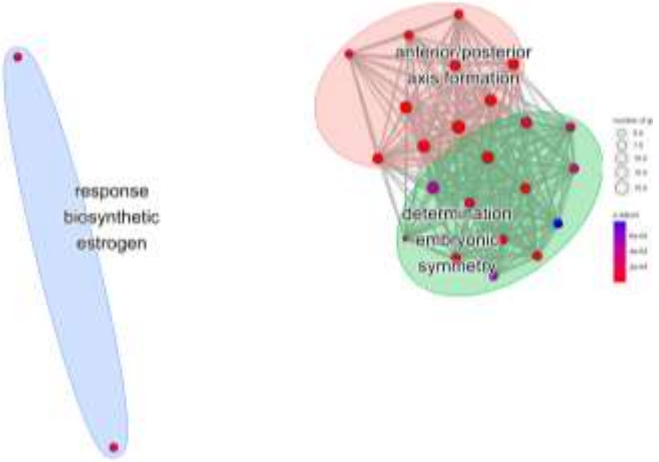


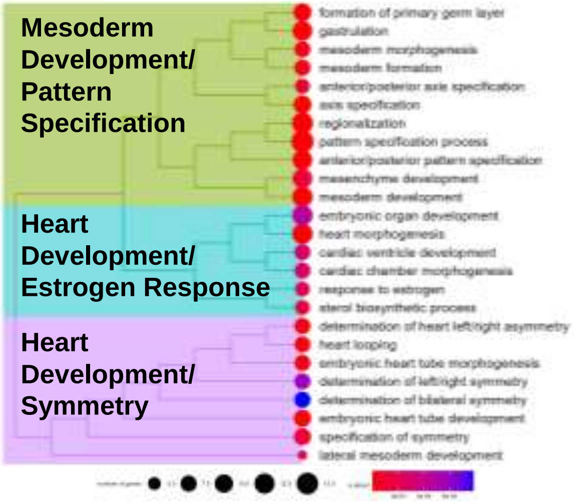

Fig. S12
Lineage DEG SS

| Control | D04 | CMESO/CPROG | 2,229 cells | Lineage DE Analyses & Enrichment |
| --- | --- | --- | --- | --- |

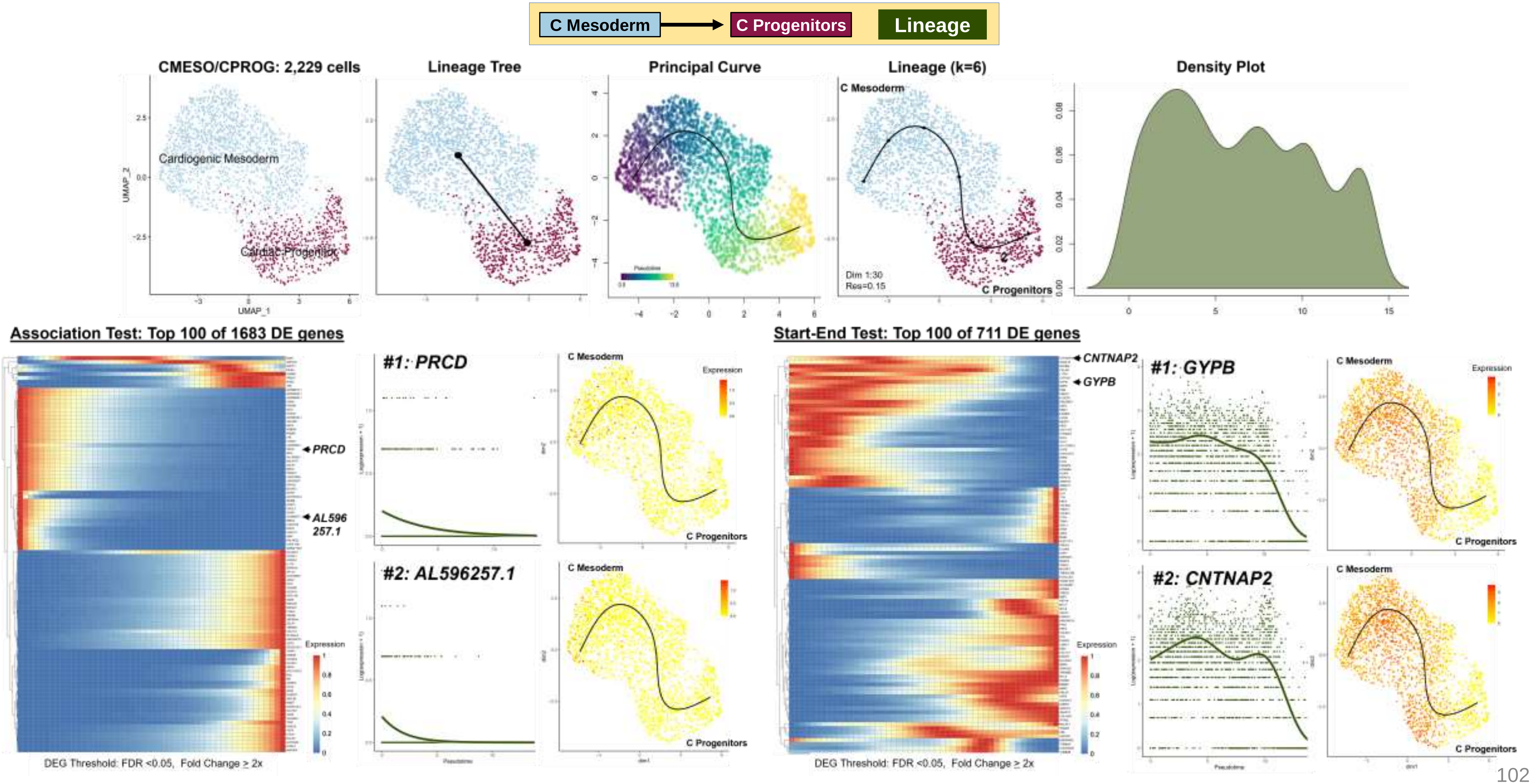

**Fig. S12**
**Lineage DEG SS**

| Control | D04 | CMESO/CPROG | 2,229 cells | Lineage DE Analyses & Enrichment |
|---|---|---|---|---|

**Over-Representation Analysis**

C Mesoderm → C Progenitors | Lineage

| Enrichment Plots: Top 5 Gene Sets | Enrichment Maps Top 25 Gene Sets | Enrichment Trees Top 25 Gene Sets |
|---|---|---|

**Top 100 of 1683 Association DEG**

**Enriched Gene Sets = 0**

| No Enrichment: GO Biological Process Gene Sets | No Enrichment: GO Biological Process Gene Sets | No Enrichment: GO Biological Process Gene Sets |
|---|---|---|

**Top 100 of 711 Start-End DEG**

**Enriched Gene Sets = 114**

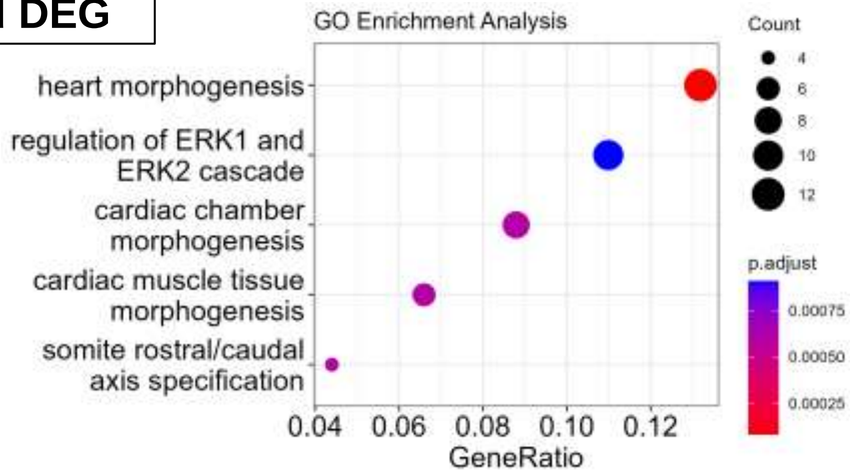


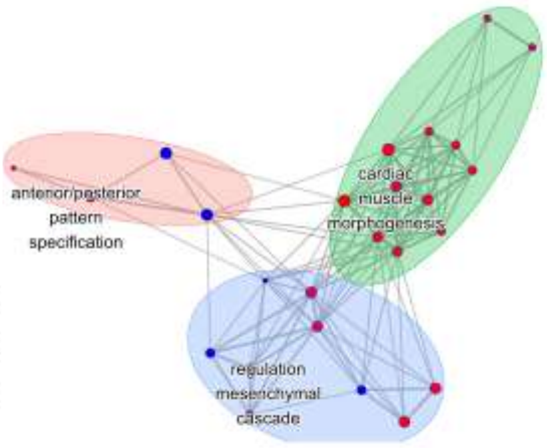


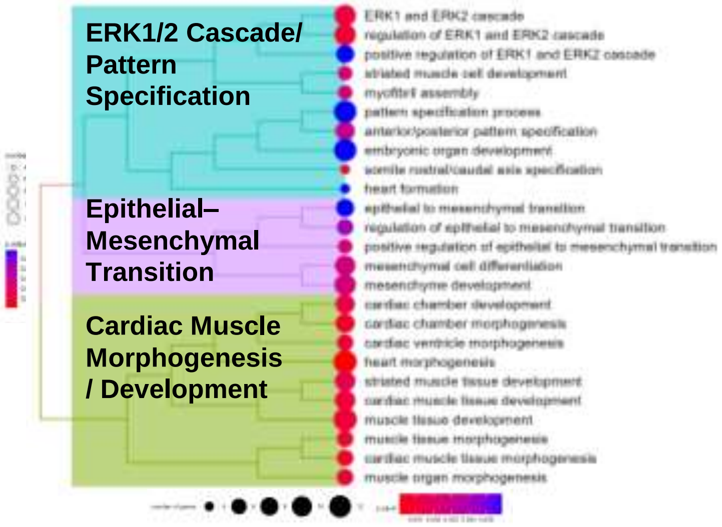

Fig. S12
Lineage DEG SS

| Control | D09A | CPROG-A/CM-A | 1,671 cells | Lineage DE Analyses & Enrichment |
|---|---|---|---|---|

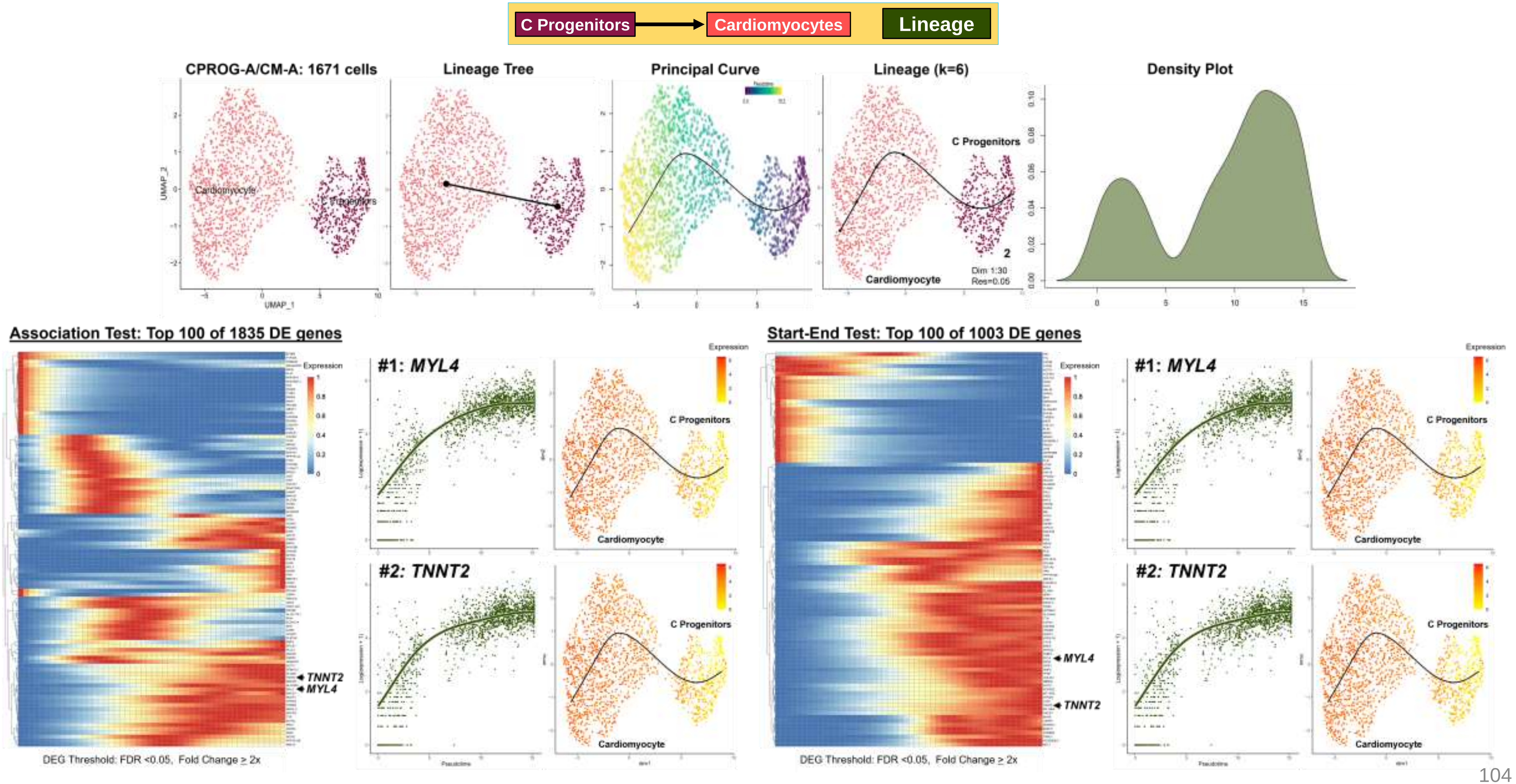

**Fig. S12**
**Lineage DEG SS**

| Control | D09A | CPROG-A/CM-A | 1,671 cells | Lineage DE Analyses & Enrichment |
|---|---|---|---|---|

#### Over-Representation Analysis

C Progenitors → Cardiomyocytes — Lineage

**Enrichment Plots: Top 5 Gene Sets**

**Enrichment Maps Top 25 Gene Sets**

**Enrichment Trees Top 25 Gene Sets**

**Top 100 of 1835 Association DEG**

**Enriched Gene Sets = 139**

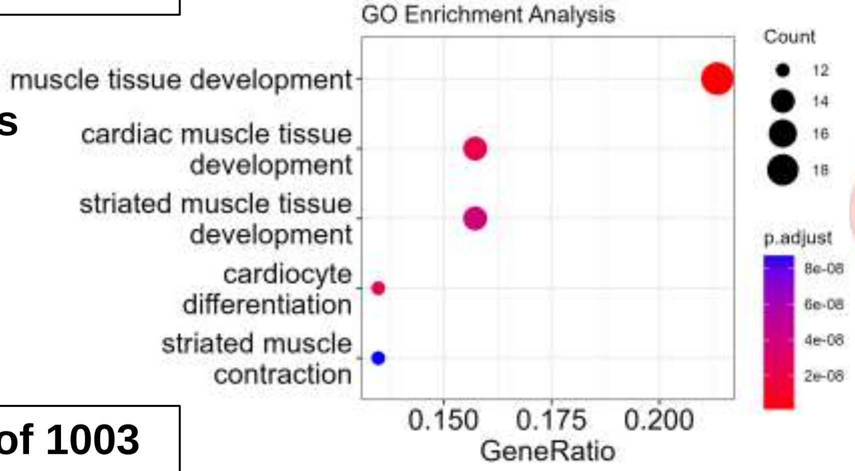


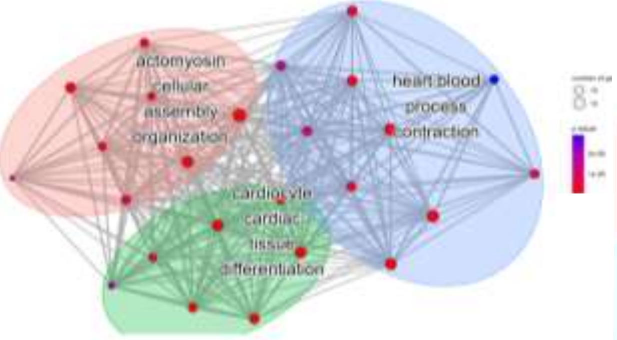


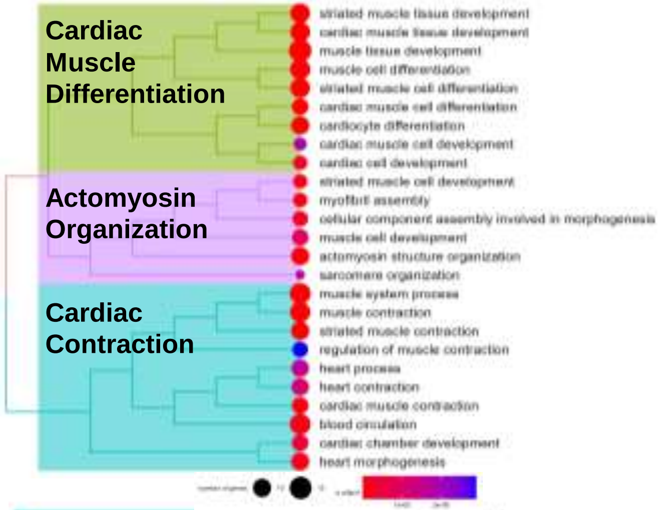


**Top 100 of 1003 Start-End DEG**

**Enriched Gene Sets = 239**

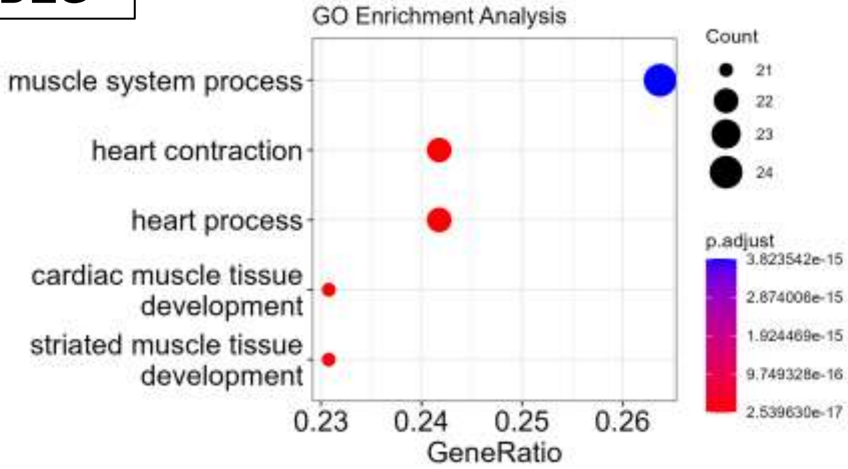


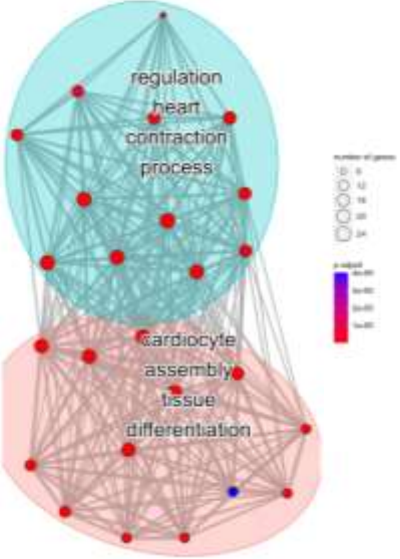


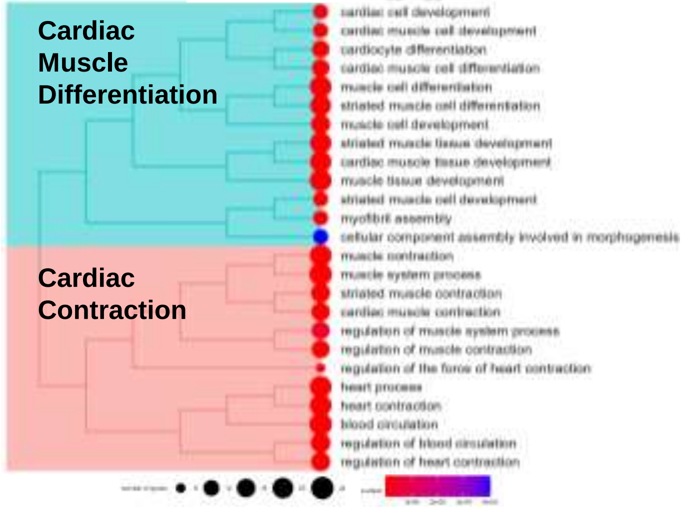

**Fig. S12**
**Lineage DEG SS**

| Control | D09B | CPROG-A/CM-A | 5,293 cells | Lineage DE Analyses & Enrichment |
|---|---|---|---|---|

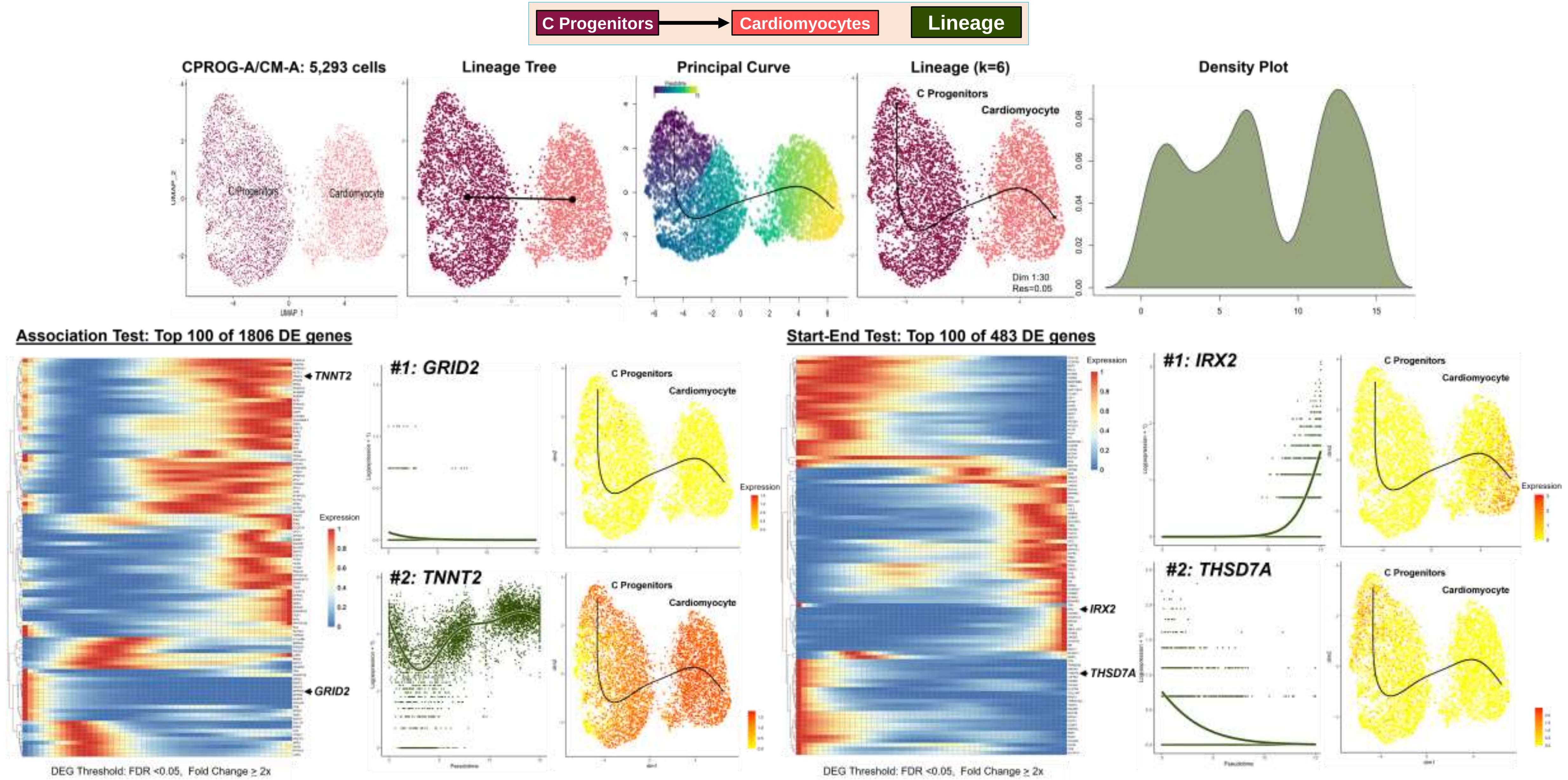

**Fig. S12**
**Lineage DEG SS**

| Control | D09B | CPROG-A/CM-A | 5,293 cells | Lineage DE Analyses & Enrichment |
|---|---|---|---|---|

#### Over-Representation Analysis

C Progenitors → Cardiomyocytes Lineage

**Top 100 of 1806 Association DEG**

**Enriched Gene Sets = 153**

**Enrichment Plots: Top 5 Gene Sets**

**Enrichment Maps Top 25 Gene Sets**

**Enrichment Trees Top 25 Gene Sets**

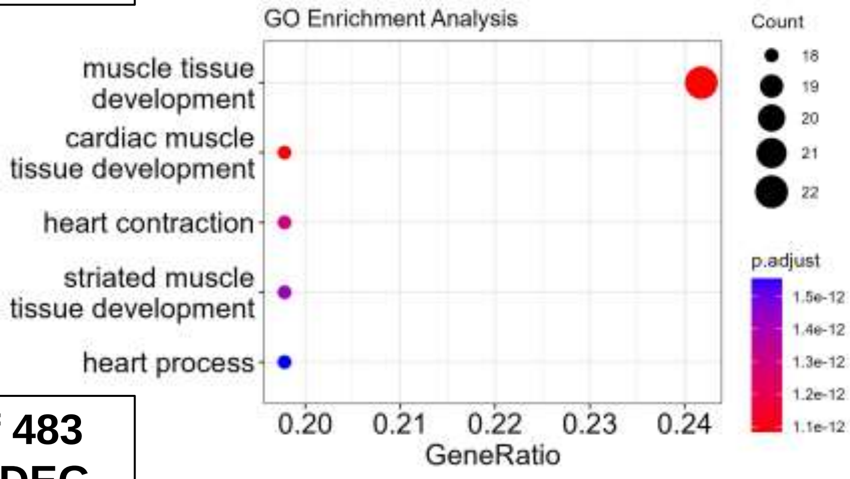


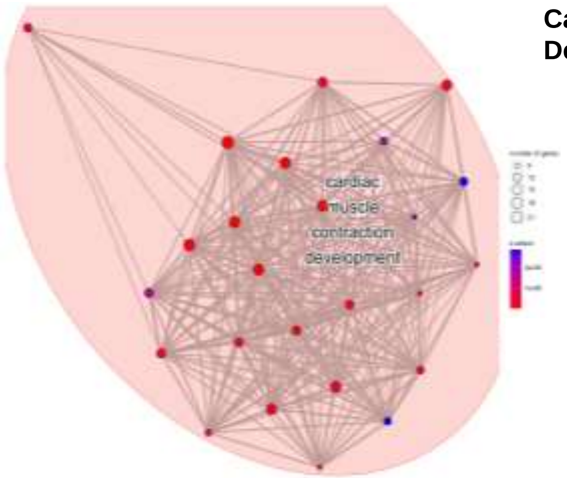


**Cardiac Function/ Development**

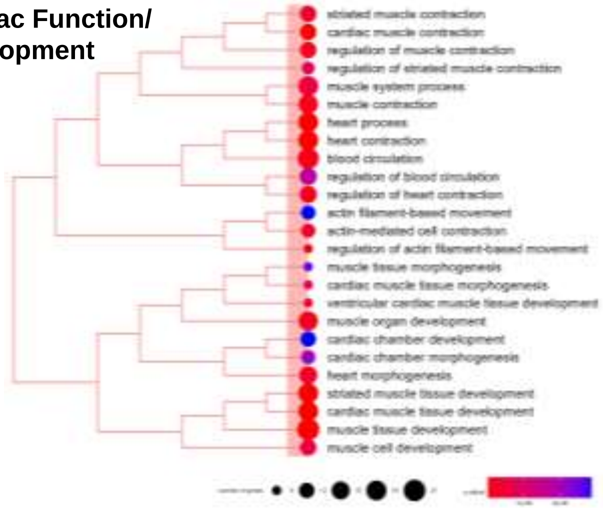


**Top 100 of 483 Start-End DEG**

**Enriched Gene Sets = 305**

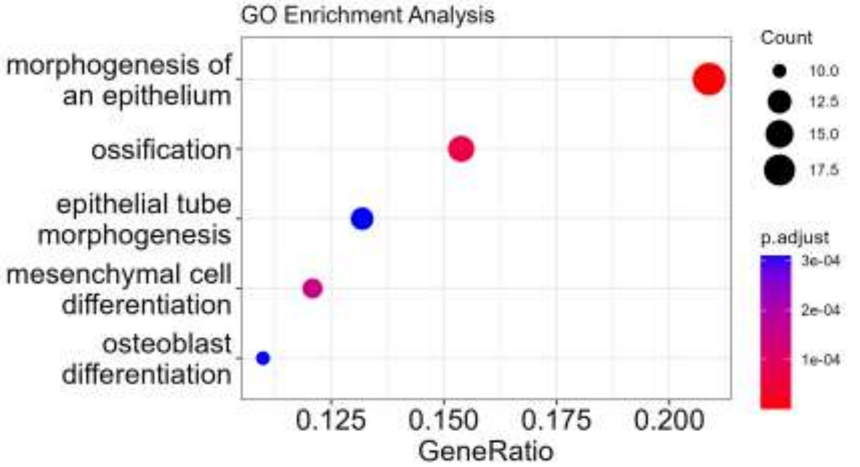


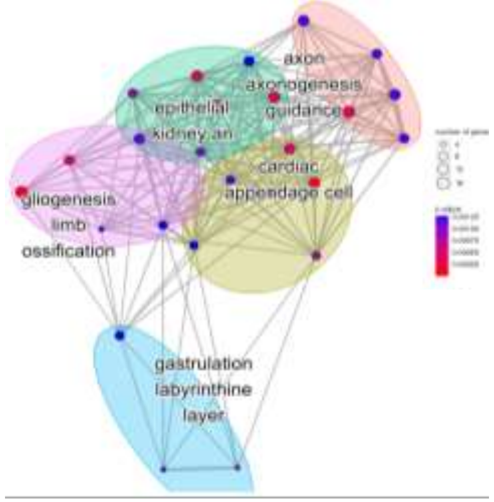


**Mesenchyme/ Cardiac/Limb Morphogenesis**

**Epithelium/ Urogen Dev**

**Ossification/ Osteoblast Diff**

**Placenta Dev**

**Axon Dev**

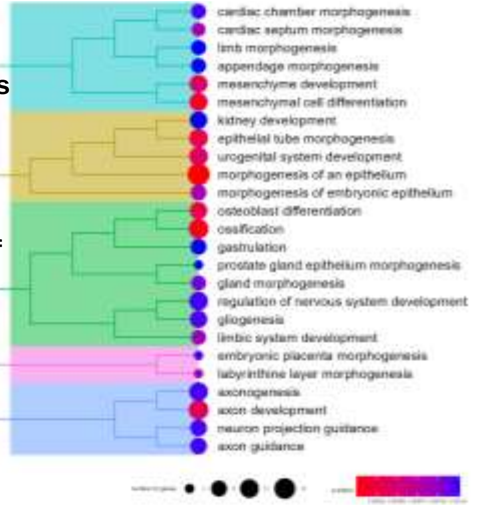

Fig. S12
Lineage DEG SS

| Patient | D09B | CPROG-A/CM-A | 4,091 cells | Lineage DE Analyses & Enrichment |
|---|---|---|---|---|

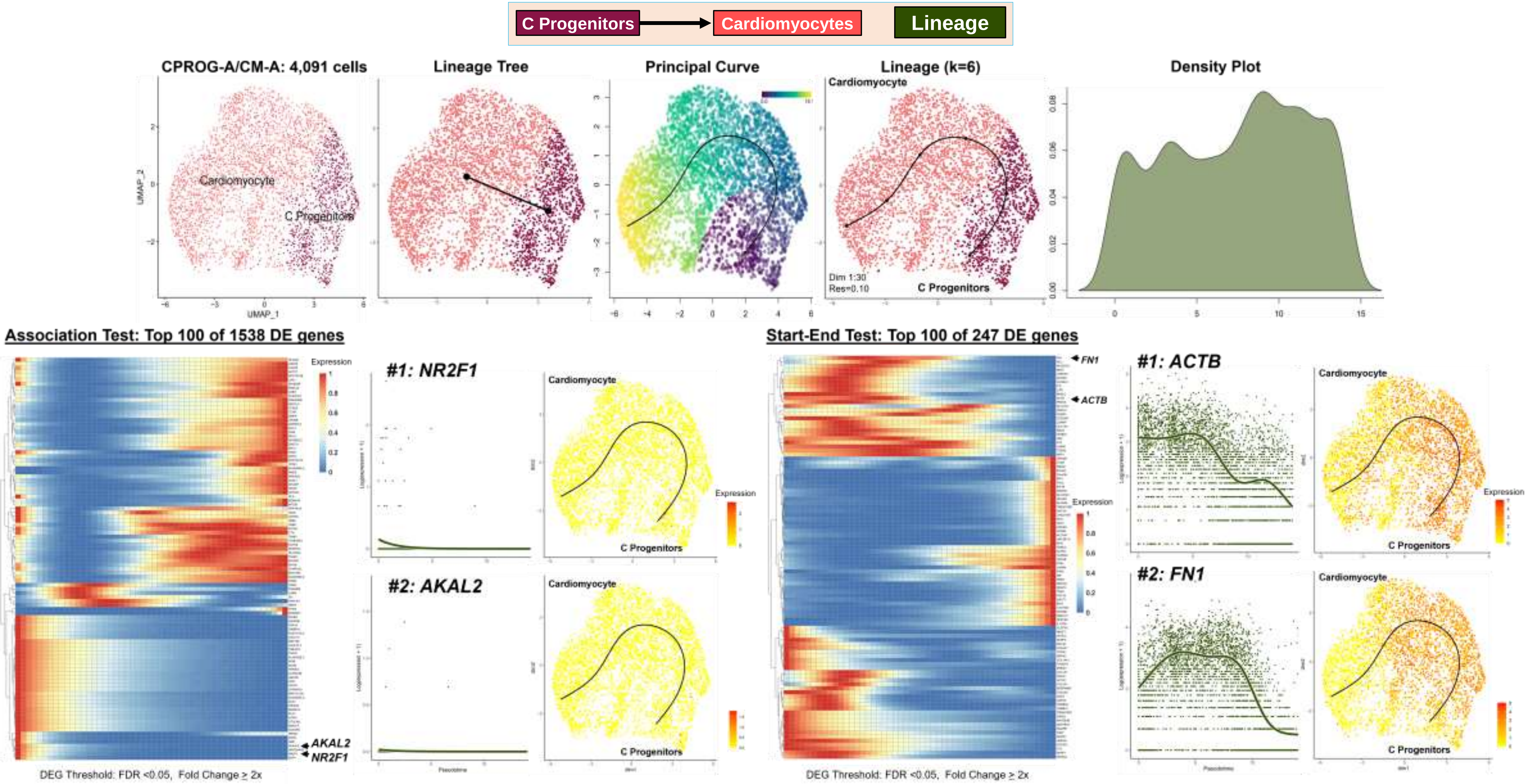

Fig. S12
Lineage DEG SS

| Patient | D09B | CPROG-A/CM-A | 4,091 cells | Lineage DE Analyses & Enrichment |
|---|---|---|---|---|

#### Over-Representation Analysis

C Progenitors → Cardiomyocytes | Lineage

| Enrichment Plots: Top 5 Gene Sets | Enrichment Maps Top 25 Gene Sets | Enrichment Trees Top 25 Gene Sets |
|---|---|---|

**Top 100 of 1538 Association DEG**

**Enriched Gene Sets = 117**

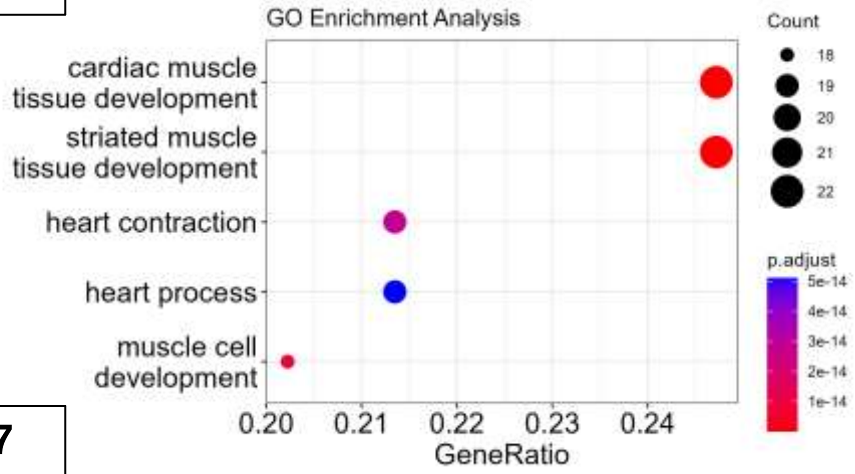


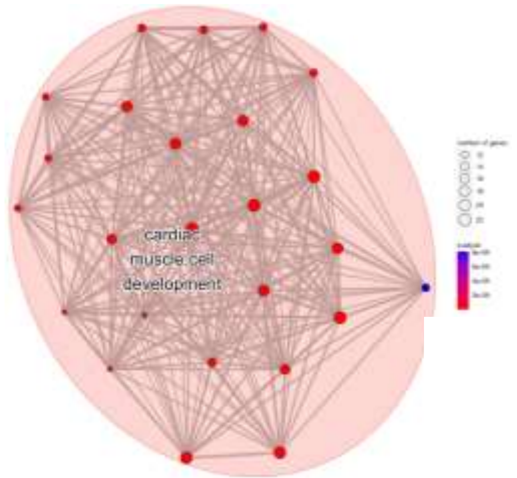


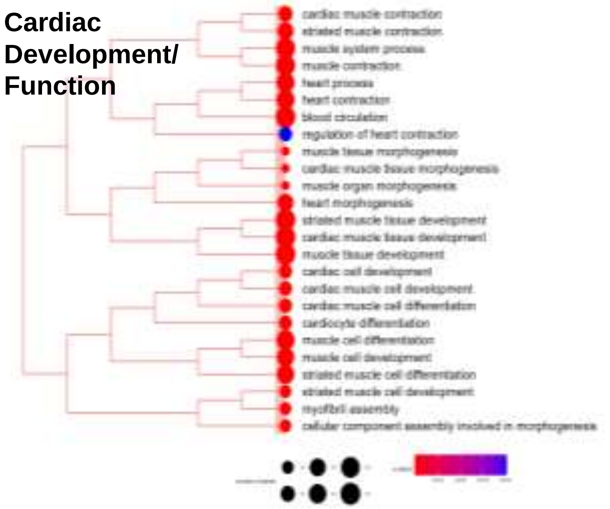


**Top 100 of 247 Start-End DEG**

**Enriched Gene Sets = 195**

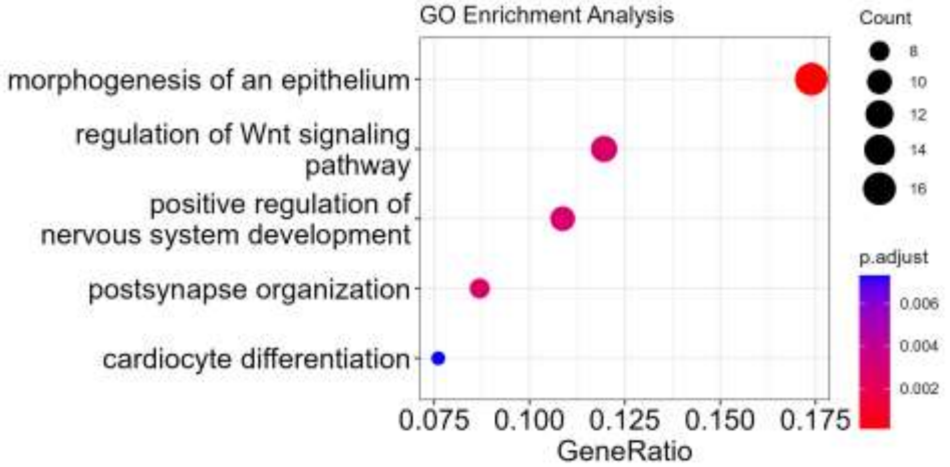


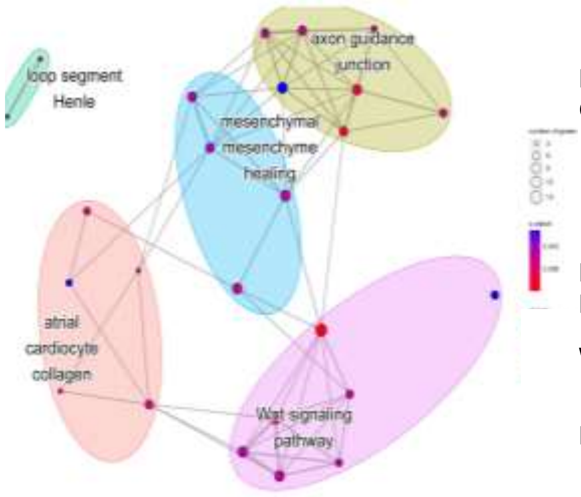


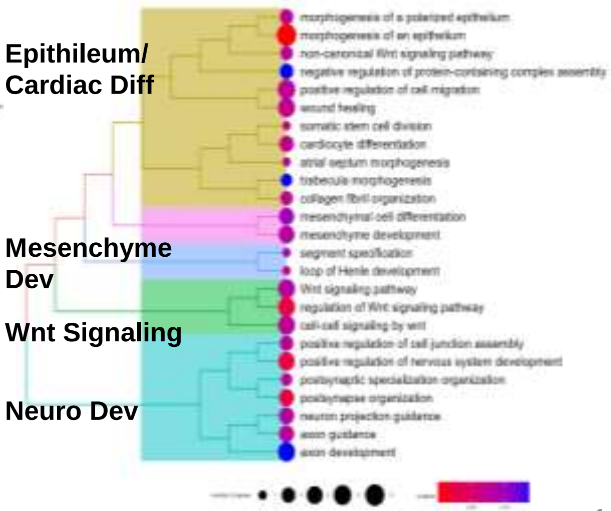

**Fig. S13 Lineage DEG INTEGR**

### Workflow Step-III: Paired Subset Data Results

#### Trajectory Analyses for Lineage-Specific Differential Expression and Enrichment

##### A. Summary: Lineage Differentially Expressed Genes (Lineage DEG)

**Integrated Subset Data: 62,488 Total Cells**

| Dataset | Toplogy/ Lineage(s) | Condiments Tests (p=) Topology | Progress | Differ | DEG Type[a] | ORA[b] DEGs | GS | Top GO BP Enriched GS | GSEA[b] DEGs | GS | MSigDB Hallmark Enriched GS |
|---|---|---|---|---|---|---|---|---|---|---|---|
| DAY 0: Integrated BALANCED Subsets (Pluripotent-A, Pluripotent-B) 19,346 cells | Single Trajectory | Different (2.2 e-16) | Different (1.5 e-14) | N/A | AT | 152 | 43 | Metal Homeostasis | 1130 | 0 | None |
| | | | | | CT | 97 | 4 | Metal Homeostasis | 827 | 0 | None |
| | 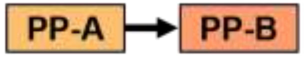 | | | | Top 100 | | | | | | |
| | | | | | Gr-A | 74 | 22 | Metal Homeostasis | | | |
| | | | | | Gr-B | 23 | 2 | Transcription | | | |
| Day 9B,16,19: Integrated BALANCED Subsets (C Prog, CM-A, EPDC) 43,142 cells | Bifurcating Trajectory | Different (6.9 e-4) | Different (4.0 e-12) | Different (6.8 e-4) | AT | 2452 | 218 | Mitosis/Cell Cycle | 2880 | 6 | E2F Targets |
| | | | | | CT | 45 | 0 | None | 2290 | 0 | G2M Checkpoint |
| | | | | | L-1 CT | 391 | 19 | | 1404 | 0 | Spermatogenesis<br>Mitotic Spindle<br>DNA Repair<br>MYC Targets V1 |
| | 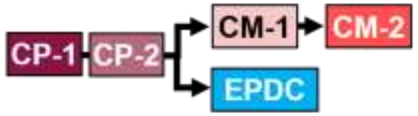 | | | | Top 100 | | | | | | |
| | | | | | Gr-A | 10 | 3 | Muscle Adaption<br>Aerobic Respiration | | | |
| | | | | | Gr-B | 10 | 0 | None | | | None |
| | | | | | Gr-C | 59 | 6 | Growth Factor Response<br>Metal Homeostasis | | | |
| | | | | | Gr-D | 21 | 61 | BMP Signaling | | | |
| | | | | | L-2 CT | 25 | 0 | | 2191 | 0 | None |
| | | | | | Top 25 | | | | | | |
| | | | | | Gr-A | 19 | 2 | Phagocytosis | | | |
| | | | | | Gr-B | 3 | 10 | Antigen Processing<br>Dosage Compensation | | | |
| | | | | | Gr-C | 3 | 12 | Cyclin Kinase | | | |

[a]Lineage DEG Type: Association Test (AT) and Condition Test (CT)
[b]Parameters: 1. ORA: DEG Threshold= FDR<0.05 & FC>2x, rank by Wald stat; min #DEGs= 3, adj p-value<0.05, q-value cutoff 0.20; 2. GSEA: DEG Threshold= FDR<0.05 & FC>2x, WaldStat>0.0; scoreType "pos", Adj p-value cutoff= 0.05, GS size 10-500
Abbreviations: PP, pluripotent; C Prog/CP, Cardiac Progenitors; CM, Cardiomyocyte; EPDC, Epicardium-derived cells; Progress, Progression; Differ, Differentiation; DEG, differentially expressed gene; ORA, Over-Representation Analysis; GO BP, Gene Ontology Biological Processes; GS, gene set; GSEA, Gene Set Enrichment Analysis; MSigDB, Molecular Signatures Database; Gr, Group; L, Lineage

**Fig. S13 Lineage DEG**
**INTEGR**

#### B. Summary: UMAP Plots and Cell Lineages (n=2: 62,488 Total Cells)

**Control Subset Data** ✚ **Patient Subset Data** → **Integrated Subset Data** → **Cell Lineages**

### D00 PP-AB

**19,346 cells**

**Integrated**

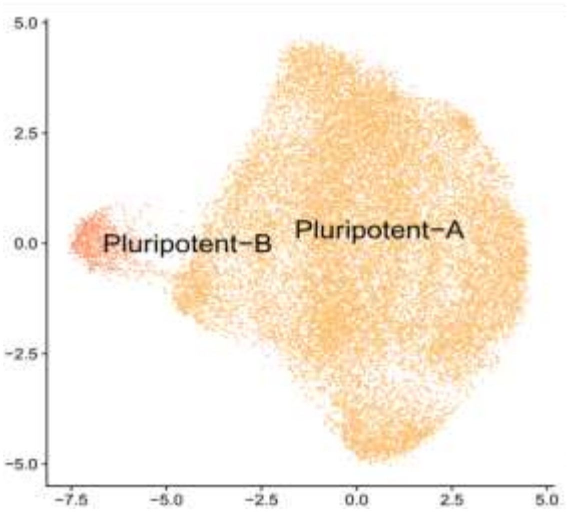


**Lineages**

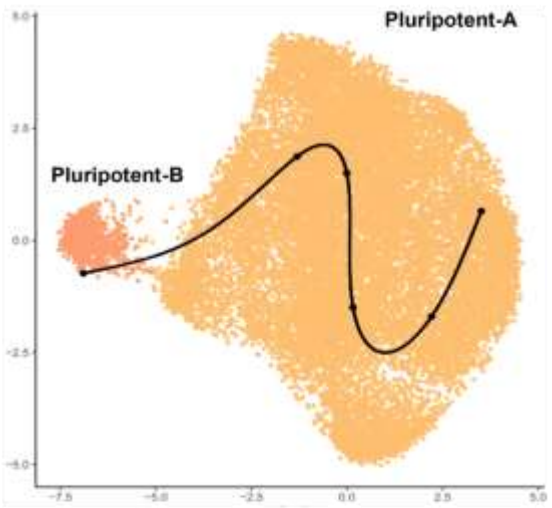


##### D09B-D16-D19 CPROG/CM-A/EPDC

**43,142 cells**

**Integrated**

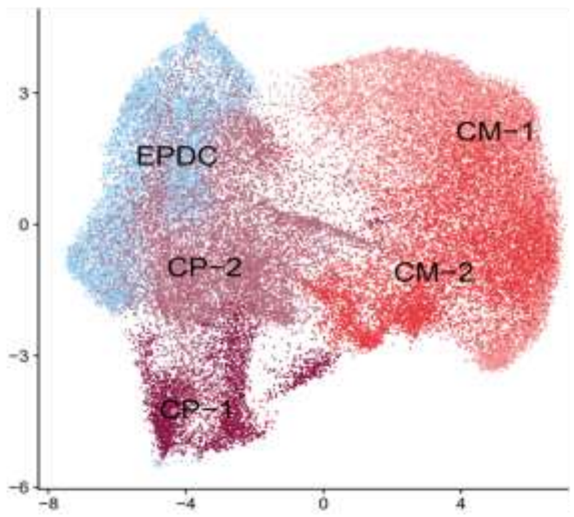


**Lineages**

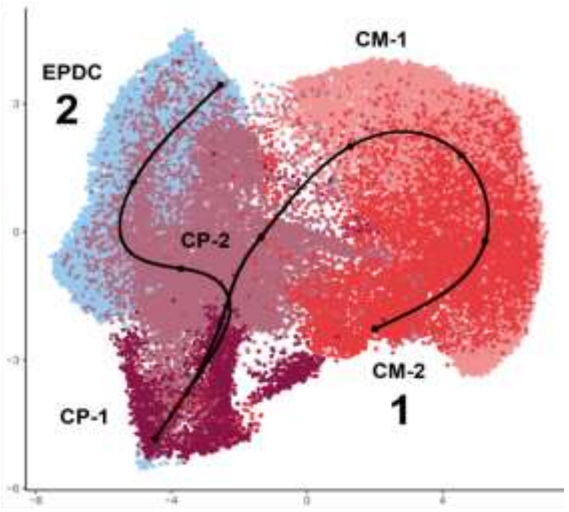

**Fig. S13 Lineage DEG**
**INTEGR**

### C. Paired Subset Data Analyses: Pluripotent Cell Lineage (19,346 Total Cells)

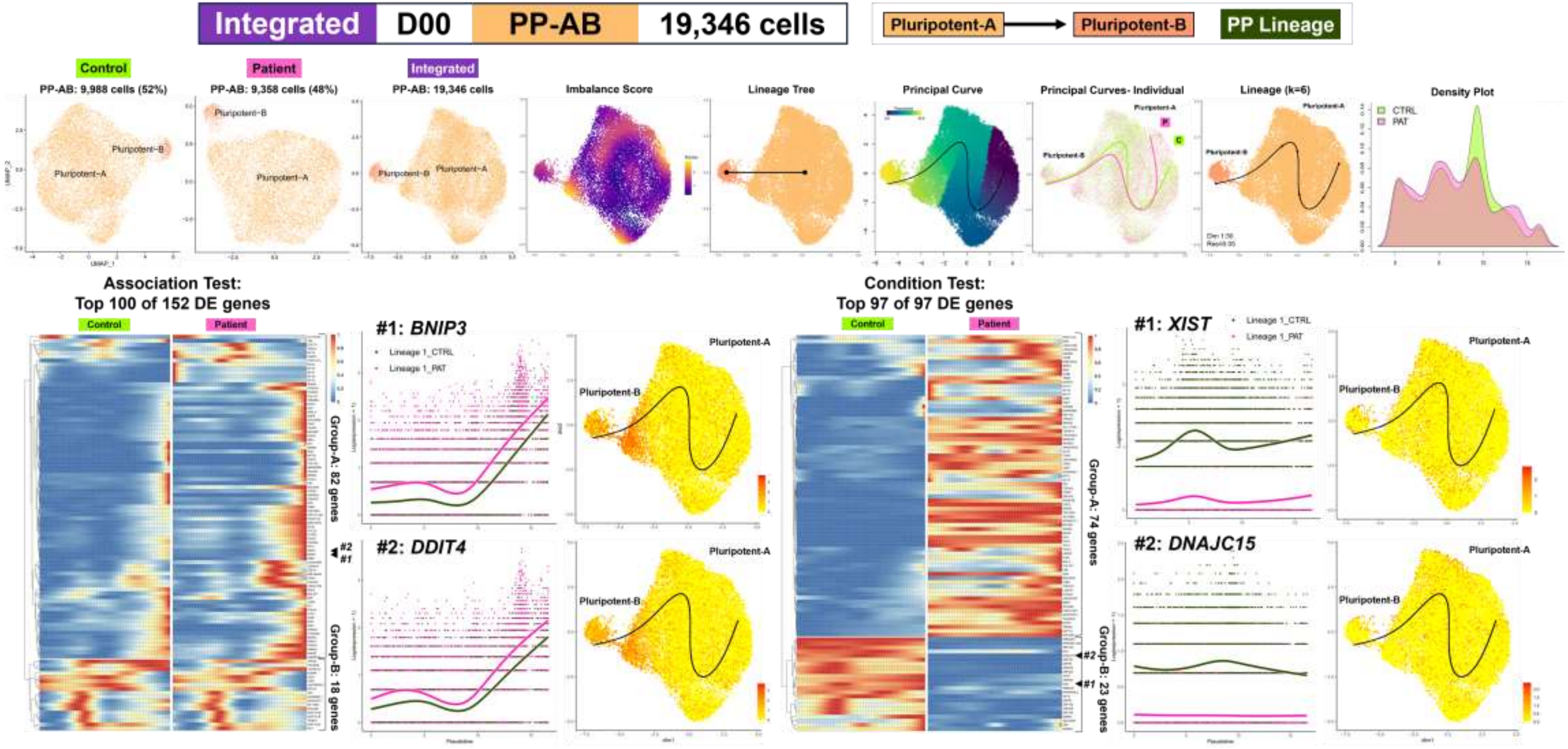

**Fig. S13 Lineage DEG INTEGR**

| Integrated | D00 | PP-AB | 19,346 cells |
|---|---|---|---|

Pluripotent-A → Pluripotent-B | PP Lineage

### ASSOCIATION Test DEG Enrichment

#### Over-Representation Analysis

**Top 100 of 152 Association DEG**

**Enrichment Plots: Top 5 Gene Sets**

**Enrichment Maps Top 25 Gene Sets**

**Enrichment Trees Top 25 Gene Sets**

**Group-A**
**82 genes**
**Enriched Gene Sets = 48**

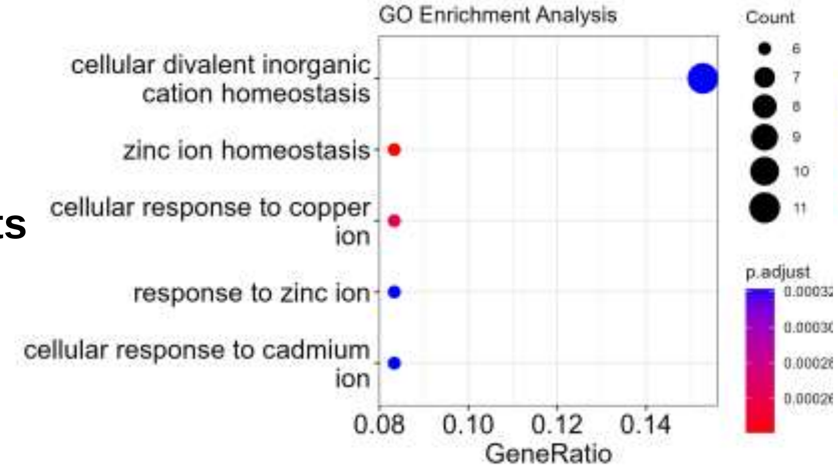


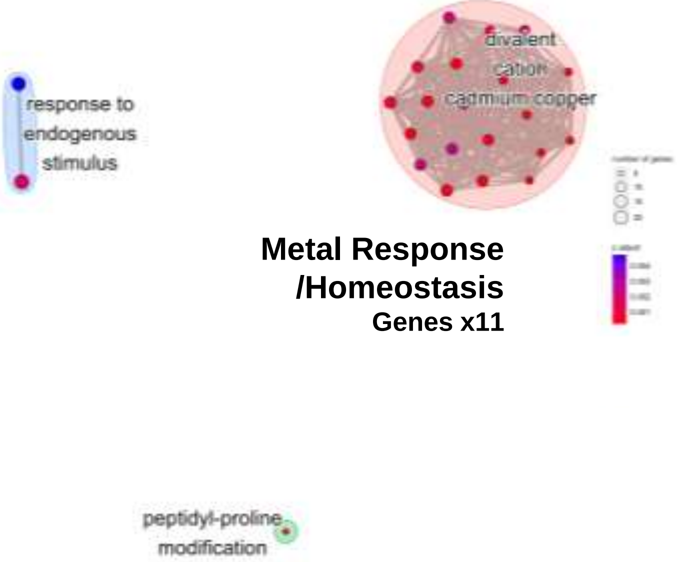


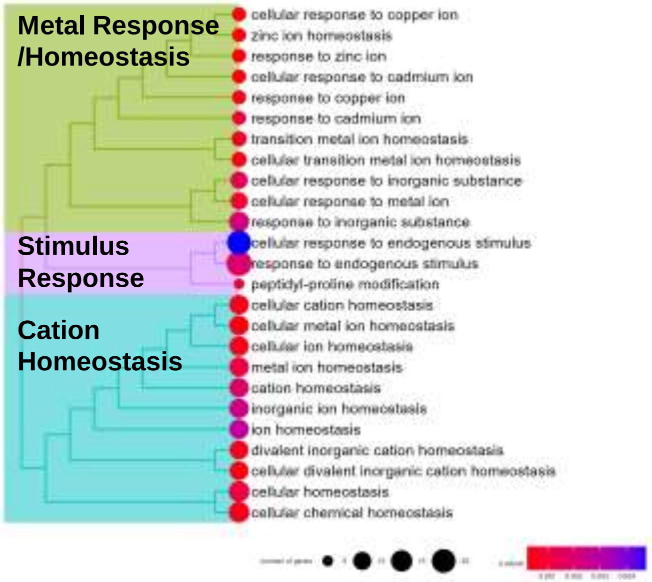


**Group-B**
**18 genes**
**Enriched Gene Sets = 0**

**No Enriched GO BP Gene Sets**

**No Enriched GO BP Gene Sets**

**No Enriched GO BP Gene Sets**

---

#### Gene Set Enrichment Analysis

**Association Test: 1130 DEG Total**

**No Enriched MSigDB Hallmark Gene Sets**

**Fig. S13 Lineage DEG**
**INTEGR**

| Integrated | D00 | PP-AB | 19,346 cells |
|---|---|---|---|

Pluripotent-A → Pluripotent-B  PP Lineage

#### CONDITION Test DEG Enrichment

##### Over-Representation Analysis

**Top 97 of 97 Condition DEG**

**Enrichment Plots: Top 5 Gene Sets**

**Enrichment Maps Top 25 Gene Sets**

**Enrichment Trees Top 25 Gene Sets**

**Group-A**
**74 genes**
**Enriched Gene Sets = 22**

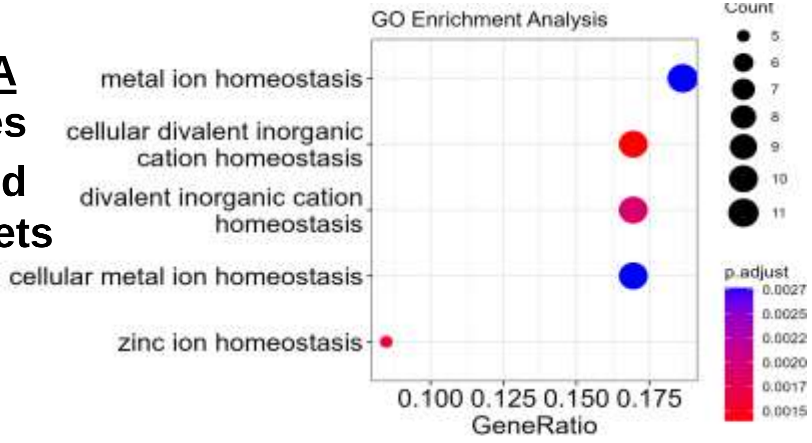


**Metal/Cation Homeostasis**
**Genes x11**

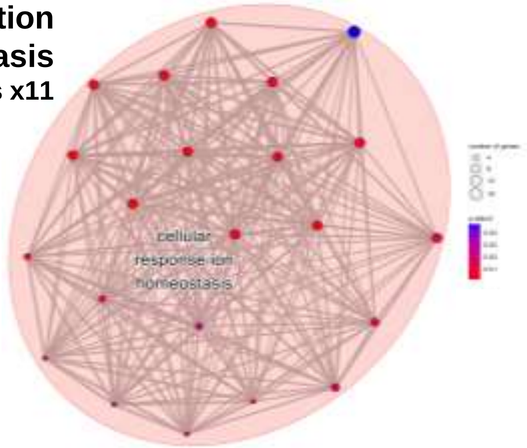


**Metal/Cation Homeostasis**

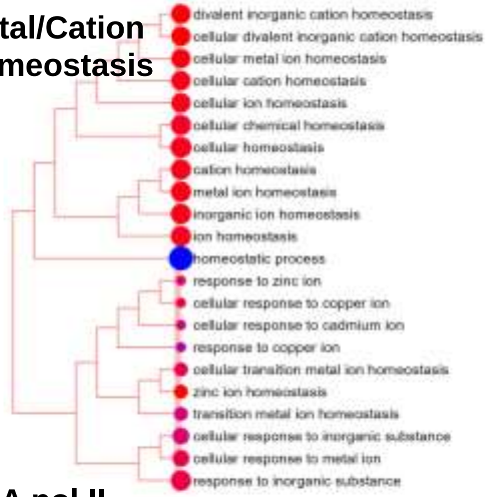


**Group-B**
**23 genes**
**Enriched Gene Sets = 2**

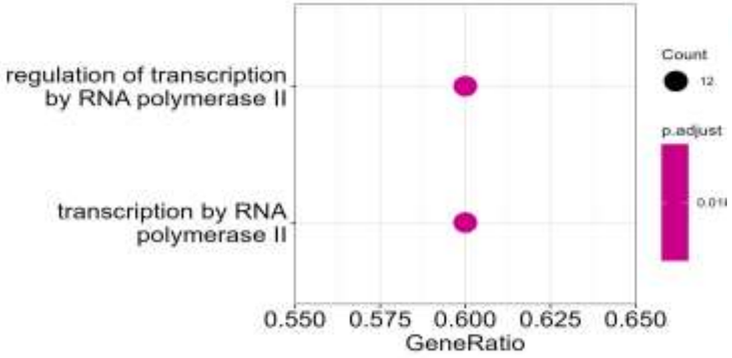


**RNA pol II transcription**
**Genes x12**

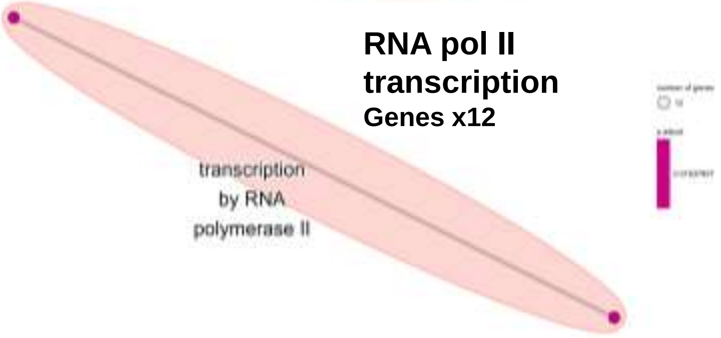


**RNA pol II transcription**

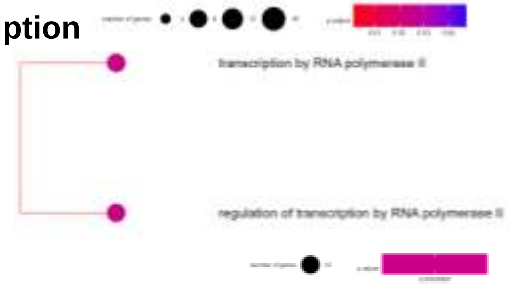


**Gene Set Enrichment Analysis**

**Condition Test: 827 DEG Total**

**No Enriched MSigDB Hallmark Gene Sets**

**Fig. S13 Lineage DEG**
**INTEGR**

#### D. Paired Subset Data Analyses: Cardiac Progenitor Lineages (43,142 Total Cells)

| Integrated | D09B-D16-D19 | CPROG/CM-A/EPDC | 43,142 cells |
|---|---|---|---|

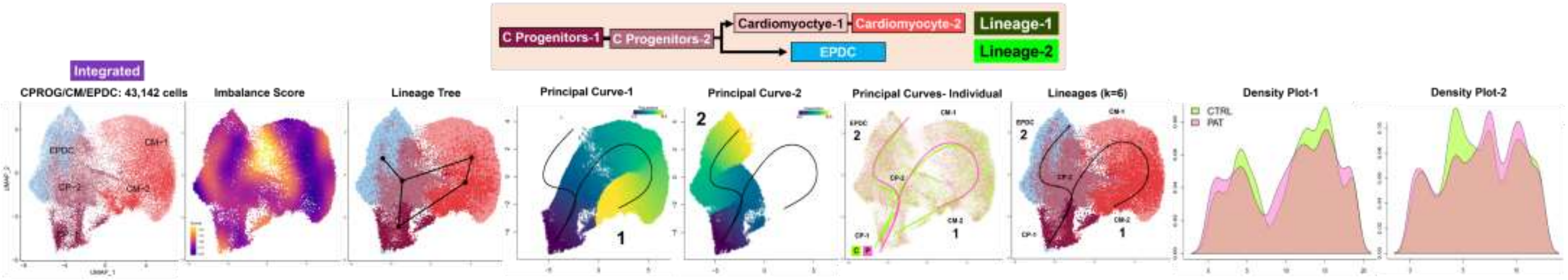


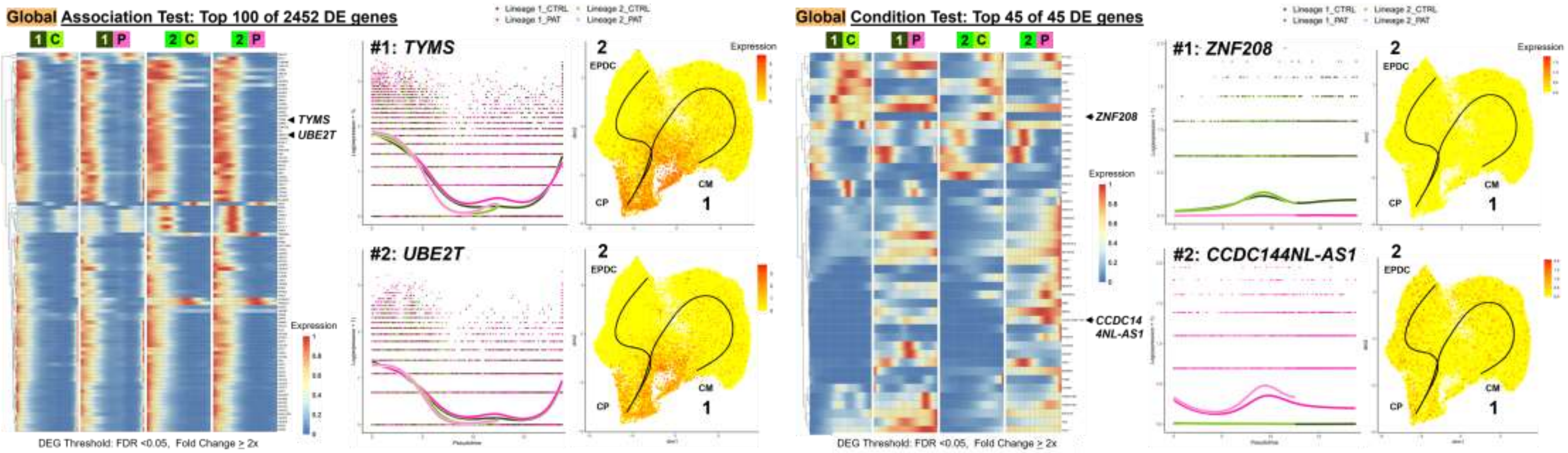

**Fig. S13 Lineage DEG INTEGR**

| Integrated | D09B-D16-D19 | CPROG/CM-A/EPDC | 43,142 cells |
|---|---|---|---|

#### Over-Representation Analysis

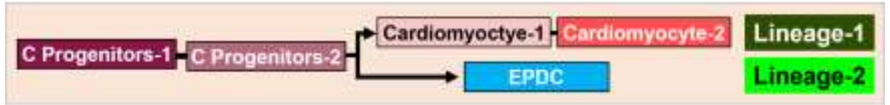


**Top 100 of 2452 Association DEG**

**Enrichment Plots: Top 5 Gene Sets**

**Enrichment Maps Top 25 Gene Sets**

**Enrichment Trees Top 25 Gene Sets**

**Enriched Gene Sets = 218**

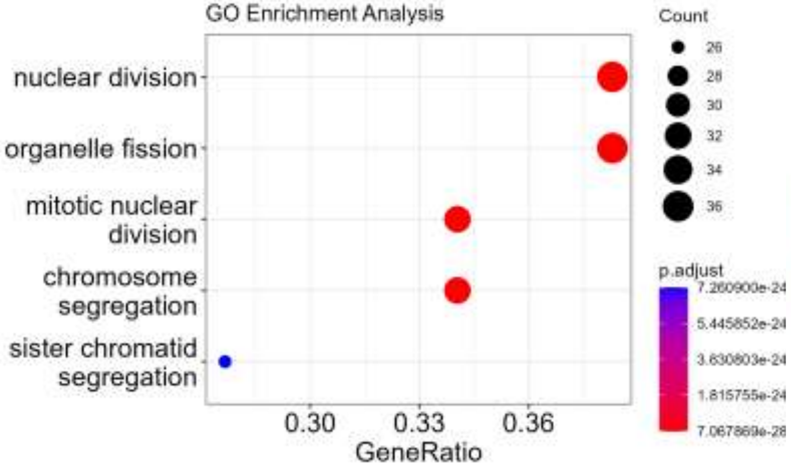


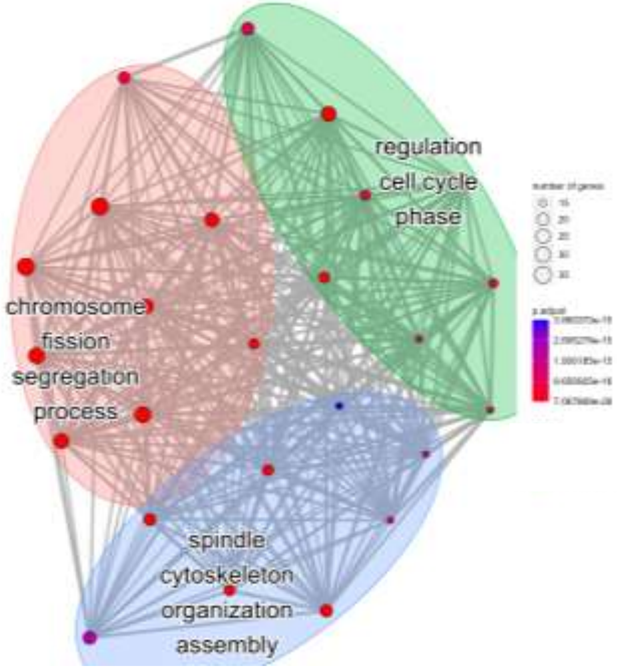


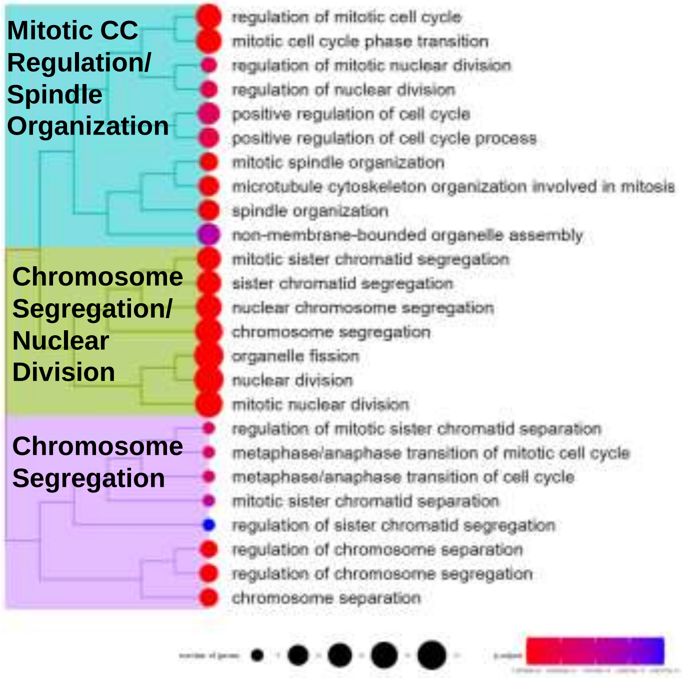


**Top 45 of 45 Condition DEG**

**Enriched Gene Sets = 0**

**No Enriched GO BP Gene Sets**

**No Enriched GO BP Gene Sets**

**No Enriched GO BP Gene Sets**

**Fig. S13 Lineage DEG INTEGR**

| Integrated | D09B-D16-D19 | CPROG/CM-A/EPDC | 43,142 cells |
|---|---|---|---|

#### Gene Set Enrichment Analysis

**Association Test: 2880 DEG Total**

**Enriched Gene Sets = 6**

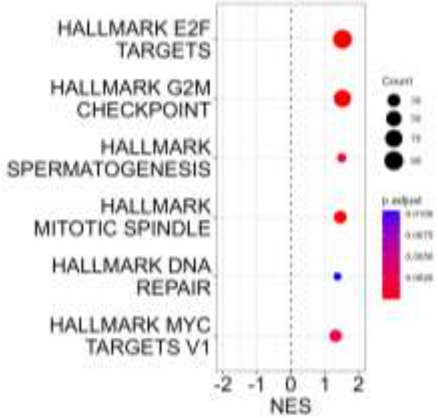


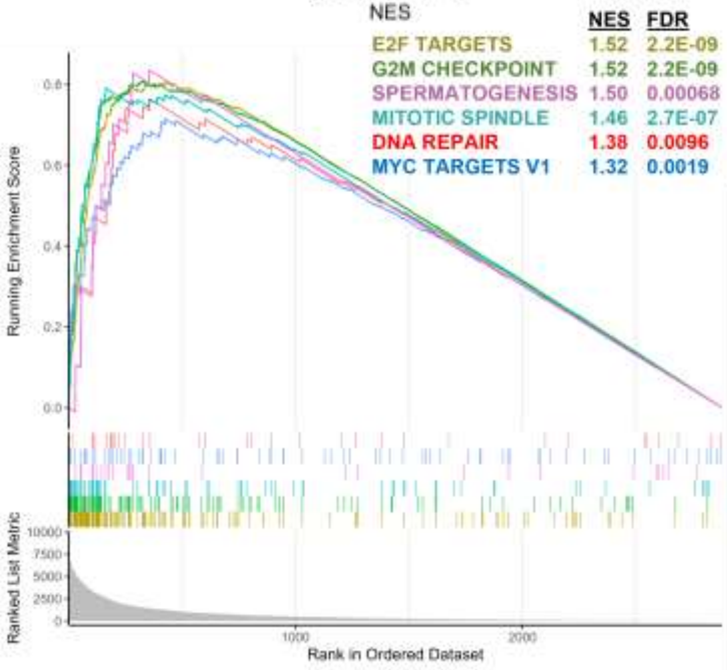


**MODULE SCORING**

**E2F TARGETS**

**G2M CHECKPOINT**

**MITOTIC SPINDLE**

**MYC TARGETS V1**

**SPERMATO-GENESIS**

**DNA REPAIR**

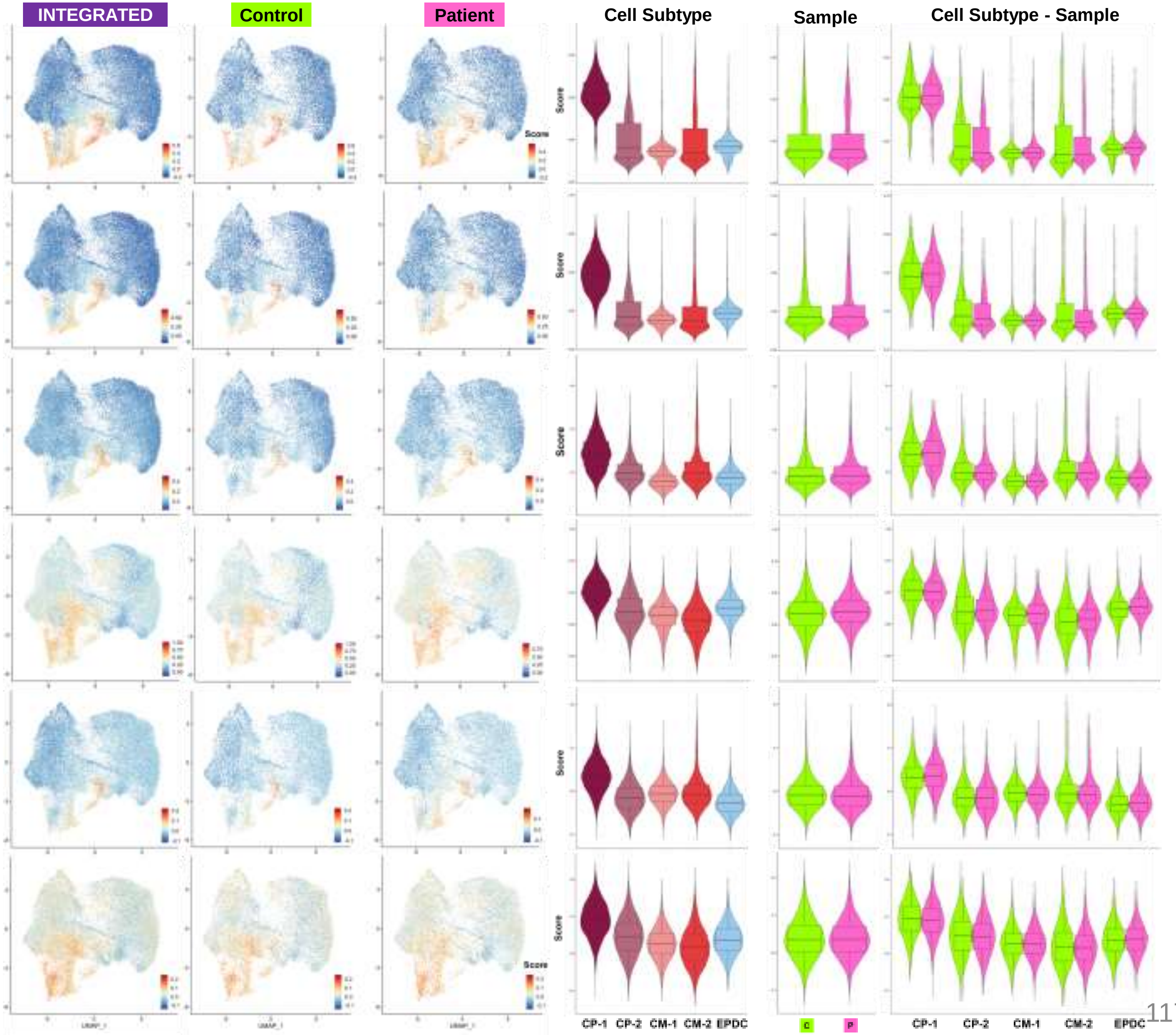

**Fig. S13 Lineage DEG INTEGR**

| Integrated | D09B-D16-D19 | CPROG/CM-A/EPDC | 43,142 cells |
| --- | --- | --- | --- |

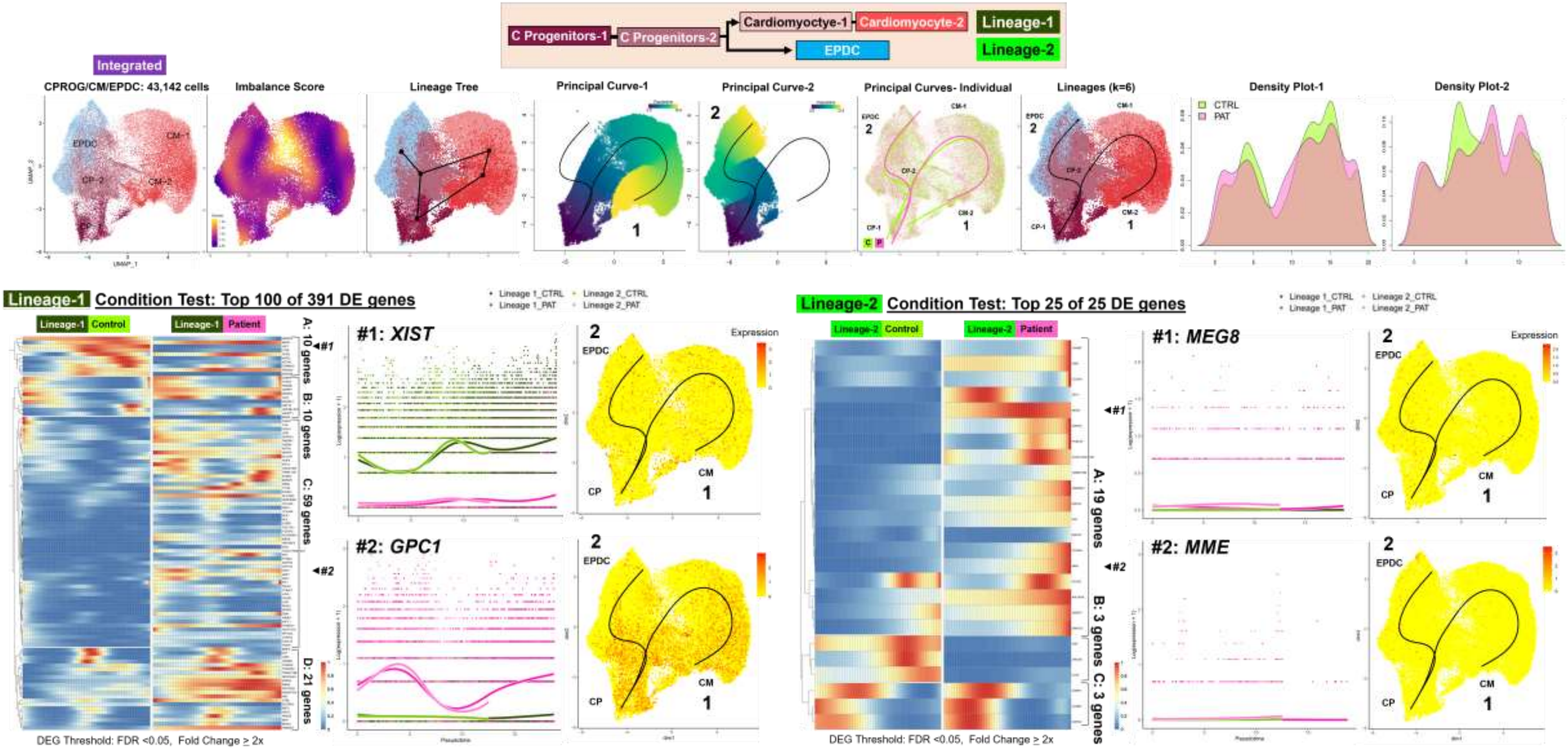

**Fig. S13 Lineage DEG INTEGR**

| Integrated | D09B-D16-D19 | CPROG/CM-A/EPDC | 43,142 cells |
|---|---|---|---|

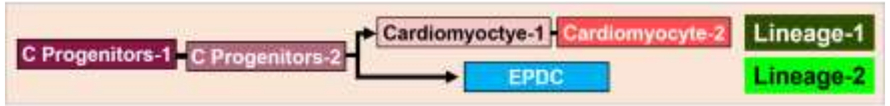


#### Over-Representation Analysis

##### Lineage-1

**Top 100 of 391 Condition DEG**

**Enrichment Plots: Top 5 Gene Sets**

**Enrichment Maps Top 25 Gene Sets**

**Enrichment Trees Top 25 Gene Sets**

**Group A:**
**10 genes**

**Enriched Gene Sets = 3**

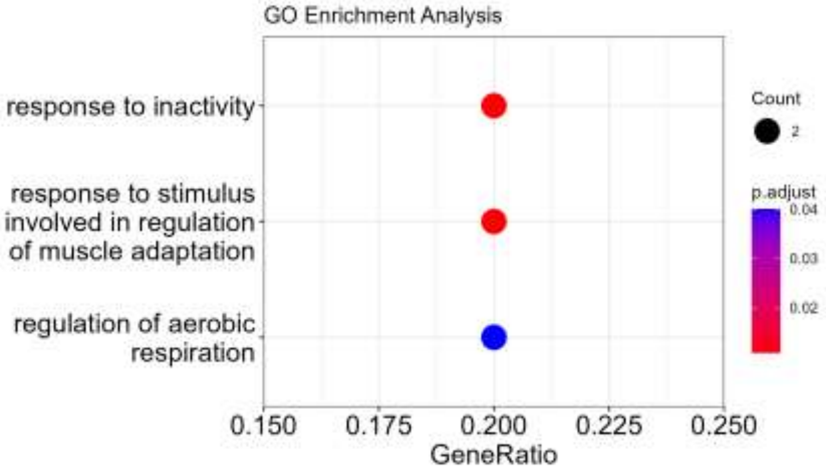


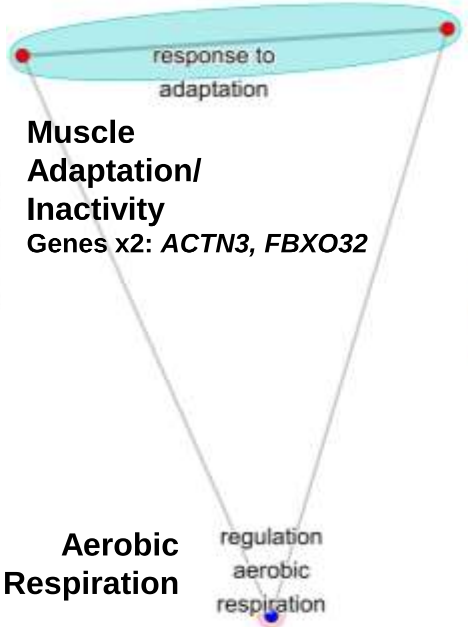


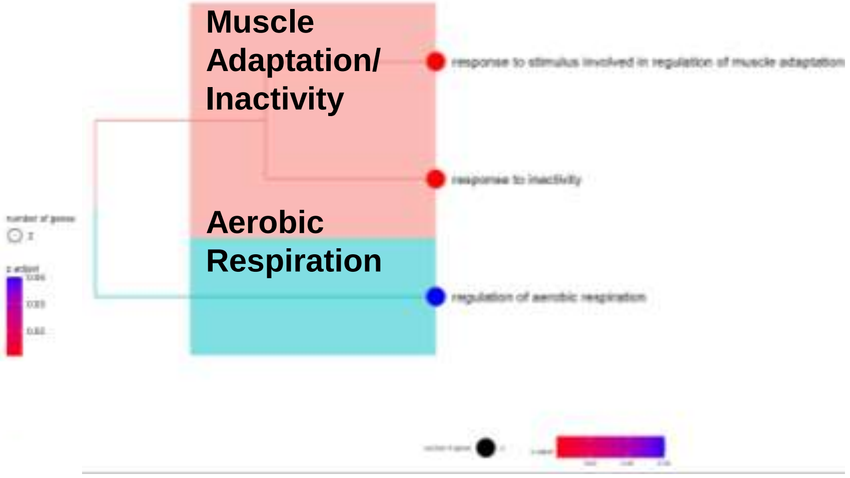


**Group B:**
**10 genes**

**Enriched Gene Sets = 0**

No Enriched GO BP Gene Sets

No Enriched GO BP Gene Sets

No Enriched GO BP Gene Sets

Fig. S13 Lineage DEG
INTEGR

| Integrated | D09B-D16-D19 | CPROG/CM-A/EPDC | 43,142 cells |
|---|---|---|---|

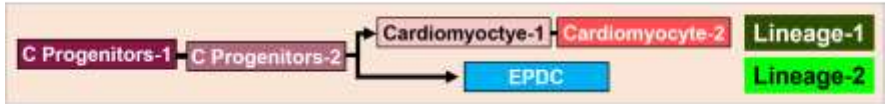


#### Over-Representation Analysis

#### Lineage-1

**Top 100 of 391 Condition DEG**

**Enrichment Plots: Top 5 Gene Sets**

**Enrichment Maps Top 25 Gene Sets**

**Enrichment Trees Top 25 Gene Sets**

**Group C: 59 genes**

**Enriched Gene Sets = 6**

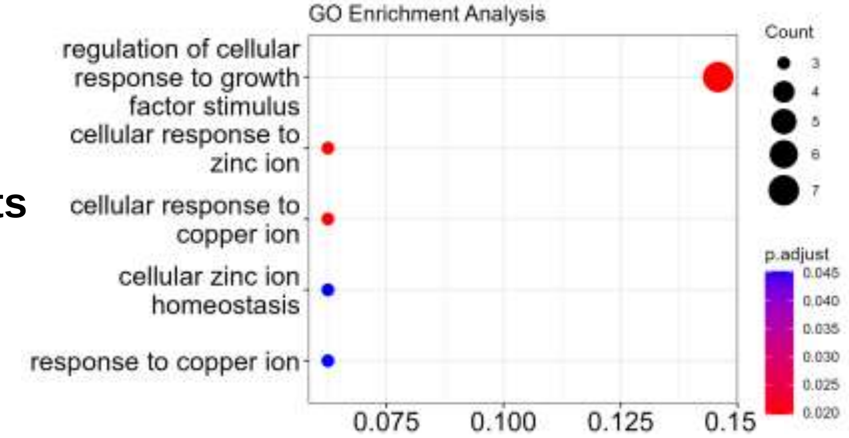


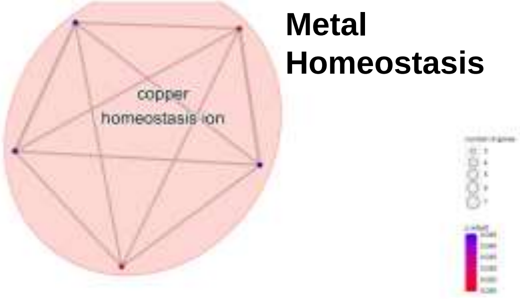


**Metal Homeostasis**

**Growth Factor Response**

**Genes x7:** ***FOXD1, PMEPA1, CCBE1, DCNMT3, MT3, GPC1, SFRP5***

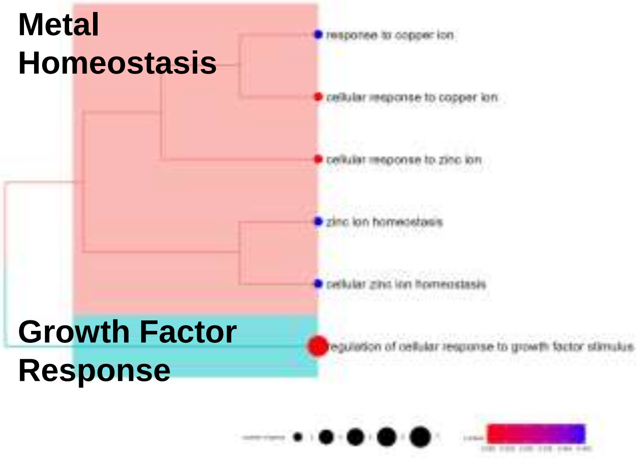


**Metal Homeostasis**

**Growth Factor Response**

**Group D: 21 genes**

**Enriched Gene Sets = 61**

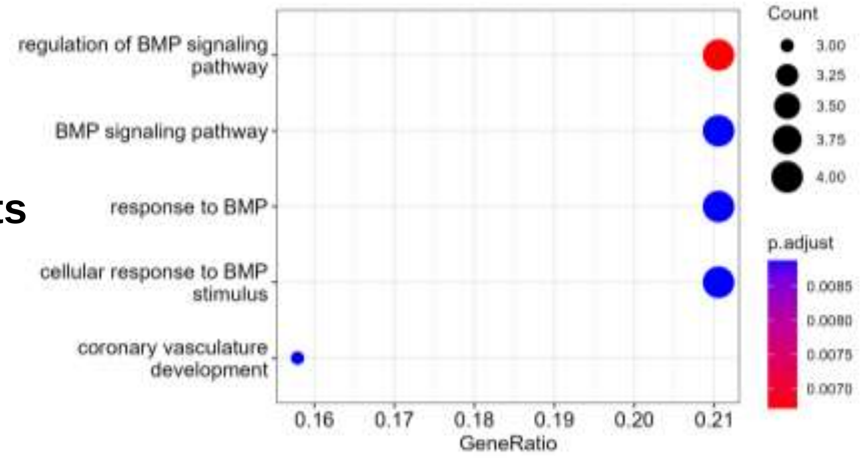


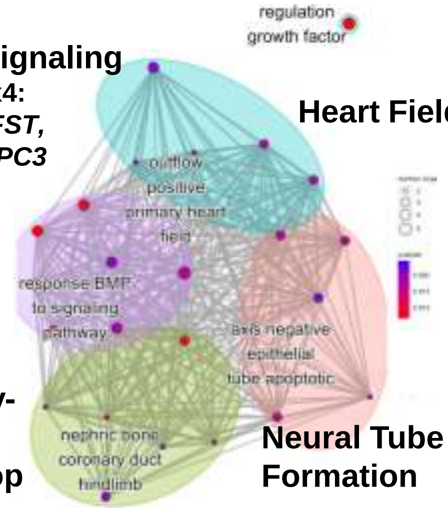


**BMP Signaling**
**Genes x4:** ***BMP4, FST, LRP2,GPC3***

**Heart Field**

**Kidney-Limb Develop**

**Neural Tube Formation**

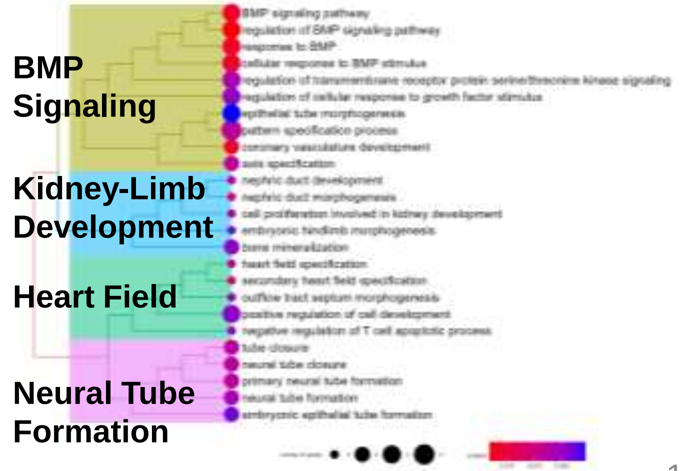


**BMP Signaling**

**Kidney-Limb Development**

**Heart Field**

**Neural Tube Formation**

**Fig. S13 Lineage DEG**
**INTEGR**

| Integrated | D09B-D16-D19 | CPROG/CM-A/EPDC | 43,142 cells |
|---|---|---|---|

#### Over-Representation Analysis

**Lineage-2**

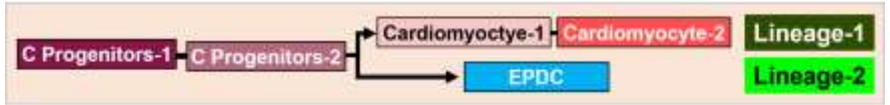


**Top 25 of 25 Condition DEG**

**Enrichment Plots: Top 5 Gene Sets**

**Enrichment Maps Top 25 Gene Sets**

**Enrichment Trees Top 25 Gene Sets**

**Group A:**
**19 genes**

**Enriched Gene Sets = 2**

**Group B:**
**3 genes**

**Enriched Gene Sets = 10**

**Group C:**
**3 genes**

**Enriched Gene Sets = 12**

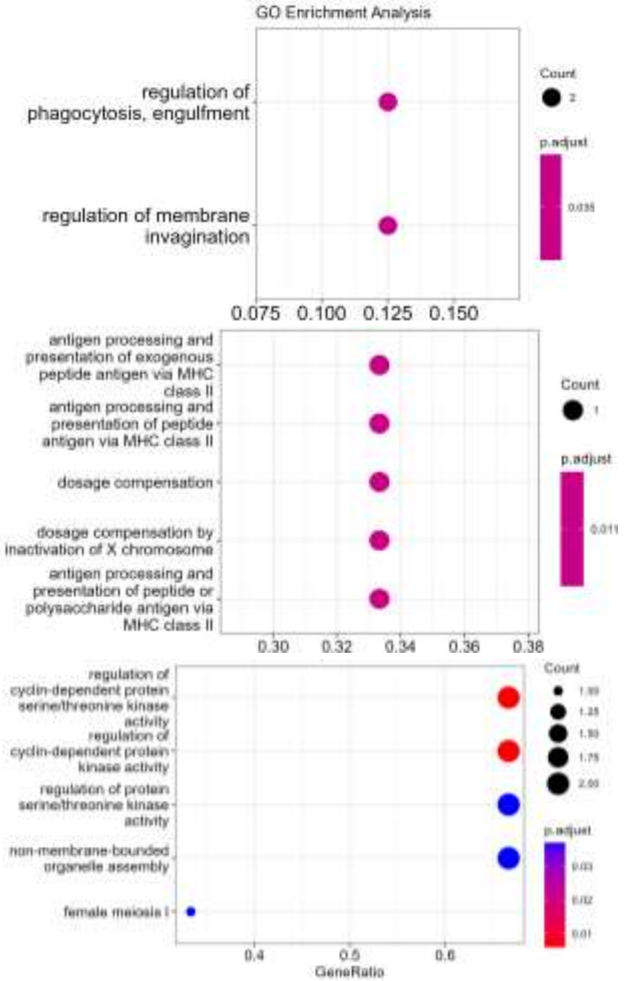


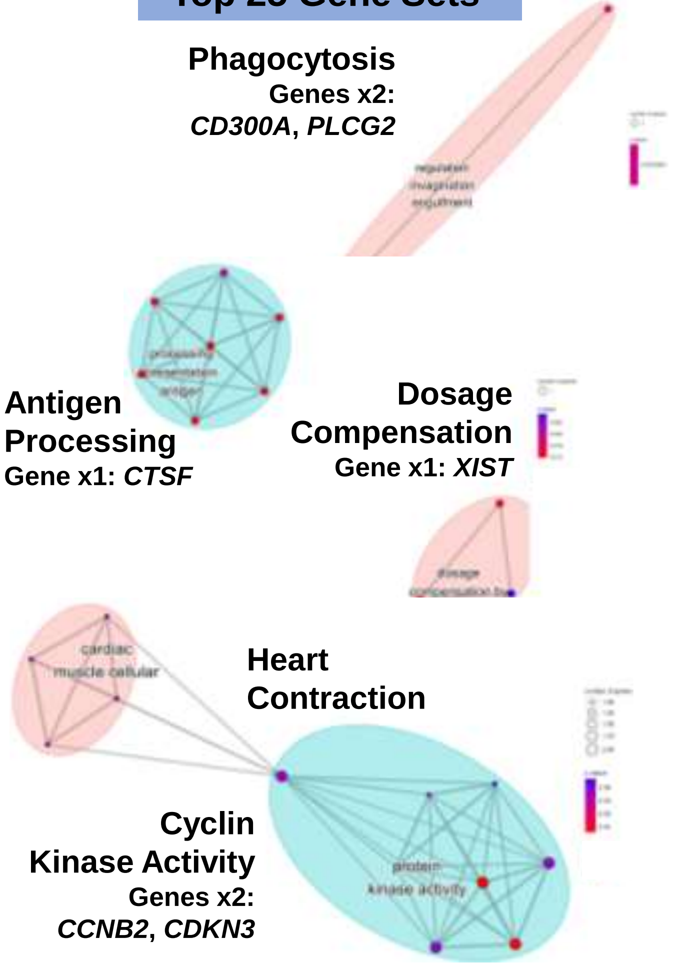


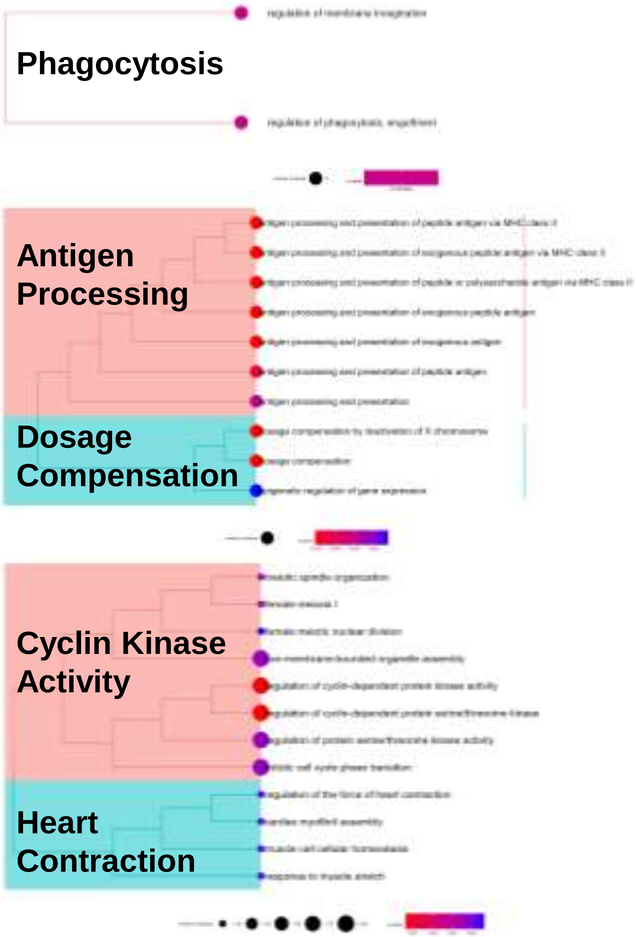

Fig. S14

#### Blot 1

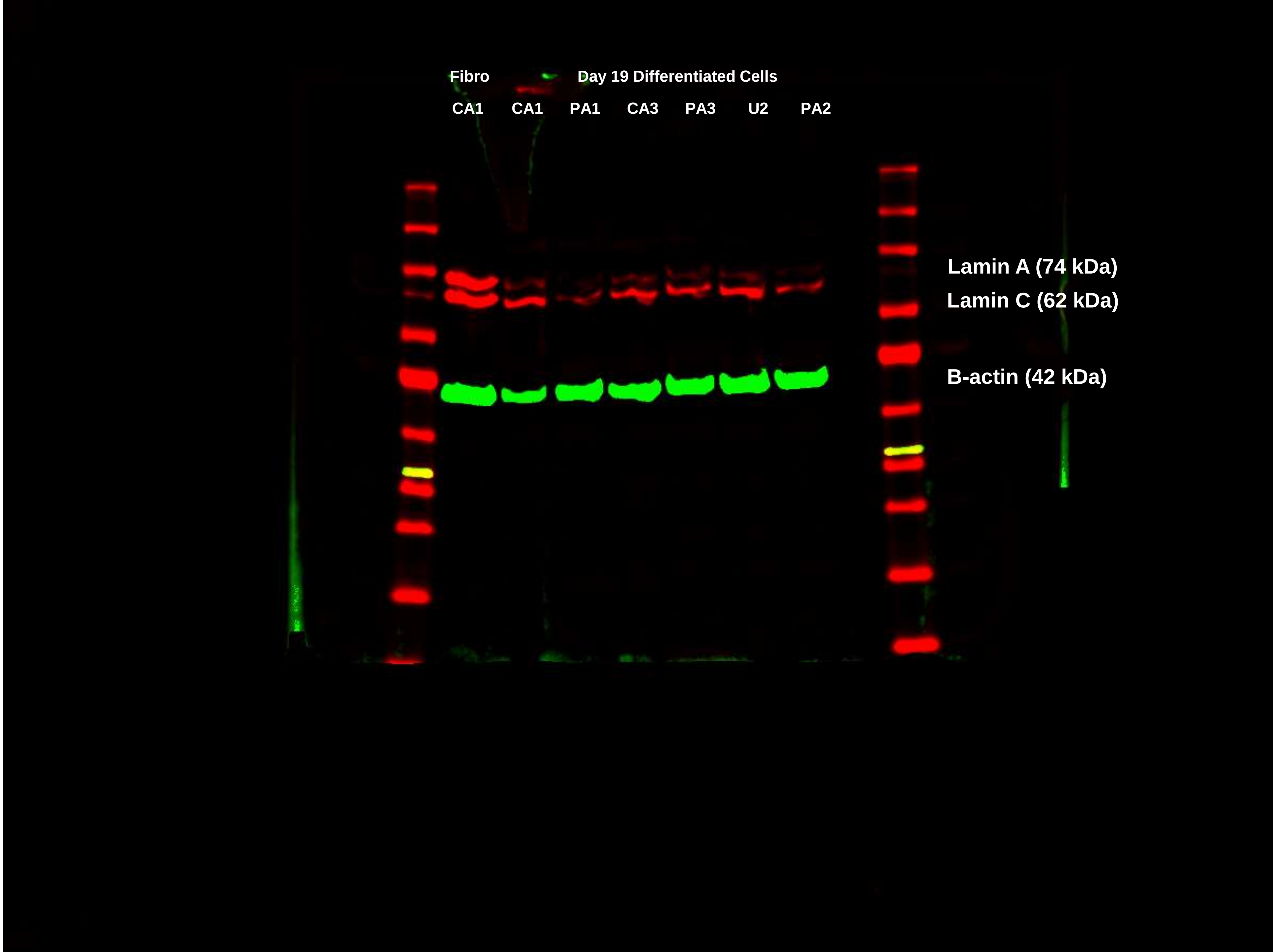

Fig. S14

#### Blot 2

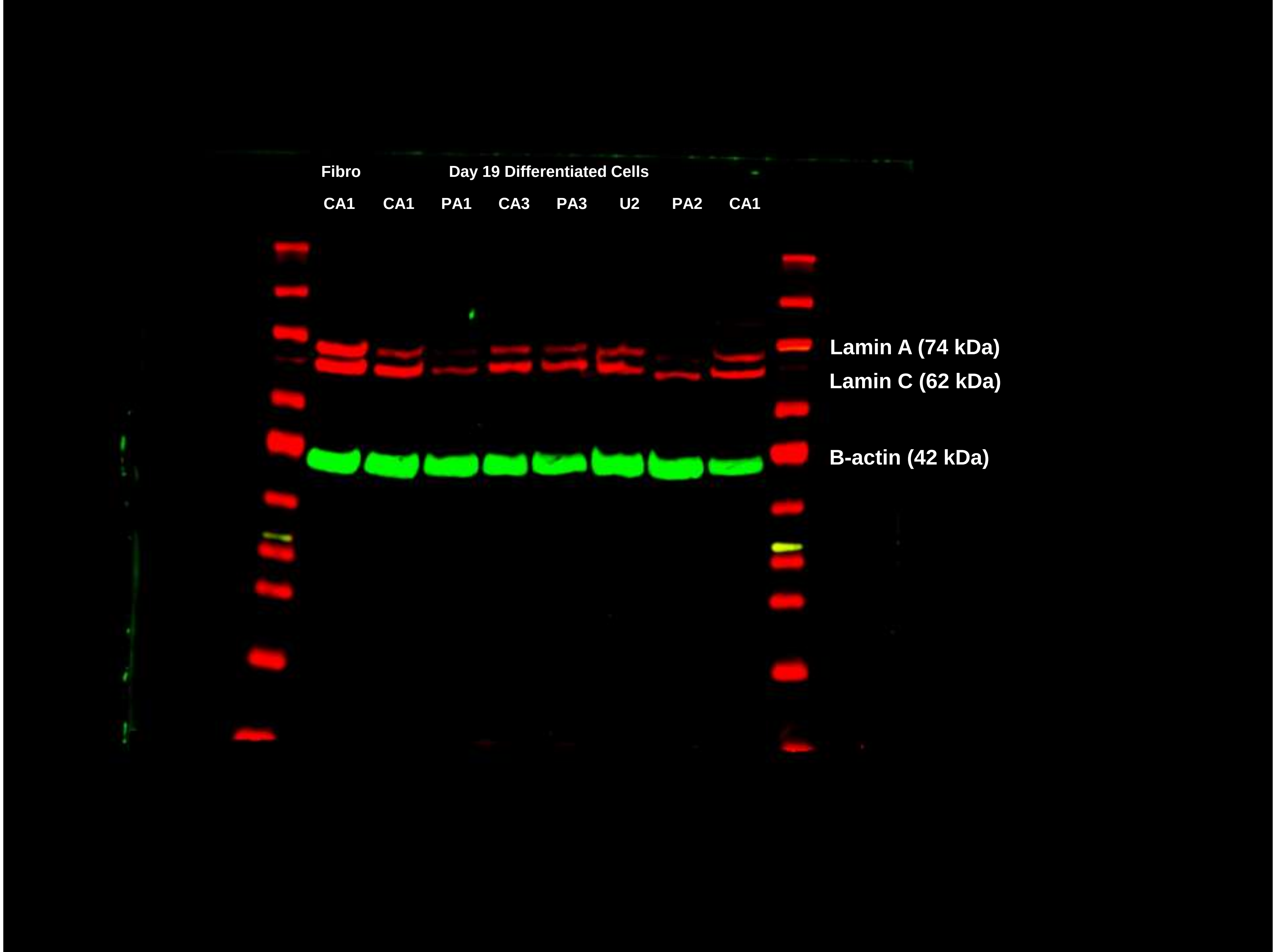

Fig. S14

#### Blot 3

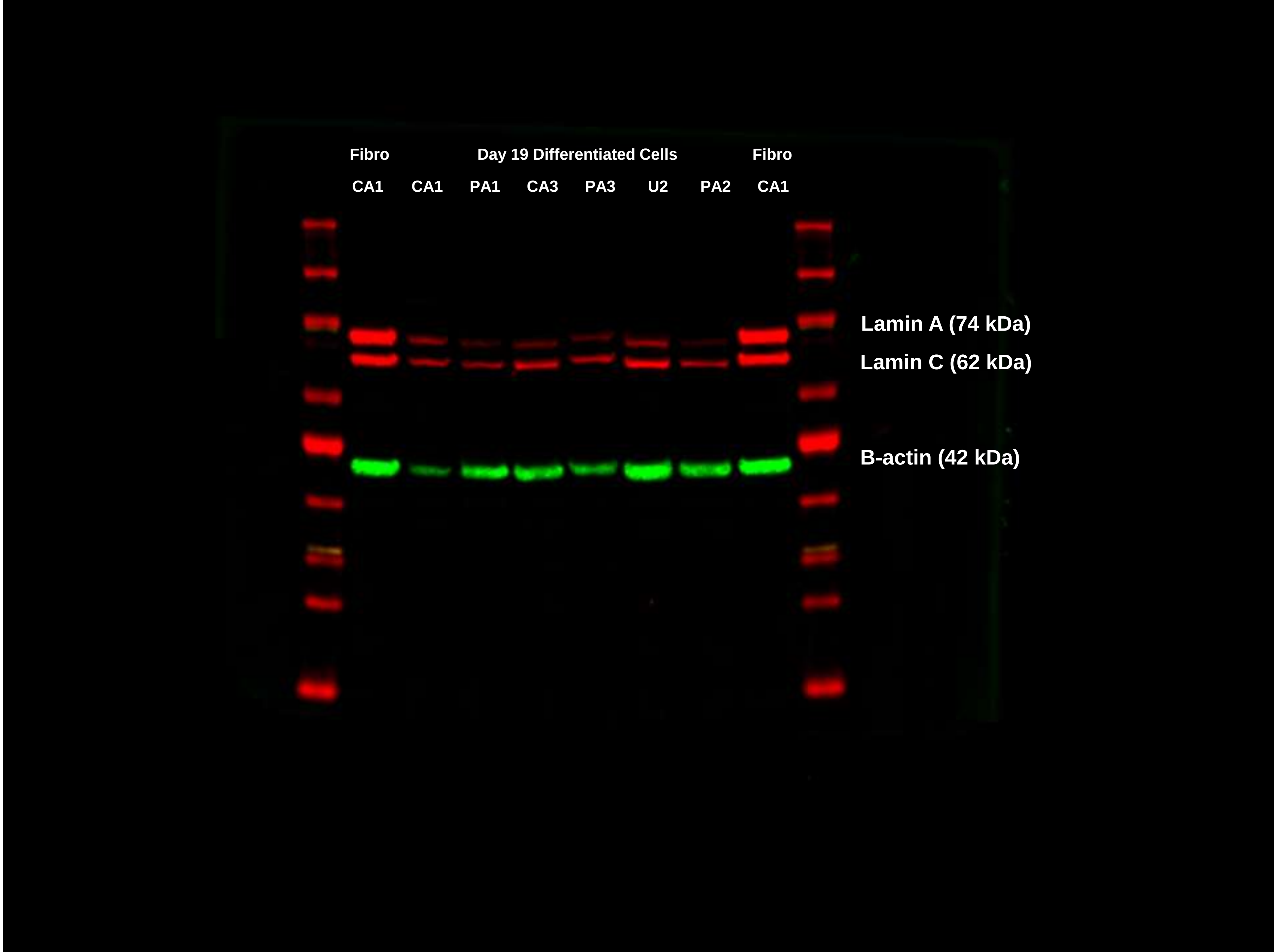
