## Supplemental Tables for "*LMNA*-Related Dilated Cardiomyopathy: Single-Cell Transcriptomics during Patient-derived iPSC Differentiation Support Cell type and Lineage-specific Dysregulation of Gene Expression and Development for Cardiomyocytes and Epicardium-Derived Cells with Lamin A/C Haploinsufficiency"


| | | Page |
|---|---|---|
| **Table S1.** | **Human subjects and results for fibroblast _LMNA_ sequencing and iPSC validation.** | **2** |
| **Table S2.** | **Methods: Primary and secondary antibodies.** | **3** |
| **Table S3.** | **Methods: Main software packages.** | **4** |
| **Table S4.** | **scRNA-seq Summary Metrics using Cell Ranger.** | **5** |
| **Table S5.** | **Single Sample Data: QC data processing using SoupX, Seurat, and DoubletFinder.** | **6** |
| **Table S6.** | **Single Sample and Combined Data: QC processing of Raw to Singlet Data Matrices.** | **7** |
| **Table S7.** | **Single Sample Data: Individual and Subcluster Analyses for Cell Annotation.** | **8 - 10** |
| **Table S8.** | **Gene marker panels for cell annotation.** | **11 - 12** |
| **Table S9.** | **Paired Sample Data: Individual and Subcluster Analyses.** | **13 - 14** |
| **Table S10.** | **Single Subset Data: Trajectory Inference, Lineage DEG, and Enrichment.** | **15** |
| **Table S11.** | **Paired Subset Data: Integration, Trajectory Inference, Lineage DEG, and Enrichment.** | **16** |
| **Table S12.** | **Lamin A/C Western Blot Quantification Data and Statistical Analyses.** | **17 - 19** |
| | **References:** | **20 - 22** |

**Table S1. Human subjects and results for fibroblast *LMNA* sequencing and iPSC validation.**

| ***Human Subjects*** | | | ***Fibroblast*** | | ***iPSC*** | | |
|---|---|---|---|---|---|---|---|
| **Identification[a]** | **Sex** | **Age At Bx** | ***LMNA* Mutation Genotype[b]** | ***LMNA* Coding SNV[c]** | **Clone** | **Karyotype (Passage #)[d]** | **ICC-DIFF[e]** |
| **CONTROLS** | | | | | | | |
| Control A1 (CA1): IV-2 | F | 49 | +/+ | None | CA1-A | 46,XX (P13) | Validated |
| | | | | | CA1-B | 46,XX (P13) | Validated |
| Control A2 (CA2): III-2 | M | 69 | +/+ | None | CA2[f] | 46,XY (P09) | Validated |
| Control A3 (CA3): III-3 | F | 68 | +/+ | None | CA3 | 46,XX (P13) | Validated |
| Unrelated Control (U2)[f] | M | 51 | +/+ | rs538089 Exon 5: T>C<br>rs505058 Exon 7: T>C | U2 | 46,XY (P12) | Validated |
| **PATIENTS** | | | | | | | |
| Patient A1 (PA1): IV-5 | F | 38 | -/+ | rs4641 Exon 10: C>T | PA1 | 46,XX (P19) | Validated |
| Patient A2 (PA2): III-5 | M | 62 | -/+ | rs4641 Exon 10: C>T | PA2 | 46,XY (P11) | Validated |
| Patient A3 (PA3): III-1 | F | 70 | -/+ | rs4641 Exon 10: C>T | PA3 | 46,XX (P10) | Validated |

[a]Identification: Study family pedigree [1]
[b]*LMNA* Genotype: +/+ homozygous normal allele; +/- heterozygous *LMNA* c.357-2A>G mutation
[c]*LMNA* Coding SNV: detected in fibroblast gDNA and cDNA [1] and used for allelic expression.
[d]Karyotype: G-banded metaphase chromosome constitution [2]
[e]ICC-DIFF: Differentiation capability by immunocytochemistry staining for germ layer markers in embryoid bodies (EBs).
[f]CA2 had poor viability and failed CM differentiation; therefore, U2 was used instead.
Abbreviations: iPSC, induced pluripotent stem cell; Bx, biopsy; SNV, single nucleotide variant; F, female; M, male; P, Passage

**Table S2. Methods: Primary and secondary antibodies**.

| Antigen (host) | Company, Catalog No. | Dilution | Antigen (host) | Company, Catalog No. | Dilution | Assay |
|---|---|---|---|---|---|---|
| **Primary antibodies** | | | **Secondary antibodies** | | | |
| AFP/alpha-fetoprotein (mouse) | Thermo Fisher Sci, A25530* | 1:500 | Anti-mouse (goat) AF-488 | Thermo Fisher Sci, A25536* | 1:250 | ICC-DIFF ENDO |
| FOXA2/Forkhead Box A2 (mouse) | Thermo Fisher Sci, BDB561580 | 1:100 | Anti-mouse (goat) AF-488 | Abcam, ab150113 | 1:250 | ICC-DIFF ENDO |
| SMA/smooth muscle actin (mouse) | Thermo Fisher Sci, A25531* | 1:100 | Anti-mouse (goat) AF-594 | Thermo Fisher Sci, A25534* | 1:250 | ICC-DIFF MESO |
| Beta-III tubulin (rabbit) | Thermo Fisher Sci, A25532* | 1:500 | Anti-rabbit (donkey) AF-647 | Thermo Fisher Sci, A25537* | 1:250 | ICC-DIFF ECTO |
| Lamin A/C N-term (E-1) monoclonal (mouse) | SC Biotech, sc-376248 | 1:500 | Anti-mouse (goat) AF-680 | Thermo Fisher Sci, A21057 | 1:2000 | WB |
| Beta-Actin (13E5) monoclonal (rabbit) | Cell Signaling, 4970S | 1:1000 | Anti-rabbit (goat) AF-790 | Thermo Fisher Sci, A11369 | 1:2000 | WB-control |

*3-Germ Layer Immunocytochemistry Kit (A25538, Life Technologies)

Abbreviations: AF, Alexa Fluor; ICC-DIFF, differentiation capability by immunocytochemistry; ENDO, endoderm; MESO, mesoderm; ECTO, ectoderm

**Table S3. Methods: Main software packages.**

| Name | Version | Source | Reference |
|---|---|---|---|
| AzureSpot | 2.1 | azurebiosystems.com | |
| bcl2fastq2 | 2.20 | support.illumina.com | |
| BioRender | | www.biorender.com | |
| Cell Ranger | 3.0.2 | www.10xgenomics.com | [3] |
| Cluster Profiler | 4.6.2 | bioconductor.org | [4] |
| Condiments | 1.6 | bioconductor.org | [5] |
| DoubletFinder | 2.0 | github.com | [6] |
| EnhancedVolcano | 1.16.0 | github.com | [7] |
| Integrative Genomics Viewer | 2.9.2 | igv.org | [8] |
| Molecular Signatures Database R | 7.5.1 | cran.r-project.org | [9] |
| Pretty heatmaps (pheatmap) | 1.0.12 | cran.r-project.org | |
| R | 4.2.2 | www.r-project.org | |
| RStudio | 2023.06.1+524 | posit.co | |
| Seurat | 4.3.0 | cran.r-project.org | [10, 11] |
| Slingshot | 2.6.0 | bioconductor.org | [12] |
| SoupX | 1.6.2 | cran.r-project.org | [13] |
| Tidyverse includes ggplot2 | 1.3.2 | cran.r-project.org | |
| TradeSeq | 1.12.0 | bioconductor.org | [14] |
| VennDiagram | 1.7.3 | cran.r-project.org | [15] |

**Table S4. scRNA-seq Summary Metrics using Cell Ranger [3].**

| Sample | Set | Shallow Sequencing | | | Deep Sequencing | | | |
|---|---|---|---|---|---|---|---|---|
| | | Estimated No. Cells Captured | Mean Reads Per Cell | Median Genes Per Cell | Estimated No. Cells Captured | Mean Reads Per Cell | Median Genes Per Cell | % Reads Mapped Confidently to Genome |
| Control A1 D00 | B | 11,825 | 3,072 | 820 | 14,080 | 40,657 | 4,482 | 93.0 |
| Control A1 D02 | A | 5,684 | 6,015 | 1,298 | 6,360 | 44,140 | 4,396 | 94.2 |
| Control A1 D04 | A | 4,899 | 5,724 | 1,166 | 5,570 | 36,259 | 3,698 | 93.3 |
| Control A1 D09 | A | 6,517 | 5,913 | 1,057 | 7,306 | 47,307 | 3,908 | 93.6 |
| Control A1 D09 | B | 10,311 | 4,200 | 853 | 13,727 | 29,371 | 3,236 | 92.0 |
| Control A1 D16 | B | 8,054 | 7,580 | 1,383 | 9,822 | 46,790 | 4,068 | 92.9 |
| Control A1 D19 | B | 10,077 | 5,571 | 1,076 | 12,601 | 41,416 | 3,550 | 93.9 |
| Control A1 D30 | A | 4,195 | 9,957 | 1,502 | 4,631 | 48,000 | 3,682 | 94.0 |
| Patient A1 D00 | B | 10,764 | 3,386 | 856 | 13,471 | 39,451 | 4,410 | 93.1 |
| Patient A1 D09 | B | 10,367 | 4,441 | 846 | 14,116 | 30,951 | 3,201 | 91.6 |
| Patient A1 D16 | B | 8,806 | 6,519 | 1,269 | 10,836 | 43,074 | 4,040 | 92.9 |
| Patient A1 D19 | B | 10,401 | 5,975 | 1,070 | 13,034 | 44,325 | 3,588 | 93.6 |
| All Samples (n=12) | AB | 101,900 | 5,696 | | 125,554 | 40,978 | | 93.2 |
| Control Samples (n=8) | AB | 60,501 | 6,043 | | 74,097 | 41,743 | | 93.4 |
| Patient Samples (n=4) | B | 41,399 | 5,002 | | 51,457 | 39,450 | | 92.8 |
| Paired Samples (n=8) | B | 80,605 | 5,093 | | 101,687 | 39,504 | | 92.9 |

**Table S5. Single Sample Data: QC data processing using SoupX, Seurat, and DoubletFinder.**

| Single Sample Data (n=12) | Set | SoupX[a] | | | Seurat[b] | | | DoubletFinder[c] | | |
|---|---|---|---|---|---|---|---|---|---|---|
| | | Clustering Dims and Resolution | Method: Contamination Fraction (rho) | Non-expressed Genes for rho Calculation | nFeature | nCount | %Mt | pK | Exp | nExp, nExp-adj |
| Control A1 D00 | B | 1:30, 0.15 | Auto: 0.01[a]<br>Manual: Not done | N/A | < 500 | > 40000 | N/A | 0.23 | 0.10 | 1318, 489 |
| Control A1 D02 | A | 1:30, 0.25 | Auto: 0.16<br>Manual: 0.45 | *SOX17, FOXA2* | < 900 | > 30000 | N/A | 0.04 | 0.05 | 285, 206 |
| Control A1 D04 | A | 1:30, 0.25 | Auto: 0.03<br>Manual: 0.37 | *SOX17, FOXA2* | < 900 | > 30000 | N/A | 0.04 | 0.05 | 241, 178 |
| Control A1 D09 | A | 1:30, 0.25 | Auto: 0.04<br>Manual: 0.13 | *TTN, MYL7, MYL2* | < 300 | > 60000 | N/A | 0.04 | 0.06 | 423, 359 |
| Control A1 D09 | B | 1:30, 0.25 | Auto: 0.12<br>Manual: 0.33 | *TTN, MYL7, MYL2* | < 100 | > 30000 | N/A | 0.21 | 0.10 | 1312, 1073 |
| Control A1 D16 | B | 1:30, 0.15 | Auto: 0.04<br>Manual: 0.53 | *TTN, MYL7, MYL2* | < 100 | > 30000 | N/A | 0.17 | 0.08 | 740, 537 |
| Control A1 D19 | B | 1:30, 0.05 | Auto: 0.04<br>Manual: 0.47 | *TTN, MYL7, MYL2* | < 100 | > 40000 | N/A | 0.16 | 0.10 | 1140, 784 |
| Control A1 D30 | A | 1:30, 0.05 | Auto 0.05<br>Manual: 0.19 | *TTN, MYL7, MYL2* | < 200 | > 70000 | N/A | 0.30 | 0.04 | 182, 134 |
| Patient A1 D00 | B | 1:30, 0.15 | Auto: 0.13<br>Manual: Not done | N/A | < 500 | > 40000 | N/A | 0.16 | 0.10 | 1252, 495 |
| Patient A1 D09 | B | 1:30, 0.25 | Auto: 0.01<br>Manual: 0.30 | *TTN, MYL7, MYL2* | < 100 | > 30000 | N/A | 0.17 | 0.10 | 1403, 1127 |
| Patient A1 D16 | B | 1:30, 0.20 | Auto: 0.06<br>Manual: 0.38 | *TTN, MYL7, MYL2* | < 100 | > 30000 | N/A | 0.10 | 0.08 | 823, 611 |
| Patient A1 D19 | B | 1:30, 0.07 | Auto: 0.04<br>Manual: 0.42 | *TTN, MYL7, MYL2* | < 100 | > 40000 | N/A | 0.005 | 0.10 | 1136, 768 |

[a]SoupX [13]: manually calculated rho values were used except for Day 0 due to lack of negative gene markers. For Day 0, automatically estimated global rho values were used. For CA1, this value was low (0.01); therefore, the greater rho value (0.13) estimated for PA1 was used instead.
[b]Seurat [10, 11]: QC covariate thresholds used to identify low-quality cells.
High %Mt thresholds were used after clustering/annotation for non-CM cells.
[c]DoubletFinder [6]: values for pK were obtained with 'paramSweep_v3' (PCs = 1:30). Expected proportion of doublets (Exp) was obtained from the 10x Chromium User Guide. Expected number of doublets (nExp) was adjusted (nExp-adj) using an estimated proportion of homotypic doublets.
Abbreviations: D, day; dims, dimensions; Auto, automatic; N/A, Not Applicable; Mt, Mitochondrial DNA genes; CM, cardiomyocyte

**Table S6. Single Sample and Combined Data: QC processing of Raw to Singlet Data Matrices.**

| Single Sample Data (n=12) | Set | Total Number of Cells in Processed Data (gene-barcode matrix) | | | | | |
|---|---|---|---|---|---|---|---|
| | | Raw[a] | Corrected[b] (%Raw) | Filtered: LQ Removed[c] (%Corrected) | Filtered: HQ Retained[c] (%Corrected) | Doublet Cells Removed[d] (%Filtered-HQ) | Singlet Cells Retained[d] (%Filtered-HQ) |
| Control A1 D00 | B | 14,080 | 14,080 (100%) | 901 (6%) | 13,179 (94%) | 489 (4%) | 12,690 (96%) |
| Control A1 D02 | A | 6,360 | 6,360 (100%) | 664 (10%) | 5,696 (90%) | 206 (4%) | 5,490 (96%) |
| Control A1 D04 | A | 5,570 | 5,570 (100%) | 741 (13%) | 4,829 (87%) | 178 (4%) | 4,651 (96%) |
| Control A1 D09 | A | 7,306 | 7,306 (100%) | 253 (3%) | 7,053 (97%) | 359 (5%) | 6,694 (95%) |
| Control A1 D09 | B | 13,727 | 13,727 (100%) | 600 (4%) | 13,127 (96%) | 1,073 (8%) | 12,054 (92%) |
| Control A1 D16 | B | 9,822 | 9,822 (100%) | 573 (6%) | 9,249 (94%) | 537 (6%) | 8,712 (94%) |
| Control A1 D19 | B | 12,601 | 12,601 (100%) | 1,201 (10%) | 11,400 (90%) | 784 (7%) | 10,616 (93%) |
| Control A1 D30 | A | 4,631 | 4,631 (100%) | 80 (2%) | 4,551 (98%) | 134 (3%) | 4,417 (97%) |
| Patient A1 D00 | B | 13,471 | 13,471 (100%) | 953 (7%) | 12,518 (93%) | 495 (4%) | 12,023 (96%) |
| Patient A1 D09 | B | 14,116 | 14,116 (100%) | 90 (1%) | 14,026 (99%) | 1,127 (8%) | 12,899 (92%) |
| Patient A1 D16 | B | 10,836 | 10,836 (100%) | 547 (5%) | 10,289 (95%) | 611 (6%) | 9,678 (94%) |
| Patient A1 D19 | B | 13,034 | 13,034 (100%) | 1,669 (13%) | 11,365 (87%) | 768 (7%) | 10,597 (93%) |
| **Combined Data** | | | | | | | |
| All 12 Samples- Merged | AB | 125,554 | 125,554 (100%) | 8,272 (7%) | 117,282 (93%) | 6,761 (6%) | 110,521 (94%) |
| Eight Controls- Merged | AB | 74,097 | 74,097 (100%) | 5,013 (7%) | 69,084 (93%) | 3,760 (5%) | 65,324 (95%) |
| Four Patients- Merged | B | 51,457 | 51,457 (100%) | 3,259 (6%) | 48,198 (94%) | 3,001 (6%) | 45,197 (94%) |
| Eight Paired- Integrated | B | 101,687 | 101,687 (100%) | 6,534 (6%) | 95,153 (94%) | 5,884 (6%) | 89,269 (94%) |

[a]Raw Data Matrix: total cells from Cell Ranger filtered gene-barcode matrix

[b]Corrected Data Matrix: total cells from SoupX corrected gene-barcode matrix

[c]Filtered Data Matrix: total high-quality cells after low-quality cell removal using QC covariate thresholds for nFeature and nCounts in Seurat

[d]Singlet Data Matrix: total DoubletFinder predicted singlet cells after removal of predicted heterotypic doublet cells

### Table S7. Single Sample Data: Individual and Subcluster Analyses for Cell Annotation

| A. Individual Analyses of Singlet Data for Main Cell Types | | | | | | B. Subcluster Analyses of Subset Data for Possible Cell Subtypes | | | | | | | | | |
|---|---|---|---|---|---|---|---|---|---|---|---|---|---|---|---|
| Single Sample Data (Singlet): n=12 | Total Cells | Res | Vars.to regress | Main Cell Types: n=10 | Cells (%Total) | Subset Data: n=32 | QC Thresholds (LQ Cells) %Mt | nFeature | nCount | LQ Cells Removed (%Pre-filter) | Post-filter HQ Cells (%Pre-filter) | Res | Vars.to. regress | All Possible Subtypes: n=30* | Cells (%Post-filter) |
| Control A1 D00 | 12,690 | 0.15 | %Mt & CC | PP | 11,241 (89%) | A | >25% | N/A | N/A | 185 (2%) | 11,056 (98%) | 0.15 | CC | PP-A<br>PP-B<br>PP-C<br>PP-D | 9670 (88%)<br>318 (3%)<br>722 (6%)<br>346 (3%) |
| | | | | UNK | 1,449 (11%) | B | N/A | N/A | N/A | N/A | 1,449 (100%) | 0.15 | CC | UNK-A<br>UNK-B<br>UNK-C | 566 (39%)<br>679 (47%)<br>204 (14%) |
| Control A1 D02 | 5,490 | 0.25 | %Mt & CC | ME<br>CMESO<br>ENDO | 5,292 (96%) | A | >20% | N/A | N/A | 8 (0.2%) | 5,284 (99.8%) | 0.25 | CC | ME-A<br>ME-B<br>ME-C<br>CMESO<br>ENDO | 1,954 (37%)<br>185 (3%)<br>1,356 (26%)<br>1,609 (31%)<br>180 (3%) |
| | | | | UNK | 198 (4%) | B | N/A | N/A | N/A | N/A | 198 (100%) | 0.25 | CC | UNK-A1<br>UNK-A2<br>UNK-B | 90 (46%)<br>82 (41%)<br>26 (13%) |
| Control A1 D04 | 4,651 | 0.25 | %Mt & CC | PP<br>CMESO<br>ENDO<br>CP | 4,572 (98%) | A | >20% | N/A | N/A | 40 (1%) | 4,532 (99%) | 0.25 | CC | CMESO<br>CP<br>ENDO-A<br>ENDO-B<br>PP | 1,175 (26%)<br>1,054 (23%)<br>1,515 (33.5%)<br>23 (0.5%)<br>765 (17%) |
| | | | | UNK | 79 (2%) | B | N/A | N/A | N/A | N/A | 79 (100%) | 0.25 | CC | UNK-A | 79 (100%) |
| Control A1 D09A | 6,694 | 0.25 | %Mt & CC | CP<br>CM<br>EPDC | 4,568 (69%) | A | N/A | N/A | N/A | N/A | 4,568 (100%) | 0.25 | CC | CP-A<br>CM-A<br>CM/UNK-B<br>CP-B<br>EPDC | 421 (9%)<br>1,250 (28%)<br>334 (7%)<br>1,090 (24%)<br>1,473 (32%) |
| | | | | ENDO<br>ENDOTH<br>UNK | 1,823 (27%) | B | >20% | N/A | N/A | 20 (1%) | 1,803 (99%) | 0.25 | CC | ENDO-A1<br>ENDO-A2<br>ENDO-A3<br>ENDO-A4<br>ENDO-B<br>ENDOTH<br>UNK-C1<br>UNK-C2 | 658 (37%)<br>510 (28%)<br>238 (13%)<br>178 (10%)<br>42 (2%)<br>98 (5%)<br>62 (3%)<br>17 (1%) |
| | | | | UNK | 303 (4%) | C | N/A | N/A | N/A | N/A | 303 (100%) | 0.25 | CC | UNK-A | 303 (100%) |
| Control A1 D09B | 12,054 | 0.15 | %Mt & CC | CP<br>CM<br>EPDC | 10,612 (88%) | A | N/A | N/A | N/A | N/A | 10,612 (100%) | 0.25 | CC | CP-A<br>CM-A<br>CP-B<br>EPDC<br>CM/UNK-B1<br>CM/UNK-B2 | 3,077 (29%)<br>2,216 (21%)<br>396 (4%)<br>3,033 (28%)<br>1,258 (12%)<br>632 (6%) |
| | | | | ENDO<br>ENDOTH<br>ECTO<br>UNK | 532 (5%) | B | >20% | N/A | N/A | 13 (2%) | 519 (98%) | 0.25 | CC | ECTO-A<br>ECTO-B<br>ENDOTH<br>ENDO | 159 (31%)<br>131 (25%)<br>97 (19%)<br>87 (17%) |

| | | | | | | | | | | | | | | | |
|---|---|---|---|---|---|---|---|---|---|---|---|---|---|---|---|
| | | | | | | | | | | | | | | UNK-C | 45 (8%) |
| | | | | UNK | 910 (7%) | C | N/A | N/A | N/A | N/A | 910 (100%) | 0.25 | CC | UNK-A1<br>UNK-A2<br>UNK-A3 | 700 (77%)<br>118 (13%)<br>92 (10%) |
| Control A1 D16 | 8,712 | 0.15 | %Mt & CC | CM | 6,508 (75%) | A | N/A | N/A | N/A | N/A | 6,508 (100%) | 0.15 | CC | CM-A1<br>CM-A2<br>CM-A3<br>CM/UNK-B | 4,656 (72%)<br>240 (4%)<br>218 (3%)<br>1,394 (21%) |
| | | | | EPDC | 1,172 (13%) | B | >15% | <1500 | >15000 | 154 (13%) | 1,018 (87%) | 0.15 | CC | EPDC-A<br>EPDC-B | 867 (85%)<br>151 (15%) |
| | | | | UNK | 1,032 (12%) | C | N/A | N/A | N/A | N/A | 1,032 (100%) | 0.15 | CC | UNK-A1<br>UNK-A2<br>UNK-B | 522 (51%)<br>328 (32%)<br>182 (17%) |
| Control A1 D19 | 10,616 | 0.05 | %Mt & CC | CM | 7,831 (74%) | A | N/A | N/A | N/A | N/A | 7,831 (100%) | 0.10 | CC | CM-A1<br>CM-A2<br>CM/UNK-B | 4,914 (63%)<br>526 (7%)<br>2,391 (30%) |
| | | | | EPDC | 1,359 (13%) | B | >15% | <1500 | >30000 | 234 (17%) | 1,125 (83%) | 0.10 | CC | EPDC | 1,125 (100%) |
| | | | | UNK | 1,426 (13%) | C | N/A | N/A | N/A | N/A | 1,426 (100%) | 0.10 | CC | UNK-A<br>UNK-B | 904 (63%)<br>522 (37%) |
| Control A1 D30 | 4,417 | 0.05 | %Mt & CC | CM | 1,675 (38%) | A | N/A | N/A | N/A | N/A | 1,675 (100%) | 0.15 | CC | VENTR-CM<br>ATRIAL-CM | 1,070 (64%)<br>605 (36%) |
| | | | | EPDC | 1,960 (44%) | B | >15% | <1500 | >40000 | 243 (12%) | 1,717 (88%) | 0.15 | CC | CFIBRO<br>EPROG<br>EPDC-UNSP<br>VSM | 797 (46%)<br>460 (27%)<br>382 (22%)<br>78 (5%) |
| | | | | UNK | 782 (18%) | C | N/A | N/A | N/A | N/A | 782 (100%) | 0.15 | CC | UNK-A<br>UNK-B<br>UNK-C | 444 (57%)<br>245 (31%)<br>93 (12%) |
| Patient A1 D00 | 12,023 | 0.15 | %Mt & CC | PP | 9,878 (82%) | A | >25% | N/A | N/A | 118 (1%) | 9,760 (99%) | 0.15 | CC | PP-A<br>PP-B<br>PP-C<br>PP-D | 8,976 (92%)<br>382 (4%)<br>338 (3%)<br>64 (1%) |
| | | | | UNK | 2,145 (18%) | B | N/A | N/A | N/A | N/A | 2,145 (100%) | 0.15 | CC | UNK-A<br>UNK-B<br>UNK-C | 764 (36%)<br>1,004 (47%)<br>377 (17%) |
| Patient A1 D09B | 12,899 | 0.25 | %Mt & CC | CP<br>CM<br>EPDC | 10,588 (82%) | A | N/A | N/A | N/A | N/A | 10,588 (100%) | 0.25 | CC | CP-A<br>CM-A<br>CP-B<br>CP-C<br>EPDC<br>CM/UNK-B | 819 (8%)<br>3,272 (31%)<br>2,160 (20%)<br>801 (8%)<br>1,314 (12%)<br>2,222 (21%) |
| | | | | ENDO<br>ENDOTH<br>ECTO<br>UNK | 1,066 (8%) | B | >20% | N/A | N/A | N/A | 1,012 (95%) | 0.25 | CC | ECTO-A<br>ECTO-B<br>ECTO-C<br>ECTO-D<br>ENDO<br>ENDOTH<br>UNK-A | 245 (24%)<br>247 (24%)<br>159 (16%)<br>188 (19%)<br>85 (8%)<br>31 (3%)<br>57 (6%) |

| | | | | | | | | | | | | | | | |
|---|---|---|---|---|---|---|---|---|---|---|---|---|---|---|---|
| | | | | UNK | 1,245 (10%) | C | N/A | N/A | N/A | N/A | 1,245 (100%) | 0.25 | CC | UNK-A1<br>UNK-A2<br>UNK-A3<br>UNK-C | 746 (60%)<br>173 (14%)<br>47 (4%)<br>279 (22%) |
| | | | | CM | 6,192 (64%) | A | N/A | N/A | N/A | N/A | 6,192 (100%) | 0.20 | CC | CM-A1<br>CM-A2<br>CM-A3<br>CM-A4<br>CM/UNK-B<br>UNK-B | 3,425 (55%)<br>1,222 (20%)<br>216 (3.5%)<br>117 (2%)<br>991 (16%)<br>221 (3.5%) |
| Patient A1 D16 | 9,678 | 0.20 | %Mt & CC | EPDC | 2,175 (22%) | B | >15% | <1500 | >20000 | 60 (3%) | 2,115 (97%) | 0.20 | CC | EPDC-A1<br>EPDC-A2<br>EPDC-B<br>EPDC-C | 1,040 (49%)<br>889 (42%)<br>141 (7%)<br>45 (2%) |
| | | | | UNK | 1,311 (14%) | C | N/A | N/A | N/A | N/A | 1,311 (100%) | 0.20 | CC | UNK-A1<br>UNK-A2<br>UNK-C1<br>UNK-C2<br>UNK-C3<br>UNK-C4 | 643 (49%)<br>284 (22%)<br>219 (17%)<br>95 (7%)<br>37 (3%)<br>33 (2%) |
| | | | | CM | 7,392 (70%) | A | N/A | N/A | N/A | N/A | 7,392 (100%) | 0.10 | CC | CM-A<br>CM/UNK-B<br>UNK-B | 5,129 (69%)<br>2,201 (30%)<br>62 (1%) |
| Patient A1 D19 | 10,597 | 0.07 | %Mt & CC | EPDC | 1,164 (11%) | B | >15% | <2000 | >40000 | 31 (3%) | 1,133 (97%) | 0.10 | CC | EPDC | 1,133 (100%) |
| | | | | UNK | 2,041 (19%) | C | N/A | N/A | N/A | N/A | 2,041 (100%) | 0.10 | CC | UNK-A1<br>UNK-A2<br>UNK-B<br>UNK-C | 1,326 (65%)<br>164 (8%)<br>382 (19%)<br>169 (8%) |

*Possible Cell subtypes (n=18 of 30 total, highlighted in blue) re-subsetted, integrated, and used for trajectory inference.

Abbreviations: D, day; Res, resolution; Mt, MtDNA genes; CC, cell cycle; PP, pluripotent; UNK, unknown; CM, cardiomyocyte; EPDC, Epicardium-derived cells; N/A, Not Applicable; LQ, low-quality; ME, Mesendoderm; CMESO, Cardiogenic Mesoderm; ENDO, Endoderm; CP, Cardiac Progenitors; ENDOTH, Endothelium; ECTO, Ectoderm; VENTR-CM, Ventricular-Cardiomyocytes; ATRIAL-CM, Atrial Cardiomyocytes; CFIBRO, Cardiac Fibroblasts; EPROG, Epicardial Progenitors; EPDC-UNSP, EPDC-Unspecified; VSM, Vascular Smooth Muscle

**Table S8. Gene marker panels for cluster annotation.**

| Panel | No. | Gene Names | References |
|---|---|---|---|
| **Primary Marker Panel** | 25 | *POU5F1, SOX2, NANOG, CNMD, EOMES, MESP1, SOX17, FOXA2, PAX6, EGFL7, HAND1, HAPLN1, TMEM88, MYL7, TNNI1, NKX2-5, TTN, MYH7, MYL2, TBX18, TCF21, WT1, COL3A1, LUM, FBN1* | |
| Pluripotent cells (PP) | 3 | *POU5F1, SOX2, NANOG* | [16, 17] |
| Undifferentiated cells | 1 | *CNMD* | [18] |
| Early Differentiated cells | 6 | Mesendoderm: *EOMES*<br>Cardiogenic Mesoderm: *MESP1*<br>Endoderm: *SOX17, FOXA2*<br>Ectoderm: *PAX6*<br>Endothelium: *EGFL7* | [19-22] |
| Cardiac Progenitors (CP) | 3 | *HAND1, HAPLN1, TMEM88* | [20, 23] |
| Cardiomyocyte (CM) | 6 | Early CM: *MYL7, TNNI1, NKX2-5*<br>CM: *TTN, MYH7, MYL2* | [16, 19, 24, 25] |
| Epicardium-derived cell (EPDC) | 6 | Early EPDC/PEO: *TBX18, TCF21, WT1*<br>EPDC: *COL3A1, LUM, FBN1* | [19, 26, 27] |
| **Expanded Marker Panel** | 38 | *POU5F1, SOX2, NANOG, DNMT3B, NODAL, UTF1, LIN28A, LEFTY1, GDF3, SDC2, CNMD, EOMES, MIXL1, TBXT, SOX17, FOXA2, AFP, FOXA3, HHEX, IHH, APOA1, GATA6, HAND1, HAPLN1, TMEM88, GATA4, HCN4, TBX5, ISL1, TBX1, HAND2, PAX6, SOX1, EGFL7, SELE, PECAM1, VWF, CD34* | |
| Pluripotent cells | 10 | *POU5F1, SOX2, NANOG, DNMT3B, NODAL, UTF1, LIN28A, LEFTY1, GDF3, SDC2* | [17] |
| Undifferentiated cells | 1 | *CNMD* | [18] |

| | | | |
|---|---|---|---|
| Mesendoderm (ME) | 3 | *EOMES*, *MIXL1, TBXT* | [20] |
| Endoderm (ENDO) | 8 | *SOX17, FOXA2*, *AFP, FOXA3, HHEX, IHH, APOA1, GATA6* | [21, 28, 29] |
| Cardiac Progenitor (CP) | 9 | *HAND1, HAPLN1, TMEM88, GATA4, HCN4, TBX5, ISL1, TBX1, HAND2* | [19, 20, 23] |
| Ectoderm (ECTO) | 2 | *PAX6, SOX1* | [21] |
| Endothelium (ENDOTH) | 5 | *EGFL7*, *SELE, PECAM1, VWF, CD34* | [27] |
| **Cardiomyocyte Subtype Marker Panel** | 27 | *NKX2-5, TBX5, NPPA, GATA4, NR2F2, MYH6, MYL7, MYL4, PITX2, HAND1, IRX4, IRX5, MYL2, MYH7, MYL3, ACTN2, VDR, HAND2, ISL1, MYH3, TBX3, HCN4, IRX3, ID2, GJC1, GJD3, TBX18* | [21, 30, 31] |
| Cardiomyocyte-shared | 3 | *NKX2-5, TBX5, NPPA* | |
| Atrial CM (ATR-CM) | 6 | *GATA4, NR2F2, MYH6, MYL7, MYL4, PITX2,* | |
| Ventricular CM (VENTR-CM) | 11 | *HAND1, IRX4, IRX5, MYL2, MYH7, MYL3, ACTN2, VDR, HAND2, ISL1, MYH3* | |
| Nodal-shared | 2 | *TBX3, HCN4* | |
| Atrioventricular node (AVN) | 4 | *IRX3, ID2, GJC1, GJD3* | |
| Sinoatrial node (SAN) | 1 | *TBX18* | |
| **EPDC Subtype Marker Panel** | 27 | *TBX18, TCF21, WT1, TBX5, ETS1, VCAM1, BVES, GJA1, ALDH1A2, ITGA4, PDGFRB, PLAU, ISL1, ITLN1, EFEMP1, UPK3B, PDGFRA, VIM, SNAI2, THY1, S100A4, DDR2, PECAM1, ACTA2, SRF, KDR, NFATC1* | [21, 26, 32] |
| PEO/EPI-shared | 6 | *TBX18, TCF21, WT1, TBX5, ETS1, VCAM1* | |
| Proepicardial Progenitor (PEO) | 2 | *BVES, GJA1* | |
| Epicardial Progenitor (EPI) | 8 | *ALDH1A2, ITGA4, PDGFRB, PLAU, ISL1, ITLN1, EFEMP1, UPK3B* | |
| Mesenchymal/ EPDC-shared | 3 | *PDGFRA, VIM, SNAI2* | |
| Cardiac Fibroblast (CFIBRO) | 4 | *THY1, S100A4, DDR2, PECAM1* | |
| Vascular Smooth Muscle (VSM) | 2 | *ACTA2, SRF* | |
| Angioblasts/ Endothelial cells | 2 | *KDR, NFATC1* | |

### Table S9. Paired Sample Data: Individual and Subcluster Analyses.

#### A. Individual Analyses of Paired Singlet Data for Shared Cell Types: 89,269 total cells

| Paired Sample Data (Singlet): n=4 pairs | Cells (%Total) | Integrated Data: n=4 | Total Cells | Res | Vars.to. regress | Shared Cell Types: n=10 | Cells (%Total) | CA1 Cells | PA1 Cells | CA1% | PA1% | Balanced Cell Type |
|---|---|---|---|---|---|---|---|---|---|---|---|---|
| CA1 D00<br>PA1 D00 | 12,690 (51%)<br>12,023 (49%) | CA1PA1 D00 | 24,713 | 0.15 | %Mt & CC | PP | 21,150 (86%) | 11,149 | 10,001 | 53% | 47% | No |
| | | | | | | UNK | 3,563 (14%) | 1,541 | 2,022 | 43% | 57% | No |
| CA1 D09B<br>PA1 D09B | 12,054 (48%)<br>12,899 (52%) | CA1PA1 D09 | 24,953 | 0.25 | %Mt & CC | CM | 7,419 (30%) | 3,878 | 3,541 | 52% | 48% | Yes |
| | | | | | | CP | 7,037 (28%) | 3,512 | 3,525 | 50% | 50% | Yes |
| | | | | | | EPDC | 6,449 (26%) | 2,998 | 3,451 | 46% | 54% | Yes |
| | | | | | | UNK | 2,567 (10%) | 1,179 | 1,388 | 46% | 54% | No |
| | | | | | | ECTO | 1,148 (4%) | 291 | 857 | 25% | 75% | No |
| | | | | | | ENDO | 197 (1%) | 94 | 103 | 48% | 52% | No |
| | | | | | | ENDOTH | 136 (1%) | 102 | 34 | 75% | 25% | Yes |
| CA1 D16<br>PA1 D16 | 8,712 (47%)<br>9,678 (53%) | CA1PA1 D16 | 18,390 | 0.15 | %Mt & CC | CM | 12,555 (68%) | 6,510 | 6,045 | 52% | 48% | Yes |
| | | | | | | EPDC | 3,419 (19%) | 1,149 | 2,270 | 34% | 66% | Yes |
| | | | | | | UNK | 2,416 (13%) | 1,053 | 1,363 | 44% | 56% | No |
| CA1 D19<br>PA1 D19 | 10,616 (50%)<br>10,597 (50%) | CA1PA1 D19 | 21,213 | 0.05 | %Mt & CC | CM | 15,188 (72%) | 7,828 | 7,360 | 52% | 48% | Yes |
| | | | | | | EPDC | 2,893 (13%) | 1,364 | 1,529 | 47% | 53% | Yes |
| | | | | | | UNK | 3,132 (15%) | 1,424 | 1,708 | 45% | 55% | No |

#### B. Subcluster Analyses of Paired Subset Data for Possible Shared Cell Subtypes: 88,420 total cells

| Paired Subset Data: n=11 pairs | Cells (%Total) | Integrated Subset Data: n=11* | Total Cells | Res | Vars.to. regress | Shared Subtypes: n=19* | Cells (%Total) | CA1 Cells | PA1 Cells | CA1% | PA1% | Balanced Subtype |
|---|---|---|---|---|---|---|---|---|---|---|---|---|
| CA1 D00 Subset-A (PP)<br>PA1 D00 Subset-A (PP) | 11,056 (53%)<br>9,760 (47%) | CA1PA1 D00 Subset-A (PP) | 20,816 | 0.10 | CC | PP-A1 | 17,677 (85%) | 9,333 | 8,344 | 53% | 47% | Yes |
| | | | | | | PP-A2 | 877 (4%) | 296 | 581 | 34% | 66% | Yes |
| | | | | | | PP-B | 716 (3%) | 326 | 390 | 46% | 54% | Yes |
| | | | | | | PP-C | 1,184 (6%) | 751 | 433 | 63% | 37% | No |
| | | | | | | PP-D | 362 (2%) | 350 | 12 | 97% | 3% | No |
| CA1 D00 Subset-B (UNK)<br>PA1 D00 Subset-B (UNK) | 1,449 (40%)<br>2,145 (60%) | CA1PA1 D00 Subset-B (UNK) | 3,594 | 0.15 | CC | UNK-A | 1,353 (38%) | 487 | 866 | 36% | 64% | No |
| | | | | | | UNK-B | 1,776 (49%) | 762 | 1,014 | 43% | 57% | No |
| | | | | | | UNK-C | 465 (13%) | 200 | 265 | 43% | 57% | No |
| CA1 D09B Subset-A<br>PA1 D09B Subset-A | 10,612 (50%)<br>10,588 (50%) | CA1PA1 D09 Subset-A (CP/CM /EPDC) | 21,200 | 0.25 | CC | CP-A | 5,946 (28%) | 3,053 | 2,893 | 51% | 49% | Yes |
| | | | | | | CP-B | 4,750 (23%) | 2,252 | 2,498 | 47% | 53% | Yes |
| | | | | | | CP-C | 1,216 (6%) | 437 | 779 | 36% | 64% | Yes |
| | | | | | | CM-A | 3,433 (16%) | 2,219 | 1,214 | 65% | 35% | Yes |
| | | | | | | CM/UNK-B1 | 2,191 (10%) | 1,244 | 947 | 57% | 43% | Yes |
| | | | | | | CM/UNK-B2 | 1,936 (9%) | 641 | 1,295 | 33% | 67% | Yes |
| | | | | | | EPDC | 1,728 (8%) | 766 | 962 | 44% | 56% | Yes |
| CA1 D09B Subset-B<br>PA1 D09B Subset-B | 519 (34%)<br>1,012 (66%) | CA1PA1 D09 Subset-B (ENDO/ ENDOTH/ ECTO/UNK) | 1,531 | 0.25 | CC | ECTO-A | 286 (19%) | 83 | 203 | 29% | 71% | Yes |
| | | | | | | ECTO-B | 289 (19%) | 88 | 201 | 30% | 70% | Yes |
| | | | | | | ECTO-C | 205 (13%) | 47 | 158 | 23% | 77% | No |
| | | | | | | ECTO-D | 191 (12%) | 13 | 178 | 7% | 93% | No |
| | | | | | | ENDO | 166 (11%) | 81 | 85 | 49% | 51% | No |
| | | | | | | ENDOTH | 132 (9%) | 101 | 31 | 77% | 23% | No |
| | | | | | | UNK-A | 93 (6%) | 19 | 74 | 20% | 80% | No |

| | | | | | | | | | | | | |
|---|---|---|---|---|---|---|---|---|---|---|---|---|
| | | | | | | UNK-C1<br>UNK-C2 | 116 (8%)<br>53 (3%) | 67<br>20 | 49<br>33 | 58%<br>38% | 42%<br>62% | No<br>No |
| CA1 D09B Subset-C (UNK)<br>PA1 D09B Subset-C (UNK) | 910 (42%)<br>1,245 (58%) | CA1PA1 D09 Subset-C (UNK) | 2,155 | 0.25 | CC | UNK-A1a<br>UNK-A1b<br>UNK-A2<br>UNK-A3<br>UNK-A4<br>UNK-C | 806 (37%)<br>623 (29%)<br>195 (9%)<br>165 (8%)<br>91 (4%)<br>275 (13%) | 145<br>505<br>100<br>98<br>44<br>18 | 661<br>118<br>95<br>67<br>47<br>257 | 18%<br>81%<br>51%<br>59%<br>48%<br>7% | 82%<br>19%<br>49%<br>41%<br>52%<br>93% | No<br>No<br>No<br>No<br>Yes<br>No |
| CA1 D16 Subset-A (CM)<br>PA1 D16 Subset-A (CM) | 6,508 (51%)<br>6,192 (49%) | CA1PA1 D16 Subset-A (CM) | 12,700 | 0.15 | CC | CM-A1<br>CM-A2<br>CM-A3<br>CM/UNK-B<br>UNK-B | 5,198 (41%)<br>3,441 (27%)<br>1,368 (11%)<br>2,472 (19%)<br>221 (2%) | 2,601<br>1,794<br>712<br>1,393<br>8 | 2,597<br>1,647<br>656<br>1,079<br>213 | 50%<br>52%<br>52%<br>56%<br>4% | 50%<br>48%<br>48%<br>44%<br>96% | Yes<br>Yes<br>No<br>Yes<br>No |
| CA1 D16 Subset-B (EPDC)<br>PA1 D16 Subset-B (EPDC) | 1,018 (32%)<br>2,115 (68%) | CA1PA1 D16 Subset-B (EPDC) | 3,133 | 0.15 | CC | EPDC-A1<br>EPDC-A2<br>EPDC-B<br>EPDC-C | 2,126 (68%)<br>723 (23%)<br>263 (8.5%)<br>21 (0.5%) | 672<br>227<br>98<br>21 | 1454<br>496<br>165<br>0 | 32%<br>31%<br>37%<br>100% | 68%<br>69%<br>63%<br>0% | No<br>No<br>No<br>No |
| CA1 D16 Subset-C (UNK)<br>PA1 D16 Subset-C (UNK) | 1,032 (44%)<br>1,311 (56%) | CA1PA1 D16 Subset-C (UNK) | 2,343 | 0.15 | CC | UNK-A1<br>UNK-A2<br>UNK-A3<br>UNK-A4<br>UNK-B<br>UNK-C1<br>UNK-C2 | 889 (38%)<br>569 (24%)<br>243 (10%)<br>84 (4%)<br>280 (12%)<br>216 (9%)<br>62 (3%) | 386<br>258<br>126<br>63<br>197<br>2<br>0 | 503<br>311<br>117<br>21<br>83<br>214<br>62 | 43%<br>45%<br>52%<br>75%<br>70%<br>1%<br>0% | 57%<br>55%<br>48%<br>25%<br>30%<br>99%<br>100% | No<br>Yes<br>No<br>No<br>No<br>No<br>No |
| CA1 D19 Subset-A (CM)<br>PA1 D19 Subset-A (CM) | 7,831 (51%)<br>7,392 (49%) | CA1PA1 D19 Subset-A (CM) | 15,223 | 0.10 | CC | CM-A1<br>CM-A2<br>CM/UNK-B<br>UNK-B | 9,978 (65%)<br>590 (4%)<br>4,591 (30%)<br>64 (0.4%) | 4,927<br>507<br>2,389<br>8 | 5,051<br>83<br>2,202<br>56 | 49%<br>86%<br>52%<br>13% | 51%<br>14%<br>48%<br>88% | Yes<br>No<br>Yes<br>No |
| CA1 D19 Subset-B (EPDC)<br>PA1 D19 Subset-B (EPDC) | 1,125 (50%)<br>1,133 (50%) | CA1PA1 D19 Subset-B (EPDC) | 2,258 | 0.10 | CC | EPDC | 2,258 (100%) | 1,125 | 1,133 | 50% | 50% | Yes |
| CA1 D19 Subset-C (UNK)<br>PA1 D19 Subset-C (UNK) | 1,426 (41%)<br>2,041 (59%) | CA1PA1 D19 Subset-C (UNK) | 3,467 | 0.10 | CC | UNK-A1<br>UNK-A2<br>UNK-B<br>UNK-C | 2,005 (58%)<br>360 (10%)<br>935 (27%)<br>167 (5%) | 740<br>154<br>531<br>1 | 1,265<br>206<br>404<br>166 | 37%<br>43%<br>57%<br>1% | 63%<br>57%<br>43%<br>99% | No<br>No<br>No<br>No |

*Integrated Subset Data with primarily ‘balanced” cell types (n=6) and ‘balanced’ cell subtypes (n=14): highlighted in yellow used for cell type analyses of differential expression.

Abbreviations: D, day; PP, pluripotent; UNK, unknown; CM, cardiomyocyte; EPDC, Epicardium-derived cells; Res, resolution; Mt, MtDNA genes; CC, cell cycle; CP, Cardiac Progenitors; ECTO, Ectoderm; ENDO, Endoderm; ENDOTH, Endothelium

**Table S10. Single Subset Data: Trajectory Inference, Lineage DEG, and Enrichment.**

| Single Subset Data (n=7) | Total Cells | Res | k | Lineage Topology using Slingshot [12] | Cell Lineage(s) | Lineage DEG[a] | | ORA GO BP GS[b] | |
|---|---|---|---|---|---|---|---|---|---|
| | | | | | | AT | SET | AT | SET |
| CA1 D00 PP-AB | 9,988 | 0.05 | 6 | Single Trajectory | PP-A → PP-B | 367 | 83 | 36 | 45 |
| CA1 D02 ME/CMESO/ENDO | 3,145 | 0.15 | 6 | Bifurcating Trajectory | 1. ME → CMESO<br>2. ME → ENDO | 907 | 469 | 27 | 187 |
| CA1 D04 CMESO/CP | 2,229 | 0.15 | 6 | Single Trajectory | CMESO → CP | 1,683 | 711 | 0 | 114 |
| CA1 D09A CP-A/CM-A | 1,671 | 0.05 | 6 | Single Trajectory | CP-A → CM-A | 1,835 | 1,003 | 139 | 239 |
| CA1 D09B CP-A/CM-A | 5,293 | 0.05 | 6 | Single Trajectory | CP-A → CM-A | 1,806 | 483 | 153 | 305 |
| PA1 D00 PP-AB | 9,358 | 0.05 | 6 | Single Trajectory | PP-A → PP-B | 301 | 99 | 174 | 246 |
| PA1 D09B CP-A/CM-A | 4,091 | 0.10 | 6 | Single Trajectory | CP-A → CM-A | 1,538 | 247 | 117 | 195 |

[a]Lineage DEG using TradeSeq [14]: Threshold = FDR <0.05 & Fold Change > 2x;
For bifurcating trajectories, global lineage DEG were identified across both lineages.
[b]ORA Parameters: DEG = top 100 DEG ranked by Wald statistic, minimum #DEG = 3, adjusted p-value < 0.05, q-value cutoff = 0.20
Abbreviations: PP, pluripotent; ME, mesendoderm; CMESO, cardiogenic mesoderm; ENDO, endoderm; CP, Cardiac Progenitors; CM, cardiomyocyte; Res, resolution; DEG, differentially expressed gene; AT, Association Test; SET, Start-End Test; ORA, Over-Representation Analysis; GO BP, Gene Ontology Biological Processes; GS, gene set

**Table S11. Paired Subset Data: Integration, Trajectory Inference, Lineage DEG and Enrichment.**

| Paired Subset Data (n=2) | Cells (%Total) | Integrated Subset Data (n=2) | Total Cells | Patient (%Total) | Control (%Total) | Res | Vars.to. regress | k* | Lineage Topology [12] | Cell Lineage(s) |
|---|---|---|---|---|---|---|---|---|---|---|
| CA1 D00 PP-AB<br>PA1 D00 PP-AB | 9,988 (52%)<br>9,358 (48%) | CA1PA1 D00 PP-AB | 19,346 | 9,988 (52%) | 9,358 (48%) | 0.05 | CC | 6 | Single-Trajectories | PP-A→PP-B |
| CA1 D09B CP-AB/CM-A/EPDC<br>CA1 D16 CM-A123<br>CA1 D16 EPDC-AB<br>CA1 D19 CM-A12<br>CA1 D19 EPDC<br>PA1 D09B CP-ABC/CM-A/EPDC<br>PA1 D16 CM-A1234<br>PA1 D16 EPDC-A12BC<br>PA1 D19 CM-A<br>PA1 D19 EPDC | 8,722 (20%)<br>5,114 (12%)<br>1,018 (2%)<br>5,440 (13%)<br>1,125 (3%)<br>8,366 (19%)<br>4,980 (11%)<br>2,115 (5%)<br>5,129 (12%)<br>1,133 (3%) | CA1PA1 D09BD16D19 CP/CM-A/EPDC | 43,142 | 21,419 (50%) | 21,723 (50%) | 0.20 | CC | 6 | Bifurcating-Trajectories | 1. CP1→CP2→CM1→CM2<br>2. CP1→CP2→EPDC |
| Totals | 62,488 | | 62,488 | 31,407 (50%) | 31,081 (50%) | | | | | |

*Tradeseq [14] parameters: number of knots (k) determined using ‘evaluateK (k = 3:10, nGenes = 200).

Abbreviations: PP, pluripotent; CP, Cardiac Progenitors; CM, cardiomyocyte; EPDC, Epicardium-derived cells; Res, resolution; CC, cell cycle

### Table S12. Lamin A/C Western Blot Quantification Data and Statistical Analyses [33].

#### A. Technical Replicate (TR) Blots: Raw Band Volumes and Normalized Ratio (NR)

| | Raw Band Volume (BV) | | | | | | | | |
|---|---|---|---|---|---|---|---|---|---|
| Cell Type | *Fibroblast* | *Day 19 Differentiated Cells* | | | | | | | |
| Sample ID | CA1 | CA1 | PA1 | CA3 | PA3 | U2 | PA2 | CA1 | |
| Blot 1 | | | | | | | | | |
| Lamin A | 9048601.8 | 483888.56 | 173706.08 | 516436.6 | 449955.8 | 278777 | 153263.4 | | |
| Lamin C | 8877858.2 | 2529873 | 746270.23 | 2318767 | 1937147 | 2856458 | 1054945.5 | | |
| B-Actin | 13319695 | 3442580 | 7558977 | 7507949 | 6711578 | 8815425 | 7482798.55 | | |
| Blot 2 | | | | | | | | | |
| Lamin A | 439561.51 | 88750 | 33436 | 85352.5 | 60939 | 124498.31 | 30014.9 | 119215.5 | |
| Lamin C | 491520.49 | 343447 | 98663.63 | 242385.43 | 197033.5 | 249367.69 | 127044.24 | 250459 | |
| B-Actin | 792273 | 741697 | 684119 | 560653.5 | 538305.5 | 744770 | 717710 | 361561 | |
| Blot 3 | | | | | | | | | |
| Lamin A | 606177 | 116682 | 44697 | 87484 | 65595 | 155078 | 36141 | | |
| Lamin C | 426928 | 137338.5 | 112898 | 230149 | 168734 | 305713 | 161105 | | |
| B-Actin | 585176.5 | 127469.3 | 386324.5 | 410290.5 | 246465.4 | 585631.5 | 422438 | | |
| | Normalized Ratio (NR) = Lamin BV / B-Actin BV | | | | | | | | |
| Cell Type | *Fibroblast* | *Day 19 Differentiated Cells* | | | | | | | |
| Sample ID | CA1 | CA1 (#1)* | PA1 | CA3 | PA3 | U2 | PA2 | CA1 (#2)* | |
| Blot 1 | | | | | | | | | |
| Lamin A/B-Actin | 0.67934 | 0.1405599 | 0.0229801 | 0.068785 | 0.067042 | 0.031624 | 0.0204821 | | |
| Lamin C/B-Actin | 0.6665211 | 0.734877 | 0.0987264 | 0.308842 | 0.288628 | 0.324029 | 0.14098275 | | |
| Lamin A+C/B-Actin | 1.3458611 | 0.8754369 | 0.1217065 | 0.377627 | 0.355669 | 0.355653 | 0.16146484 | | |
| Blot 2 | | | | | | | | | Mean CA1* |
| Lamin A/B-Actin | 0.224691 | 0.119658027 | 0.048874538 | 0.152237523 | 0.113205234 | 0.167163433 | 0.041820373 | 0.329724 | 0.224691 |
| Lamin C/B-Actin | 0.57788571 | 0.463055668 | 0.144219982 | 0.432326615 | 0.366025426 | 0.3348251 | 0.177013334 | 0.692716 | 0.57788571 |
| Lamin A+C/B-Actin | 0.802576946 | 0.582713696 | 0.19309452 | 0.584564138 | 0.47923066 | 0.501988533 | 0.218833707 | 1.02244 | 0.802576946 |
| Blot 3 | | | | | | | | | |
| Lamin A/B-Actin | 1.035887 | 0.915373 | 0.115698 | 0.213225 | 0.266143 | 0.264805 | 0.085553 | | |
| Lamin C/B-Actin | 0.729571 | 1.077424 | 0.292236 | 0.560942 | 0.684615 | 0.522023 | 0.38137 | | |
| Lamin A+C/B-Actin | 1.765459 | 1.992797 | 0.407934 | 0.774166 | 0.950758 | 0.786828 | 0.466923 | | |

*Mean CA1 NR calculated using CA1 (#1) NR and CA1 (#2) NR for Blot 2

### B. Patient vs. Control Samples: Mean Normalized Ratio (NR) of Biological Replicates

| | NR Lamin A | | | | | |
|---|---|---|---|---|---|---|
| | *Day 19 Differentiated Cells* | | | | | |
| | Control | | | Patient | | |
| **Technical Replicate (TR) / Sample ID** | **CA1** | **U2** | **CA3** | **PA1** | **PA2** | **PA3** |
| **Blot 1** | 0.1405599 | 0.031624 | 0.068785 | 0.0229801 | 0.0204821 | 0.067042 |
| **Blot 2** | 0.224691* | 0.167163433 | 0.152237523 | 0.048874538 | 0.041820373 | 0.113205234 |
| **Blot 3** | 0.915373205 | 0.264805 | 0.213225 | 0.115698 | 0.085553 | 0.266143 |
| **Mean NR of TR** | 0.426874767 | 0.15453065 | 0.14474912 | 0.062517572 | 0.049285284 | 0.148796616 |
| **SD of TR** | 0.42513829 | 0.072510204 | 0.117102655 | 0.047840931 | 0.104213127 | 0.033171701 |
| **Mean NR ± SD of BR (n=3)** | 0.242051512 ± 0.160136336 | | | 0.08686649 ± 0.054039603 | | |
| | NR Lamin C | | | | | |
| | *Day 19 Differentiated Cells* | | | | | |
| | Control | | | Patient | | |
| **Technical Replicate (TR) / Sample ID** | **CA1** | **U2** | **CA3** | **PA1** | **PA2** | **PA3** |
| **Blot 1** | 0.73487704 | 0.324029471 | 0.308841661 | 0.098726353 | 0.140982748 | 0.288627604 |
| **Blot 2** | 0.57788571* | 0.3348251 | 0.432326615 | 0.144219982 | 0.177013334 | 0.366025426 |
| **Blot 3** | 1.077423964 | 0.522022808 | 0.560941577 | 0.29223619 | 0.381369574 | 0.177013334 |
| **Mean NR of TR** | 0.796728905 | 0.393625793 | 0.434036618 | 0.178394175 | 0.446422836 | 0.233121885 |
| **SD of TR** | 0.255448344 | 0.111326014 | 0.126058657 | 0.101180141 | 0.209879505 | 0.129644065 |
| **Mean NR ± SD of BR (n=3)** | 0.541463772 ± 0.221987555 | | | 0.285979632 ± 0.141616725 | | |
| | NR Lamin A+C | | | | | |
| | *Day 19 Differentiated Cells* | | | | | |
| | Control | | | Patient | | |
| **Technical Replicate (TR) / Sample ID** | **CA1** | **U2** | **CA3** | **PA1** | **PA2** | **PA3** |
| **Blot 1** | 0.875436899 | 0.355653244 | 0.377626966 | 0.121706457 | 0.161464844 | 0.35566934 |
| **Blot 2** | 0.802576946* | 0.501988533 | 0.584564138 | 0.19309452 | 0.218833707 | 0.47923066 |
| **Blot 3** | 1.992797169 | 0.786827553 | 0.774166109 | 0.407934263 | 0.466922957 | 0.950758354 |
| **Mean NR of TR** | 1.223603671 | 0.548156443 | 0.578785738 | 0.240911747 | 0.282407169 | 0.595219451 |
| **SD of TR** | 0.667136508 | 0.219263376 | 0.198332714 | 0.148984746 | 0.162349479 | 0.314042646 |
| **Mean NR ± SD of BR (n=3)** | 0.783515284 ± 0.381435289 | | | 0.372846122 ± 0.193695355 | | |

*NR value = Mean CA1 NR for Blot 2
Abbreviations: SD, Standard Deviation

### C. Pairwise Comparisons: Relative Normalized Ratio (RNR) and Mean Fold Change (FC)

| | | | | |
|---|---|---|---|---|
| | ***CA1 vs PA1*** | | | |
| | **NR Lamin A+C** | | **RNR Lamin A+C** | |
| **Technical Replicate / Sample ID** | **CA1** | **PA1** | **CA1/CA1** | **PA1/CA1** |
| **Blot 1** | 0.875436899 | 0.121706457 | 1 | 0.139023677 |
| **Blot 2** | 0.802576946 | 0.19309452 | 1 | 0.240593156 |
| **Blot 3** | 1.992797169 | 0.407934263 | 1 | 0.204704357 |
| | | | **Mean FC of TR** | 0.19477373 |
| | | | **SD of TR** | 0.051507793 |
| | | | **% Change** | -80.52262701 |
| | | | **CV (%)** | 26.44493842 |
| | | | **% Change/CV** | -3.044916412 |
| | ***U2 vs PA2*** | | | |
| | **NR Lamin A+C** | | **RNR Lamin A+C** | |
| **Technical Replicate / Sample ID** | **U2** | **PA2** | **U2/U2** | **PA2/U2** |
| **Blot 1** | 0.355653244 | 0.161464844 | 1 | 0.453995139 |
| **Blot 2** | 0.501988533 | 0.218833707 | 1 | 0.435933677 |
| **Blot 3** | 0.786827553 | 0.466922957 | 1 | 0.593424766 |
| | | | **Mean FC of TR** | 0.494451194 |
| | | | **SD of TR** | 0.086188051 |
| | | | **% Change** | -50.55488059 |
| | | | **CV (%)** | 17.43105324 |
| | | | **% Change/CV** | -2.900276873 |
| | ***CA3 vs PA3*** | | | |
| | **NR Lamin A+C** | | **RNR Lamin A+C** | |
| **Technical Replicate / Sample ID** | **CA3** | **PA3** | **CA3/CA3** | **PA3/CA3** |
| **Blot 1** | 0.377626966 | 0.35566934 | 1 | 0.941853661 |
| **Blot 2** | 0.584564138 | 0.47923066 | 1 | 0.819808518 |
| **Blot 3** | 0.774166109 | 0.950758354 | 1 | 1.228106401 |
| | | | **Mean FC of TR** | 0.996589527 |
| | | | **SD of TR** | 0.20958006 |
| | | | **% Change** | -0.341047311 |
| | | | **CV (%)** | 21.0297273 |
| | | | **% Change/CV** | -0.016217391 |

Abbreviations: CV, coefficient of variation
